# Identification of potent and selective *N*-myristoyltransferase inhibitors of *Plasmodium vivax* liver stage hypnozoites and schizonts

**DOI:** 10.1101/2023.01.27.525941

**Authors:** Diego Rodríguez-Hernández, Kamalakannan Vijayan, Rachael Zigweid, Michael K. Fenwick, Banumathi Sankaran, Wanlapa Roobsoong, Jetsumon Sattabongkot, Elizabeth K.K. Glennon, Peter J. Myler, Per Sunnerhagen, Bart L. Staker, Alexis Kaushansky, Morten Grøtli

## Abstract

New drugs targeting multiple stages of the malaria-causing parasite, *Plasmodium*, are needed to reduce and eliminate malaria worldwide. *N*-Myristoyltransferase (NMT) is an essential eukaryotic enzyme, and a validated chemically tractable drug target for malaria. Previous efforts have failed to target NMT owing to the low selectivity for the *Plasmodium* enzyme compared with human NMTs. Herein, we applied a structure-guided approach using previously reported NMT inhibitors as scaffolds to develop a new generation of *Plasmodium vivax* NMT (*Pv*NMT) targeting compounds. We report a series of compounds with IC_50_ values in the nM range and an order of magnitude improved selectivity to *Plasmodium* NMT over human NMT (*Hs*NMT). X-ray co-crystallization of *Pv*NMT with a representative lead compound, **12b**, supported the prevailing hypothesis that a conformational difference in a key tyrosine residue of *Pv*NMT and *Hs*NMT drives the selectivity between these enzymes. The compounds were triaged based on their selectivity for *Pv*NMT. They significantly decreased *P. falciparum* blood-stage parasite load, with IC_50_ values ranging from 0.36 μM to 1.25 μM. The compounds exhibited a dose-dependent inhibition of *P. vivax* liver stage schizont and hypnozoite infection, consistently, in three different *P. vivax* isolates with IC_50_ values ranging from 2.2 μM to 6 μM and from 1.2 μM to 12 μM. Our data provide evidence that NMT inhibitors could be multistage antimalarials, targeting both dormant and developing liver stage parasites, which is essential for malaria elimination.

**One Sentence Summary:** Potent and selective *N*-myristoyltransferase inhibitors of *Plasmodium vivax* hypnozoites and schizonts were synthesized and tested.

## INTRODUCTION

Malaria is a mosquito-borne disease caused by a parasitic infection. In 2020, there were approximately 241 million patients affected by malaria worldwide. Of those, 627,000 people died (*1*). There are five different species of parasites that cause malaria in humans, among which *Plasmodium falciparum* and *Plasmodium vivax* pose the greatest threat. In sub-Saharan Africa most cases of malaria and mortality are caused by *P. falciparum*, while *P. vivax* is globally ubiquitous and causes persistent infections (*2*). *P. vivax* is highly prevalent in the Americas and Southeast Asia, accounting for 75% and 47% of all malaria cases in these regions, respectively (*3*). Antimalarials currently available in the clinic include artemisinin-based combination therapy (ACTs), chloroquine phosphate, sulfadoxine/pyrimethamine, mefloquine, primaquine phosphate, halofantrine, and quinine (*4*). However, the emergence and persistence of resistance to almost all these drugs, and the threat of the continued robust spread of drug tolerance and resistance, necessitates the urgent development of new antimalarials. Additionally, even without the development of resistance, many of the current drugs do not successfully target the *P. vivax* dormant form, called hypnozoites, which is the source of all relapsing infections. Transmission from the *Anopheles* mosquito host to the human host occurs when parasites are deposited in the dermis. The parasites then travel through the skin and ultimately enter a blood vessel. The blood stream then carries the parasites to the liver, where they cross the sinusoidal endothelium and invade a single hepatocyte. For parasites of most *Plasmodium* species, liver stage development takes 2-10 days, during which time the parasites rapidly divide within the host hepatocyte. After liver replication, called schizogony, the parasites exit the liver, invade erythrocytes, and initiate the asexual replication cycle. This cycle is the source of the morbidity and mortality associated with malaria disease. When blood-stage parasites differentiate into sexual gametocyte forms productive human-to-mosquito transmission occurs.

The life-cycles of *P. vivax* and *P. ovale* are slightly altered and include a dormant liver stage form called the hypnozoite, which represents a significant hurdle for malaria eradication efforts. Hypnozoites remain in the liver for weeks, months, or even years and later reactivate, leading to relapse and symptomatic blood-stage infection. Between 20% and 100% of *P. vivax* human cases result from hypnozoite relapse, depending on the location and intensity of transmission (*5, 6*). The current pharmacological options for the elimination of hypnozoites are limited to primaquine and tafenoquine, and the use of these drugs is hampered by their severe toxicity in individuals with glucose-6-phosphate dehydrogenase (G6PD) deficiency. Targeting regulatory proteins within the parasites that are essential for multiple stages of the complex life cycle, including the hypnozoite stage, might be the most robust approach to eliminating malaria.

Multiple antimalarials and combination therapies that can be used as single-dose regimens are being developed pre-clinically and clinically. Ambitious efforts to eliminate malaria, such as those by the Bill and Melinda Gates Foundation and the Medicines for Malaria Venture (MMV), include approaches that could target multiple stages of the parasitic life cycle. Yet, the goal of malaria eradication is far from being achieved, and several knowledge gaps remain in the current pharmacopeia of antimalarial compounds. Antimalarials that are part of several classes are needed to support a robust malaria eradication campaign. Specifically, candidate drugs targeting the asexual blood stage (target candidate profile [TCP]-1), anti-relapse/hypnozoites (TCP-3), liver schizonts (TCP-4), and transmission-blocking (TCP-5 and TCP-6) are essential for reducing malaria cases worldwide (*7, 8*). Accordingly, compounds that can achieve multiple of these objectives would be of particular interest.

*N*-Myristoyltransferase (NMT) is a ubiquitous enzyme in eukaryotes that catalyzes the transfer of myristate from myristoyl-coenzyme A (myrCoA) to the *N*-terminal glycine residue of a nascent polypeptide at the ribosome (*9*). The myristoylation of a protein generally aids in membrane localization and stability by increasing the protein’s hydrophobicity. We and other researchers have previously developed highly active *Plasmodium* NMT inhibitors (NMTis) with moderate selectivity over the human enzyme capable of killing the parasite, some of which have been demonstrated to eliminate *Plasmodium* liver stage and blood-stage parasites (*9*–*14*). Schlott *et al*. discovered **IMP-1002** with an IC_50_ value of 3 nM and up to fourfold selectivity over *Hs*NMT1/2 with an EC_50_ value of 10 nM in a *P. falciparum* growth assay (*13*). In addition, Wright *et al.* reported the activity of **DDD85646**, a potent inhibitor of *P. falciparum* NMT (*Pf*NMT) (*12*), with an IC_50_ value of 40 nM and selectivity <1 over *Hs*NMT1/2 and an EC_50_ value of 69 nM in a *P. falciparum* growth assay (*12*, *13*). Rackham *et al*. also report high-affinity compounds that kill blood- and liver stage parasites; *Pf*NMT with an IC_50_ value of 8 nM, up to fourfold selectivity over *Hs*NMT1/2, *P. falciparum* blood stage EC_50_ value of 302 nM, and *P. berghei* liver stage EC_50_ value of 372 nM (*14*). Pharmacological inhibition of NMT prevents the completion of blood-stage development of *P. falciparum* partly owing to the inhibition of the inner membrane complex formation (*15*). Thus, selective targeting of NMT could aid in the development of a drug that targets multiple life cycle stages.

Although multiple inhibitors of *Plasmodium* NMT have been identified, the selectivity against human NMTs remains a major challenge as *Plasmodium* and human NMTs exhibit a high degree of homology at the active site of the enzyme, where all known inhibitors bind. This challenge has been cited as a reason to deprioritize the targeting of parasitic NMT sites in antimalarial efforts (*16*). In this study, we report the development of *Plasmodium-*specific NMT inhibitors (NMTis) with strong selectivity (~270x) towards *P. vivax* over their human counterpart. Furthermore, we provide a structural basis for such high selectivity through crystal structure elucidation of *Pv*NMT bound to one of our most selective inhibitors. Selected *Pv*NMTis were evaluated in schizont and hypnozoite infections of liver-stage *P. vivax* parasites and blood-stage *P. falciparum* parasites.

## RESULTS

### Design and synthesis of selective *Pv*NMT inhibitors

*Pv*NMT inhibitors **DDD85646** (*12*) and **IMP-1002** (*13*) (Fig. 1A) were used as the starting point for a structure-based approach to develop more potent and selective *Pv*NMTis. Both NMTis occupy different parts of the *P. vivax* NMT binding pocket, with excellent affinity and potency (Fig. 1B). Based on the crystal structures of **IMP-1002** and **DDD85646** bound to *Pv*NMT (Fig. 1B), we designed inhibitors using a part of **IMP-1002** (Fig. 1A, red circle) as a scaffold and fragments of **DDD85646** (Fig. 1A, blue and brown circles) to build hybrid compounds. The molecular hybridization strategy was based on a “headgroup” region linked to core A and a “tail” region linked to core B (Fig. 1C), whose combination formed the biaryl scaffold. Our optimization efforts focused on altering these regions, and different aromatic rings were used to modify the biaryl scaffold (Fig. 1C, cores A and B). In the tail region (Fig. 1C, highlighted in blue), the substituents were designed to interact with Ser319 (12, 14). In the head region (Fig. 1C, highlighted in brown), the substituents were selected to interact ionically with the carboxylate of the C-terminal residue (Leu410), which plays an essential role in myristate transfer and is crucial to the inhibitor’s potency against Plasmodium NMT (12, 13). Fig. 1D shows a particular hybrid compound (12e) that was designed and synthesized to dock into the crystal structure of PvNMT with the target hydrogen bonding and ionic interactions formed with Ser319 and Leu410.

**Fig. 1.**
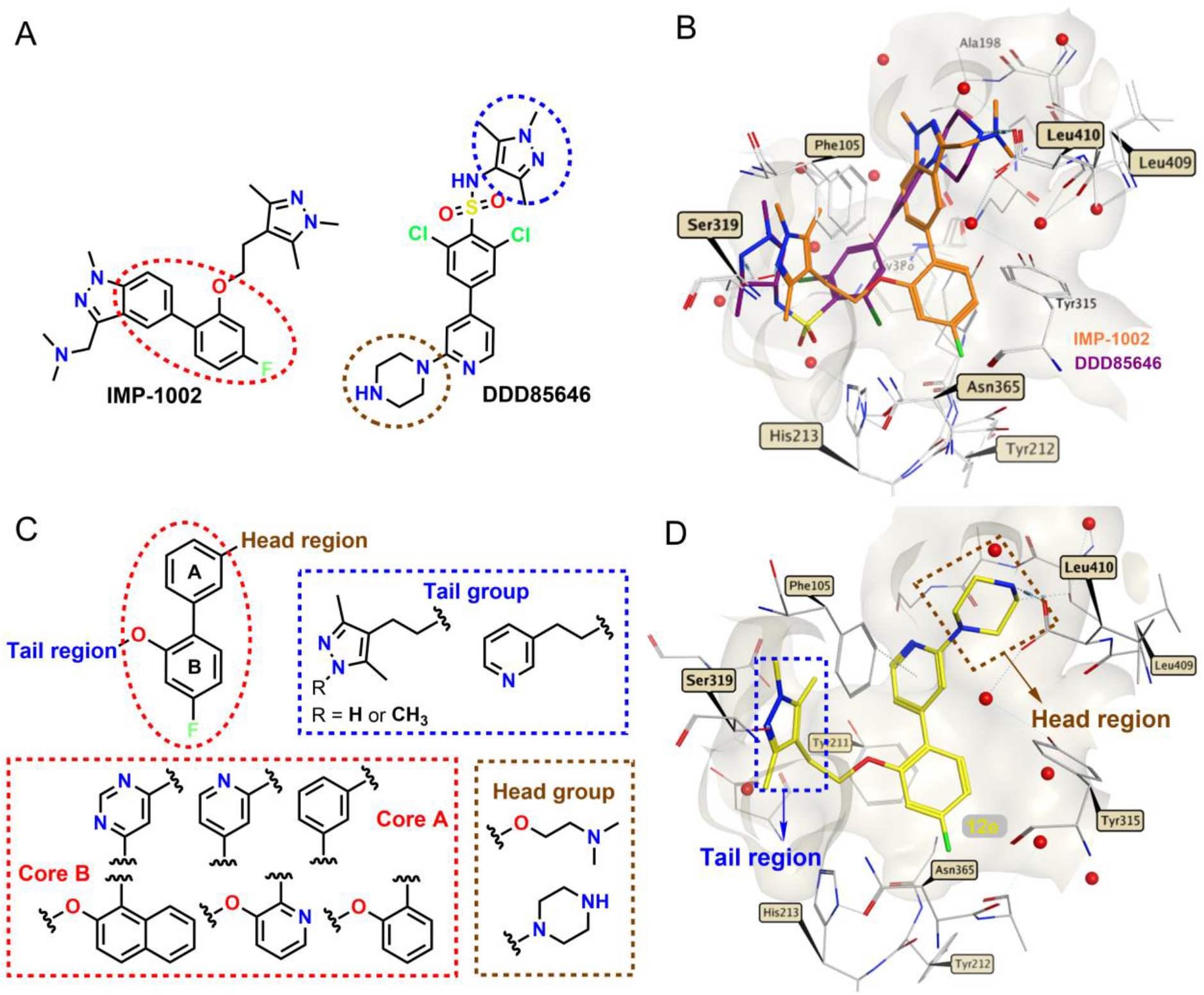
Hybridization approach. (**A**) Chemical structures of **IMP-1002** (*13*) and **DDD85646** (*12*). (**B**) **IMP-1002** and **DDD85646** bound to *Pv*NMT superimposed (PDB codes 6MB1 and 2YND, respectively). (**C**) Design of target molecules: cores A and B (red), tail group (blue), and head group (brown). (**D**) Example of a designed hybrid compound (**12e**) docked into the crystal structure of *Pv*NMT (PDB code 6MB1). Surface representations in front of the binding site are removed for clarity. Interactions involving the atoms of **IMP-1002**, **DDD85646**, and compound **12e** are drawn using dashed lines, and water molecules are represented by red spheres.

### Piperazine-hybrid compounds show high affinity and >145-fold selectivity

Table 1 summarizes the synthetic route used to prepare the hybrid compounds (**12a**-**q** and **16a**-**f**). These structural fragments were prepared using a three-step synthesis strategy (Table 1A), with a condensation reaction as a key step. The synthesis of hybrid compounds **12a**-**q** began with the formation of different blocks of aryl ethers (**3**-**11**) through a sulfonyl transfer (*17*) or the Mitsunobu reaction (*18*) (Table 1B). The intermediate blocks (**3**-**11**) were obtained from commercially available aryl-hydroxyl or pyridine-hydroxyl with pyrazoles **1** and **2** or with commercially available 3-(2-hydroxyethyl)pyridine. From these aryl-ethers blocks, hybrid synthesis was accomplished in two steps (Table 1B). A Suzuki cross-coupling reaction was used to couple cores A and B together (Fig. 1C). This was followed by Boc-deprotection using 4 M HCl in dioxane to furnish compounds **12a**-**q** in sufficient overall yields (60-91%).

**Table 1.** Synthesis and biochemical activity of hybrid compounds bearing a piperazine moiety as a head group.

| Code | core | Y | Z | X | R | Tail | PvNMT IC<sub>50</sub> (nM)* | HsNMT IC<sub>50</sub> (nM)* | SI** |
| --- | --- | --- | --- | --- | --- | --- | --- | --- | --- |
| **12a** |  | N | CH | N | H |  | 36.8 | 5400 | 146.7 |
| **12b** |  | CH | CH | N | H |  | 80.15 | 20780 | 259.2 |
| **12c** |  | N | CH | CH | H |  | 9.48 | 599 | 63.1 |
| **12d** |  | CH | CH | CH | H |  | 48.5 | 7440 | 153.4 |
| **12e** |  | N | CH | CH | F |  | 15.8 | 124 | 7.8 |
| **12f** |  | CH | CH | CH | F |  | 15.9 | 699 | 43.9 |
| **16a** |  | N | N | CH | H |  | 9390 | - | - |
| **16b** |  | N | N | CH | F |  | 168 | 27620 | 164.4 |
| **12g** |  | CH | CH | N | H |  | 104 | 15900 | 152.8 |
| **12h** |  | N | CH | N | H |  | 440.6 | >40000 | 90.9 |
| **12i** |  | CH | CH | CH | H |  | 1540 | - | - |
| **12j** |  | N | CH | CH | H |  | 124 | 27360 | 220.6 |
| **12k** |  | CH | CH | CH | F |  | 3120 | - | - |
| **12l** |  | N | CH | CH | F |  | 23.4 | 1710 | 73 |
| **16c** |  | N | N | CH | H |  | >40000 | - | - |
| **16d** |  | N | N | CH | F |  | 3390 | - | - |
| **12m** |  | CH | CH | N | H |  | 12540 | - | - |
| **12n** |  | N | CH | CH | H |  | 604 | 39730 | 65.7 |
| **12o** |  | N | CH | N | H |  | 3116 | - | - |
| **12p** |  | CH | CH | CH | F |  | 368 | 23080 | 62.7 |
| **12q** |  | N | CH | CH | F |  | 73 | 1632 | 22 |
| **16e** |  | N | N | CH | F |  | 5070 | - | - |
| **16f** |  | N | N | CH | H |  | 14033 | - | - |

\*PvNMT and HsNMT IC<sub>50</sub> values are shown as mean values of two or more determinations. \*\*Enzyme selectivity is calculated as HsNMT IC<sub>50</sub>/PvNMT IC<sub>50</sub> (nM).

NMT activity was indirectly measured through the detection of free CoA by the thiol-reactive probe 7-diethylamino-3-(4′-maleimidylphenyl)-4-methylcoumarin (CPM) (*19*). The inhibition of enzymatic activity of the hybrid compounds **12a**-**q** are summarized in Table 1. Five compounds in this series having either a 1,3,5-trimethylpyrazole (**12a, 12b,** and **12d**) or pyridine (**12g** and **12j**) tail group yielded a selectivity index (SI) of >145, computed as the ratio of IC_50_ values for the *Hs*NMT1 and *Pv*NMT. Moreover, the three compounds bearing a 1,3,5-trimethylpyrazole tail group had IC_50_ values below 100 nM. These results demonstrate that designing potent inhibitors with high selectivity is achievable using the combinatorial design strategy, despite the high degree of active site conservation of *Pv*NMT and *Hs*NMT1.

### Increasing the polarity of the core scaffold with pyrimidine yields mixed results

Compounds **16a**-**f** had a pyrimidine moiety as their core A (Fig. 1C). The introduction of a pyrimidine group increased the polarity of the hybrid compounds. To attach the piperazine head to this core, we reacted commercially available 2,4-dichloropyrimidine with Boc-protected-piperazine to generate compound **13** (Table 1C). Subsequent Suzuki coupling with the appropriate boronic acid yielded biaryl intermediates **14** and **15**. The tail components were introduced via the Mitsunobu reaction, followed by Boc-deprotection using 4M HCl in dioxane, to produce the pyrimidine hybrid compounds (**16a**-**16f)** in good yields.

The introduction of a pyrimidine group in core A, motivated by the hypothesis that this group increases the polarity of the hybrid compounds, had no impact on the affinity over *Pv*NMT (Table 1). However, compound **16b,** having the same groups in the head and tail regions, was more selective over *Hs*NMT (SI = 164) than **12e** or **12f**.

### Dimethylaminoethanol head group shows an increased potency

With the goal of increasing potency against *Pv*NMT, the piperazine head group was replaced with the 2-(dimethylamino)ethanol moiety. This increased the cLogP value by approximately 0.3-0.9 log units for the different compounds presented in Table 1. The protonatable nitrogen from the dimethylamino group was expected to form a favorable ionic interaction with the carboxylate group of the *C*-terminal residue. Table 2A shows the synthetic route used to prepare this series of hybrid compounds (**26a**-**i**). The synthesis began with a Suzuki coupling reaction using the different blocks of aryl ethers (**3**-**11**) with the commercially available 3-hydroxyphenylboronic acid to form the biaryl scaffold (**17**-**25**). Finally, the commercially available 2-(dimethylamino)-ethanol moiety was reacted with a corresponding biaryl framework using the Mitsunobu reaction to produce the desired hybrid compounds (**26a**–**i**) in good overall yields. The in vitro IC50 values of the hybrid compounds 26a-i are summarized in Table 2. The three compounds of this series having a 1,3,5-trimethylpyrazole group in the tail region (26a, 26b, and 26c) exhibited an SI of >115. Although the compounds with the dimethylaminoethanol group had higher cLogP values than the hybrids that had the piperazine group in the head region (12f vs 26c or 12d vs 26b), the affinity for PvNMT was similar to that of compounds with the 1,3,5-trimethylpyrazole group in the tail region with IC50 values below 100 nM (see Tables 1 and 2).

**Table 2.** Synthesis and biochemical activity of hybrid compounds bearing a dimethylamino-ethanol moiety as a head group and hybrid compounds bearing a naphthol moiety in core B of the biaryl scaffold.

| Code | core | X | R | Tail | *Pv*NMT<br>IC<sub>50</sub> (nM)* | *Hs*NMT<br>IC<sub>50</sub> (nM)* | SI** |
| --- | --- | --- | --- | --- | --- | --- | --- |
| 26a |  | N | H |  | 266 | >40000 | 153.8 |
| 26b |  | CH | H |  | 83.2 | 9850 | 118.4 |
| 26c |  | CH | F |  | 29.4 | 4790 | 162.3 |
| 26d |  | N | H |  | 14800 | - | - |
| 26e |  | CH | H |  | 12760 | - | - |
| 26f |  | CH | F |  | 622 | >40000 | 62.3 |
| 26g |  | N | H |  | 19400 | - | - |
| 26h |  | CH | H |  | 1563 | - | - |
| 26i |  | CH | F |  | 242 | 13860 | 57 |

| Code | core | X | Head | Tail | *Pv*NMT<br>IC<sub>50</sub> (nM)* | *Hs*NMT<br>IC<sub>50</sub> (nM)* | SI** |
| --- | --- | --- | --- | --- | --- | --- | --- |
| 30a |  | N |  |  | 89 | 24010 | 269.8 |
| 30b |  | CH |  |  | 19830 | - | - |
| 30c |  | N |  |  | 248 | >40000 | 161 |
| 30d |  | CH |  |  | 7210 | - | - |
| 30e |  | N |  |  | 2880 | - | - |
| 30f |  | CH |  |  | 25400 | - | - |

\**Pv*NMT and *Hs*NMT IC<sub>50</sub> values are shown as mean values of two or more determinations. \*\*Enzyme selectivity calculated as *Hs*NMT IC<sub>50</sub>/*Pv*NMT IC<sub>50</sub> (nM).

### Increasing the lipophilicity of the core scaffold with naphthol improves selectivity

Further synthetic modifications in core B were explored by introducing a polycyclic aromatic hydrocarbon such as naphthol. Through molecular docking of compound 12e (Fig. 1D), a space was observed in the pocket of the binding site, indicating that it was appropriate to introduce another ring in core B. The introduction of the naphthol group increased lipophilicity, improving potency and selectivity. For these hybrids, different groups of heterocycles were preserved, including the tail moiety and the piperazino-pyridine and 2-(phenoxy)-N,N-dimethyl-ethane-1-amine moiety as core A and the head moiety. To synthesize these compounds (30a-f), we employed the same reactions presented in Tables 1A-C and 2A. Table 2B displays the synthetic route used to prepare this series of hybrid compounds (30a-f) in good overall yields.

With the introduction of a naphthol group into core B, the enzymatic activity had no impact when the compounds carried dimethylaminoethanol (**26b** vs. **30b**) in the head region. However, the affinity enhancement imparted by the piperazine group in the head region (**30a** vs **30b**) was accompanied by an improved selectivity (Table 2). Even compound **30a** had less affinity over *Pv*NMT than compound **12c**. However, compound **30a** showed a better degree of selectivity over *Hs*NMT (SI = 269) than did all other hybrid compounds (Tables 1 and 2).

### Crystal structure shows hybrid compound bound in a selective binding mode

We attempted to determine the X-ray co-crystal structures of *Pv*NMT, myrCoA, and compounds with an SI over 200 and nano-molar affinity but were successful in obtaining only high-resolution diffraction from the co-crystals with **12b**, which diffracted to 1.65 Å (Fig. 2A and Table S1). Similar to other *Pv*NMT inhibitors, **12b** binds in a hydrophobic pocket formed by residues from the *N*- and *C*-terminal domains, including the Ab loop (*10-13, 20*). However, the changes in the conformation of *Pv*NMT that take place upon binding **12b**, particularly in residues 217-247 (Fig. 2A), are more significant than ones usually observed in *Pv*NMT-inhibitor complexes.

**Fig. 2.**
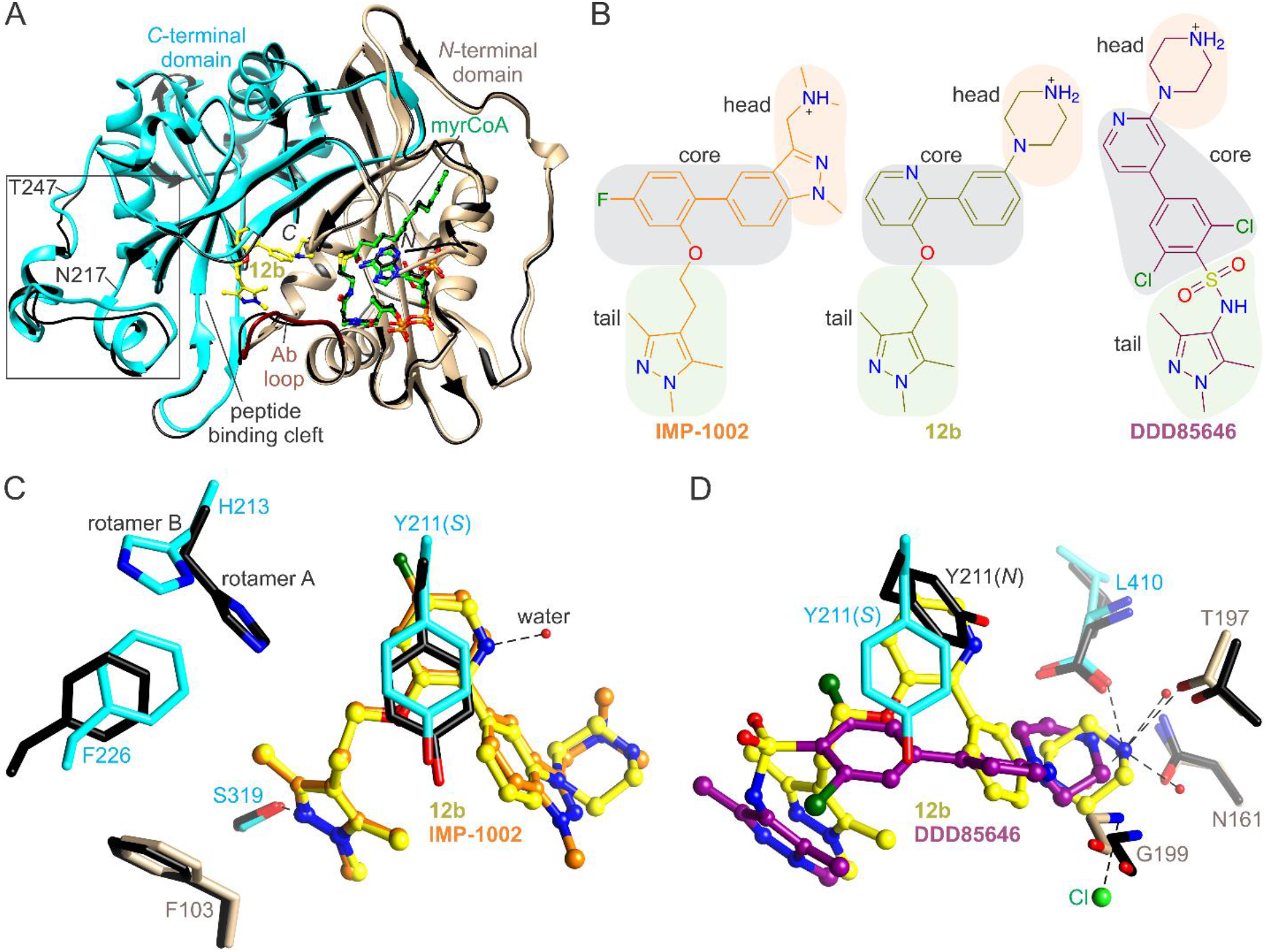
Selective architecture of *Pv*NMT bound to 12b. (**A**) Overall architecture. **12b** binds in the substrate peptide binding cleft at the interface of the NMT *N*- and *C*-terminal domains (tan, residues 1-204, truncated at residue 27, and cyan, residues 205-410, respectively), with the Ab loop (dark red, residues 95-102) in the closed conformation. Comparison with the inhibitor-free state (thin black ribbons, PDB entry 4B10 (*11*)) shows major conformational changes in several regions of *Pv*NMT surrounding **12b** (*e.g.*, residues 217-247). (**B**) **12b** contains the head group of **DDD85646**, the tail group of **IMP-1002**, and a similar core topology as **IMP-1002**. (**C**) **12b** and **IMP-1002** adopt a similar pose but induce distinct binding site architectures. Both stabilize the *S* state of Tyr211, but the pyridine core moiety of **12b** incorporates an additional water molecule, which is accompanied by a change in ligand orientation and conformational differences at Tyr211, His213, and Phe226. (**D**) The piperazine groups of **12b** and **DDD85646** occupy similar binding site locations but interact differently with nearby polar sites of *Pv*NMT via *N4*. In panels **C**and **D**, aligned residues of **IMP-1002**- and **DDD85646-**bound *Pv*NMT are colored black (PDB entries 6MB1 (*13*) and 2YND (*12*), respectively). Hydrogen bonding and salt bridge interactions involving **12b** are drawn using dashed lines.

The **12b** binding site architecture is similar to those of its hybridization precursors, especially **IMP-1002** [(*13*), root mean square deviation of binding site residues of 0.9 Å; Fig. 2B and C]. The phenyl-pyridine core of **12b** stabilizes a highly rotated conformation of Tyr211 that is also observed in the **IMP-1002** bound structure of *Pv*NMT and which was previously proposed to represent a selective (*S*) state Fig. 2C) (*10, 11, 13, 21*). This differs from the **DDD85646**-bound structure, which shows Tyr211 in a nonrotated conformation that corresponds to a nonselective state (*N*) (*12*) (Fig. 2D). The 1,3,5-trimethylpyrazole tail group of **12b** forms a hydrogen bond with Ser319, similar to that of **IMP-1002** and as predicted from the docking calculations performed using **12e**. In addition, the piperazine group of **12b** binds in a similar location as that of **DDD85646**. However, *N4* makes weaker electrostatic interactions with the *C*-terminal carboxylate group conformational change in residues 217-247 is associated with a narrowing of the substrate binding cleft at pocket 8, with peptide excluding distances separating Phe226 and His213 (Figs. 2A and S1) (separation distance of 4.0) and is instead positioned more closely to Thr197, forming a hydrogen bond with the side chain hydroxyl group.

The gain in selectivity achieved via swapping the phenyl and pyridine rings within the core groups of 12b and 12c does not involve changes in polar contacts with PvNMT. Instead, the nitrogen of the pyridine ring of 12b forms a hydrogen bond with an additional water molecule accommodated in this region of the binding site relative to that of IMP-1002, which like 12c, replaces the pyridine of 12b with a phenyl group. This difference, together with the misalignment of the piperazine with Leu410, is associated with an orientational change in the inhibitor and displacement of Tyr211 (Fig. 2C). In addition, the phenyl group, together with the piperazine group, form part of a proximal chloride binding site that would be expected to be replaced with water in the binding site of 12c. Thus, the increased selectivity of 12b over IMP-1002 and 12c is predicted to be related to a difference in the water structure surrounding the inhibitors and the accompanying orientational change in the head and core moieties. The structure also shows that 12b induces a displacement in the side chain of Phe226 to a position that sterically favors rotamer B of His213 (13), whereas IMP-1002 binding favors rotamer A (Fig. 2C). Remarkably, the

### NMTis show minimal toxicity in human hepatoma cells and are active against blood-stage parasites

A major concern in developing *Plasmodium* NMT inhibitors is the potential toxicity in host cells due to cross-reactivity with human NMTs, given the structural overlap between the active site of *Plasmodium* and human NMTs (*9*). Therefore, we assessed cytotoxicity in the human hepatoma HepG2 cell line. We required a *Pv*NMT to *Hs*NMT selectivity index > 20 for further screening. HepG2 cells were exposed to the compounds at concentrations ranging from 1 μM to 20 μM for 48 hours and toxicity was assessed via live-dead staining. Staurosporine, a promiscuous kinase inhibitor and known inducer of cell death in HepG2 cells, was used as a positive control; as expected, it reduced the cell viability at 10-20 μM. None of the NMT-targeting compounds induced >30% cell death in the HepG2 cells, even at their highest concentration of 20 μM (Fig. 3A and Table 3).

**Fig. 3.**
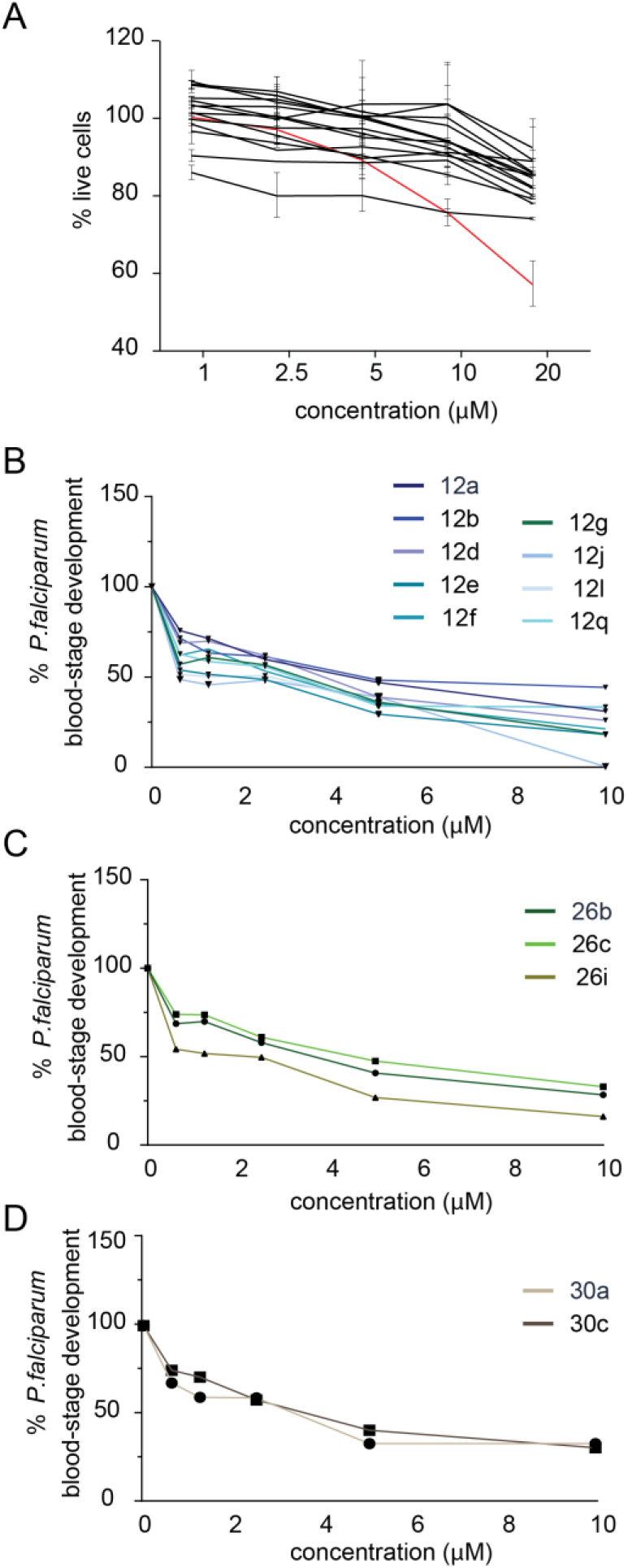
NMT inhibitors show minimal toxicity in human hepatoma cells and reduce *Plasmodium falciparum* blood-stage development. (**A**) Percentage of live HepG2 cells after 48-hour treatment with each NMT inhibitor (shown in black) compared with that of controls treated with diluent. The staurosporine-positive control (shown in red). (**B**-**D**) Percentage change in *P. falciparum* blood-stage development as measured by parasite DNA level fluorescence assay after 72-hour treatment with NMT inhibitors from each of the three prioritized compound families. Data are normalized to untreated controls set at 100%.

**Table 3.**
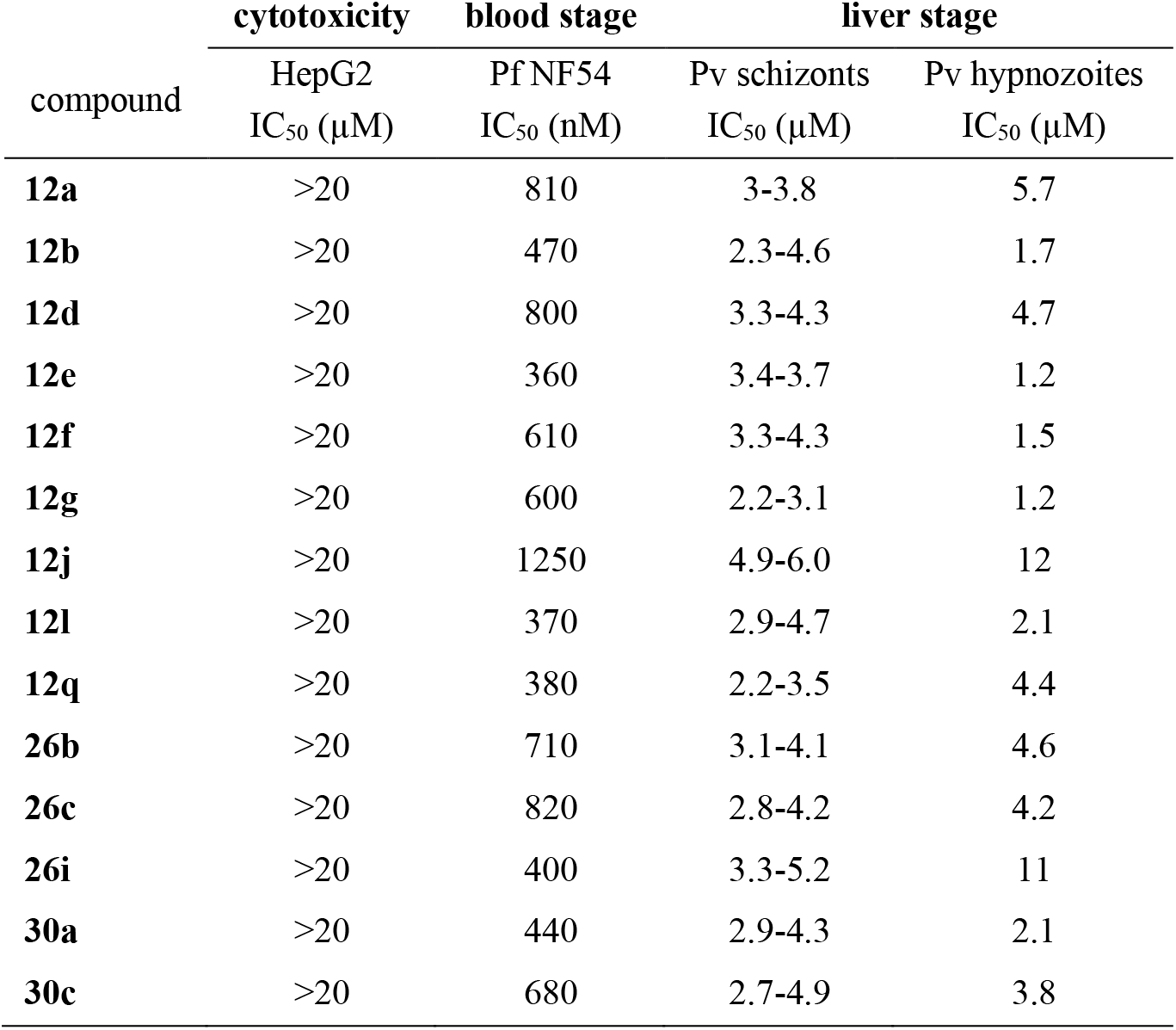
IC_50_ values of NMT inhibitor compounds for cytotoxicity and effect on *Plasmodium* blood and liver stage infection.

To assess whether any of the *Pv*NMTis developed here could qualify as TCP-1 candidates, we assessed the effect of each compound on blood-stage *P. falciparum* parasites, as there is currently no well-established *in vitro* platform for screening *P. vivax* blood-stage parasites. Synchronized *P. falciparum* NF54 ring stage parasites were treated for 72 h with each compound at various concentrations, ranging from 0.625 μM to 10 μM, and parasite DNA replication was measured using a fluorescent DNA binding dye (SYBR Green). The most active compounds against blood stage parasites had the pyridine moiety in core A and the *p*-fluorophenyl group in core B in the biaryl scaffold (**12e**, **12l,** and **12i**) (Fig. 3B-D). The relative IC_50_ values for each compound varied from 360 nM to 1.25 μM (Table 3).

### NMTis are active against *P. vivax* liver stage schizonts

We next evaluated the candidate compounds against liver stage *P. vivax* (Fig. 4). *P*. *vivax* sporozoites were obtained from three independent patient isolates and cultured in primary human hepatocytes from a single donor lot using a 384-well microculture system as described previously (*22*). Cells were infected and treated with NMTis at 20 μM beginning day 5 post-infection until day 8, when the infection rates were quantified by fluorescent microscopy. We defined schizonts as any liver stage parasites that exhibited circumferential *P. vivax* <u>U</u>pregulated in <u>I</u>nfectious <u>S</u>porozoites – 4 (*Pv*UIS4) staining, had multiple nuclear masses, and were ≥10 μm in diameter (Fig. 4A). We observed robust schizont infections in each of the three independent patient isolates (Fig. 4B) and found that all the tested compounds reduced the parasite load by at least 90% when administered at 20 μM (Fig. 4B, Table 3). Accordingly, we applied dose-response curves of all the tested NMTis on parasite isolates 2 and 3 over concentrations up to 20 μM (Fig. 4C and D). We observed similar kinetics for all inhibitors, with a gradual decrease in parasite load upon increasing the drug concentration. The IC_50_ values were calculated for each compound and ranged from 2.2 to 6 μM (Table 3). The most active compounds in this stage (**12b** and **12g**) shared the same scaffold and the piperazine moiety in the tail region; both compounds showed an SI of >150 in the biochemical assay (Table 1).

**Fig. 4.**
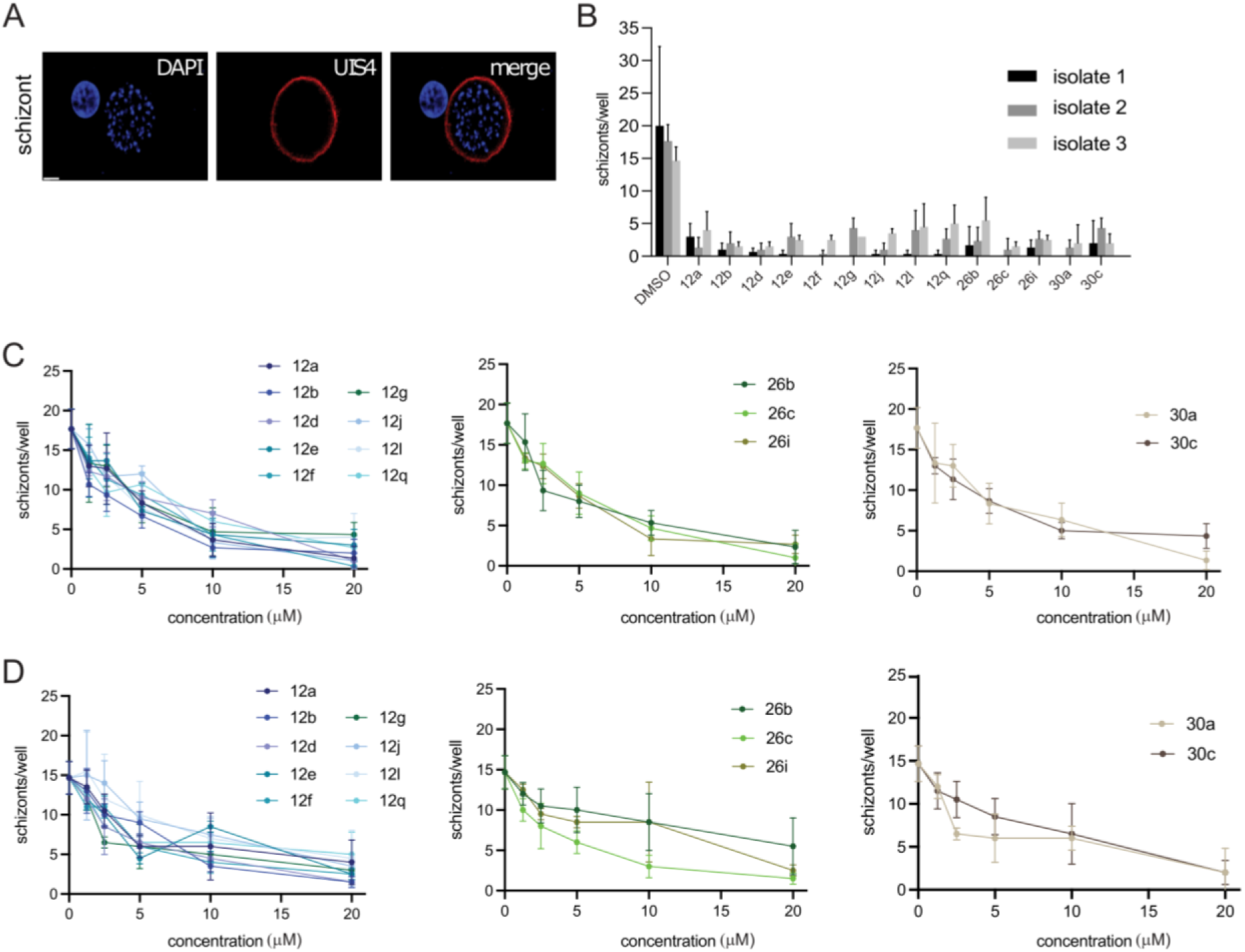
NMT inhibitors reduce *Plasmodium vivax* schizont infection in vitro. (**A**) Representative image of schizont parasite at 8 dpi. Scale bar: 10μm. (**B**) Effect of NMT inhibitor compounds on schizont infection levels at 8 dpi in three independent parasite isolates. All compounds were used at 20 μM, with DMSO as a control. Dose-response curves for each compound, grouped by family, for (**C**) isolates 2 and (**D**) 3. Error bars represent the standard deviation of two to three technical replicates.

### NMTis are active against *P. vivax* liver stage hypnozoites

Finally, we evaluated the NMT-binding compounds against non-developing *P. vivax* hypnozoites. Hypnozoites were defined as parasites having a single nucleus, diameter of <10 mM, and exhibiting prominent staining for *Pv*UIS4 at a point within the parasite periphery (termed the ‘prominence’ (*23*)) (Fig. 5A). Of the three parasite isolates we used to infect primary human hepatocytes, two isolates exhibited robust numbers of *P. vivax* hypnozoites as defined by these criteria (Fig. 5B). We therefore evaluated the impact of all tested compound on *P. vivax* hypnozoites using these two isolates. All tested compounds except **26i** and **30c** exhibited >90% inhibition of hypnozoites at 20 μM (Fig. 5C). Compound **30c** exhibited >75% inhibition at 20μM. The IC_50_ values were calculated for each compound and ranged from 1.2 to 12 μM (Fig. 5D, Table 3). Six compounds showed IC_50_ values < 2.1 μM in this stage, two of which (**12b** and **30a**) had an SI of > 250 in the enzymatic assay, both having the same group in the tail and head regions (Table 1 and 2). Interestingly, the efficacy of NMTis on schizont (Fig. 4) and hypnozoite (Fig. 5) *P. vivax* liver stage forms, as well as efficacy against *P. falciparum* asexual blood stages (Fig. 3), was strongly correlated (Fig. 5E). NMTis also exhibited consistent effects across *P. vivax* parasite isolates. Multiple comparison tests for each compound revealed no significant difference in the effect of NMTis on schizonts between isolates, except for compound **12q** (Table S2). A significant difference in the effect on hypnozoites between isolates was observed for compound **26i** only. Notably, compound **26i** was the only inhibitor that showed a significantly different effect on the two parasite forms, suggesting that *Pv*NMT targeting compounds might form the basis for a drug that is effective against multiple life cycle stages of *Plasmodium*, and across evolved field isolates.

**Fig. 5.**
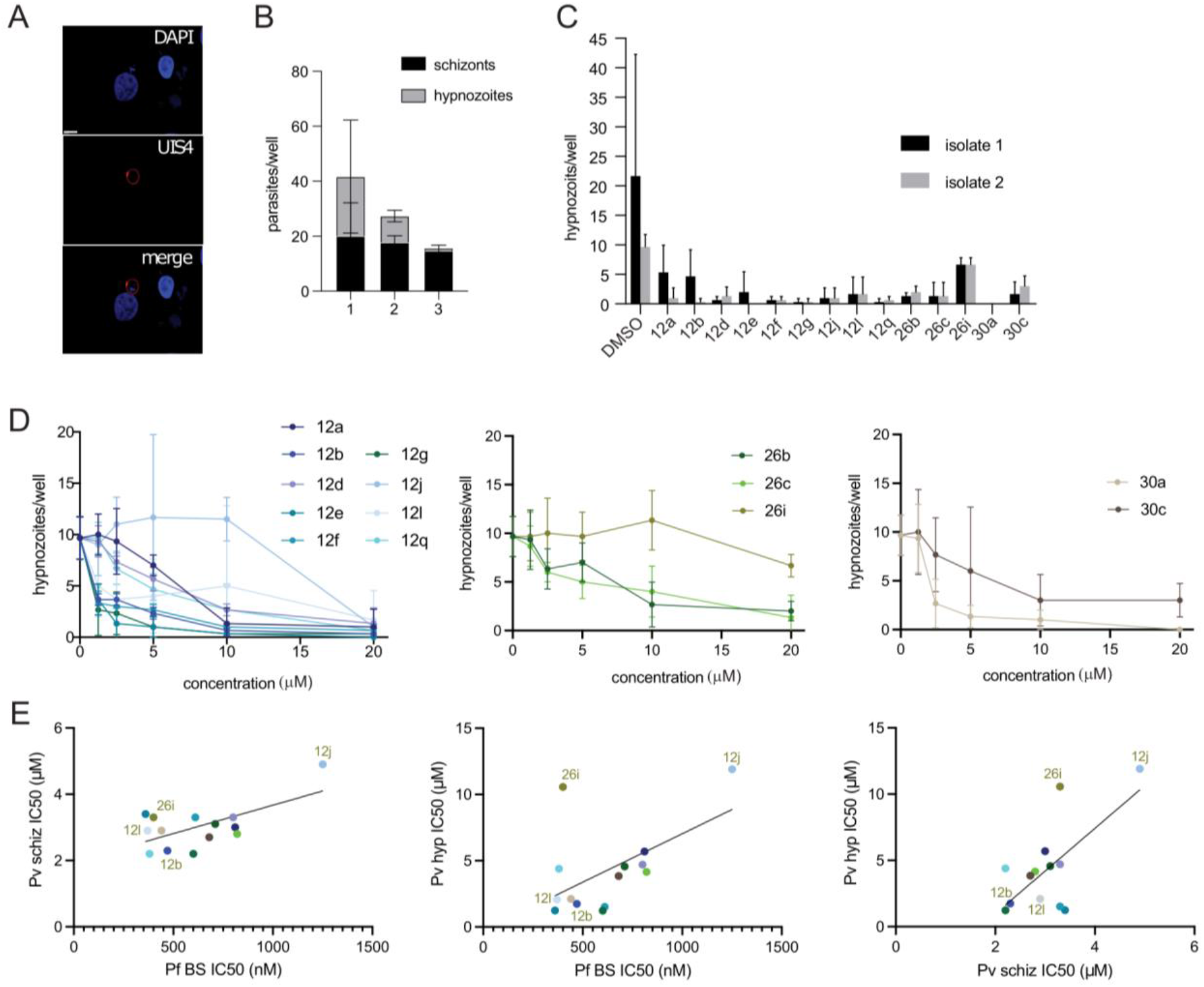
NMT inhibitors reduce *P. vivax* hypnozoite forms *in vitro*. (**A**) Representative image of hypnozoite form 8 dpi. Scale bar: 10μm. (**B**) Schizont and hypnozoite infection levels for each isolate. (**C**) Effect of NMT inhibitor compounds on hypnozoite infection levels 8 dpi for 2 independent parasite isolates. All compounds were administered at 20μM with DMSO as a control. Error bars represent the standard deviation of 2-3 technical replicates. (**D**) Dose-response curves for each compound, grouped by family, for isolate 2. (**E**) Correlations between IC_50_ against *P. falciparum* blood-stage (Pf BS), *P. vivax* schizont (Pv schiz), and *P. vivax* hypnozoite (Pv hyp) infection plotted against each other for each NMTi.

## DISCUSSION

The development of new and potent anti-malarial compounds that target the complete life cycle of *Plasmodium* is needed for eradication. As most of the currently licensed antimalarials target only the erythrocytic stage, expanding our antimalarial arsenal is crucial. Drugs that can effectively target the liver stage have been a particular challenge to develop. The liver stage is a clinically silent and obligatory developmental phase that occurs before parasites can infect erythrocytes and cause malaria symptoms. Targeting the hepatic stage is therefore highly desirable in the context of malaria eradication, not only because its asymptomatic nature makes it ideally suited for prophylactic intervention but also because the liver can serve as a reservoir for *P. vivax* hypnozoites, the dormant parasite forms that cause relapses long after the initial blood infection has been treated.

The enzyme NMT is expressed throughout the *Plasmodium* life cycle and is a promising target for antimalarial drug development. Schlott *et al*. (2021) showed that NMT inhibition disrupts at least 3 vital pathways of the parasite lifecycle: early schizont development, merozoite formation, and merozoite egress (*15*). Despite this promise, the enzyme has been deprioritized as an anti-malarial target due to challenges in identifying parasite-selective inhibitors and a slow mechanism of action (*16*). The impact of NMT inhibition on hypnozoites has not been fully investigated. Here we demonstrate that NMT inhibitors reduce the growth of *P. vivax* schizonts and hypnozoites derived from clinical isolates, validating *Pv*NMT as a target for eliminating dormant forms of the parasite. Combining NMT inhibitors with fast killing blood stage compounds is an attractive strategy for complete parasite elimination (*16*). To develop these selective inhibitors, we utilized a hybrid approach to combine previous inhibitor designs to maximize the affinity and selectivity for the parasite over the human enzyme. We developed a series of inhibitors that exhibit strong selectivity towards *Pv*NMT over *Hs*NMT1/2, combining fragments of **IMP-1002** (*13*) and **DDD85646** (*12*) to synthesize compounds (Fig. 1 and Tables 1 and 2). Functional group substitution was selected to interact with Ser319 at the tail region and the C-terminal residue of Leu410 at the head region (Fig. 1).

We solved a high-resolution cocrystal structure and demonstrated that one of the inhibitors (**12b**) binds within the active site of *Pv*NMT. Specifically, we sought to confirm the mechanism of action, identify the structural determinants associated with the binding of the inhibitors to the active site, and ultimately harness the accumulated structural information and insights gained to optimize pharmacological activity. The observed pose of **12b** in the crystal structure bound to *Pv*NMT validates the design strategy we employed. Compound **12b** adopts the selective pose displayed by **IMP-1002** and the binding of the piperazine moiety near the *C*-terminal carboxylate largely recapitulates the targeted part of the **DDD85646**-binding mode. However, the unique features of the binding-site architecture provide a molecular basis for the approximately 100-fold increase in the selectivity of compound **12b** over **IMP-1002**, which can guide future structure-aided drug development.

Our study not only provides insights into how selective *Pv*NMT inhibitors can be developed but also underscores the potential value of NMTis as antimalarial drugs. While several *P. vivax* targets have been identified, strategies to interfere with the non-developing *P. vivax* hypnozoite form remain a major gap in drug development. Therefore, it is particularly impactful that the compounds we describe here target both developing and dormant liver forms of *P. vivax*. We have validated *P. vivax* NMT as a new target for anti-relapse therapy and developed new NMTis that show promise as multi-stage targeting antimalarials.

### Limitations of this study

In this study, we demonstrate that a new class of *Plasmodium* NMT-targeting compounds are active against multiple stages of the *Plasmodium* life cycle, including asexual blood stages and developing and dormant liver stages of *P. vivax*. However, there are limitations to the assays we use throughout the study. First, all assessments were performed *in vitro*, and as such, we cannot make any statement about the bioavailability or *in viv*o efficacy of these compounds. Additionally, we only evaluate *P. vivax* liver forms for initial rather than relapsing infection. While it is commonly believed that small forms (which we show are reduced in response to the small molecules in this study) are the origin of relapsing infection, we have not directly tested this hypothesis. Finally, our study uses three independent isolates of *P. vivax*, all from Thailand. There is known variability across *Plasmodium* isolates across the globe, so the efficacy of these compounds would need to be assessed for *Plasmodium* isolates from a wider range of geographical origins to understand better how broadly these compounds act against *Plasmodium* parasites.

## MATERIALS AND METHODS

### Study design

This study aimed to identify NMTis with high selectivity for the *P. vivax* enzyme compared with human NMTs and high potency against the different stages of the *P. vivax* parasite. For this purpose, we applied a structure-guided approach using previously reported NMT inhibitors as scaffolds to develop a new generation of *Pv*NMT-targeting compounds. We first measured the NMT activity of these inhibitors using biochemical assays through the detection of free CoA by the thiol-reactive probe 7-diethylamino-3-(4′-maleimidylphenyl)-4-methylcoumarin (CPM) (*19)*. We further evaluated their antiparasitic activity by measuring blood-stage parasite load and inhibition of *P. vivax* liver-stage schizont and hypnozoite infection. X-ray co-crystallization of *Pv*NMT with a representative lead compound, **12b**, was used to rationalize the observed selectivity for *Pv*NMT over *Hs*NMT. Cytotoxicity was assessed in the human hepatoma HepG2 cell line. Synchronized *P. falciparum* NF54 ring-stage parasites were used for measuring blood-stage parasite load. *P. vivax* sporozoites were obtained from three independent patient isolates. Liver-stage infections were cultured in primary human hepatocytes from a single donor lot using a 384-well microculture system (*22*). In vitro experiments were repeated at least twice. A single concentration of each inhibitor was initially used to evaluate the inhibition of *P. vivax* liver stage schizont and hypnozoite infection, followed by the determination of the dose-response relationship.

### Detailed Synthetic Procedure and Characterization of Compounds

All the hybrid compounds were synthesized as described in Supplementary Materials and Methods, and the NMR spectra for the final compounds (**12a-12q**, **16a-16f**, **26a-26i,** and **30a-30f**) are illustrated in Figs. S2-S77.

### Docking of compounds (12e) in the *Pv*NMT (PDB: 6MB1) model binding-pocket

The hybrid compound **12e** was rendered in the form of 2D images using the ChemDraw Professional (Version 19.1.1.21) software package, converted to SDF format, and then prepared for docking using the Molecular Operating Environment (MOE 2019.01) software package. After loading the SDF files, it was processed as follows: the compound was energy-minimized and partial charges added (Amber10 forcefield) using QuickPrep. To prepare the receptor protein, the *Pv*NMT PDB file was loaded into MOE and processed using QuickPrep. The docking simulation was set up by setting the receptor to “receptor+solvent”. The SDF file containing the processed ligands to be docked was loaded. Ligand placement and refinement were performed using the Alpha PMI and rigid receptor methods, with 30 and 3 poses, respectively.

### Cloning, Expression and Purification of *Pv*NMT *P. vivax* and *H. sapiens* NMT enzymes Cloning, Expression and Purification of *Pv*NMT

Cloning, expression and purification were conducted as part of the Seattle Structural Genomics Center for Infectious Disease (SSGCID) *(24, 25*) following protocols described previously (*10, 13, 26, 27*). A region of the *Pv*NMT gene encoding residues 27-410 with a *N*-terminus 6xHis sequence and PreScission cleavage site was cloned into a pET11a expression vector. The *N*-terminal sequence is MGSSHHHHHHSAALEVLFQ/GP-ORF, where cleavage occurs between the glutamine and glycine residues. Plasmid DNA was transformed into chemically competent *E. coli* Rosetta 2 (DE3) pRARE cells. Cells were expression tested and 4-12 liters of culture were grown using auto-induction media (*28*) in the LEX bioreactor for 18-22 h at 18°C. The expression clone was assigned the SSGCID target identifier PlviB.18219.a.FR2.GE44010 and is available at https://www.ssgcid.org/available-materials/ssgcid-proteins/. Protein was purified following a 5-step procedure as previously described (*10, 13*) consisting of a Ni^2+^-affinity chromatography (IMAC), cleavage of the 6xHis-tag and pass through over a second Ni^2+^-affinity chromatography (IMAC) column to remove cleaved tag and protease. The eluted protein was purified further using an anion exchange HiTRAP Q HP 5 mL column. Peak fractions were concentrated to 5 mL and applied to a Superdex 75 10/300 column. The final buffer was composed of 0.3 M NaCl, 20 mM HEPES, 5 %(v/v) glycerol, 1 mM TCEP, pH 7.0. Fractions were analyzed on an SDS-PAGE gel and fractions containing the target protein were concentrated and flash frozen and stored at – 80°C until further use.

### Crystallization and structure determination of *P. vivax* NMT

Purified *Pv*NMT (27-410) concentrated to 8 mg/mL was incubated with 1 mM myrCoA (MedChem101 LLC.) and compound **12b** for 20 minutes at room temperature and set up in 96-well sitting drop crystallization screens JCSG+ HT96 (Molecular Dimensions) and Morpheus HT96 (Molecular Dimensions). Crystals formed within 2 weeks in JCSG+ condition A1 composed of 0.2 M lithium sulfate, 0.1 M sodium acetate, and 50% PEG 400, pH 4.5. Crystals were harvested directly, and flash frozen in liquid nitrogen without cryo-protectant exchange. Frozen crystals were shipped to the Advanced Light Source (ALS), Berkeley National Laboratory as part of the Collaborative Crystallography program of ALS-ENABLE. Data were collected at 100° K on ALS-ENABLE beamlines as described in Supplementary Table S1. Raw X-ray diffraction images are available at the Integrated Resource for Reproducibility in Macromolecular Crystallography at www.proteindiffraction.org (*29*). Data were indexed and integrated with HKL2000 (*30*) and scaled with XSCALE (*31*). The structure was solved with Phaser (*32*) using PDB 6NXG as a search model. The model was refined with iterative rounds of refinement with Phenix (*33*) and manual model building in Coot (*34*). The quality of the structure was checked with Molprobity (*35*). Molecular graphics of this structure were produced using Chimera (*36*).

### NMT Activity Assay

To measure the activity of the purified *Pv*NMT an assay was adapted from Goncalves et al (*10, 13, 19*). The assay buffer was prepared in a 4x stock solution consisting of 9.2 mM potassium phosphate, 69.7 mM sodium phosphate, 2 mM EDTA and 10 % TritonX-100 at pH 7.0. Working stock solutions were made fresh adding DMSO for final concentrations of either 1 % or 5 % DMSO. The *Pv*NMT enzyme was diluted in assay buffer containing 1 % DMSO for a final concentration of 25 nM. Ten μL of the test compound or 10% (v/v) DMSO /water were dispensed into a 96 well plate (Greiner Bio-One) and 50 μL of the enzyme (in assay buffer containing 1 % DMSO) were added for a final concentration of 25 nM per well. The plate was incubated for 30 min at room temperature. The enzymatic reaction was initiated by adding 50 μL of reaction substrate containing 10 μM myrCoA and *Pf*ARF, as well as 8 μM CPM. Fluorescent readings were taken on a Spectra M2 plate reader (Molecular Devices) with excitation at 385 nm and emission at 485 nm. Fluorescent intensity was measured continuously in one-minute intervals for 45 minutes. Background fluorescence and noise were determined by replacing each constituent of the reaction individually with assay buffer containing 1 % DMSO and values were deducted from experimental samples. The enzymatic reactions were set up with varying pHs (6.0-8.5) of the assay buffer to determine optimal reaction conditions with minimal background and off target reactions. In lieu of compound, 10 μL of 10% DMSO/H_2_O was added to each well. Fluorescence signal was obtained continuously for 45 min. A linear reaction rate was observed during the first 30 minutes and used to determine all values. The synthetic peptide (*Pf*ARF) Gly-Leu-Tyr-Val-Ser-Arg-Leu-Phe-Asn-Arg-Leu-Phe-Gln-Lys-Lys-NH2 was purchased from Innopep (San Diego, California). 7-Diethylamino-3-(4'-Maleimidylphenyl)-4-Methylcoumarin (CPM) was purchased from Thermo Scientific Life Technologies (Grand Island, New York) and the co-factor myrCoA was purchased from Med Chem 101 LLC (Plymouth Meeting, Pennsylvania). IC_50_ calculations were calculated using Prism (GraphPad Software, Inc).

### HepG2 cytotoxicity assay

Viability of the HepG2 cell line following exposure to the compounds was determined by a Live/Dead cell assay kit (Invitrogen). Briefly, cells were added to a 96-well plate at a concentration of 5000 cells per well, excluding the exterior wells. Cells were exposed to the compounds at different concentrations ranging from 1 to 20 μM. Cells were washed every 24 h and supplemented with fresh doses of compounds. At 72 h post treatment, 1 μM calcein-AM and 1 μM ethidium homodimer were added to the wells and incubated for 10 min at 37 °C. After washing with PBS, the cells were visualized with fluorescent microscopy (Keyence BZ-X700).

### *P. falciparum* blood stage assay

Sorbitol-synchronized *P. falciparum* NF54 ring-stage parasites were cultured at a parasitemia of 0.5% and hematocrit of 1.5% in a 96-well microtiter plate. Parasites were treated with each NMT inhibitors compounds at 0.625 μM, 1.25 μM, 2.5 μM, 5 μM or 10 μM for 72 h. Direct lysis of the blood cells to release the parasite DNA was performed by adding 100 μl of LBS buffer containing SYBR green I DNA binding dye to each well. Plates were incubated at −20°C overnight for complete lysis and fluorescence was measured at 485 nm (excitation) and 528 nm (emission). ICEstimator regression analysis (http://www.antimalarial-icestimator.net/) was used to obtain relative IC_50_ values for each compound.

### Generation of *P. vivax* sporozoites

*Plasmodium vivax* infected blood was collected from patients attending malaria clinics in Tak and Yala provinces, Thailand, under the approved protocol by the Ethics Committee of the Faculty of Tropical Medicine, Mahidol University (MUTM 2018-016-03). Written Informed consent was obtained from each patient before sample collection. The infected blood was washed once with RPMI1640 incomplete medium before resuspended with AB serum to a final 50% hematocrit and fed to female *Anopheles dirus* through membrane feeding. The engorged mosquitoes were maintained in 10% sugar solution until used. The infected mosquitoes at day 14-21 post feeding were used for harvesting sporozoites.

### *P. vivax* liver stage assay

*Plasmodium vivax* liver stage assays were performed as described previously (*22*). Briefly, we seeded 384-well plates with primary hepatocytes from a single donor lot at a density of 25,000 cells per well. We then infected the hepatocytes with 14,000 freshly hand-dissected sporozoites per well. Cells were exposed to the compounds at indicated concentrations from day 5 post-infection. We then proceeded to feed cultures every day, replacing the compounds until day 8 post-infection. Cells were fixed with 4% PFA, permeabilized with 1% Triton X-100, blocked with 2% BSA, and stained with DAPI and *Pv*UIS4 antibodies to detect schizonts and hypnozoites forms by confocal microscopy. The images were processed and analyzed using IMARIS (Bitplane Inc.) image analysis software.

### Statistical analysis

A 2-way ANOVA with Dunnett’s multiple comparisons test (unpaired) was used to compare the effect of each NMTi on schizonts compared to DMSO controls and to compare the effect of each NMTi on schizonts between parasite isolates. Sidak’s multiple comparisons test (paired) was used to compare the effect of each NMTi on hypnozoites and schizonts within an isolate.

## Acknowledgments

We thank Dr. Noah Sather for the *P. vivax* UIS4 antibody.

## Author contributions

Conceptualization: BLS, AK, MG

Methodology: DRH, KV, MG, WR, JS, BLS, AK

Investigation: DRH, KV, RZ, MF, BS, EKKG, WR, JS, BLS

Visualization: DRH, MF, KV, EKKG

Funding acquisition: PM, BLS, AK, MG, PS

Project administration: BLS, MG, AK

Supervision: BLS, MG, AK

Writing – original draft: DRH, KV, BLS, MF, MG, AK

Writing – review & editing: all authors

## Data and materials availability

Structure coordinates of *Pv*NMT bound to myrCoA and Compound **12b** are available in the Protein Data Bank, accession number: 8FBQ. Raw diffraction images are available at protein diffraction.org using the PDB code 8FBQ. Protein expression plasmids of *Pv*NMT and *Hs*NMT1 and *Hs*NMT2 are available at https://www.ssgcid.org/available-materials/ssgcid-proteins/ through material transfer agreement (MTA).

