## Supplemental Material for "Identification of potent and selective *N*-myristoyltransferase inhibitors of *Plasmodium vivax* liver stage hypnozoites and schizonts"

The PDF file includes:

Materials and Methods

Figs. S1 to S78

Tables S1 to S2

References (12-14, 36-39)

### Materials and Methods

#### General Experimental Information for Synthesis and Compound Characterization

General reagents and solvents for synthesizing compounds were purchased from commercial sources and used as supplied unless otherwise stated.

Purification by flash column chromatography was performed on a Selekt (Biotage, U.K.) automated instrument with Sfär KP-amino D or Sfär silica D cartridges (Biotage, U.K.), mobile phase consist of pentane (solvent A) and ethyl acetate (solvent B) or by reverse phase flash column chromatography performed on an Isolera (Biotage, U.K.) automated instrument with Sfär C18 D cartridges (Biotage, U.K.), mobile phase consisting of water (solvent A) and acetonitrile (solvent B). The standard gradient consisted of *x*% solvent B for one column volume, *x*% to *y*% B for 10 column volumes, and then *y*% B for 2 column volumes. *x* and *y* are defined in the characterization section of the compound the interest.

All NMR spectra ($^{1}$H and $^{13}$C) were recorded on a Varian 400 MHz spectrometer at 25 °C. Samples were dissolved (0.5 mL) in deuterated chloroform, methanol, or dimethylsulfoxide ($CDCl_3$, $CD_3OD$, DMSO-$d_6$). The residual solvent peaks specific to the deuterated solvent was used as an internal reference; $CDCl_3$: 7.26 ppm ($^{1}$H NMR) and 77.20 ppm ($^{13}$C NMR); $CD_3OD$: 3.31 ppm ($^{1}$H NMR) and 49.00 ppm ($^{13}$C NMR); DMSO-$d_6$: 2.50 ppm ($^{1}$H NMR) and 39.52 ppm ($^{13}$C NMR). Data are presented as follows: chemical shift in ppm, multiplicity (br = broad, s = singlet, d = doublet, dd = doublet of doublet, t = triplet, q = quartet, m = multiplet), coupling constants in Hz and integration.

Purity determination of selected compounds was performed in analytical HPLC (Waters 2690 Separations Module; Atlantis® T3, 5 µm column, 4.6 × 250 mm; $H_2O$/ACN (0.1% TFA)). High-resolution mass spectra (HRMS) were recorded on an Agilent 1290 infinity LC system in tandem with an Agilent 6520 Accurate Mass Q-TOF spectrometer.

**Detailed Synthetic Procedure and Characterization of Compounds**

**General Procedure A** – Reduction of Ester to 1° Alcohol

To a solution of lithium aluminum hydride (2.5 equiv.) in anhydrous tetrahydrofuran (15 mL) was cooled to 0 °C and a solution of the selected ester (1 equiv.) in anhydrous tetrahydrofuran (25 mL) was added dropwise. The mixture was stirred a room temperature for 12 h. Then the reaction mixture was cooled to 0 °C, and 2 mL of water was slowly added. The mixture was stirred for 10 min, filtered through celite and rinsed with EtOAc (2x50 mL) before the residue was concentrated under reduced pressure to afford the desired alcohol.

**General Procedure B** – Mesyl transfer

The solution of the alcohol (1 equiv.) and $NEt_3$ (1.5 equiv.), in DCM (10 mL) was cooled to 0°-5°C and methanesulfonyl chloride (1.2 equiv.) was added dropwise. The mixture was stirred at room temperature for 3 h. The reaction was quenched with water (50 mL), and the organic compound was extracted with DCM. The organic layer was dried over $Na_2SO_4$, filtered, and evaporated under reduced pressure to afford the desired phenylmethane sulfonate. The phenylmethane sulfonate (2 equiv.) was dissolved in dry acetonitrile (3 mL), in a microwave vial (2-5 mL) and a solution of the appropriate alcohol (1 equiv.) in solvent (ACN) followed by solid sodium *t*-butoxide (1.2 equiv.) were added. The vial was sealed and then heated under microwave irradiation at 140 °C for 30 min. The reaction mixture was cooled to room temperature and partitioned between EtOAc (20mL) and saturated sodium carbonate solution (10mL). The organic phase was separated, dried over $Na_2SO_4$, filtered, concentrated under reduced pressure, and the crude product was purified by flash column chromatography using a pentane/EtOAc gradient (0-60%) as eluent.

**General Procedure C** – Mitsunobu reaction

A solution of the alcohol (1 equiv.) in toluene (10 mL) was reacted with selected aryl or pyridine-alcohol (1.25 equiv.) and cyanomethyiene tributylphosphorane (1.5 equiv.) at 100°C for 16 h. The reaction mixture was then diluted with ethyl acetate (50 mL) and washed with water (1X25 mL) and brine (2X25 mL). The organic phase separated, dried over $Na_2SO_4$, filtered and concentrated under reduced pressure. The crude product was purified by flash column chromatography eluting with a gradient of pentane/EtOAc or by reverse-phase chromatography eluting with a gradient $H_2O$/ACN.

**General Procedure D** – Suzuki cross-coupling reaction

A solution of the aryl-bromide (1 equiv.), aryl boronic acid pinacol ester (1.25 equiv.) and tetrakis(triphenylphosphine) palladium(0) (0.05 equiv.) in 1,4-dioxane (10 mL) in a microwave vial (10-20 mL) was mixed with a solution of potassium phosphate (2.7 equiv.) in water (3 mL) under $N_2$. The reaction mixture was heated at 100 °C for 1-3 h. The solution was cooled to room temperature and evaporated under reduced pressure. The residue was partitioned between EtOAc (20 mL) and saturated sodium bicarbonate solution (20 mL). The organic phase was dried over $Na_2SO_4$, filtered and concentrated under reduced pressure. The crude product was purified by flash column chromatography eluting with a gradient of pentane/EtOAc or by reverse-phase chromatography eluting with a gradient of $H_2O$/ACN.

**General Procedure E** – Boc-Deprotection

The Boc-protected amine was dissolved in dioxane (1 mL) and treated with a solution of HCl in dioxane (4 M, 2mL). The reaction mixture was stirred at room temperature overnight. All volatiles were removed under reduced pressure, and the product was triturated with ether and DCM, redissolved in water, and freeze-dried to afford the desired compound.

**2-(1,3,5-Trimethyl-1*H*-pyrazol-4-yl)ethan-1-ol (1)**

Ethyl 3-acetyl-4-oxopentanoate (**1a**) and ethyl 2-(1,3,5-trimethyl-1*H*-pyrazol-4-yl)acetate (**1b**) were prepared as described previously (*14*). Compound **1b** was reacted according to General Procedure A. Afforded the title compound as a colorless oil (85.1 %). $^1$H NMR ($CDCl_3$, 400MHz, δ, ppm) 3.68 (3H, s), 3.63 (2H, t, $J$ = 6.8 Hz), 2.59 (2H, t, $J$ = 6.8 Hz), 2.16 (3H, s), 2.15 (3H, s).

**2-(3,5-Dimethyl-1*H*-pyrazol-4-yl)ethan-1-ol (2)**

Ethyl 3-acetyl-4-oxopentanoate (**1a**) and ethyl 2-(3,5-dimethyl-1*H*-pyrazol-4-yl)acetate (**2b**) were prepared as described previously (*14*). Compound **2b** was reacted according to General Procedure A. Afforded the title compound as a colorless oil (80.1 %). $^1$H NMR ($CD_3OD$, 400MHz, δ, ppm) 3.52 (2H, t, $J$ = 7.2 Hz), 2.55 (2H, t, $J$ = 7.2 Hz), 2.14 (6H, s).

**4-(2-(2-Bromophenoxy)ethyl)-1,3,5-trimethyl-1*H*-pyrazole (3)**

Following General Procedure B, 2-bromophenol (250 µL, 2.15 mmol) was reacted with mesyl-chloride (208 µL, 2.69 mmol) and $NEt_3$ (450 µL, 3.23 mmol) to afford 500 mg (92%) of 2-bromophenyl methanesulfonate. $^1$H NMR ($CDCl_3$, 400MHz, δ, ppm) 7.64 (1H, dd, $J$ = 8.0, 1.6 Hz), 7.47 (1H, dd, $J$ = 8.2, 1.6 Hz), 7.37 (1H, ddd, $J$ = 8.2, 7.4, 1.6 Hz), 7.20 (1H, ddd, $J$ = 8.0, 7.4, 1.6 Hz), 2.27 (3H, s). Next, 2-bromophenyl methanesulfonate (488 mg, 1.94 mmol) was reacted with **1** (150 mg, 0.97 mmol) to afford 122 mg (40%) of the title compound as a yellow oil. $^1$H NMR ($CDCl_3$, 400MHz, δ, ppm) 7.52 (1H, dd, *J* = 7.8, 1.6 Hz), 7.22 (1H, ddd, *J* = 8.3, 7.4, 1.6 Hz), 6.85 – 6.78 (2H, m), 4.00 (2H, t, *J* = 7.0 Hz), 3.71 (3H, s), 2.89 (2H, t, *J* = 7.0 Hz), 2.23 (6H, s).

**2-Bromo-3-(2-(1,3,5-trimethyl-1*H*-pyrazol-4-yl)ethoxy)pyridine (4)**

Following General Procedure C, **1** (100 mg, 0.71 mmol) was reacted with 2-bromo-3-hydroxypyridine (155 mg, 0.89 mmol). The crude was purified by flash column chromatography (gradient of pentane/EtOAc) to afford 145 mg (63%) of the title compound as a yellow pale solid. $^1$H NMR ($CDCl_3$, 400MHz, δ, ppm) 7.87 (1H, dd, *J* = 4.6, 1.6 Hz), 7.11 (1H, ddd, *J* = 8.1, 4.6, 0.9 Hz), 7.01 (1H dd, *J* = 8.1, 1.5 Hz), 3.95 (2H, t, *J* = 6.8 Hz), 3.62 (3H, s), 2.84 (2H, t, *J* = 6.6 Hz,), 2.18 (3H, s), 2.16 (3H, s).

**4-(2-(2-Bromo-5-fluorophenoxy)ethyl)-1,3,5-trimethyl-1*H*-pyrazole (5)**

Following General Procedure B, 2-bromo-5-fluorophenol (250 µL, 2.24 mmol) was reacted with mesyl-chloride (217 µL, 2.80 mmol) and $NEt_3$ (470 µL, 3.37 mmol) to afford 530 mg (87%) of 2-bromo-5-fluorophenyl methanesulfonate. $^1$H NMR ($CDCl_3$, 400MHz, δ, ppm) 7.61 (1H, dd, *J* = 8.9, 5.7 Hz), 7.25 (1H, dd, *J* = 8.7, 2.9 Hz), 6.97 (1H, ddd, *J* = 8.9, 7.6, 2.9 Hz), 3.29 (3H, s). Next, 2-bromo-5-fluorophenyl methanesulfonate (500 mg, 1.85 mmol) was reacted with **1** (143 mg, 0.92 mmol) to afford 113 mg (37%) of the title compound as a yellow oil. $^1$H NMR ($CDCl_3$, 400MHz, δ, ppm) 7.49 – 7.42 (1H, m), 6.60 – 6.53 (2H, m), 3.96 (2H, t, *J* = 6.9 Hz), 3.71 (3H, s), 2.89 (2H, t, *J* = 6.9 Hz,), 2.24 (3H, s), 2.23 (3H, s).

**4-(2-(2-Bromophenoxy)ethyl)-3,5-dimethyl-1*H*-pyrazole (6)**

Following General Procedure C, **2** (100 mg, 0.71 mmol) was reacted with 2-bromophenol (103 µL, 0.89 mmol). The crude was purified by flash column chromatography (gradient of pentane/EtOAc) to afford 130 mg (61%) of the title compound as a yellow pale solid. $^1$H NMR ($CDCl_3$, 400MHz, δ, ppm) 7.51 (1H, dd, *J* = 7.8, 1.6 Hz), 7.21 (1H, ddd, *J* = 8.3, 7.4, 1.6 Hz), 6.84 – 6.78 (2H, m), 4.00 (2H, t, *J* = 7.1 Hz), 2.90 (2H, t, *J* = 7.1 Hz), 2.28 (6H, s).

**2-Bromo-3-(2-(3,5-dimethyl-1*H*-pyrazol-4-yl)ethoxy)pyridine (7)**

Following General Procedure C, **2** (100 mg, 0.71 mmol) was reacted with 2-bromo-3-hydroxypyridine (155 mg, 0.89 mmol). The crude was purified by flash column chromatography (gradient of pentane/EtOAc) to afford 140 mg (66%) of the title compound as a yellow pale solid. $^1$H NMR ($CDCl_3$, 400MHz, δ, ppm) 7.96 (1H, dd, *J* = 4.6, 1.6 Hz), 7.18 (1H, dd, *J* = 8.1, 4.6 Hz), 7.06 (1H, dd, *J* = 8.1, 1.6 Hz), 4.03 (2H, t, *J* = 6.8 Hz), 2.93 (2H, t, *J* = 6.8 Hz,), 2.29 (s, 6H).

**4-(2-(2-Bromo-5-fluorophenoxy)ethyl)-3,5-dimethyl-1*H*-pyrazole (8)**

Following General Procedure C, **2** (78 mg, 0.55 mmol) was reacted with 2-bromo-5-fluorophenol (80 µL, 0.71 mmol). The crude was purified by flash column chromatography (gradient of pentane/EtOAc) to afford 120 mg (68%) of the title compound as a yellow pale solid. $^1$H NMR ($CDCl_3$, 400MHz, δ, ppm) 7.44 (1H, dd, *J* = 8.8, 6.2 Hz), 6.60 – 6.52 (2H, m), 3.98 (2H, t, *J* = 6.9 Hz), 2.91 (2H, t, *J* = 6.9 Hz), 2.30 (6H, s).

**3-(2-(2-Bromophenoxy)ethyl)pyridine (9)**

Following General Procedure B, 2-bromophenyl methanesulfonate (500 mg, 1.99 mmol) was reacted with 3-(2-hydroxyethyl)pyridine (112 µL, 0.99 mmol) to afford 125 mg (45%) of the title compound as a yellow solid. $^1$H NMR ($CDCl_3$, 400MHz, δ, ppm) 8.61 (1H, d, *J* = 2.1 Hz), 8.50 (1H, dd, *J* = 4.8, 1.6 Hz), 7.78 – 7.72 (1H, m), 7.52 (1H, dd, *J* = 7.8, 1.6 Hz), 7.27 – 7.24 (2H, m), 7.24 – 7.21 (1H, m), 6.87 – 6.79 (2H, m), 4.22 (2H, t, *J* = 6.3 Hz), 3.15 (2H, t, *J* = 6.3 Hz).

**2-Bromo-3-(2-(pyridin-3-yl)ethoxy)pyridine (10)**

Following General Procedure C, 3-(2-hydroxyethyl)pyridine (112 µL, 0.99 mmol) was reacted with 2-bromo-3-hydroxypyridine (138 mg, 0.79 mmol). The crude was purified by flash column chromatography (gradient of pentane/EtOAc) to afford 118 mg (69%) of the title compound as a yellow pale solid. $^1$H NMR ($CDCl_3$, 400MHz, δ, ppm) 8.61 (1H, d, *J* = 2.3 Hz), 8.52 (1H, dd, *J* = 4.8, 1.6 Hz), 7.98 (1H, dd, *J* = 4.6, 1.6 Hz), 7.76 (1H, dt, *J* = 7.8, 2.0 Hz), 7.27 (2H, dd, *J* = 7.8, 4.8 Hz), 7.18 (1H, ddd, *J* = 8.1, 4.6, 0.4 Hz), 7.08 (1H, dd, *J* = 8.1, 1.5 Hz), 4.22 (2H, t, *J* = 6.3 Hz), 3.18 (2H, t, *J* = 6.3 Hz).

**3-(2-(2-Bromo-5-fluorophenoxy)ethyl)pyridine (11)**

Following General Procedure C, 3-(2-hydroxyethyl)pyridine (68 µL, 0.60 mmol) was reacted with 2-bromo-5-fluorophenol (84 µL, 0.75 mmol). The crude was purified by flash column chromatography (gradient of pentane/EtOAc) to afford 130 mg (72%) of the title compound as a yellow pale solid. $^1$H NMR ($CDCl_3$, 400MHz, δ, ppm) 8.55 (1H, d, *J* = 2.2 Hz), 8.44 (1H, dd, *J* = 4.8, 1.6 Hz), 7.71 – 7.63 (1H, m), 7.37 (1H, ddd, *J* = 8.0, 6.2, 0.7 Hz), 7.18 (1H, ddd, *J* = 7.8, 4.8, 0.7 Hz), 6.55 – 6.45 (2H, m), 4.09 (2H, t, *J* = 6.2 Hz), 3.07 (2H, t, *J* = 6.2 Hz).

**2'-(Piperazin-1-yl)-3-(2-(1,3,5-trimethyl-1*H*-pyrazol-4-yl)ethoxy)-2,4'-bipyridine (12a)**

Following General Procedure D, **4** (35 mg, 0.11 mmol) was reacted with 2-(4-*tert*-butoxycarbonylpiperazin-1-yl)pyridine-4-boronic acid, pinacol ester (51 mg, 0.14 mmol), the crude was purified by reverse phase chromatography (6g C18 column; MeCN in water 0-100%). The resulting residue was reacted according to General

Procedure E. Afforded the title compound as a fluffy colorless solid (7 mg, 16%). $^1$H NMR ($CD_3OD$, 400MHz, δ, ppm) 8.20 (1H dd, *J* = 4.7, 1.3 Hz), 8.13 (1H, dd, *J* = 5.3, 0.7 Hz), 7.57 (1H, dd, *J* = 8.5, 1.3 Hz), 7.41 (1H, dd, *J* = 8.5, 4.7 Hz), 7.12 – 7.11 (1H, m), 6.95 (1H, dd, *J* = 5.3, 1.3 Hz), 4.15 (2H, t, *J* = 6.5 Hz), 3.63 (3H, s), 3.54 – 3.50 (4H, m), 2.97 – 2.93 (4H, m), 2.83 (2H, t, *J* = 6.5 Hz), 2.05 (3H, s), 2.01 (3H, s). $^{13}$C NMR ($CD_3OD$, 100MHz, δ, ppm) 161.35, 154.85, 148.38, 148.12, 147.57, 146.90, 141.92, 139.20, 125.97, 122.31, 115.31, 113.29, 109.38, 69.80, 47.34, 46.31, 35.77, 24.38, 11.61, 9.40. HRMS (ESI), found 393.2403 $C_{22}H_{28}N_6O$, $[M + H]^+$, requires 393.2403.

**1-(3-(3-(2-(1,3,5-Trimethyl-1*H*-pyrazol-4-yl)ethoxy)pyridin-2-yl)phenyl)piperazine (12b)**

Following General Procedure D, **4** (100 mg, 0.32 mmol) was reacted with (3-(4-(*tert*-butoxycarbonyl)piperazin-1-yl)phenyl)boronic acid (123 mg, 0.40 mmol), the crude was purified by reverse phase chromatography (6g C18 column; MeCN in water 0-100%). The resulting residue was reacted according to General Procedure E. Afforded the title compound as a fluffy yellow solid (70 mg, 83%). $^1$H NMR ($CD_3OD$, 400MHz, δ, ppm) 8.40 (1H, dd, *J* = 5.7, 1.1 Hz), 8.33 (1H, dd, *J* = 8.8, 1.0 Hz), 7.99 (1H, dd, *J* = 8.8, 5.7 Hz), 7.51 (1H, t, *J* = 7.9 Hz), 7.38 – 7.32 (2H, m), 7.18 – 7.14 (1H, m), 4.39 (2H, t, *J* = 6.0 Hz), 3.86 (3H, s), 3.56 (4H, dd, *J* = 6.3, 4.1 Hz), 3.41 (4H, dd, *J* = 6.3, 4.1 Hz), 2.99 (2H, t, *J* = 6.0 Hz), 2.15 (3H, s), 2.11 (3H, s). $^{13}$C NMR ($CD_3OD$, 100MHz, δ, ppm) 156.36, 152.02, 146.03, 145.13, 144.75, 134.15, 131.19, 131.16, 130.54, 128.24, 122.85, 120.55, 118.55, 116.47, 70.77, 47.26, 44.63, 35.58, 23.24, 9.59, 9.33. HRMS (ESI), found 392.2454 $C_{23}H_{29}N_5O$, $[M + H]^+$, requires 392.2453.

**1-(4-(2-(2-(1,3,5-Trimethyl-1*H*-pyrazol-4-yl)ethoxy)phenyl)pyridin-2-yl)piperazine (12c)**

Following General Procedure D, **3** (100 mg, 0.32 mmol) was reacted with 2-(4-*tert*-butoxycarbonylpiperazin-1-yl)pyridine-4-boronic acid, pinacol ester (154 mg, 0.40 mmol), the crude was purified by reverse phase chromatography (6g C18 column; MeCN in water 0-100%). The resulting residue was reacted according to General Procedure E. Afforded the title compound as a fluffy pale yellow solid (65 mg, 51%). $^1$H NMR ($CD_3OD$, 400MHz, δ, ppm) 8.02 (1H, dd, *J* = 5.3, 0.6 Hz), 7.27 (1H, ddd, *J* = 8.3, 7.4, 1.8 Hz), 7.22 (1H, dd, *J* = 7.6, 1.8 Hz), 7.00 – 6.97 (1H, m), 6.95 (1H, td, *J* = 7.6, 1.1 Hz), 6.80 (1H, s), 6.66 (1H, dd, *J* = 5.3, 1.1 Hz), 3.97 (2H, t, *J* = 6.5 Hz), 3.57 (3H, s), 3.45 – 3.40 (4H, m), 2.90 – 2.86 (4H, m), 2.69 (2H, t, *J* = 6.5 Hz), 1.98 (3H, s), 1.91 (3H, s). $^{13}$C NMR ($CD_3OD$, 100MHz, δ, ppm) 161.29, 157.12, 150.22, 147.85, 146.79, 139.06, 131.36, 130.90, 130.31, 122.10, 116.19, 113.94, 113.58, 109.72, 69.57, 47.41, 46.33, 35.75, 24.56, 11.65, 9.38. HRMS (ESI), found 392.2451 $C_{23}H_{29}N_5O$, $[M + H]^+$, requires 392.2450.

**1-(2'-(2-(1,3,5-Trimethyl-1*H*-pyrazol-4-yl)ethoxy)-[1,1'-biphenyl]-3-yl)piperazine (12d)**

Following General Procedure D, **3** (100 mg, 0.32 mmol) was reacted with (3-(4-(*tert*-butoxycarbonyl)piperazin-1-yl)phenyl)boronic acid (123 mg, 0.40 mmol), the crude was purified by reverse phase chromatography (6g C18 column; MeCN in water 0-100%). The resulting residue was reacted according to General Procedure E. Afforded the title compound as a fluffy brownish white solid (60 mg, 58%). $^{1}$H NMR ($CD_3OD$, 400MHz, δ, ppm) 7.34 – 7.25 (2H, m), 7.18 (1H, dd, *J* = 7.5, 1.7 Hz), 7.14 – 7.10 (2H, m), 7.05 – 7.02 (1H, m), 6.98 (1H, td, *J* = 7.5, 0.8 Hz), 6.93 (1H, d, *J* = 7.7 Hz), 4.09 (2H, t, *J* = 5.8 Hz), 3.84 (3H, s), 3.54 (4H, dd, *J* = 6.4, 3.4 Hz), 3.45 (4H, dd, *J* = 6.4, 3.3 Hz), 2.86 (2H, t, *J* = 5.8 Hz), 2.09 (3H, s), 2.06 (3H, s). $^{13}$C NMR ($CD_3OD$, 100MHz, δ, ppm) 156.83, 150.29, 146.30, 144.45, 141.88, 132.43, 131.87, 130.10, 130.08, 124.93, 122.35, 119.89, 117.68, 117.11, 114.03, 68.77, 48.68, 44.51, 35.49, 23.68, 9.40, 9.22. HRMS (ESI), found 391.2499 $C_{24}H_{30}N_4O$, $[M + H]^+$, requires 391.2498.

**1-(4-(4-Fluoro-2-(2-(1,3,5-trimethyl-1*H*-pyrazol-4-yl)ethoxy)phenyl)pyridin-2-yl)piperazine (12e)**

Following General Procedure D, **5** (50 mg, 0.15 mmol) was reacted with 2-(4-*tert*-butoxycarbonylpiperazin-1-yl)pyridine-4-boronic acid, pinacol ester (75 mg, 0.19 mmol), the crude was purified by reverse phase chromatography (6g C18 column; MeCN in water 0-100%). The resulting residue was reacted according to General Procedure E. Afforded the title compound as a fluffy pale yellow solid (11 mg, 52%). $^{1}$H NMR ($CD_3OD$, 400MHz, δ, ppm) 8.11 (1H, dd, *J* = 5.2, 0.7 Hz), 7.28 (1H, dd, *J* = 8.5, 6.7 Hz), 6.89 – 6.84 (2H, m), 6.77 – 6.71 (2H, m), 4.05 (2H, t, *J* = 6.4 Hz), 3.79 – 3.73 (4H, m), 3.31 – 3.27 (4H, m), 3.61 (3H, s), 2.76 (2H, t, *J* = 6.3 Hz), 1.97 (3H, s), 1.96 (3H, s). $^{13}$C NMR ($CD_3OD$, 100MHz, δ, ppm) 165.14 (d, *J* = 245 Hz), 160.06, 158.50 (d, *J* = 10.1 Hz), 149.79, 148.24, 146.94, 139.15, 132.49 (d, *J* = 10.1 Hz), 126.16 (d, *J* = 3.3 Hz), 117.36, 113.54, 109.96, 108.25 (d, *J* = 21.5 Hz), 101.70 (d, *J* = 25.90 Hz), 69.89, 44.53, 44.13, 35.76, 24.29, 11.56, 9.38. HRMS (ESI), found 410.2356 $C_{23}H_{28}FN_5O$, $[M + H]^+$, requires 410.2356.

**1-(4'-Fluoro-2'-(2-(1,3,5-trimethyl-1*H*-pyrazol-4-yl)ethoxy)-[1,1'-biphenyl]-3-yl)piperazine (12f)**

Following General Procedure D, **5** (100 mg, 0.30 mmol) was reacted with (3-(4-(*tert*-butoxycarbonyl)piperazin-1-yl)phenyl)boronic acid (121 mg, 0.39 mmol), the crude was purified by reverse phase chromatography (6g C18 column; MeCN in water 0-100%). The resulting residue was reacted according to General Procedure E. Afforded the title compound as a fluffy brownish white solid (60 mg, 58%). $^{1}$H NMR ($CD_3OD$, 400MHz, δ, ppm) 7.31 – 7.26 (1H, m), 7.17 (1H, dd, *J* = 8.4, 6.8 Hz), 7.04 (1H, ddd, *J* = 8.2, 2.5, 0.8 Hz), 7.00 – 6.97 (1H, m), 6.88 – 6.81 (2H, m), 6.72 (1H, td, *J* = 8.3, 2.5 Hz), 4.09 (2H, t, *J* = 5.8 Hz), 3.84 (3H, s), 3.47 (4H, dd, *J* = 6.7, 3.5 Hz), 3.40

(4H, dd, $J = 6.6, 3.6$ Hz), 2.87 (2H, t, $J = 5.8$ Hz), 2.08 (3H, s), 2.05 (3H, s). $^{13}$C NMR ($CD_3OD$, 100MHz, δ, ppm) 164.40 (d, $J = 243.3$), 158.05 (d, $J = 9.9$), 151.43, 146.24, 144.53, 140.81, 132.73 (d, $J = 9.9$), 130.04, 128.74 (d, $J = 3.3$), 123.86, 119.42, 117.45, 116.77, 108.27 (d, $J = 21.2$), 101.69 (d, $J = 25.85$), 69.06, 48.03, 44.76, 35.46, 23.48, 9.37, 9.17. HRMS (ESI), found 429.2407 $C_{24}H_{30}FN_4O$, $[M + H]^+$, requires 429.2404.

**1-(3-(3-(2-(Pyridin-3-yl)ethoxy)pyridin-2-yl)phenyl)piperazine (12g)**

Following General Procedure D, **10** (100 mg, 0.35 mmol) was reacted with (3-(4-(*tert*-butoxycarbonyl)piperazin-1-yl)phenyl)boronic acid (137 mg, 0.44 mmol), the crude was purified by reverse phase chromatography (6g C18 column; MeCN in water 0-100%). The resulting residue was reacted according to General Procedure E. Afforded the title compound as a fluffy yellow solid (70 mg, 83%). $^1$H NMR ($CD_3OD$, 400MHz, δ, ppm) 8.73 (2H, s), 8.41 (2H, d, $J = 7.4$ Hz), 8.37 – 8.32 (1H, m,), 7.98 (1H, dd, $J = 8.7, 5.7$ Hz,), 7.95 – 7.88 (1H, m), 7.51 – 7.45 (1H, m), 7.33 – 7.28 (2H, m), 7.15 – 7.10 (1H, m), 4.63 (2H, t, $J = 6.0$ Hz), 3.54 (4H, dd, $J = 6.3, 4.0$ Hz), 3.44 – 3.37 (6H, m). $^{13}$C NMR ($CD_3OD$, 100MHz, δ, ppm) 154.49, 150.46, 147.37, 143.83, 141.36, 139.58, 133.29, 129.64, 129.49, 128.99, 126.67, 121.56, 119.16, 117.19, 69.05, 45.85, 43.23, 31.48. HRMS (ESI), found 361.2029 $C_{22}H_{24}N_4O$, $[M + H]^+$, requires 361.2028.

**2'-(Piperazin-1-yl)-3-(2-(pyridin-3-yl)ethoxy)-2,4'-bipyridine (12h)**

Following General Procedure D, **10** (88 mg, 0.31 mmol) was reacted with 2-(4-*tert*-butoxycarbonylpiperazin-1-yl)pyridine-4-boronic acid, pinacol ester (153 mg, 0.39 mmol), the crude was purified by reverse phase chromatography (6g C18 column; MeCN in water 0-100%). The resulting residue was reacted according to General Procedure E. Afforded the title compound as a fluffy yellow solid (40 mg, 85%). $^1$H NMR ($CD_3OD$, 400MHz, δ, ppm) 8.81 (1H, s), 8.75 (1H, d, $J = 5.5$ Hz), 8.60 (1H, d, $J = 8.1$ Hz), 8.39 (1H, d, $J = 4.5$ Hz), 8.15 (1H, d, $J = 6.4$ Hz), 8.03 (1H, dd, $J = 8.0, 5.9$ Hz), 7.94 (1H, dd, $J = 8.7, 0.9$ Hz), 7.79 – 7.76 (1H, m), 7.70 (1H, dd, $J = 8.6, 4.9$ Hz), 7.44 (1H, dd, $J = 6.4, 1.2$ Hz), 4.58 (2H, t, $J = 6.2$ Hz), 4.09 – 4.03 (4H, m), 3.51 – 3.46 (4H, m), 3.44 (2H, t, $J = 6.2$ Hz). $^{13}$C NMR ($CD_3OD$, 100MHz, δ, ppm) 155.73, 154.62, 148.91, 142.74, 142.18, 141.06, 141.05, 140.95, 140.62, 139.16, 128.74, 128.51, 125.44, 115.90, 113.73, 69.68, 44.58, 43.76, 33.02. HRMS (ESI), found 362.1986 $C_{21}H_{23}N_5O$, $[M + H]^+$, requires 362.1981.

**1-(2'-(2-(Pyridin-3-yl)ethoxy)-[1,1'-biphenyl]-3-yl)piperazine (12i)**

Following General Procedure D, **9** (100 mg, 0.36 mmol) was reacted with (3-(4-(*tert*-butoxycarbonyl)piperazin-1-yl)phenyl)boronic acid (137 mg, 0.44 mmol), the crude was purified by reverse phase chromatography (6g C18 column; MeCN in water 0-100%). The resulting residue was reacted according to General Procedure E. Afforded the title compound as a fluffy off-white solid (76 mg, 74%). $^1$H NMR ($CD_3OD$, 400MHz, δ, ppm) 8.63 (1H, d, *J* = 5.7 Hz), 8.59 (1H, d, *J* = 1.8 Hz), 8.31 (1H, dt, *J* = 8.1, 1.6 Hz,), 7.85 – 7.80 (1H, m), 7.48 (1H, d, *J* = 1.9 Hz), 7.44 – 7.40 (2H, m), 7.29 (1H, ddd, *J* = 8.3, 7.4, 1.7 Hz), 7.22 (1H, dd, *J* = 7.6, 1.7 Hz), 7.20 – 7.17 (1H, m), 7.06 (1H, dd, *J* = 8.3, 0.9 Hz), 6.99 (1H, td, *J* = 7.5, 1.0 Hz), 4.30 (2H, t, *J* = 6.0 Hz), 3.78 (4H, dd, *J* = 6.5, 4.0 Hz), 3.64 (4H, dd, *J* = 6.5, 4.0 Hz,), 3.28 (2H, t, *J* = 6.0 Hz). $^{13}$C NMR ($CD_3OD$, 100MHz, δ, ppm) 156.46, 148.87, 146.35, 142.48, 141.98, 141.47, 140.54, 131.70, 131.15, 130.66, 130.62, 128.76, 128.04, 122.68, 121.57, 118.77, 114.09, 68.66, 50.65, 43.52, 33.38. HRMS (ESI), found 360.2077 $C_{23}H_{25}N_3O$, $[M + H]^+$, requires 360.2076.

**1-(4-(2-(2-(Pyridin-3-yl)ethoxy)phenyl)pyridin-2-yl)piperazine (12j)**

Following General Procedure D, **9** (100 mg, 0.35 mmol) was reacted with 2-(4-*tert*-butoxycarbonylpiperazin-1-yl)pyridine-4-boronic acid, pinacol ester (174 mg, 0.45 mmol), the crude was purified by reverse phase chromatography (6g C18 column; MeCN in water 0-100%). The resulting residue was reacted according to General Procedure E. Afforded the title compound as a fluffy pale yellow solid (70 mg, 77%). $^1$H NMR ($CD_3OD$, 400MHz, δ, ppm) 8.76 (2H, s), 8.57 (1H, d, *J* = 8.0 Hz), 8.03 (2H, d, *J* = 6.6 Hz), 7.53 – 7.48 (2H, m), 7.44 (1H, d, *J* = 1.2 Hz), 7.25 – 7.18 (2H, m), 7.13 (1H, td, *J* = 7.6, 1.0 Hz), 4.44 (2H, t, *J* = 6.1 Hz), 4.11 – 4.05 (4H, m), 3.54 – 3.49 (4H, m), 3.38 (2H, t, *J* = 6.1 Hz,). $^{13}$C NMR ($CD_3OD$, 100MHz, δ, ppm) 157.11, 156.86, 153.88, 148.73, 142.62, 140.77, 136.87, 133.45, 131.79, 128.46, 127.09, 122.96, 117.43, 114.08, 113.55, 68.80, 44.68, 43.73, 33.35. HRMS (ESI), found 361.2028 $C_{22}H_{24}N_4O$, $[M + H]^+$, requires 361.2028.

**1-(4'-Fluoro-2'-(2-(pyridin-3-yl)ethoxy)-[1,1'-biphenyl]-3-yl)piperazine (12k)**

Following General Procedure D, **11** (100 mg, 0.33 mmol) was reacted with (3-(4-(*tert*-butoxycarbonyl)piperazin-1-yl)phenyl)boronic acid (129 mg, 0.42 mmol), the crude was purified by reverse phase chromatography (6g C18 column; MeCN in water 0-100%). The resulting residue was reacted according to General Procedure E. Afforded the title compound as a fluffy yellow solid (76 mg, 74%). $^1$H NMR ($CD_3OD$, 400MHz, δ, ppm) 8.69 (1H, d, *J* = 5.6 Hz), 8.60 (1H, s), 8.28 (1H, dt, *J* = 8.1, 1.6 Hz), 7.84 (1H, dd, *J* = 8.0, 5.8 Hz), 7.35 – 7.28 (1H, m), 7.22 (1H, dd, *J* = 8.5, 6.7 Hz), 7.07 (1H, ddd, *J* = 8.3, 2.5, 0.8 Hz), 7.00 – 6.96 (1H, m), 6.90 (1H, dd, *J* = 10.9, 2.5 Hz), 6.87 – 6.83 (1H, m), 6.76 (1H, td, *J* = 8.4, 2.5 Hz), 4.33 (2H, t, *J* = 5.8 Hz), 3.48 (4H, dd, *J* = 6.8, 3.1 Hz), 3.42 (4H, dd, *J* = 6.8, 3.0 Hz), 3.27 (2H, t, *J* = 5.8 Hz). $^{13}$C NMR ($CD_3OD$, 100MHz, δ, ppm) 164.41 (d, *J* = 243.7 Hz), 157.67 (d, *J* = 10 Hz), 157.62, 151.11, 148.94, 142.47, 141.29, 140.57, 140.32, 132.53 (d, *J* = 10 Hz), 129.94, 128.62 (d, *J* = 3.3 Hz), 127.97, 124.10, 119.47, 116.88, 108.62 (d, *J* = 21.2 Hz), 101.86 (d, *J* = 25.7 Hz), 68.87, 48.10, 44.73, 33.26. HRMS (ESI), found 378.1986 $C_{23}H_{24}FN_3O$, $[M + H]^+$, requires 378.1982.

**1-(4-(4-Fluoro-2-(2-(pyridin-3-yl)ethoxy)phenyl)pyridin-2-yl)piperazine (12l)**

Following General Procedure D, **11** (90 mg, 0.30 mmol) was reacted with (3-(4-(*tert*-butoxycarbonyl)piperazin-1-yl)phenyl)boronic acid (117 mg, 0.38 mmol), the crude was purified by reverse phase chromatography (6g C18 column; MeCN in water 0-100%). The resulting residue was reacted according to General Procedure E. Afforded the title compound as a fluffy off-white solid (50 mg, 69%). $^{1}$H NMR ($CD_3OD$, 400MHz, δ, ppm) 8.76 – 8.72 (2H, m), 8.58 – 8.54 (1H, m), 8.06 – 7.99 (2H, m), 7.54 (1H, dd, *J* = 8.6, 6.5 Hz), 7.44 – 7.41 (1H, m), 7.21 (1H, dd, *J* = 6.6, 1.5 Hz), 7.03 (1H, dd, *J* = 10.9, 2.4 Hz), 6.90 – 6.83 (1H, m), 4.42 (2H, t, *J* = 6.1 Hz), 4.10 – 4.04 (4H, m), 3.53 – 3.46 (4H, m), 3.37 (2H, t, *J* = 6.1 Hz,). $^{13}$C NMR ($CD_3OD$, 100MHz, δ, ppm) 166.54 (d, *J* = 248.6 Hz), 158.47 (d, *J* = 10.5 Hz), 156.32, 153.70, 148.89, 142.65, 140.93, 140.82, 136.64, 133.44 (d, *J* = 10.5 Hz), 128.49, 123.20 (d, *J* = 3.3 Hz), 117.23, 113.66, 109.49 (d, *J* = 21.9 Hz), 102.20 (d, *J* = 26.5 Hz), 69.26, 44.71, 43.71, 33.11. HRMS (ESI), found 379.1925 $C_{22}H_{23}FN_4O$, $[M + H]^+$, requires 379.1934.

**1-(3-(3-(2-(3,5-Dimethyl-1*H*-pyrazol-4-yl)ethoxy)pyridin-2-yl)phenyl)piperazine (12m)**

Following General Procedure D, **7** (35 mg, 0.12 mmol) was reacted with (3-(4-(*tert*-butoxycarbonyl)piperazin-1-yl)phenyl)boronic acid (45 mg, 0.15 mmol), the crude was purified by reverse phase chromatography (6g C18 column; MeCN in water 0-100%). The resulting residue was reacted according to General Procedure E. Afforded the title compound as a fluffy off-white solid (12 mg, 47%). $^{1}$H NMR ($CD_3OD$, 400MHz, δ, ppm) 8.34 (1H, d, *J* = 5.6 Hz), 8.30 (1H, d, *J* = 8.6 Hz), 7.94 (1H, dd, *J* = 8.6, 5.7 Hz), 7.44 (1H, t, *J* = 7.9 Hz), 7.33 – 7.27 (2H, m), 7.09 (1H, d, *J* = 7.5 Hz), 4.36 (2H, t, *J* = 5.8 Hz), 3.55 – 3.47 (4H, m), 3.39 – 3.32 (4H, m), 2.95 (2H, t, *J* = 5.6 Hz), 2.09 (6H, s). $^{1}$H NMR ($CD_3OD$, 400MHz, δ, ppm) 156.38, 151.85, 145.77, 144.98, 134.02, 131.17, 131.07, 130.65, 128.31, 122.92, 120.63, 118.65, 116.20, 70.67, 47.32, 44.58, 22.80, 9.60. HRMS (ESI), found 378.2294 $C_{22}H_{27}N_5O$, $[M + H]^+$, requires 378.2294.

**1-(4-(2-(2-(3,5-Dimethyl-1*H*-pyrazol-4-yl)ethoxy)phenyl)pyridin-2-yl)piperazine (12n)**

Following General Procedure D, **6** (90 mg, 0.30 mmol) was reacted with 2-(4-*tert*-butoxycarbonylpiperazin-1-yl)pyridine-4-boronic acid, pinacol ester (148 mg, 0.38 mmol), the crude was purified by reverse phase chromatography (6g C18 column; MeCN in water 0-100%). The resulting residue was reacted according to General Procedure E. Afforded the title compound as a fluffy off-white solid (44 mg, 79%). $^{1}$H NMR ($CD_3OD$, 400MHz, δ, ppm) 8.05 (1H, d, *J* = 6.6 Hz), 7.55 – 7.49 (3H, m), 7.29 (1H, dd, *J* = 6.6, 1.3 Hz), 7.21 (1H, d, *J* = 8.0 Hz), 7.14 (1H, td, *J* = 7.5, 0.8 Hz), 4.26 (2H, t, *J* = 6.6 Hz), 4.15 – 4.08 (4H, m), 3.56 – 3.50 (4H, m), 3.00 (2H, t, *J* = 6.6 Hz), 2.32 (6H, s). $^{13}$C NMR ($CD_3OD$, 100MHz, δ, ppm) 157.49, 157.06, 153.72, 145.58, 136.77, 133.45,

131.86, 127.06, 122.79, 117.48, 116.44, 114.16, 113.63, 68.48, 49.00, 44.71, 43.73, 23.12, 9.79. HRMS (ESI), found 378.2292 $C_{22}H_{27}N_5O$, $[M + H]^+$, requires 378.2294.

**3-(2-(3,5-Dimethyl-1*H*-pyrazol-4-yl)ethoxy-2'-(piperazin-1-yl)-2,4'-bypyridine (12o)**

Following General Procedure D, **7** (50 mg, 0.17 mmol) was reacted with 2-(4-*tert*-Butoxycarbonylpiperazin-1-yl)pyridine-4-boronic acid, pinacol ester (82 mg, 0.21 mmol), the crude was purified by reverse phase chromatography (6g C18 column; MeCN in water 0-100%). The resulting residue was reacted according to General Procedure E. Afforded the title compound as a fluffy off-white solid (25 mg, 80%). $^1$H NMR ($CD_3OD$, 400MHz, δ, ppm) 8.43 (1H, brs), 8.20 (1H, d, $J$ = 5.7 Hz), 7.93 (1H, d, $J$ = 8.3 Hz), 7.78 (1H, brs), 7.73 (1H, brs), 7.46 (1H, d, $J$ = 5.6 Hz), 4.37 (2H, t, $J$ = 5.6 Hz), 4.08 (4H, brs), 3.49 (4H, brs), 3.07 (2H, t, $J$ = 5.6 Hz), 2.32 (6H, s). $^{13}$C NMR ($CD_3OD$, 100MHz, δ, ppm) 155.47, 150.83, 146.32, 145.85, 142.76, 140.73, 138.10, 128.69, 125.21, 115.98, 115.65, 114.51, 112.87, 69.38, 44.43, 43.91, 23.05, 9.87. HRMS (ESI), found 379.2247 $C_{21}H_{26}N_6O$, $[M + H]^+$, requires 379.2246.

**3-(2'-(3,5-Dimethyl-1*H*-pyrazol-4-yl)ethoxy-4'-fluoro-[1,1'-biphenyl]-3-yl)piperazine (12p)**

Following General Procedure D, **8** (75 mg, 0.23 mmol) was reacted with (3-(4-(*tert*-butoxycarbonyl)piperazin-1-yl)phenyl)boronic acid (92 mg, 0.30 mmol), the crude was purified by reverse phase chromatography (6g C18 column; MeCN in water 0-100%). The resulting residue was reacted according to General Procedure E. Afforded the title compound as a fluffy pale yellow solid (12 mg, 47%). $^1$H NMR ($CD_3OD$, 400MHz, δ, ppm) 7.28 (1H, t, $J$ = 7.8 Hz), 7.19 (1H, dd, $J$ = 8.3, 6.9 Hz), 7.06 (1H, d, $J$ = 8.1 Hz), 7.01 (1H, brs), 6.88 (1H, dd, $J$ = 11.1, 2.2 Hz), 6.82 (1H, d, $J$ = 7.5 Hz), 6.74 (1H, td, $J$ = 8.3, 2.1 Hz), 4.13 (2H, t, $J$ = 5.3 Hz), 3.52 – 3.46 (4H, m), 3.45 – 3.40 (4H, m), 2.89 (2H, t, $J$ = 5.2 Hz), 2.11 (6H, s). $^{13}$C NMR ($CD_3OD$, 100MHz, δ, ppm) 164.39 (d, $J$ = 243.3 Hz), 158.06 (d, $J$ = 9.9 Hz), 151.24, 145.64, 140.78, 132.72 (d, $J$ = 9.8 Hz), 130.06, 128.66 (d, $J$ = 3.3 Hz), 123.95, 119.50, 116.94, 116.87, 108.22 (d, $J$ = 21.2 Hz), 101.64 (d, $J$ = 25.9 Hz), 68.96, 48.13, 44.73, 23.11, 9.50. HRMS (ESI), found 395.2247 $C_{23}H_{27}FN_4O$, $[M + H]^+$, requires 395.2247.

**1-(4-(2-(2-(3,5-Dimetyl-1H-pyrazol-4-yl)ethoxy)-4-fluorophenyl)piridin-2-yl)piperazine (12q)**

Following General Procedure D, **8** (30 mg, 0.10 mmol) was reacted with 2-(4-*tert*-Butoxycarbonylpiperazin-1-yl)pyridine-4-boronic acid, pinacol ester (47 mg, 0.12 mmol), the crude was purified by reverse phase chromatography (6g C18 column; MeCN in water 0-100%). The resulting residue was reacted according to General Procedure E. Afforded the title compound as a fluffy off-white solid (44 mg, 79%). $^1$H NMR ($CD_3OD$, 400MHz, δ, ppm) 8.05 (1H, d, $J$ = 6.6 Hz), 7.58 (1H, dd, $J$ = 8.5, 6.6 Hz), 7.53 (1H, s), 7.28 (1H, dd, $J$ = 6.6, 1.1 Hz), 7.05

(1H, dd, $J$ = 11.0, 2.3 Hz), 6.90 (1H, td, $J$ = 8.3, 2.3 Hz), 4.25 (2H, t, $J$ = 6.6 Hz), 4.14 – 4.08 (4H, m), 3.56 – 3.49 (4H, m), 3.01 (2H, t, $J$ = 6.6 Hz), 2.32 (6H, s). $^{13}C$ NMR ($CD_3OD$, 100MHz, δ, ppm) 166.51 (d, $J$ = 248.3 Hz), 158.70 (d, $J$ = 10.5 Hz), 156.51, 153.71, 145.63, 136.84, 133.48 (d, $J$ = 10.5 Hz), 129.93, 123.22 (d, $J$ = 3.3 Hz), 117.32, 116.22, 113.64, 109.31 (d, $J$ = 22.0 Hz), 102.23 (d, $J$ = 26.4 Hz), 68.96, 44.72, 43.72, 22.94, 9.79. HRMS (ESI), found 396.2201 $C_{22}H_{26}FN_5O$, $[M + H]^+$, requires 396.2200.

***tert*-Butyl 4-(6-chloropyrimidin-4-yl)piperazine-1-carboxylate (13)**

To a solution of 4,6-dichloro-pyrimidine (500 mg, 3.35 mmol) was dissolved in THF (10 mL) in a microwave vial (10-20 mL) and treated with *tert*-butyl piperazine-1-carboxylate (688 mg, 3.69 mmol) and DIPEA (0.87 mL, 5.03 mmol). The reaction mixture was heated at 145°C under microwave irradiation for 30 minutes. The resulting mixture was concentrated in a vacuum, and the residue was partitioned between DCM (50 mL) and water (50 mL). The organic fraction was washed with brine (50 mL), and the organic phase was dried over $Na_2SO_4$, concentrated under reduced pressure to yield the title compound (920 mg, 92%) as a light brown solid. The intermediary **13** was used without any purification in the following reactions. $^1H$ NMR ($CDCl_3$, 400MHz, δ, ppm) $^1H$ NMR (DMSO-$d_6$, 400 MHz, δ, ppm ) 8.32 (1H, s), 6.93 (1H, s), 3.62 (4H, brs), 3.39 – 3.33 (4H, m), 1.38 (9H, s).

***tert*-Butyl 4-(6-(2-hydroxyphenyl)pyrimidin-4-yl)piperazine-1-carboxylate (14)**

Following General Procedure D, **13** (200 mg, 0.67 mmol) was reacted with 2-hydroxybenzeneboronic acid (115 mg, 0.83 mmol). The crude was purified by flash column chromatography by elution with a gradient of pentane/EtOAc to afford 130 mg (54%) of the title compound as a beige solid. $^1H$ NMR ($CDCl_3$, 400MHz, δ, ppm) 8.49 (1H, d, $J$ = 1.0 Hz), 7.65 (1H, dd, $J$ = 8.1, 1.5 Hz), 7.28 (1H, td, $J$ = 7.9, 7.2, 1.6 Hz,), 6.94 (1H, dd, $J$ = 8.3, 1.1 Hz,), 6.85 – 6.80 (2H, m), 3.68 (4H, d, $J$ = 5.0 Hz), 3.55 – 3.50 (4H, m,), 1.47 (9H, s).

***tert*-Butyl 4-(6-(4-fluoro-2-hydroxyphenyl)pyrimidin-4-yl)piperazine-1-carboxylate (15)**

Following General Procedure D, **13** (90 mg, 0.25 mmol) was reacted with 4-fluoro-2-hydroxybenzene boronic acid (75 mg, 0.48 mmol). The crude was purified by flash column chromatography by elution with a gradient of pentane/EtOAc to afford 91 mg (63 %) of the title compound as a beige solid. $^1H$ NMR ($CDCl_3$, 400MHz, δ, ppm) 8.49 (1H, d, $J$ = 1.0 Hz), 7.64 (1H, dd, $J$ = 9.0, 7.4 Hz), 6.75 (1H, d, $J$ = 1.0 Hz), 6.64 (1H, dd, $J$ = 9.1, 2.6 Hz), 6.55 (1H, ddd, $J$ = 9.0, 7.4, 2.6 Hz), 3.75 – 3.69 (4H, m), 3.58 – 3.52 (4H, m), 1.48 (9H, s).

**4-(Piperazin-1-yl)-6-(2-(2-(1,3,5-trimethyl-1*H*-pyrazol-4-yl)ethoxy)phenyl)pyrimidine (16a)**

Following General Procedure C, **14** (70 mg, 0.19 mmol) was reacted with **1** (33 mg, 0.23 mmol) the crude was purified by reverse phase chromatography (6g C18 column; MeCN in water 0-100%). The resulting residue was reacted according to General Procedure E. Afforded the title compound as a fluffy yellow solid (8 mg 38%) $^{1}$H NMR ($CD_3OD$, 400MHz, δ, ppm) 8.84 (1H, brs), 7.70 – 7.56 (2H, m), 7.38 (1H, brs), 7.24 (1H, d, *J* = 8.3 Hz), 7.19 (1H, t, *J* = 7.1 Hz), 4.45 (2H, brs), 4.24 (4H, brs), 3.91 (3H, s), 3.47 (4H, brs), 3.02 (2H, brs), 2.33 (3H, s), 2.29 (3H, s). $^{13}$C NMR ($CD_3OD$, 100MHz, δ, ppm) 162.13, 155.89, 152.83, 150.44, 144.96, 142.95, 133.93, 130.88, 121.45, 119.13, 115.31, 112.59, 102.97, 67.33, 42.74, 40.91, 34.33, 21.94, 8.27, 8.19. HRMS (ESI), found 393.2403 $C_{22}H_{28}N_6O$, $[M + H]^+$, requires 393.2403.

**4-(4-Fluoro-2-(2-(1,3,5-trimethyl-1*H*-pyrazol-4-yl)ethoxy)phenyl)-6-(piperazin-1-yl)pyrimidine (16b)**

Following General Procedure C, **15** (70 mg, 0.18 mmol) was reacted with **1** (33 mg, 0.23 mmol) the crude was purified by reverse phase chromatography (6g C18 column; MeCN in water 0-100%). The resulting residue was reacted according to General Procedure E and afforded the title compound as a fluffy brownish white solid (8 mg 39%) $^{1}$H NMR ($CD_3OD$, 400MHz, δ, ppm) 8.86 (1H, brs), 7.77 – 7.68 (1H, m), 7.39 (1H, brs), 7.10 (1H, d, *J* = 10.5 Hz), 6.97 (1H, t, *J* = 6.9 Hz), 4.49 – 4.34 (2H, m), 4.23 (4H, brs), 3.92 (3H, s), 3.49 (4H, brs), 3.04 (2H, brs), 2.34 (3H, s), 2.30 (3H, s). $^{13}$C NMR ($CD_3OD$, 100MHz, δ, ppm) 167.57 (d, *J* = 205.7 Hz), 163.50, 159.12 (d, *J* = 10.9 Hz), 153.31, 151.91, 146.39, 144.38, 134.22 (d, *J* = 10.9 Hz), 116.92 (d, *J* = 2.9 Hz), 116.49, 109.63 (d, *J* = 22.4 Hz), 104.55, 102.35 (d, *J* = 26.8 Hz), 69.38, 44.18, 42.34, 35.81, 23.20, 9.74, 9.68. HRMS (ESI), found 411.2309 $C_{22}H_{27}FN_6O$, $[M + H]^+$, requires 411.2309.

**4-(Piperazin-1-yl)-6-(2-(2-(pyridine-3-yl)ethoxy)phenyl)pyrimidine (16c)**

Following General Procedure C, **14** (66 mg, 0.18 mmol) was reacted with 3-(2-hydroxyethyl)pyridine (28 mg, 0.23 mmol) the crude was purified by reverse phase chromatography (6g C18 column; MeCN in water 0-100%). The resulting residue was reacted according to General Procedure E and afforded the title compound as a fluffy brownish white solid (22 mg 45%) $^{1}$H NMR (DMSO-$d_6$, 400MHz, δ, ppm) 10.16 (1H, brs), 10.06 (1H, brs), 8.89 (1H, s), 8.82 (1H, s), 8.79 (1H, d, *J* = 5.0 Hz), 8.46 (1H, d, *J* = 7.8 Hz), 7.95 (1H, brt, *J* = 6.6 Hz), 7.61 – 7.58 (2H, m), 7.34 (1H, s), 7.28 (1H, d, *J* = 8.5 Hz), 7.16 (1H, t, *J* = 7.4 Hz), 4.41 (2H, t, *J* = 5.8 Hz), 4.19 (4H, brs), 3.34 – 3.23 (6H, m). $^{13}$C NMR (DMSO-$d_6$, 100MHz, δ, ppm) 161.35, 155.64, 152.03, 151.26, 145.70, 142.05, 140.15, 138.35, 133.48, 131.05, 126.56, 121.19, 119.93, 113.01, 103.36, 67.83, 48.53, 41.89, 31.43. HRMS (ESI), found 362.2000 $C_{21}H_{23}N_5O$, $[M + H]^+$, requires 362.1981.

**4-(4-Fluoro-2-(2-(pyridine-3-yl)ethoxy)phenyl)-6-(piperazin-1-yl)pyrimidine (16d)**

Following General Procedure C, **15** (88 mg, 0.23 mmol) was reacted with 3-(2-hydroxyethyl)pyridine (36 mg, 0.29 mmol) the crude was purified by reverse phase chromatography (6g C18 column; MeCN in water 0-100%). The resulting residue was reacted according to General Procedure E and afforded the title compound as a fluffy brownish white solid (8 mg 39%) $^{1}$H NMR (DMSO-$d_6$, 400MHz, δ, ppm) 9.88 (1H, brs), 8.87 (1H, brs), 8.78 (2H, d, $J$ = 8.6 Hz), 8.42 (1H, d, $J$ = 7.6 Hz), 7.97 – 7.90 (1H, m), 7.66 (1H, dd, $J$ = 8.5, 6.8 Hz), 7.32 (1H, brs), 7.25 (1H, dd, $J$ = 11.3, 2.1 Hz), 7.05 (1H, td, $J$ = 8.4, 2.1 Hz), 4.43 (2H, t, $J$ = 6.3 Hz), 4.16 (4H, brs), 3.31 – 3.26 (6H, m). $^{13}$C NMR (DMSO-$d_6$, 100MHz, δ, ppm) 165.02 (d, $J$ = 248.0 Hz), 161.33, 157.43 (d, $J$ = 11.0 Hz), 151.74, 145.36, 142.41, 140.55, 137.96, 132.83 (d, $J$ = 11.0 Hz), 126.46, 116.93, 108.02 (d, $J$ = 22.1 Hz), 103.41, 101.43 (d, $J$ = 25.5 Hz), 68.40, 42.03, 31.24. HRMS (ESI), found 380.1888 $C_{21}H_{22}FN_5O$, $[M + H]^+$, requires 380.1887.

**4-(2-(2-(3,5-Dimethyl-1*H*-pyrazol-4-yl)ethoxy)-4-fluorophenyl)-6-(piperazin-1-yl)pyrimidine (16e)**

Following General Procedure C, **15** (100 mg, 0.26 mmol) was reacted with **2** (46 mg, 0.32 mmol) the crude was purified by reverse phase chromatography (6g C18 column; MeCN in water 0-100%). The resulting residue was reacted according to General Procedure E and afforded the title as a fluffy brownish white solid (15 mg 39%) $^{1}$H NMR ($CD_3OD$, 400MHz, δ, ppm) 8.85 (1H, s), 7.74 (1H, dd, $J$ = 8.5, 6.4 Hz), 7.40 (1H, s), 7.12 (1H, dd, $J$ = 8.5, 2.1 Hz), 6.96 (1H, td, $J$ = 8.4, 2.0 Hz), 4.47 (2H brs), 4.31 – 4.17 (4H, m), 3.50 (4H, brs), 3.06 (2H, t, $J$ = 6.8 Hz), 2.35 (6H, s). $^{13}$C NMR ($CD_3OD$, 100MHz, δ, ppm) 167.61 (d, $J$ = 250.9 Hz), 163.50, 159.14 (d, $J$ = 11.0 Hz), 153.37, 151.87, 145.66, 134.15 (d, $J$ = 11.0 Hz), 116.98 (d, $J$ = 3.1 Hz), 115.94, 109.66 (d, $J$ = 22.4 Hz), 104.47, 102.38 (d, $J$ = 26.8 Hz), 69.22, 44.04, 22.80, 9.77. HRMS (ESI), found 397.2154 $C_{21}H_{25}FN_6O$, $[M + H]^+$, requires 397.2152.

**4-(2-(2-(3,5-Dimethyl-1*H*-pyrazol-4-yl)ethoxy)phenyl)-6-(piperazin-1-yl)pyrimidine (16f)**

Following General Procedure C, **14** (70 mg, 0.19 mmol) was reacted with **2** (34 mg, 0.24 mmol) the crude was purified by reverse phase chromatography (6g C18 column; MeCN in water 0-100%). The resulting residue was reacted according to General Procedure E and afforded the title compound as fluffy brownish white solid (41 mg, 72%) $^{1}$H NMR ($CD_3OD$, 400MHz, δ, ppm) 8.85 (1H, brs), 7.69 (1H, dd, $J$ = 7.6, 1.4 Hz), 7.66 – 7.61 (1H, m), 7.40 (1H, brs), 7.28 (1H, d, $J$ = 8.4 Hz), 7.20 (1H, t, $J$ = 7.5 Hz), 4.48 (2H, brs), 4.29 (2H, t, $J$ = 6.8 Hz), 4.22 (2H, brs), 3.50 (4H, brs), 3.05 (2H, t, $J$ = 6.8 Hz), 2.35 (6H, s). $^{13}$C NMR ($CD_3OD$, 100MHz, δ, ppm) 163.52, 157.30, 154.24, 151.79, 145.61, 135.34, 132.22, 122.87, 120.55, 116.18, 114.04, 104.35, 68.59, 43.97, 42.26, 22.93, 9.75. HRMS (ESI), found 379.2249 $C_{21}H_{26}N_6O$, $[M + H]^+$, requires 379.2246.

**3-(3-(2-(1,3,5-Trimethyl-1*H*-pyrazol-4-yl)ethoxy)pyridin-2-yl)phenol (17)**

Following General Procedure D, **4** (100 mg, 0.32 mmol) was reacted with 3-hydroxyphenylboronic acid (56 mg, 0.39 mmol), the crude was purified by flash column chromatography by elution with a gradient of pentane/EtOAc to afford 89 mg (74%) of the title compound as a beige solid. $^1$H NMR ($CDCl_3$, 400MHz, δ, ppm) 8.79 (1H, brs), 8.24 (1H, dd, *J* = 4.6, 1.4 Hz), 7.29 (1H, dt, *J* = 7.7, 1.3 Hz), 7.24 – 7.18 (2H, m), 7.16 (1H, dd, *J* = 8.3, 4.6 Hz), 6.85 – 6.80 (2H, m), 4.01 (2H, t, *J* = 6.6 Hz), 3.68 (3H, s), 2.81 (2H, t, *J* = 6.5 Hz), 2.10 (3H, s), 2.03 (3H, s).

**2'-(2-(1,3,5-Trimethyl-1*H*-pyrazol-4-yl)ethoxy)-[1,1'-biphenyl]-3-ol (18)**

Following General Procedure D, **3** (100 mg, 0.32 mmol) was reacted with 3-hydroxyphenylboronic acid (56 mg, 0.39 mmol), the crude was purified by flash column chromatography by elution with a gradient of pentane/EtOAc to afford 80 mg (76%) of the title compound as a beige solid. $^1$H NMR ($CDCl_3$, 400MHz, δ, ppm) 7.28 – 7.22 (4H, m), 7.00 – 6.95 (1H, m), 6.94 – 6.89 (2H, m), 6.84 (1H, ddd, *J* = 8.1, 2.5, 1.0 Hz), 6.08 (1H, dd, *J* = 2.3, 1.6 Hz), 4.09 (2H, t, *J* = 6.2 Hz), 3.72 (3H, s), 2.82 (2H, t, *J* = 6.2 Hz), 2.08 (3H, s), 2.04 (3H, s).

**4'-Fluoro-2'-(2-(1,3,5-trimethyl-1*H*-pyrazol-4-yl)ethoxy)-[1,1'-biphenyl]-3-ol (19)**

Following General Procedure D, **5** (90 mg, 0.27 mmol) was reacted with 3-hydroxyphenylboronic acid (48 mg, 0.34 mmol), **and** the crude was purified by flash column chromatography by elution with a gradient of pentane/EtOAc to afford 73 mg (78%) of the title compound as a beige solid. $^1$H NMR ($CDCl_3$, 400MHz, δ, ppm) 7.25 – 7.17 (2H, m), 6.87 – 6.82 (2H, m), 6.70 – 6.62 (2H, m), 6.10 – 6.08 (1H, m), 4.04 (2H, t, *J* = 6.2 Hz), 3.72 (3H, s), 2.82 (2H, t, *J* = 6.2 Hz), 2.09 (3H, s), 2.02 (3H, s).

**3-(3-(2-(Pyridin-3-yl)ethoxy)pyridin-2-yl)phenol (20)**

Following General Procedure D, **10** (100 mg, 0.35 mmol) was reacted with 3-hydroxyphenylboronic acid (61 mg, 0.44 mmol), and the crude was purified by flash column chromatography by elution with a gradient of pentane/EtOAc to afford 75 mg (71%) of the title compound as a beige solid. $^1$H NMR ($CDCl_3$, 400MHz, δ, ppm) 8.57 (1H, d, *J* = 1.7 Hz), 8.47 (1H, dd, *J* = 4.9, 1.6 Hz), 8.23 (1H, dd, *J* = 3.8, 2.2 Hz), 7.60 – 7.55 (1H, m), 7.30 – 7.26 (1H, m), 7.23 (2H, dt, *J* = 7.7, 1.4 Hz), 7.16 – 7.13 (2H, m), 6.99 (1H, dd, *J* = 2.5, 1.5 Hz), 6.92 (1H, ddd, *J* = 7.8, 2.5, 1.4 Hz), 4.10 (2H, t, *J* = 5.8 Hz), 3.02 (2H, t, *J* = 5.7 Hz).

**2'-(2-(Pyridin-3-yl)ethoxy)-[1,1'-biphenyl]-3-ol (21)**

Following General Procedure D, **9** (100 mg, 0.36 mmol) was reacted with 3-hydroxyphenylboronic acid (62 mg, 0.45 mmol), the crude was purified by flash column chromatography by elution with a gradient of pentane/EtOAc to afford 78 mg (74%) of the title compound as a beige solid. $^{1}$H NMR ($CDCl_3$, 400MHz, δ, ppm) 10.03 (1H, brs), 8.53 (1H, s), 8.45 (1H, d, *J* = 3.9 Hz), 7.54 (1H, d, *J* = 7.8 Hz), 7.34 – 7.20 (4H, m), 7.02 – 6.95 (2H, m), 6.89 (3H, m), 4.08 (2H, t, *J* = 5.7 Hz), 2.96 (2H, t, *J* = 5.6 Hz).

**4'-Fluoro-2'-(2-(pyridin-3-yl)ethoxy)-[1,1'-biphenyl]-3-ol (22)**

Following General Procedure D, **11** (100 mg, 0.33 mmol) was reacted with 3-hydroxyphenylboronic acid (58 mg, 0.42 mmol), and the crude was purified by flash column chromatography by elution with a gradient of pentane/EtOAc to afford 90 mg (86%) of the title compound as a beige solid. $^{1}$H NMR ($CDCl_3$, 400MHz, δ, ppm) 9.82 (1H, s), 8.53 (1H, d, *J* = 2.0 Hz), 8.46 (1H, dd, *J* = 4.9, 1.6 Hz), 7.57 (1H, dt, *J* = 7.8, 1.9 Hz), 7.30 – 7.18 (3H, m), 6.93 (1H, ddd, *J* = 8.2, 2.5, 0.9 Hz), 6.82 (1H, dt, *J* = 7.6, 1.2 Hz), 6.71 (1H, dd, *J* = 2.4, 1.7 Hz), 6.66 (1H, td, *J* = 8.2, 2.4 Hz), 6.57 (1H, dd, *J* = 11.0, 2.4 Hz), 4.05 (2H, t, *J* = 5.6 Hz), 2.99 (2H, t, *J* = 5.6 Hz).

**3-(3-(2-(3,5-Dimethyl-1*H*-pyrazol-4-yl)ethoxy)pyridin-2-yl)phenol (23)**

Following General Procedure D, **7** (55 mg, 0.18 mmol) was reacted with 3-hydroxyphenylboronic acid (32 mg, 0.23 mmol), and the crude was purified by flash column chromatography by elution with a gradient of pentane/EtOAc to afford 40 mg (70%) of the title compound as a beige solid. $^{1}$H NMR ($CDCl_3$, 400MHz, δ, ppm) 8.21 (1H, d, *J* = 5.8 Hz), 7.22 – 7.11 (4H, m), 6.81 – 6.77 (2H, m), 3.98 (2H, t, *J* = 5.4 Hz), 2.75 (2H, t, *J* = 5.4 Hz), 2.00 (6H, s).

**2'-(2-(3,5-Dimethyl-1*H*-pyrazol-4-yl)ethoxy)-[1,1'-biphenyl]-3-ol (24)**

Following General Procedure D, **6** (51 mg, 0.17 mmol) was reacted with 3-hydroxyphenylboronic acid (36 mg, 0.26 mmol), and the crude was purified by flash column chromatography by elution with a gradient of pentane/EtOAc to afford 43 mg (80%) of the title compound as a beige solid. $^{1}$H NMR ($CDCl_3$, 400MHz, δ, ppm) 7.26 – 7.23 (1H, m), 7.22 – 7.16 (2H, m), 6.94 (1H, td, *J* = 7.5, 1.0 Hz), 6.90 (1H, d, *J* = 8.3 Hz), 6.86 (1H, dt, *J* = 7.6, 1.2 Hz), 6.84 – 6.79 (1H, m), 6.49 – 6.46 (1H, m), 4.00 (2H, t, *J* = 6.2 Hz), 2.72 (2H, t, *J* = 6.2 Hz), 1.99 (6H, s).

**2'-(2-(3,5-Dimethyl-1*H*-pyrazol-4-yl)ethoxy)-4'-fluoro-[1,1'-biphenyl]-3-ol (25)**

Following General Procedure D, **8** (100 mg, 0.31 mmol) was reacted with 3-hydroxyphenylboronic acid (55 mg, 0.40 mmol), and the crude was purified by flash column chromatography by elution with a gradient of pentane/EtOAc t to afford 71 mg (68%) of the title compound as a beige solid. $^{1}$H NMR ($CDCl_3$, 400MHz, δ, ppm)

7.21 – 7.16 (1H, m), 7.13 (1H, dd, $J$ = 8.3, 6.9 Hz), 6.82 (2H, dd, $J$ = 8.3, 4.2 Hz), 6.68 – 6.60 (2H, m), 6.50 – 6.46 (1H, m), 3.98 (2H, t, $J$ = 6.1 Hz), 2.73 (2H, t, $J$ = 6.1 Hz), 1.99 (6H, s).

***N*,*N*-Dimethyl-2-(3-(3-(2-(1,3,5-trimethyl-1*H*-pyrazol-4-yl)ethoxy)pyridin-2-yl)phenoxy)-ethan-1-amine (26a)**

Following General Procedure C, **17** (100 mg, 0.31 mmol) was reacted with 2-(dimethylamino)ethanol (34 mg, 0.38 mmol). The crude was purified by reverse phase chromatography (6g C18 column; MeCN in water 0-100%) to afford 33 mg (41%) of the title compound as a brown resin. $^{1}$H NMR ($CD_3OD$, 400MHz, δ, ppm) 8.14 (1H, dd, $J$ = 4.7, 1.3 Hz), 7.51 (1H, dd, $J$ = 8.4, 1.3 Hz), 7.35 – 7.29 (2H, m), 7.25 (1H, dd, $J$ = 2.5, 1.5 Hz), 7.21 (1H, dt, $J$ = 7.6, 1.1 Hz), 7.01 (1H, ddd, $J$ = 8.2, 2.5, 1.0 Hz), 4.15 (2H, t, $J$ = 5.5 Hz), 4.09 (2H, t, $J$ = 6.5 Hz), 3.61 (3H, s), 2.89 (2H, t, $J$ = 5.5 Hz), 2.79 (2H, t, $J$ = 6.5 Hz), 2.42 (6H, s), 2.02 (3H, s), 1.97 (3H, s). $^{13}$C NMR ($CD_3OD$, 100MHz, δ, ppm) 159.68, 154.52, 149.60, 146.89, 141.57, 140.12, 139.22, 130.07, 124.94, 123.27, 122.12, 116.46, 115.85, 113.38, 69.82, 66.27, 58.95, 45.63, 35.73, 24.44, 11.60, 9.38. HRMS (ESI), found 395.2446 $C_{23}H_{30}N_4O_2$, $[M + H]^+$, requires 395.2447.

***N*,*N*-Dimethyl-2-((2'-(2-(1,3,5-trimethyl-1*H*-pyrazol-4-yl)ethoxy)-[1,1'-biphenyl]-3-yl)oxy)-ethan-1-amine (26b)**

Following General Procedure C, **18** (100 mg, 0.31 mmol) was reacted with 2-(dimethylamino)ethanol (34 mg, 0.38 mmol). The crude was purified by reverse phase chromatography (6g C18 column; MeCN in water 0-100%) to afford 36 mg (45%) of the title compound as a brown resin. $^{1}$H NMR ($CD_3OD$, 400MHz, δ, ppm) 7.27 – 7.19 (3H, m), 7.01 – 6.95 (3H, m), 6.93 (1H, ddd, $J$ = 7.6, 1.5, 0.9 Hz), 6.89 (1H, ddd, $J$ = 8.3, 2.6, 0.9 Hz), 4.08 (2H, t, $J$ = 5.5 Hz), 3.96 (2H, t, $J$ = 6.5 Hz), 3.58 (3H, s), 2.82 (2H, t, $J$ = 5.5 Hz), 2.70 (2H, t, $J$ = 6.5 Hz), 2.37 (6H, s), 1.98 (3H, s), 1.91 (3H, s). $^{13}$C NMR ($CD_3OD$, 100MHz, δ, ppm) 159.67, 157.13, 146.84, 141.77, 139.18, 132.35, 131.78, 129.97, 129.79, 123.35, 122.04, 116.86, 114.10, 113.98, 113.72, 69.71, 66.36, 59.06, 45.74, 35.69, 24.64, 11.56, 9.31. HRMS (ESI), found 394.2496 $C_{24}H_{31}N_3O_2$, $[M + H]^+$, requires 394.2495.

**2-((4'-Fluoro-2'-(2-(1,3,5-trimethyl-1*H*-pyrazol-4-yl)ethoxy)-[1,1'-biphenyl]-3-yl)oxy)-*N*,*N*-dimethylethan-1-amine (26c)**

Following General Procedure C, **19** (100 mg, 0.29 mmol) was reacted with 2-(dimethylamino)ethanol (32 mg, 0.36 mmol). The crude was purified by reverse phase chromatography (6g C18 column; MeCN in water 0-100%) to afford 60 mg (44%) of the title compound as a brown resin. $^{1}$H NMR ($CD_3OD$, 400MHz, δ, ppm) 7.24 (1H, t, $J$ = 7.9 Hz), 7.18 (1H, dd, $J$ = 8.4, 6.9 Hz), 6.95 – 6.92 (1H, m), 6.89 (2H, dt, $J$ = 7.5, 2.3 Hz), 6.78 (1H, dd, $J$ =

11.2, 2.4 Hz), 6.68 (1H, td, *J* = 8.3, 2.5 Hz), 4.09 (2H, t, *J* = 5.4 Hz), 3.97 (2H, t, *J* = 6.4 Hz), 3.58 (3H, s), 2.85 (2H, t, *J* = 5.4 Hz), 2.71 (2H, t, *J* = 6.4 Hz), 2.40 (6H, s), 1.97 (3H, s), 1.90 (3H, s). $^{13}$C NMR ($CD_3OD$, 100MHz, δ, ppm) 164.35 (d, *J* = 243.1 Hz), 159.67, 158.31 (d, *J* = 9.9 Hz), 146.85, 140.89, 139.21, 132.62 (d, *J* = 9.9 Hz), 130.07, 128.32 (d, *J* = 3.3 Hz), 123.38, 116.89, 114.06, 113.56, 107.94 (d, *J* = 21.2 Hz), 101.57 (d, *J* = 25.7 Hz), 69.91, 66.24, 58.99, 45.67, 39.46, 35.70, 24.41, 11.55, 9.30. HRMS (ESI), found 412.2401 $C_{24}H_{30}FN_3O_2$, $[M + H]^+$, requires 412.2400.

***N*,*N*-Dimethyl-2-(3-(3-(2-(pyridin-3-yl)ethoxy)pyridin-2-yl)phenoxy)ethan-1-amine (26d)**

Following General Procedure C, **20** (100 mg, 0.34 mmol) was reacted with 2-(dimethylamino)ethanol (37 mg, 0.42 mmol). The crude was purified by reverse phase chromatography (6g C18 column; MeCN in water 0-100%) to afford 30 mg (28%) of the title compound as a brown resin. $^1$H NMR ($CD_3OD$, 400MHz, δ, ppm) 8.39 (1H, s), 8.36 (1H, d, *J* = 4.2 Hz), 8.16 (1H, dd, *J* = 4.8, 1.3 Hz), 7.67 – 7.63 (1H, m), 7.54 (1H, dd, *J* = 8.4, 1.3 Hz), 7.36 – 7.26 (3H, m), 7.24 (1H, dd, *J* = 2.5, 1.5 Hz), 7.15 (1H, ddd, *J* = 7.7, 1.5, 1.0 Hz), 7.01 (1H, ddd, *J* = 8.2, 2.6, 1.0 Hz), 4.31 (2H, t, *J* = 6.1 Hz), 4.13 (2H, t, *J* = 5.5 Hz), 3.10 (2H, t, *J* = 6.0 Hz), 2.81 (2H, t, *J* = 5.5 Hz), 2.36 (6H, s). $^{13}$C NMR ($CD_3OD$, 100MHz, δ, ppm) 158.37, 152.90, 149.17, 148.02, 146.65, 140.30, 138.33, 137.60, 135.01, 128.50, 123.59, 123.53, 121.86, 120.46, 115.15, 114.39, 68.51, 65.20, 57.67, 44.38, 32.18. HRMS (ESI), found 364.2026 $C_{22}H_{25}N_3O_2$, $[M + H]^+$, requires 364.2025.

***N*,*N*-Dimethyl-2-((2'-(2-(pyridin-3-yl)ethoxy)-[1,1'-biphenyl]-3-yl)oxy)ethan-1-amine (26e)**

Following General Procedure C, **21** (95 mg, 0.32 mmol) was reacted with 2-(dimethylamino)ethanol (35 mg, 0.40 mmol). The crude was purified by reverse phase chromatography (6g C18 column; MeCN in water 0-100%) to afford 50 mg (46%) of the title compound as a brown resin. $^1$H NMR ($CD_3OD$, 400MHz, δ, ppm) 8.40 (1H, dd, *J* = 4.9, 1.4 Hz), 8.39 – 8.37 (1H, m), 7.70 – 7.66 (1H, m), 7.34 – 7.24 (4H, m), 7.04 – 7.02 (1H, m), 7.01 – 6.94 (4H, m), 4.29 – 4.26 (2H, m), 4.22 (2H, t, *J* = 5.8 Hz), 3.36 – 3.32 (2H, m), 3.02 (2H, t, *J* = 5.8 Hz), 2.78 (3H, s), 2.71 (3H, s). $^{13}$C NMR ($CD_3OD$, 100MHz, δ, ppm) 158.95, 156.85, 150.77, 147.91, 141.40, 139.08, 137.16, 131.58, 131.56, 130.08, 129.95, 125.15, 123.85, 122.08, 117.74, 113.38, 113.18, 69.68, 64.19, 58.16, 44.44, 39.45, 33.82. HRMS (ESI), found 363.2073 $C_{23}H_{26}N_2O_2$, $[M + H]^+$, requires 363.2073.

**2-((4'-fluoro-2'-(2-(pyridin-3-yl)ethoxy)-[1,1'-biphenyl]-3-yl)oxy)-*N*,*N*-dimethylethan-1-amine (26f)**

Following General Procedure C, **22** (110 mg, 0.35 mmol) was reacted with 2-(dimethylamino)ethanol (40 mg, 0.44 mmol). The crude was purified by reverse phase chromatography (6g C18 column; MeCN in water 0-100%) to afford 54 mg (40%) of the title compound as a brown resin. $^1$H NMR ($CD_3OD$, 400MHz, δ, ppm) 8.26 (2H, brs), 7.49 (1H, dt, $J$ = 7.9, 1.9 Hz), 7.21 – 7.13 (3H, m), 6.88 – 6.83 (2H, m), 6.80 – 6.77 (1H, m), 6.75 (1H, dd, $J$ = 11.1, 2.5 Hz), 6.64 (1H, td, $J$ = 8.3, 2.5 Hz), 4.11 (2H, t, $J$ = 6.0 Hz), 4.02 (2H, t, $J$ = 5.5 Hz), 2.94 (2H, t, $J$ = 5.9 Hz), 2.73 (2H, t, $J$ = 5.5 Hz), 2.29 (6H, s). $^{13}$C NMR ($CD_3OD$, 100MHz, δ, ppm) 164.35 (d, $J$ = 243.3 Hz), 159.73, 158.02 (d, $J$ = 9.9 Hz), 150.54, 147.93, 140.46, 139.09, 136.58, 132.48 (d, $J$ = 9.9 Hz), 129.95, 128.09 (d, $J$ = 3.3 Hz), 124.94, 123.47, 117.03, 113.88, 108.10 (d, $J$ = 21.2 Hz), 101.33 (d, $J$ = 25.8 Hz), 69.83, 66.39, 59.05, 45.76, 33.56. HRMS (ESI), found 381.1978 $C_{23}H_{25}FN_2O_2$, $[M + H]^+$, requires 381.1978.

**2-(3-(3-(2-(3,5-Dimethyl-1*H*-pyrazol-4-yl)ethoxy)pyridin-2-yl)phenoxy)-*N*,*N*-dimethylethan-1-amine (26g)**

Following General Procedure C, **23** (40 mg, 0.13 mmol) was reacted with 2-(dimethylamino)ethanol (14 mg, 0.16 mmol). The crude was purified by reverse phase chromatography (6g C18 column; MeCN in water 0-100%) to afford 22 mg (44%) of the title compound as a brown resin. $^1$H NMR ($CD_3OD$, 400MHz, δ, ppm) 8.14 (1H, d, $J$ = 4.7 Hz), 7.53 (1H, d, $J$ = 8.4 Hz), 7.36 – 7.30 (2H, m), 7.18 (1H, d, $J$ = 8.2 Hz), 7.16 – 7.14 (1H, m), 6.99 (1H, dd, $J$ = 8.2, 2.6 Hz), 4.14 – 4.10 (4H, m), 2.84 – 2.79 (4H, m), 2.37 (6H, s), 2.03 (6H, s). $^{13}$C NMR ($CD_3OD$, 100MHz, δ, ppm) 158.28, 153.11, 148.11, 140.00, 138.57, 128.67, 123.49, 121.63, 120.47, 115.20, 113.99, 110.93, 68.19, 64.98, 57.72, 44.32, 22.53. HRMS (ESI), found 381.2290 $C_{22}H_{28}N_4O_2$, $[M + H]^+$, requires 381.2291.

**2-((2'-(2-(3,5-Dimethyl-1*H*-pyrazol-4-yl)ethoxy)-[1,1'-biphenyl]-3-yl)oxy)-*N*,*N*-dimethylethan-1-amine (26h)**

Following General Procedure C, **24** (42 mg, 0.13 mmol) was reacted with 2-(dimethylamino)ethanol (15 mg, 0.17 mmol). The crude was purified by reverse phase chromatography (6g C18 column; MeCN in water 0-100%) to afford 24 mg (46%) of the title compound as a brown resin. $^1$H NMR ($CD_3OD$, 400MHz, δ, ppm) 7.28 – 7.20 (3H, m), 7.03 – 6.99 (1H, m), 6.98 (1H, dd, $J$ = 7.4, 1.1 Hz), 6.95 – 6.91 (2H, m), 6.88 (1H, ddd, $J$ = 8.2, 2.4, 1.1 Hz), 4.07 (2H, t, $J$ = 5.4 Hz), 4.00 (2H, t, $J$ = 6.5 Hz), 2.79 (2H, t, $J$ = 5.4 Hz), 2.75 (2H, t, $J$ = 6.5 Hz), 2.36 (6H, s), 2.00 (6H, s). $^{13}$C NMR ($CD_3OD$, 100MHz, δ, ppm) 159.66, 157.14, 141.69, 132.22, 131.82, 130.00, 129.77, 123.18, 121.93, 117.02, 113.85, 113.63, 112.71, 69.47, 66.33, 59.20, 45.79, 24.16. HRMS (ESI), found 380.2338 $C_{23}H_{29}N_3O_2$, $[M + H]^+$, requires 380.2338.

**2-((2'-(2-(3,5-Dimethyl-1*H*-pyrazol-4-yl)ethoxy)-4'-fluoro-[1,1'-biphenyl]-3-yl)oxy)-*N*,*N*-dimethylethan-1-amine (26i)**

Following General Procedure C, **25** (90 mg, 0.27 mmol) was reacted with 2-(dimethylamino)ethanol (30 mg, 0.34 mmol). The crude was purified by reverse phase chromatography (6g C18 column; MeCN in water 0-100%) to afford 40 mg (36%) of the title compound as a brown resin. $^1$H NMR ($CD_3OD$, 400MHz, δ, ppm) 7.26 – 7.22 (1H, m), 7.18 (1H, dd, *J* = 8.5, 6.8 Hz), 6.88 – 6.84 (3H, m), 6.79 (1H, dd, *J* = 11.2, 2.5 Hz), 6.67 (1H, td, *J* = 8.3, 2.5 Hz), 4.05 (2H, t, *J* = 5.4 Hz), 4.00 (2H, t, *J* = 6.3 Hz), 2.79 (2H, t, *J* = 5.3 Hz), 2.74 (2H, t, *J* = 6.3 Hz), 2.36 (6H, s), 1.98 (6H, s). $^{13}$C NMR ($CD_3OD$, 100MHz, δ, ppm) 162.93 (d, *J* = 243.1 Hz), 158.24, 156.90 (d, *J* = 9.9 Hz), 139.36, 131.26 (d, *J* = 9.9 Hz), 128.70, 126.75 (d, *J* = 3.4 Hz), 121.77, 115.68, 112.19, 111.13, 106.43 (d, *J* = 21.2 Hz), 99.95 (d, *J* = 25.4 Hz), 68.25, 64.76, 57.72, 44.30, 22.52. HRMS (ESI), found 398.2244 $C_{22}H_{28}N_6O$, [M + H]$^+$, requires 398.2244.

**1,3,5-Trimethyl-4-(2-(naphthalen-2-yloxy)ethyl)-1*H*-pyrazole (27)**

Following General Procedure C, **1** (130 mg, 0.85 mmol) was reacted with 1-bromo-2-naphthol (238 mg, 1.07 mmol). The crude was purified by flash column chromatography by elution with a gradient of pentane/EtOAc to afford 200 mg (64%) of the title compound as a yellow pale solid. $^1$H NMR ($CDCl_3$, 400MHz, δ, ppm) 8.24 – 8.20 (1H, m), 7.78 (1H, d, *J* = 2.6 Hz), 7.77 – 7.75 (1H, m), 7.56 (1H, ddd, *J* = 8.4, 6.9, 1.3 Hz), 7.39 (1H, ddd, *J* = 8.1, 6.8, 1.1 Hz), 7.19 (1H, d, *J* = 9.0 Hz), 4.16 (2H, t, *J* = 7.2 Hz), 3.71 (3H, s), 2.94 (2H, t, *J* = 7.1 Hz), 2.25 (3H, s), 2.24 (3H, s).

**3-(2-(Naphthalen-2-yloxy)ethyl)pyridine (28)**

Following General Procedure C, 3-(2-hydroxyethyl)pyridine (55 mg, 0.44 mmol) was reacted with 1-bromo-2-naphthol (124 mg, 0.55 mmol). The crude was purified by flash column chromatography by elution with a gradient of pentane/EtOAc to afford 130 mg (89%) of the title compound as a yellow pale solid. $^1$H NMR ($CDCl_3$, 400MHz, δ, ppm) 8.61 (1H, d, *J* = 1.9 Hz), 8.47 (1H, dd, *J* = 4.8, 1.6 Hz), 8.20 – 8.16 (1H, m), 7.73 – 7.67 (3H, m), 7.51 (1H, ddd, *J* = 8.4, 6.9, 1.2 Hz), 7.34 (1H, ddd, *J* = 8.1, 6.9, 1.1 Hz), 7.22 – 7.18 (1H, m), 7.08 (1H, d, *J* = 9.0 Hz), 4.25 (2H, t, *J* = 6.4 Hz), 3.10 (2H, t, *J* = 6.3 Hz).

**3,5-Dimethyl-4-(2-(naphthalen-2-yloxy)ethyl)-1*H*-pyrazole (29)**

Following General Procedure C, **2** (132 mg, 0.94 mmol) was reacted with 1-bromo-2-naphthol (238 mg, 1.07 mmol). The crude was purified by flash column chromatography by elution with a gradient of pentane/EtOAc to afford 213 mg (65%) of the title compound as a yellow pale solid. $^1$H NMR ($CDCl_3$, 400MHz, δ, ppm) 8.22 (1H, dt, *J* = 8.6, 0.9 Hz), 7.77 (2H, d, *J* = 8.9 Hz), 7.59 – 7.53 (1H, m), 7.44 – 7.36 (1H, m), 7.19 (1H, d, *J* = 9.0 Hz), 4.18 (2H, t, *J* = 7.1 Hz), 2.97 (2H, t, *J* = 7.1 Hz), 2.31 (6H, s).

**1-(4-(2-(2-(1,3,5-Trimethyl-1*H*-pyrazol-4-yl)ethoxy)naphthalen-1-yl)pyridin-2-yl)piperazine (30a)**

Following General Procedure D, **27** (90 mg, 0.25 mmol) was reacted with 2-(4-*tert*-butoxycarbonylpiperazin-1-yl)pyridine-4-boronic acid, pinacol ester (117 mg, 0.30 mmol). The crude was purified by reverse phase chromatography (6g C18 column; MeCN in water 0-100%). The resulting residue was reacted according to General Procedure E. Afforded the title compound as a fluffy pale yellow solid (14 mg, 36%). $^{1}$H NMR ($CD_3OD$, 400MHz, δ, ppm) 8.18 (1H, d, *J* = 5.7 Hz), 7.95 (1H, d, *J* = 9.1 Hz), 7.83 (1H, d, *J* = 7.1 Hz), 7.45 (1H, d, *J* = 9.1 Hz), 7.38 – 7.32 (3H, m), 7.16 (1H, brs), 6.85 (1H, d, *J* = 5.6 Hz), 4.20 – 4.14 (2H, m), 3.96 (4H, brs), 3.78 (3H, s), 3.42 (4H, brs), 2.79 (2H, t, *J* = 5.8 Hz), 2.11 (3H, s), 2.06 (3H, s). $^{13}$C NMR ($CD_3OD$, 100MHz, δ, ppm) 156.37, 154.07, 153.83, 145.48, 143.23, 142.18, 133.38, 132.13, 130.58, 129.31, 128.33, 125.27, 125.02, 123.19, 119.05, 115.99, 115.89, 114.34, 70.47, 44.30, 44.02, 35.77, 24.19, 10.40, 9.42. HRMS (ESI), found 442.2606 $C_{27}H_{31}N_5O$, $[M + H]^+$, requires 442.2607.

***N*,*N*-Dimethyl-2-(3-(2-(2-(1,3,5-trimethyl-1*H*-pyrazol-4-yl)ethoxy)naphthalen-1-yl)phenoxy)ethan-1-amine (30b)**

Following General Procedure D, **27** (18 mg, 0.05 mmol) was reacted with 3-hydroxyphenylboronic acid (9 mg, 0.06 mmol). The crude was purified by flash column chromatography by elution with a gradient of pentane/EtOAc. The resulting residue was reacted (10 mg, 0.02 mmol) with 2-(dimethylamino)ethanol (4 mg, 0.04 mmol) according to General Procedure C. The crude was purified by reverse phase chromatography (6g C18 column; MeCN in water 0-100%) to afford the title compound as a brown resin (8 mg, 67%). 7.85 (1H, d, *J* = 9.0 Hz), 7.83 – 7.77 (1H, m), 7.40 – 7.33 (3H, m), 7.32 – 7.25 (2H, m), 7.04 – 6.99 (1H, m), 6.82 – 6.78 (2H, m), 4.09 (2H, td, *J* = 5.6, 1.4 Hz), 4.03 (2H, t, *J* = 6.3 Hz), 3.60 (3H, s), 2.78 (2H, t, *J* = 5.5 Hz), 2.64 (2H, t, *J* = 6.4 Hz), 2.34 (6H, s), 1.97 (3H, s), 1.87 (3H, s). $^{13}$C NMR ($CD_3OD$, 100MHz, δ, ppm) 160.09, 154.48, 146.78, 139.60, 139.28, 134.92, 130.69, 130.22, 128.89, 127.19, 126.06, 124.61, 118.13, 116.52, 114.33, 113.81, 71.13, 66.59, 59.15, 45.88, 35.67, 24.88, 11.48, 9.21. HRMS (ESI), found 444.2651 $C_{28}H_{33}N_3O_2$, $[M + H]^+$, requires 444.2651.

**1-(4-(2-(2-(Pyridin-3-yl)ethoxy)naphthalen-1-yl)pyridin-2-yl)piperazine (30c)**

Following General Procedure D, **28** (89 mg, 0.27 mmol) was reacted with 2-(4-*tert*-butoxycarbonylpiperazin-1-yl)pyridine-4-boronic acid, pinacol ester (131 mg, 0.34 mmol). The crude was purified by reverse phase chromatography (6g C18 column; MeCN in water 0-100%). The resulting residue was reacted according to General Procedure E. Afforded the title compound as a fluffy pale yellow solid (60 mg, 88%). $^{1}$H NMR ($CD_3OD$, 400MHz, δ, ppm) 8.77 (1H, brs), 8.71 (1H, brs), 8.45 (1H, d, *J* = 8.1 Hz), 8.15 (1H, d, *J* = 6.2 Hz), 8.04 – 7.98

(2H, m), 7.88 (1H, dd, $J$ = 7.5, 1.4 Hz), 7.52 (1H, d, $J$ = 9.2 Hz), 7.46 – 7.37 (3H, m), 7.33 (1H, brs), 6.92 (1H, d, $J$ = 6.1 Hz), 4.49 (2H, q, $J$ = 5.8 Hz), 4.08 – 4.01 (4H, m), 3.53 – 3.46 (4H, m), 3.31 – 3.27 (2H, m). $^{13}$C NMR ($CD_3OD$, 100MHz, δ, ppm) 155.53, 154.60, 153.56, 148.66, 142.75, 141.29, 140.86, 139.09, 133.09, 132.71, 130.64, 129.45, 128.69, 128.26, 125.48, 124.81, 122.17, 118.98, 115.66, 115.31, 69.43, 44.52, 43.80, 33.52. HRMS (ESI), found 411.2185 $C_{26}H_{26}N_4O$, $[M + H]^+$, requires 411.2185.

***N,N*-Dimethyl-2-(3-(2-(2-(pyridin-3-yl)ethoxy)naphthalen-1-yl)phenoxy)ethan-1-amine (30d)**

Following General Procedure D, **28** (100 mg, 0.30 mmol) was reacted with 3-hydroxyphenylboronic acid (52 mg, 0.38 mmol). The crude was purified by flash column chromatography by elution with a gradient of pentane/EtOAc. The resulting residue was reacted (90 mg, 0.26 mmol) with 2-(dimethylamino)ethanol (29 mg, 0.33 mmol) according to General Procedure C. The crude was purified by reverse phase chromatography (6g C18 column; MeCN in water 0-100%) to afford the title compound as a brown resin (85 mg, 79%). $^1$H NMR ($CD_3OD$, 400MHz, δ, ppm) 8.17 (1H, dd, $J$ = 4.9, 1.5 Hz), 8.08 (1H, d, $J$ = 1.7 Hz), 7.66 (1H, d, $J$ = 8.9 Hz), 7.64 – 7.60 (1H, m), 7.32 – 7.28 (1H, m), 7.26 – 7.21 (1H, m), 7.16 – 7.11 (4H, m), 6.99 (1H, ddd, $J$ = 7.8, 4.9, 0.7 Hz), 6.89 (1H, ddd, $J$ = 8.3, 2.6, 0.9 Hz), 6.67 (1H, dd, $J$ = 2.5, 1.4 Hz), 6.63 (1H, dt, $J$ = 7.5, 1.2 Hz), 3.99 (2H, t, $J$ = 5.9 Hz), 4.00 – 3.85 (2H, m), 2.69 (2H, t, $J$ = 5.8 Hz), 2.57 (2H, t, $J$ = 5.5 Hz), 2.15 (6H, s). $^{13}$C NMR ($CD_3OD$, 100MHz, δ, ppm) 159.93, 153.98, 150.33, 147.71, 139.18, 139.11, 136.59, 134.68, 130.51, 130.28, 130.22, 128.94, 127.30, 126.65, 126.04, 124.76, 124.68, 124.63, 118.19, 115.75, 114.20, 70.29, 66.47, 59.02, 45.85, 33.87. HRMS (ESI), found 413.2229 $C_{27}H_{28}N_2O_2$, $[M + H]^+$, requires 413.2229.

**1-(4-(2-(2-(3,5-Dimethyl-1*H*-pyrazol-4-yl)ethoxy)naphthalen-1-yl)pyridin-2-yl)piperazine (30e)**

Following General Procedure D, **29** (40 mg, 0.11 mmol) was reacted with 2-(4-*tert*-butoxycarbonylpiperazin-1-yl)pyridine-4-boronic acid, pinacol ester (56 mg, 0.14 mmol). The crude was purified by reverse phase chromatography (6g C18 column; MeCN in water 0-100%). The resulting residue was reacted according to General Procedure E. Afforded the title compound as a fluffy pale yellow solid (12 mg, 50%). $^1$H NMR ($CD_3OD$, 400MHz, δ, ppm) 8.21 (1H, d, $J$ = 6.4 Hz), 8.05 (1H, d, $J$ = 9.1 Hz), 7.92 – 7.88 (1H, m), 7.54 (1H, d, $J$ = 9.2 Hz), 7.52 (1H, brs), 7.50 – 7.39 (3H, m), 7.09 (1H, dd, $J$ = 6.4, 1.1 Hz), 4.32 (2H, q, $J$ = 6.3 Hz), 4.18 – 4.11 (4H, m), 3.58 – 3.52 (4H, m), 2.94 (2H, t, $J$ = 6.2 Hz), 2.29 (6H, s). $^{13}$C NMR ($CD_3OD$, 100MHz, δ, ppm) 157.09, 153.95, 153.61, 145.53, 137.48, 132.97, 132.88, 130.51, 129.43, 128.76, 125.47, 124.76, 121.73, 119.16, 116.71, 116.45, 115.54, 69.69, 44.71, 43.67, 23.33, 9.70. HRMS (ESI), found 428.2450 $C_{26}H_{29}N_5O$, $[M + H]^+$, requires 428.2450.

**2-(3-(2-(2-(3,5-Dimethyl-1*H*-pyrazol-4-yl)ethoxy)naphthalen-1-yl)phenoxy)-*N*,*N*-dimethylethan-1-amine (30f)**

Following General Procedure D, **29** (100 mg, 0.28 mmol) was reacted with 3-hydroxyphenylboronic acid (48 mg, 0.34 mmol). The crude was purified by flash column chromatography by elution with a gradient of pentane/EtOAc. The resulting residue was reacted (100 mg, 0.28 mmol) with 2-(dimethylamino)ethanol (30 mg, 0.33 mmol) according to General Procedure C. The crude was purified by reverse phase chromatography (6g C18 column; MeCN in water 0-100%) to afford the title compound as a brown resin (70 mg, 60%). $^{1}$H NMR ($CDCl_3$, 400MHz, δ, ppm) 7.83 (1H, d, *J* = 9.1 Hz), 7.81 – 7.77 (1H, m), 7.44 – 7.40 (1H, m), 7.39 – 7.34 (1H, m), 7.32 – 7.28 (3H, m), 6.93 – 6.87 (2H, m), 6.56 (1H, dd, *J* = 2.5, 1.5 Hz), 4.23 – 4.17 (1H m), 4.12 (1H, ddd, *J* = 10.7, 7.7, 3.3 Hz), 4.01 (2H, ddd, *J* = 8.1, 6.4, 3.7 Hz), 2.93 (1H, ddd, *J* = 13.3, 7.6, 3.6 Hz), 2.74 – 2.68 (3H, m), 2.46 (6H, s), 1.96 (6H, s). $^{13}$C NMR ($CDCl_3$, 100MHz, δ, ppm) 158.64, 153.15, 137.97, 133.70, 129.05, 128.89, 128.77, 127.77, 126.21, 125.40, 125.11, 123.58, 123.48, 117.97, 114.53, 111.74, 111.33, 69.18, 65.18, 58.59, 45.82, 23.55, 10.81. HRMS (ESI), found 430.2496 $C_{27}H_{31}N_3O_2$, $[M + H]^+$, requires 430.2495.

### Supplementary Figures

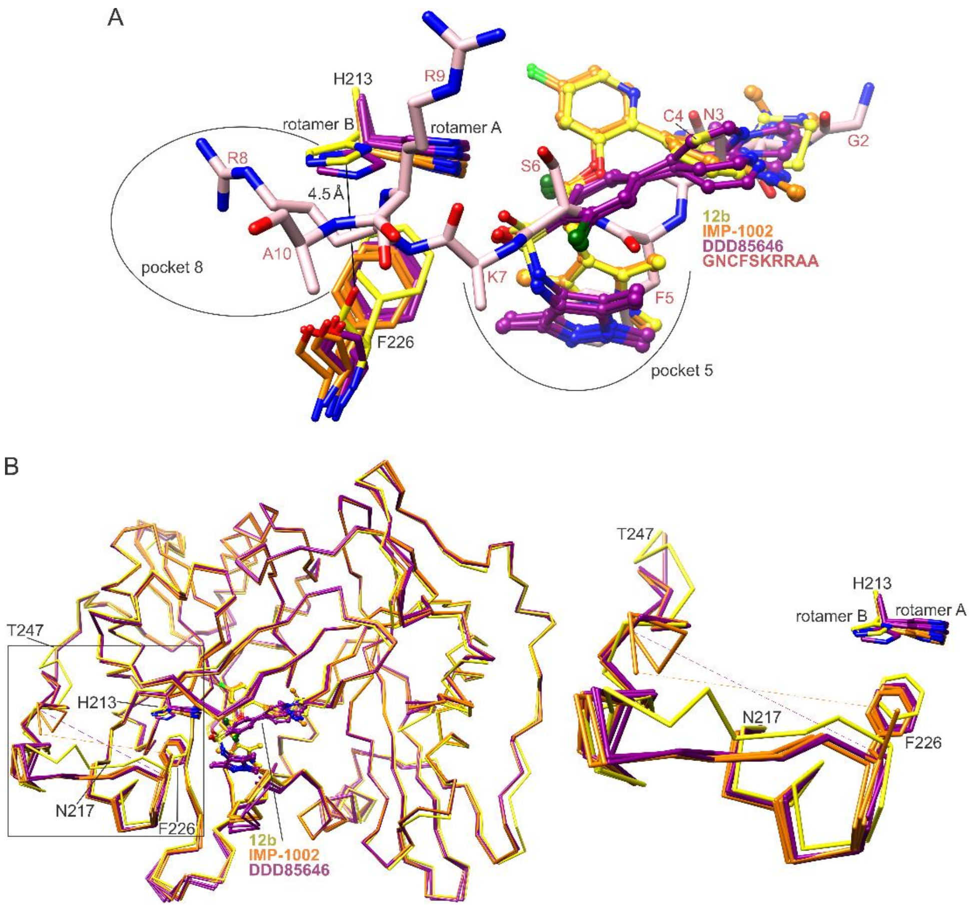

**Fig. S1.** Conformational changes in pocket 8 accompanying binding to **12b**. (**A**) Binding to **12b** induces conformational changes in Phe226 and His213 that narrow pocket 8 (*37*) to distances incompatible with peptide binding. (**B**) Associated conformational change in residues 217-247, boxed in *Cα* trace (left) and enlarged (right). In panels A and B, crystal structures of **IMP-1002**-bound *Pv*NMT (orange, PDB entry 6MB1 (*13*)), **DDD85646**-bound *Pv*NMT- (dark magenta, PDB entry 2YND (*12*)), and peptide-bound *Hs*NMT1 (pink, PDB entry 6QRM (*38*)) are superimposed (*36*) onto that of **12b–**bound *Pv*NMT (yellow).

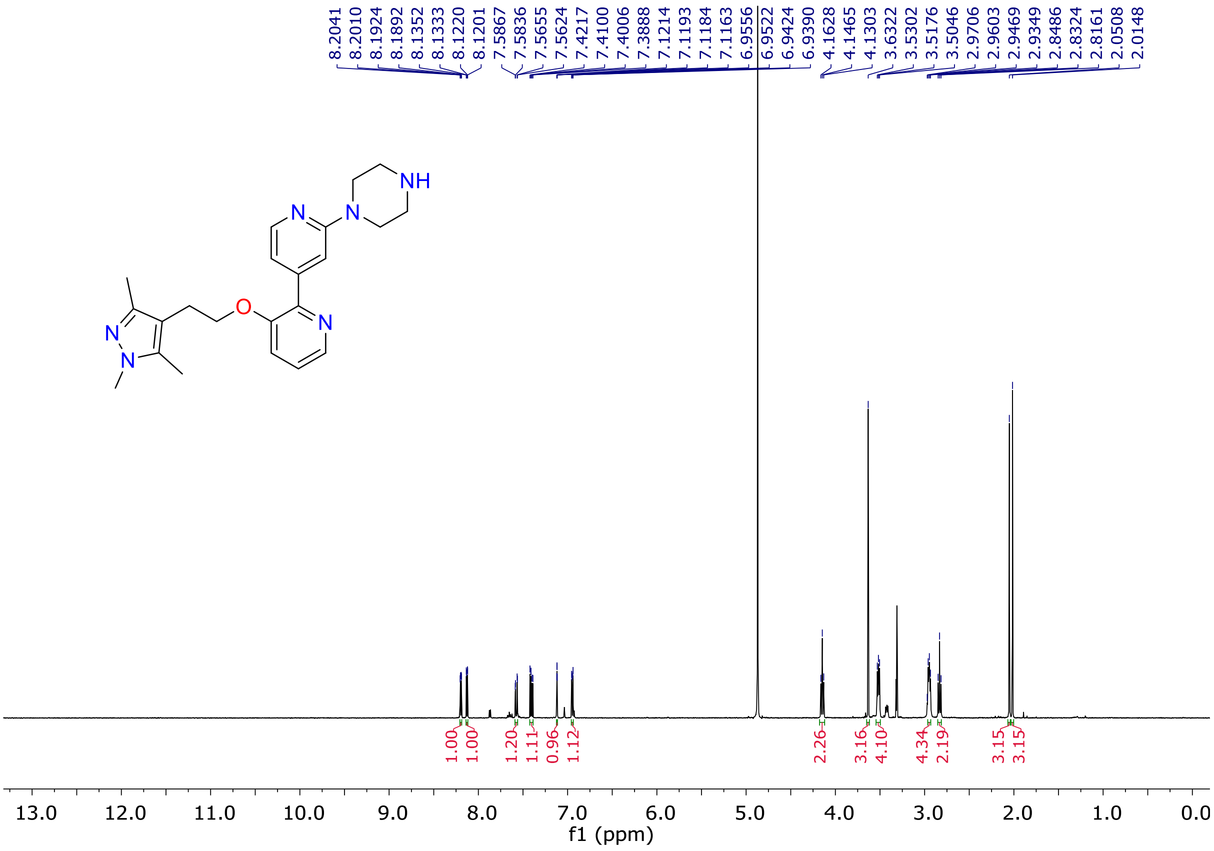


**Fig. S2.** [1]H-NMR spectrum of compound **12a** ($CD_3OD$, 400 MHz).

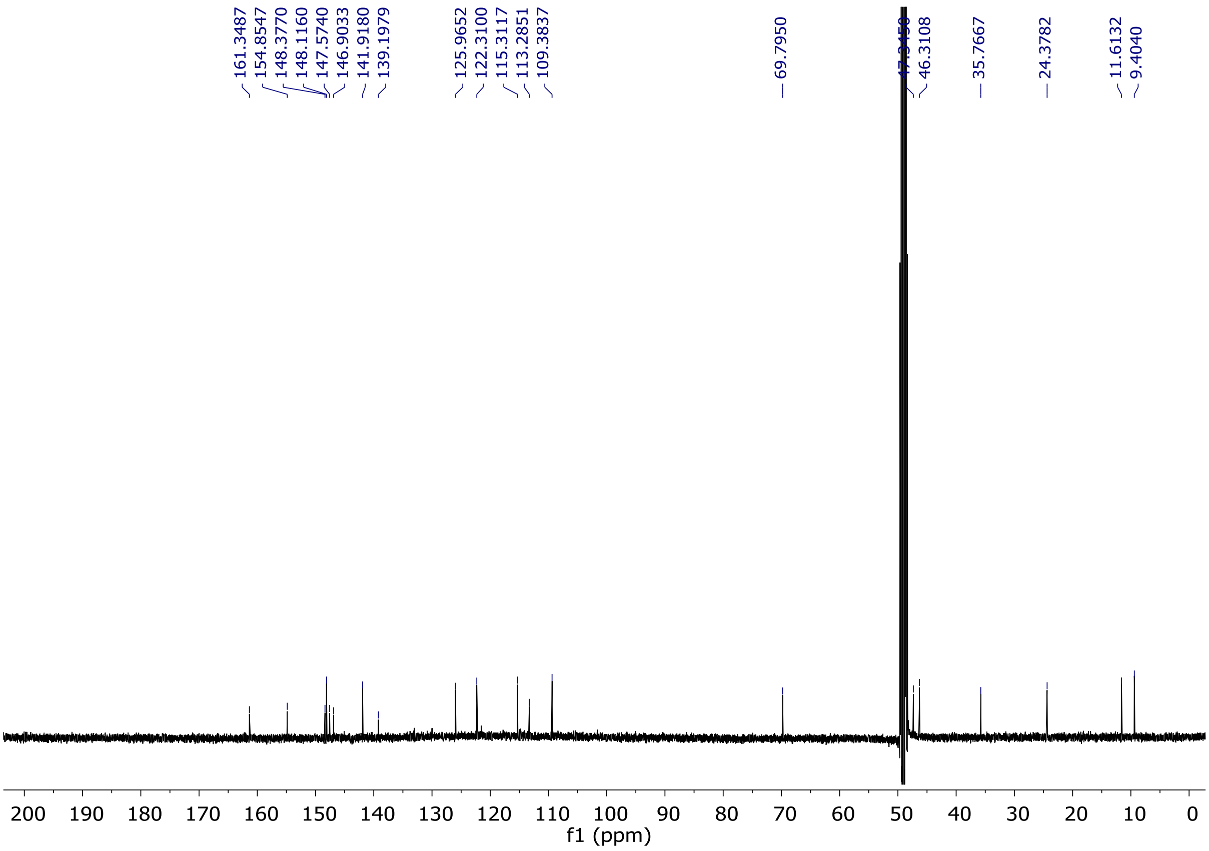


**Fig. S3.** $^{13}$C-NMR spectrum of compound **12a** ($CD_3OD$, 100 MHz).

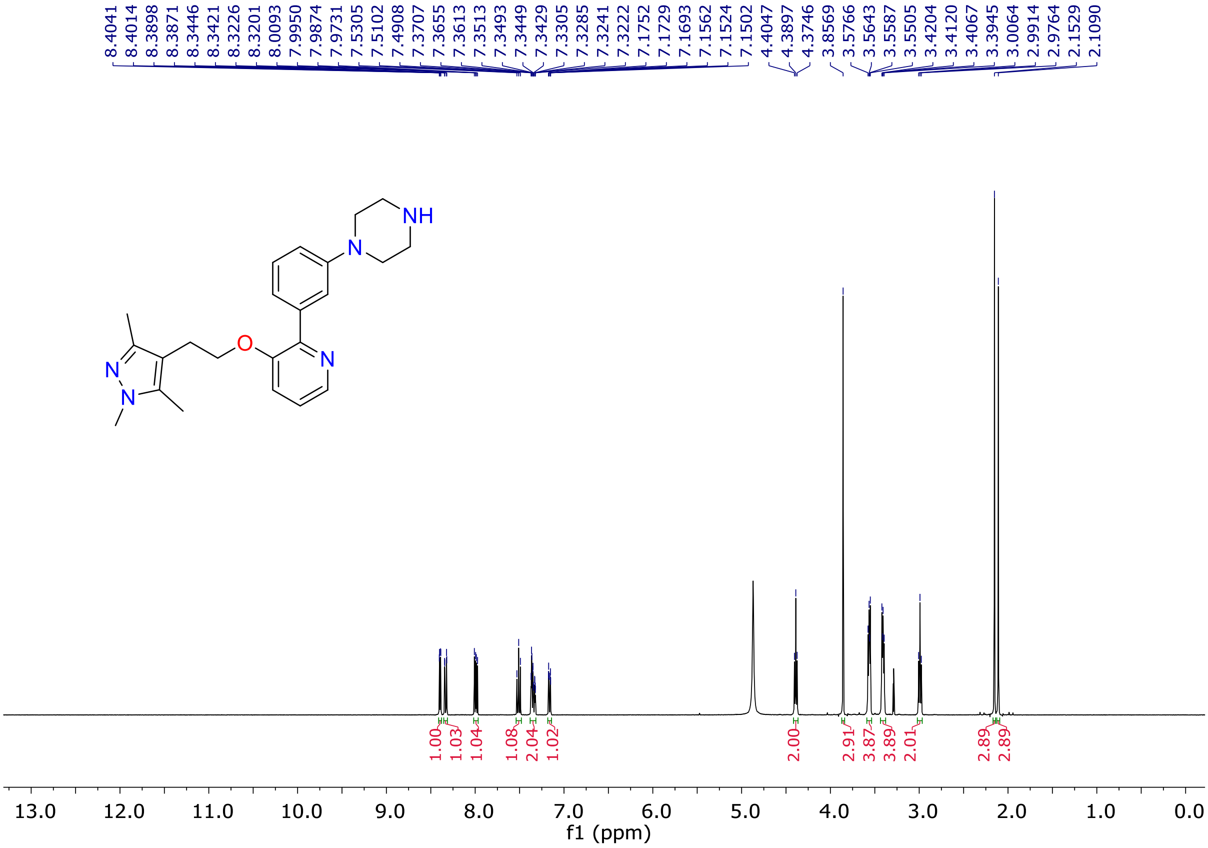


**Fig. S4.** $^{1}$H-NMR spectrum of compound **12b** ($CD_3OD$, 400 MHz).

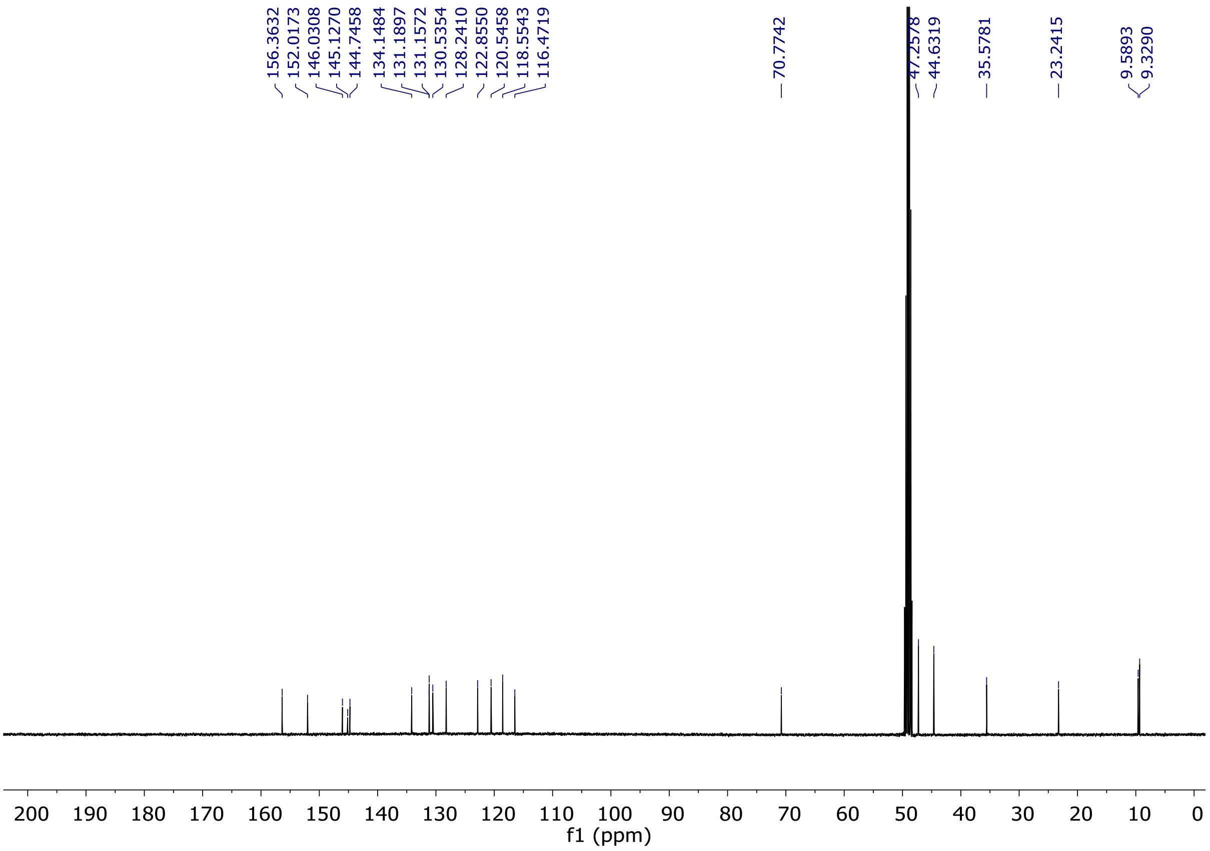


**Fig. S5.** $^{13}$C-NMR spectrum of compound **12b** ($CD_3OD$, 100 MHz).

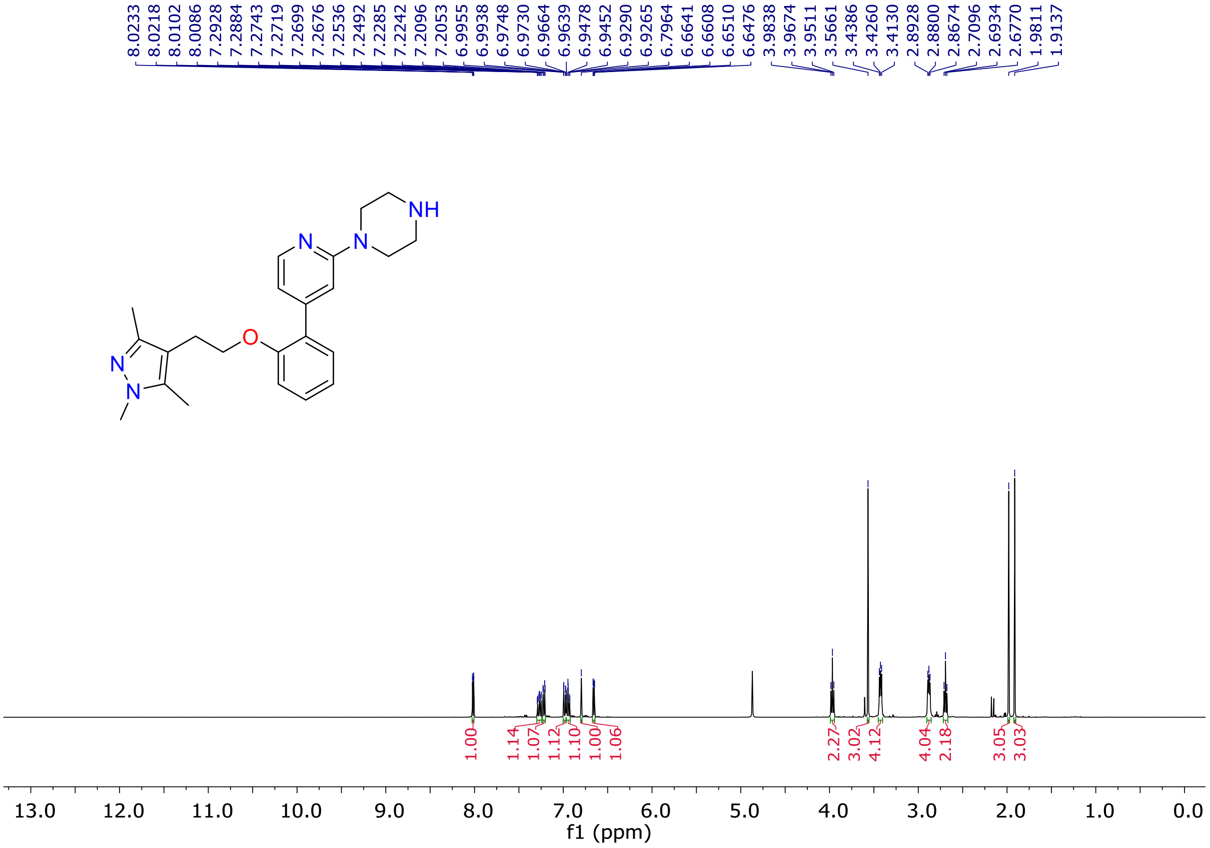


**Fig. S6.** $^{1}$H-NMR spectrum of compound **12c** ($CD_3OD$, 400 MHz).

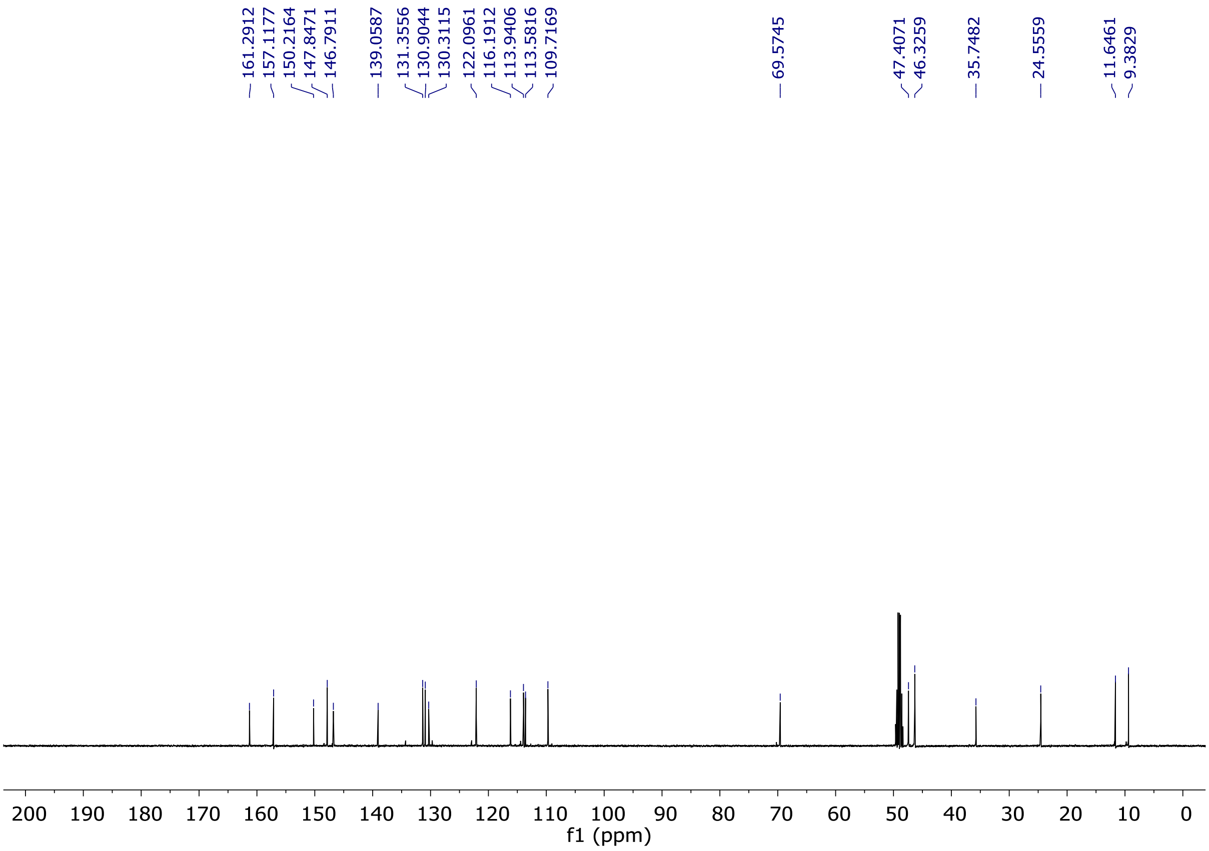


**Fig. S7.** [13]C-NMR spectrum of compound **12c** ($CD_3OD$, 100 MHz).

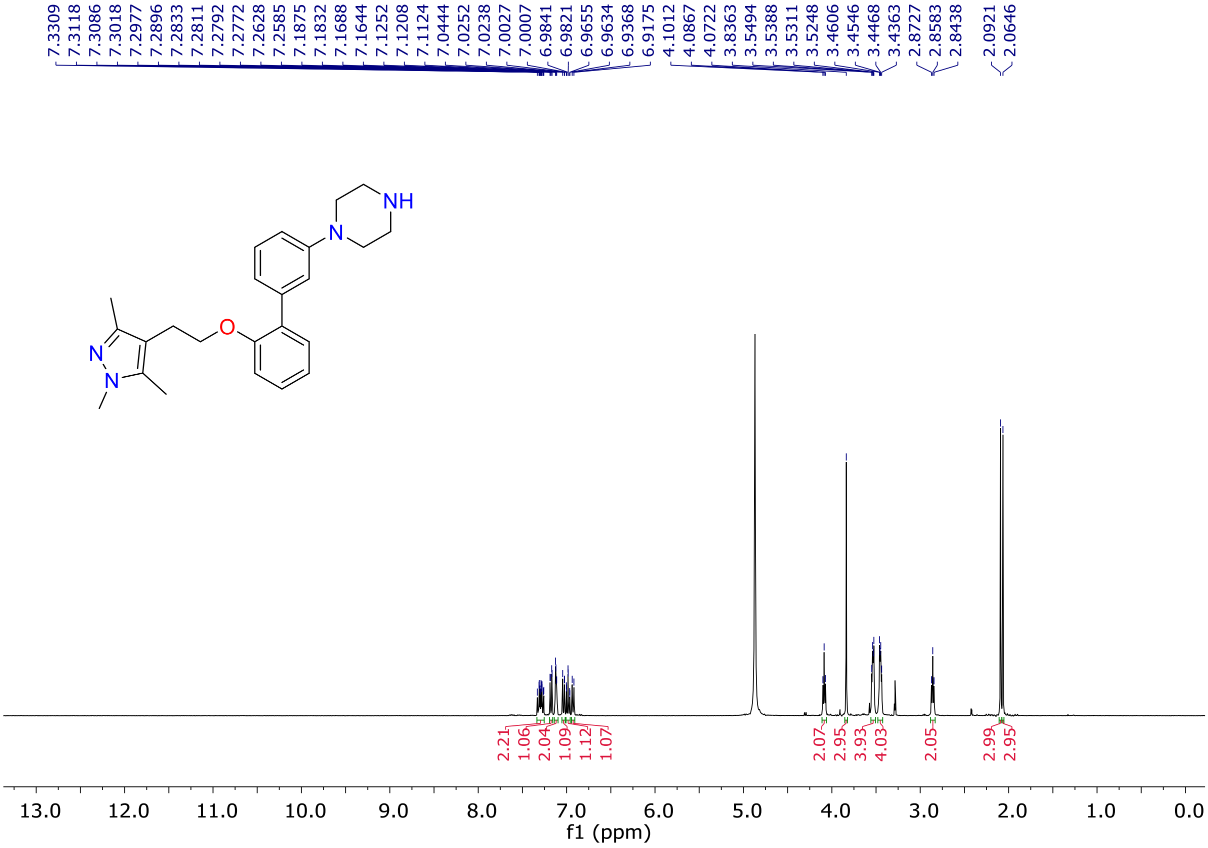


**Fig. S8.** [1]H-NMR spectrum of compound **12d** ($CD_3OD$, 400 MHz).

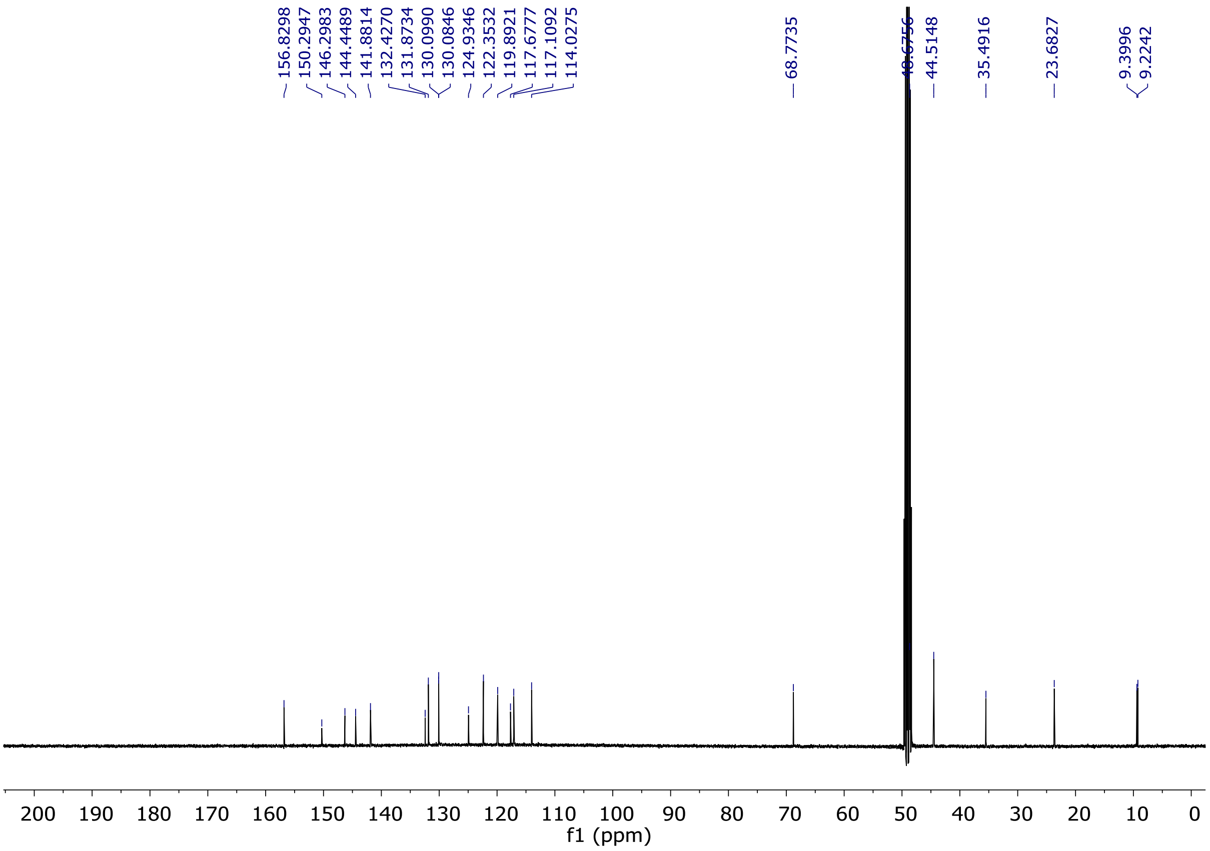


**Fig. S9.** [13]C-NMR spectrum of compound **12d** ($CD_3OD$, 100 MHz).

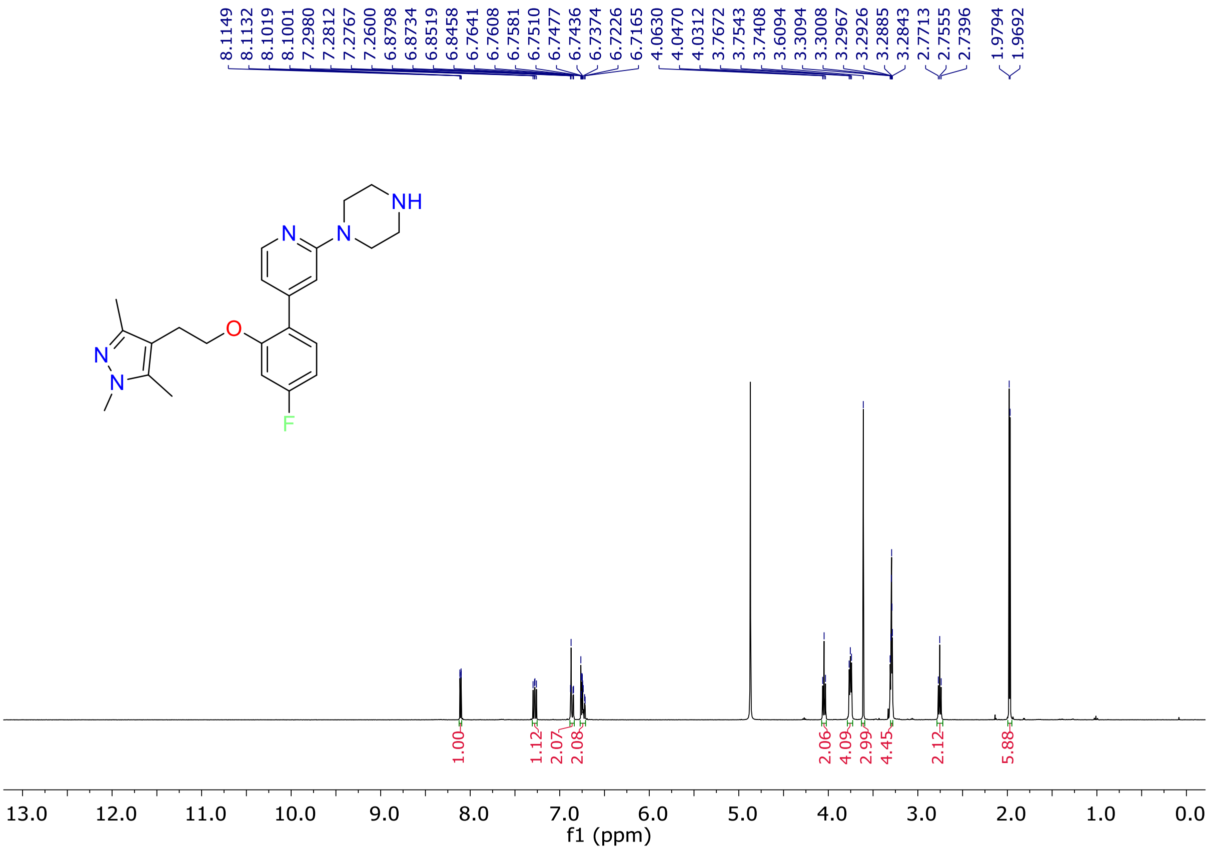


**Fig. S10.** [1]H-NMR spectrum of compound **12e** ($CD_3OD$, 400 MHz).

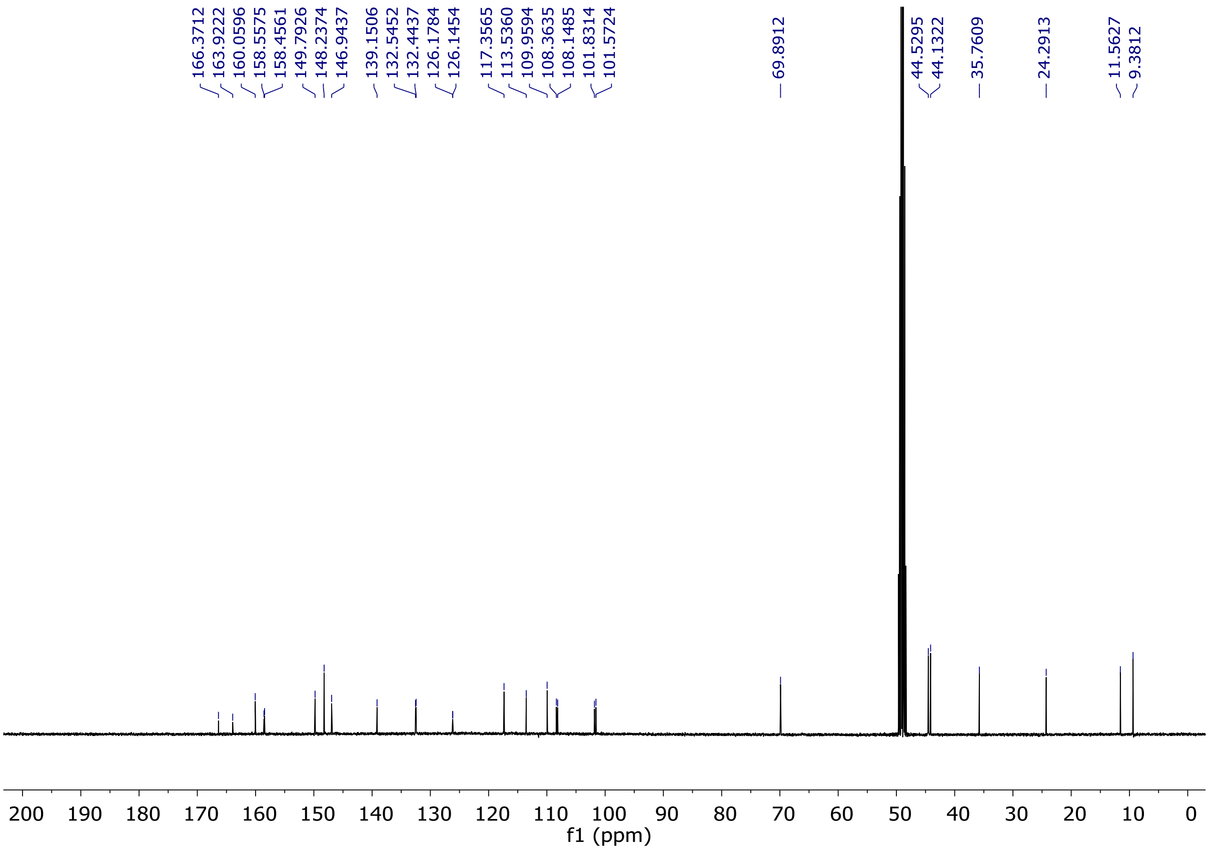


**Fig. S11.** [13]C-NMR spectrum of compound **12e** ($CD_3OD$, 100 MHz).

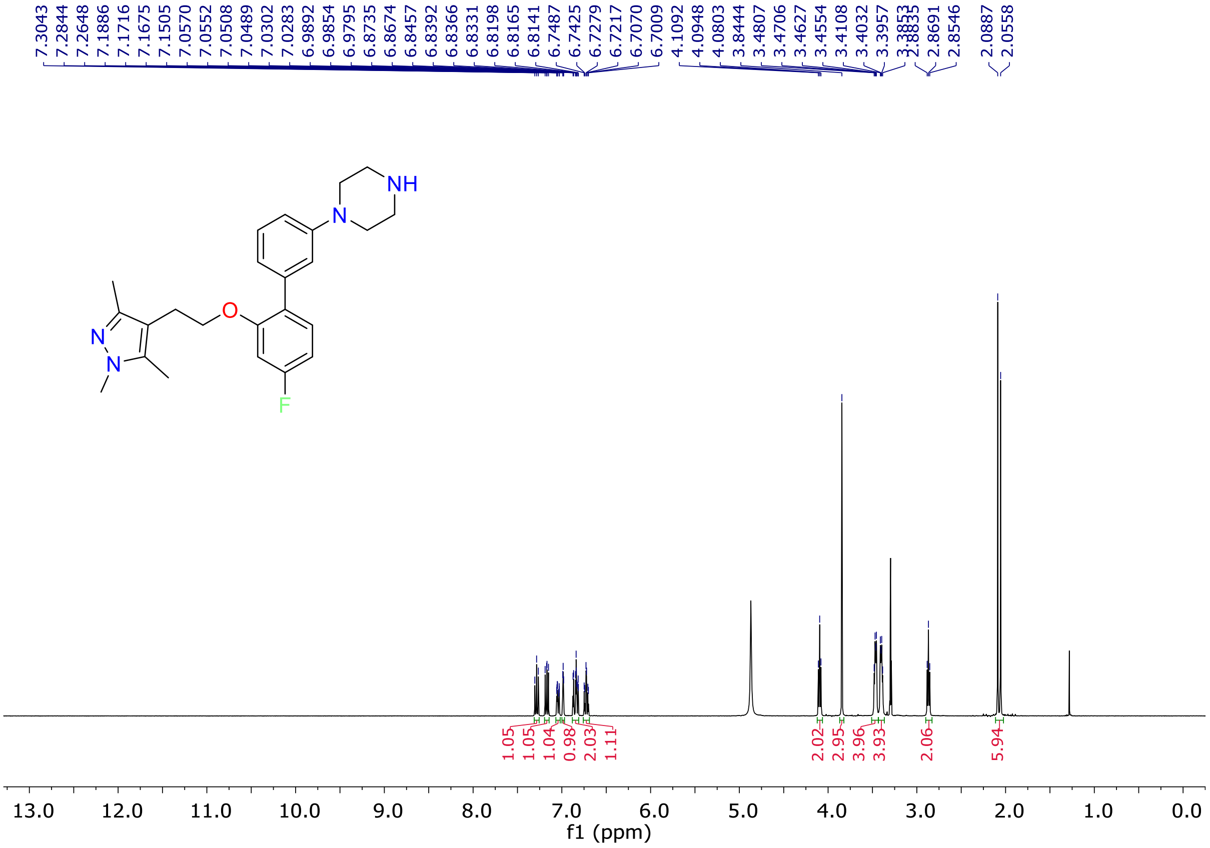


**Fig. S12.** [1]H-NMR spectrum of compound **12f** ($CD_3OD$, 400 MHz).

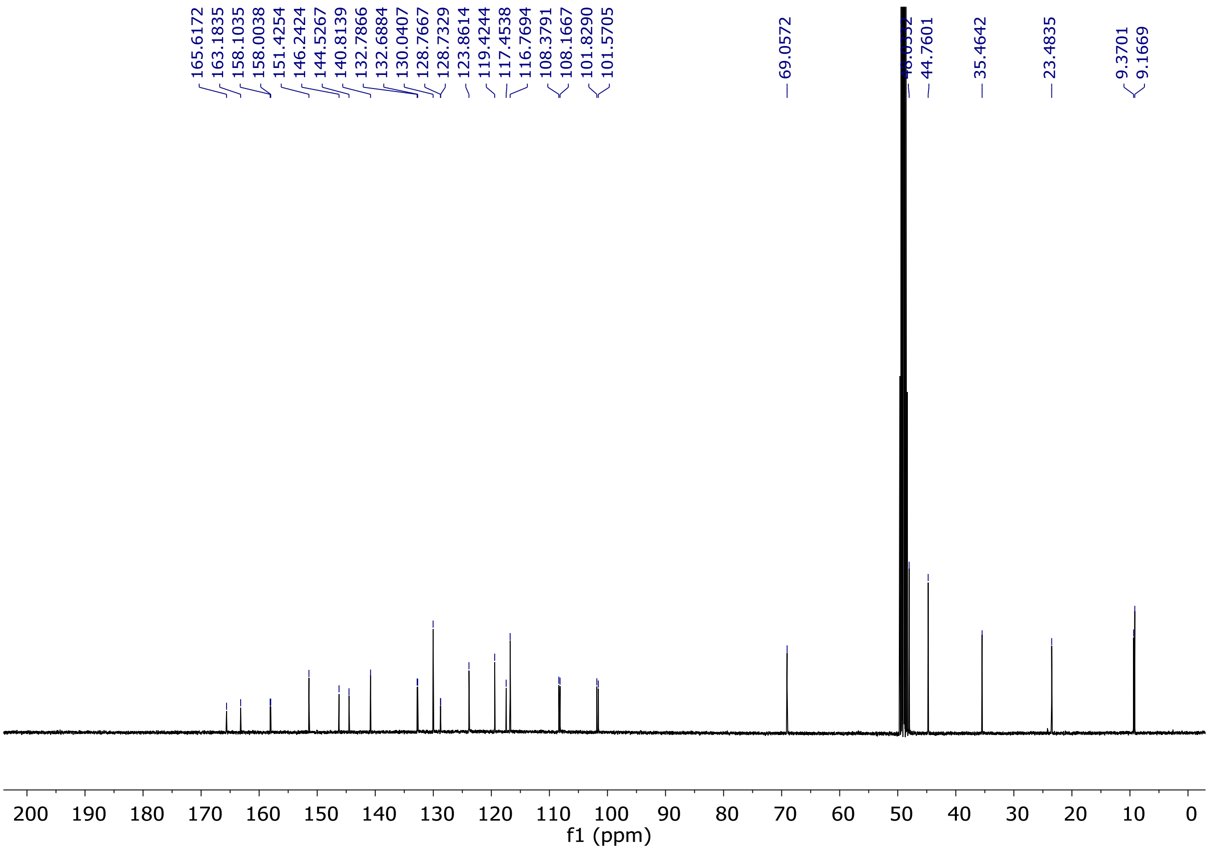


**Fig. S13.** [13]C-NMR spectrum of compound **12f** ($CD_3OD$, 100 MHz).

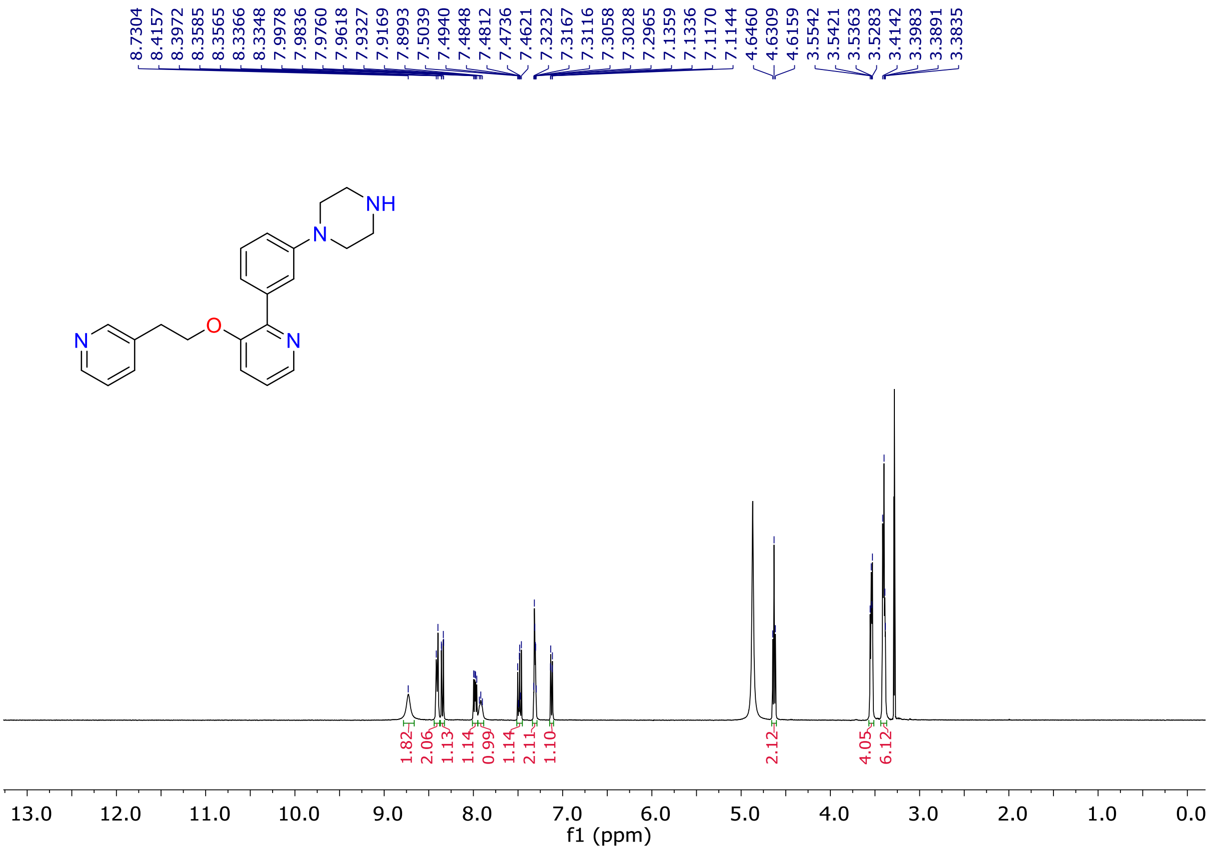


**Fig. S14.** [1]H-NMR spectrum of compound **12g** ($CD_3OD$, 400 MHz).

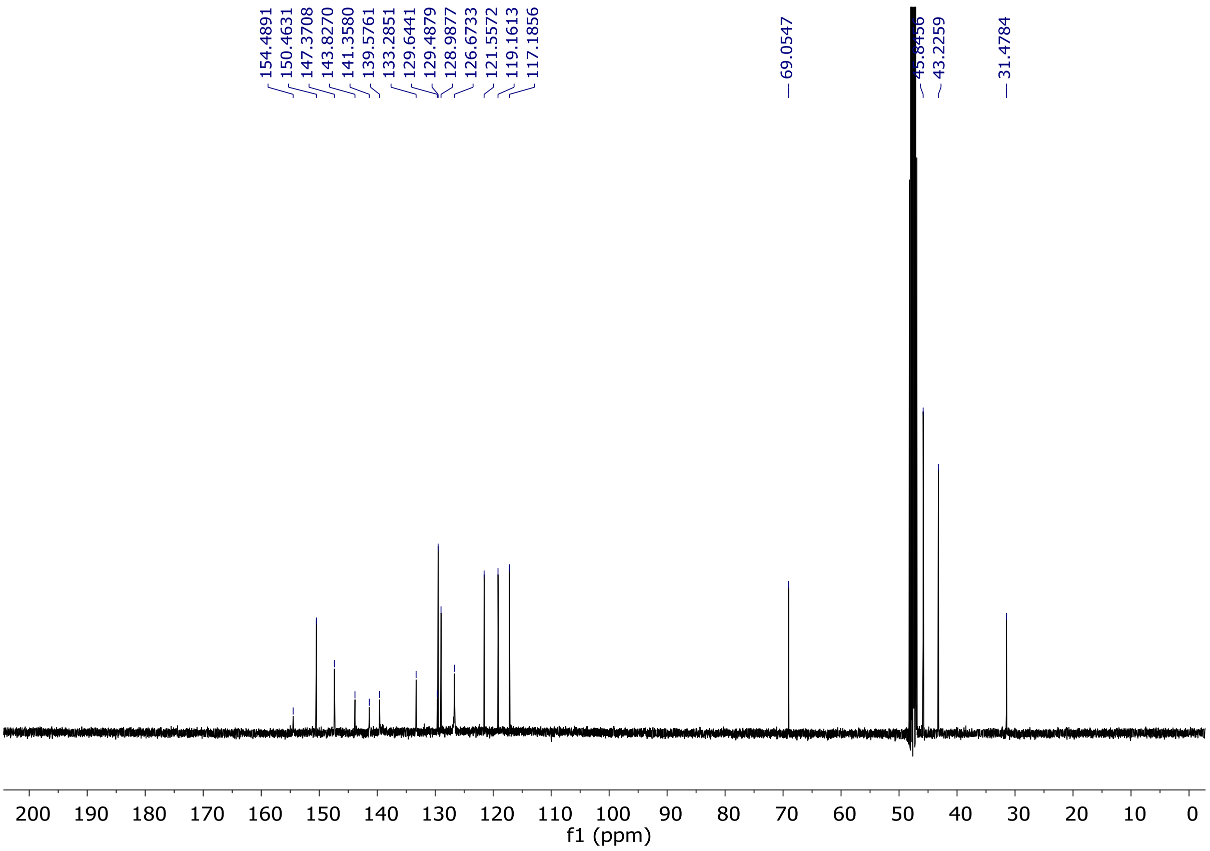


**Fig. S15.** [13]C-NMR spectrum of compound **12g** ($CD_3OD$, 100 MHz).

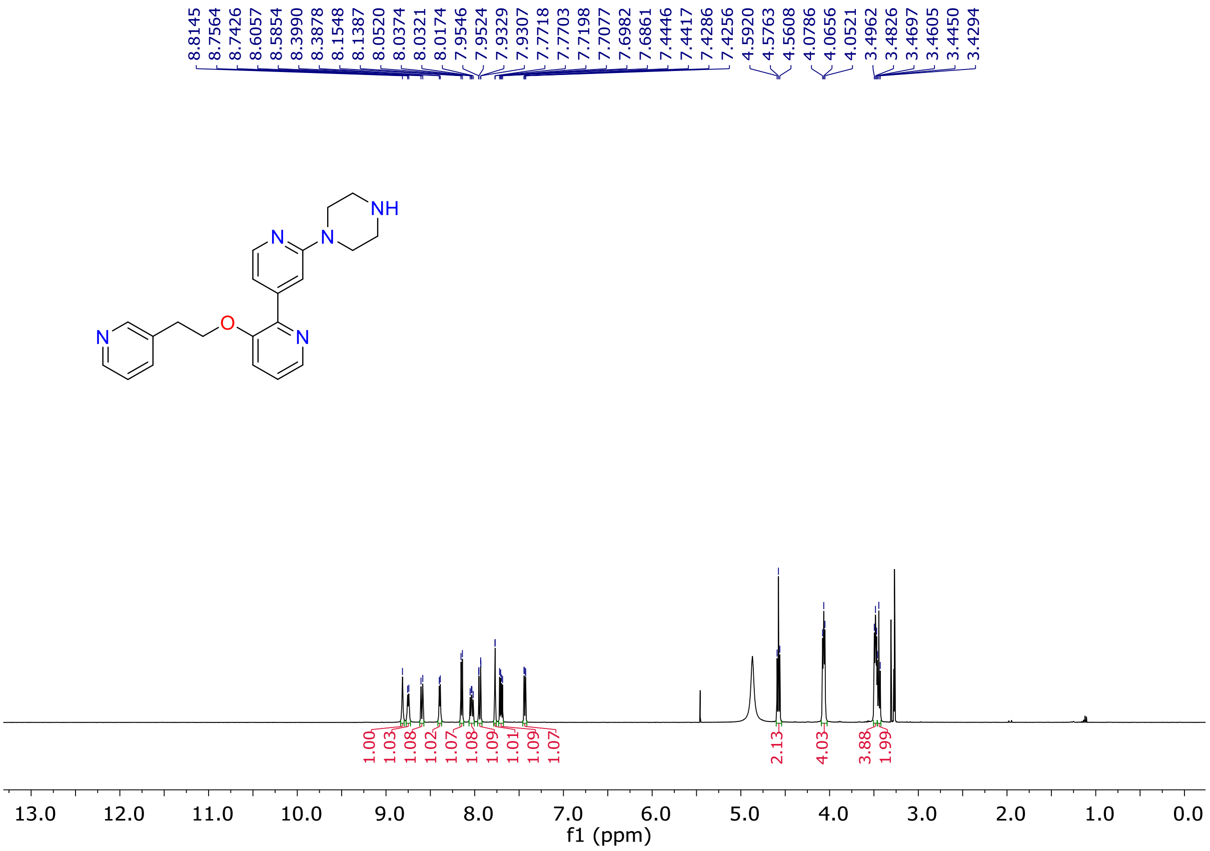


**Fig. S16.** [1]H-NMR spectrum of compound **12h** ($CD_3OD$, 400 MHz).

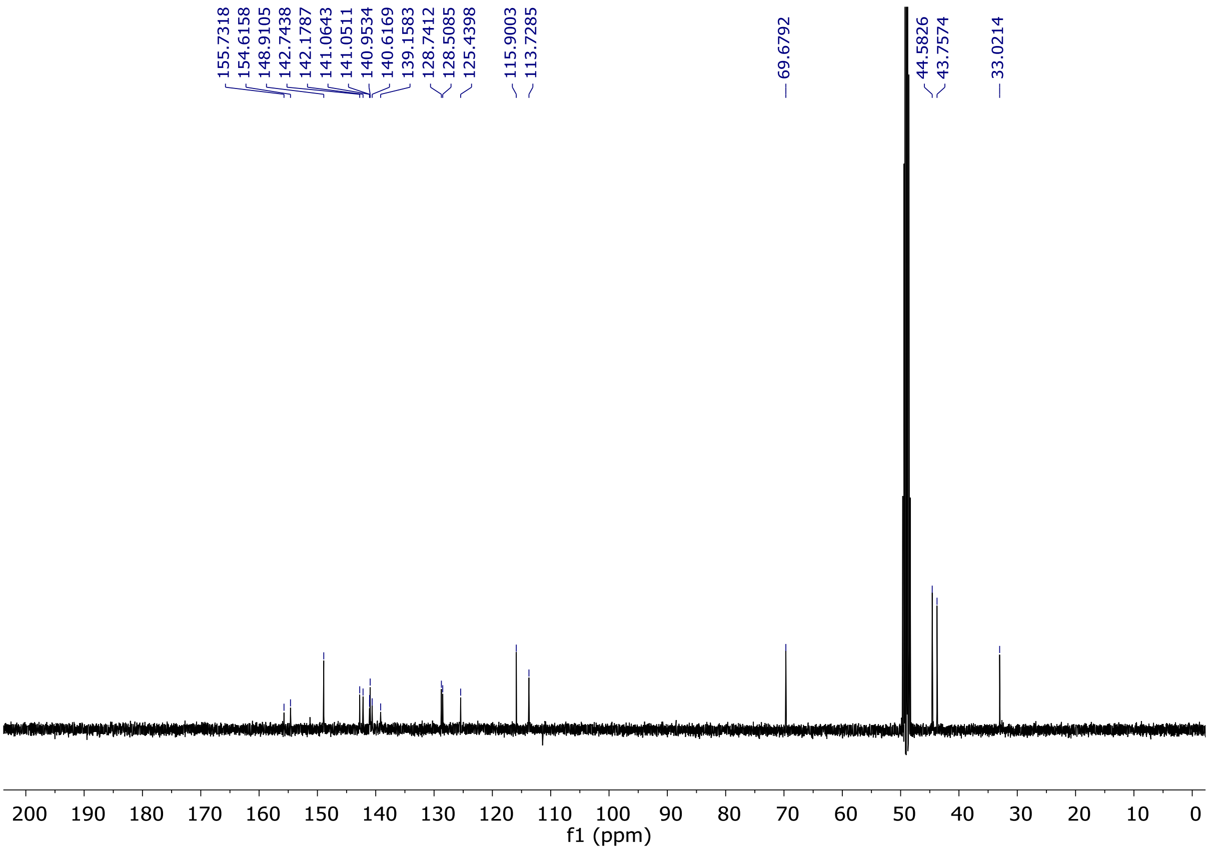


**Fig. S17.** $^{13}$C-NMR spectrum of compound **12h** ($CD_3OD$, 100 MHz).

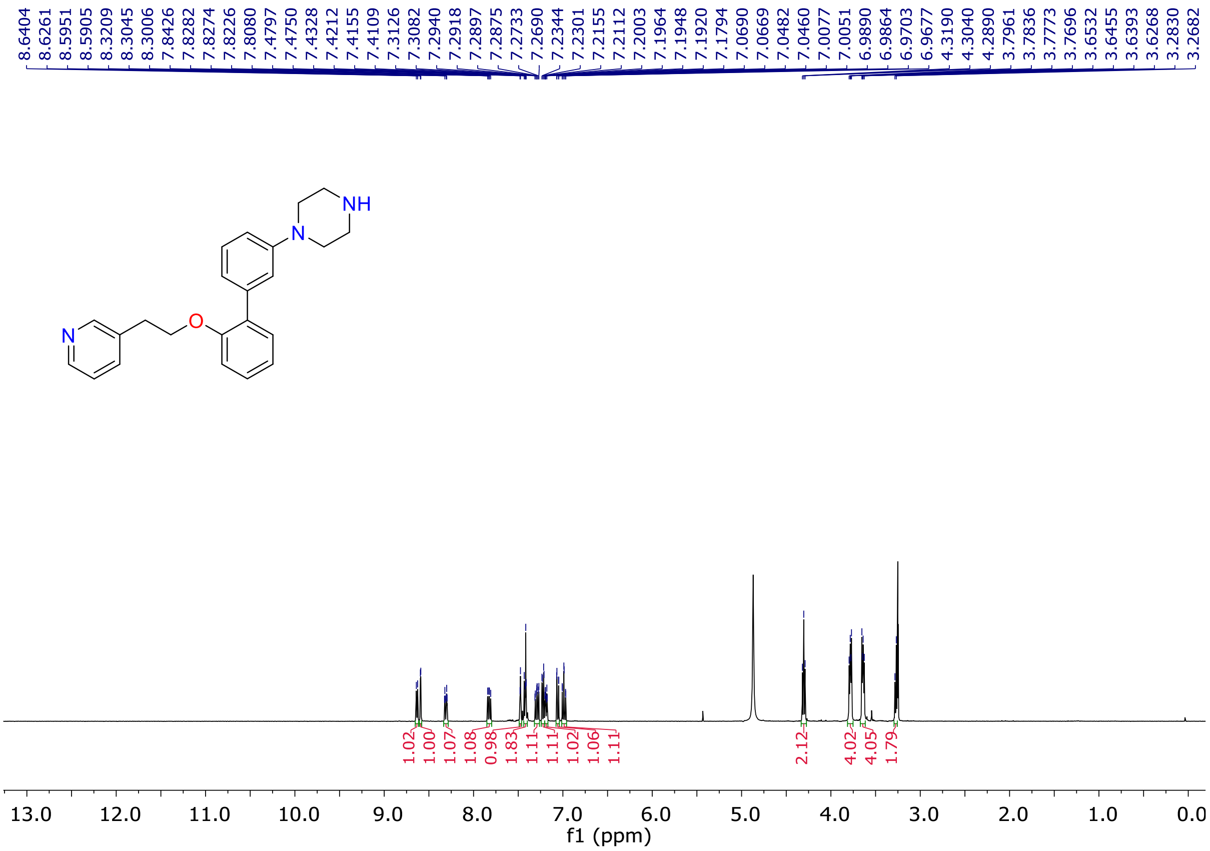


**Fig. S18.** $^{1}$H-NMR spectrum of compound **12i** ($CD_3OD$, 400 MHz).

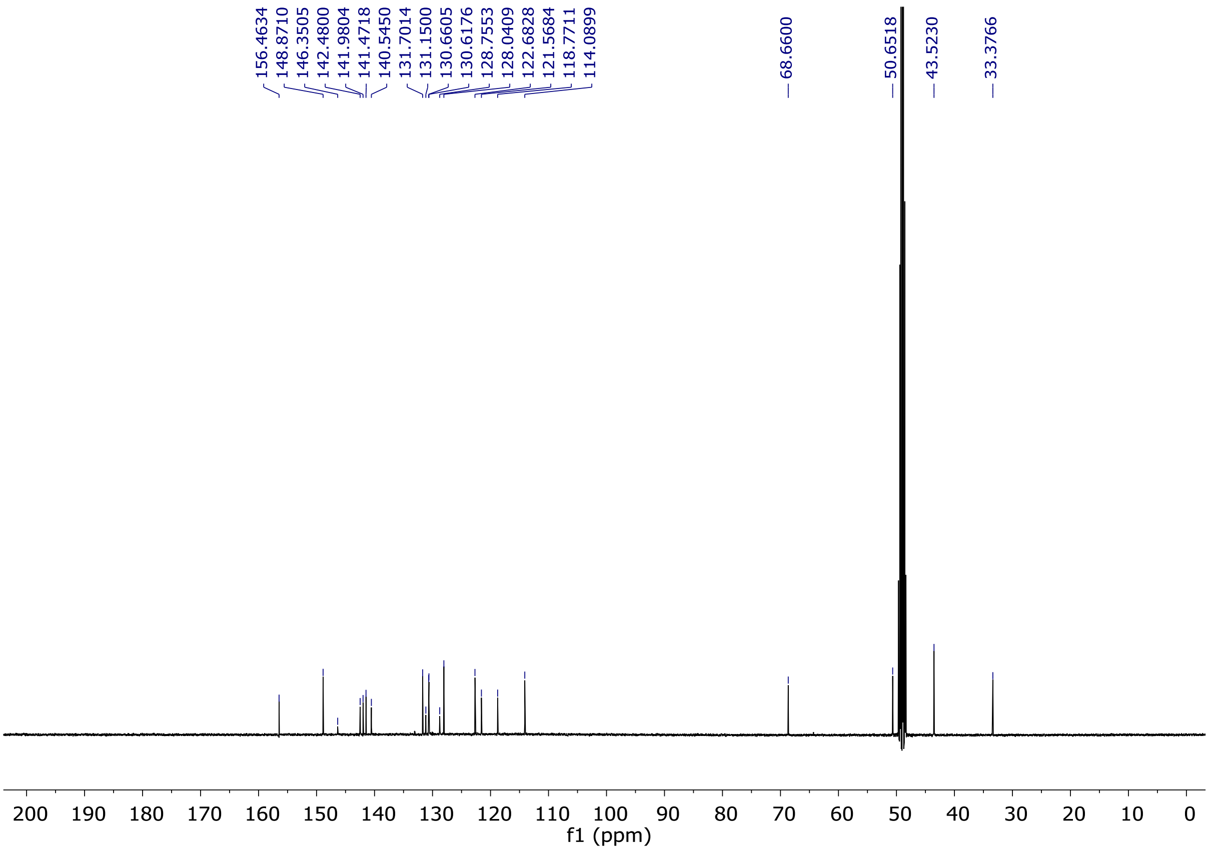

**Fig. S19.** $^{13}$C-NMR spectrum of compound **12i** ($CD_3OD$, 100 MHz).

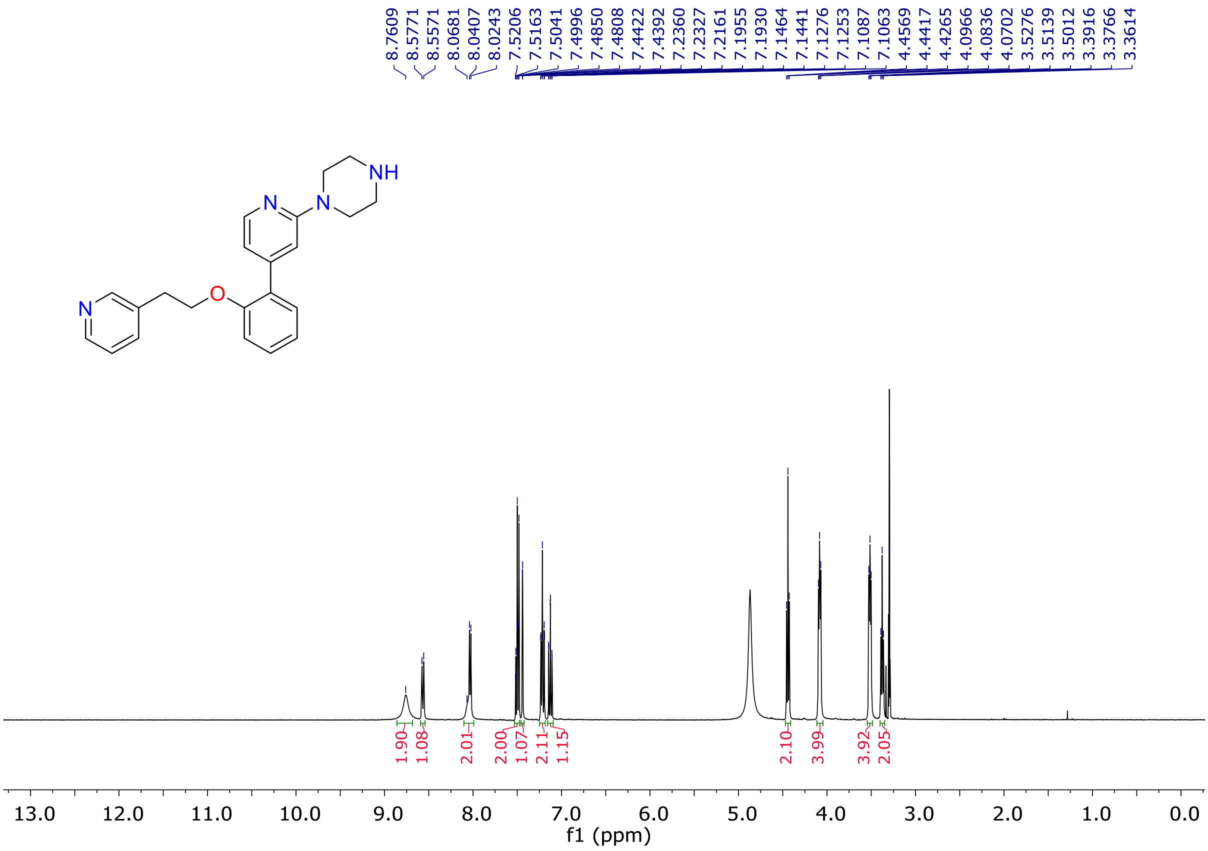

**Fig. S20.** $^{1}$H-NMR spectrum of compound **12j** ($CD_3OD$, 400 MHz).

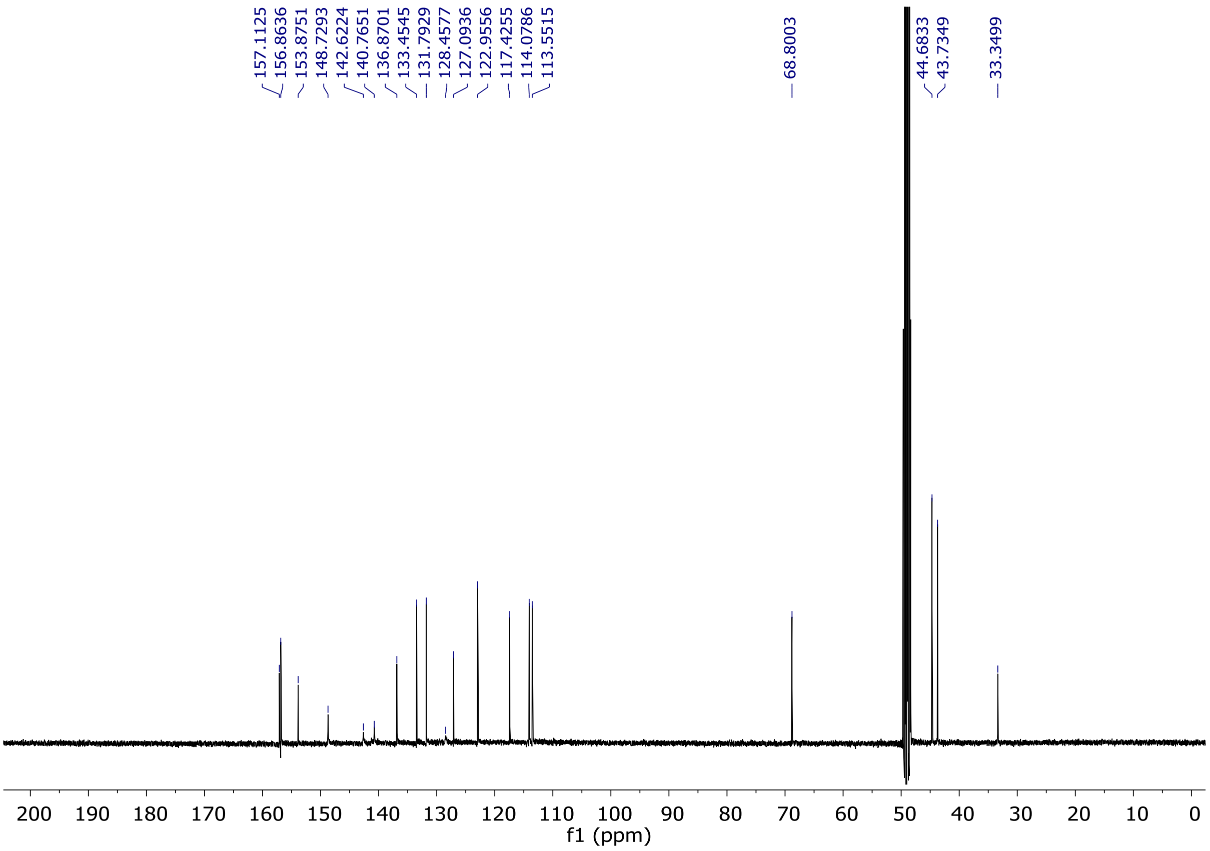


**Fig. S21.** $^{13}$C-NMR spectrum of compound **12j** ($CD_3OD$, 100 MHz).

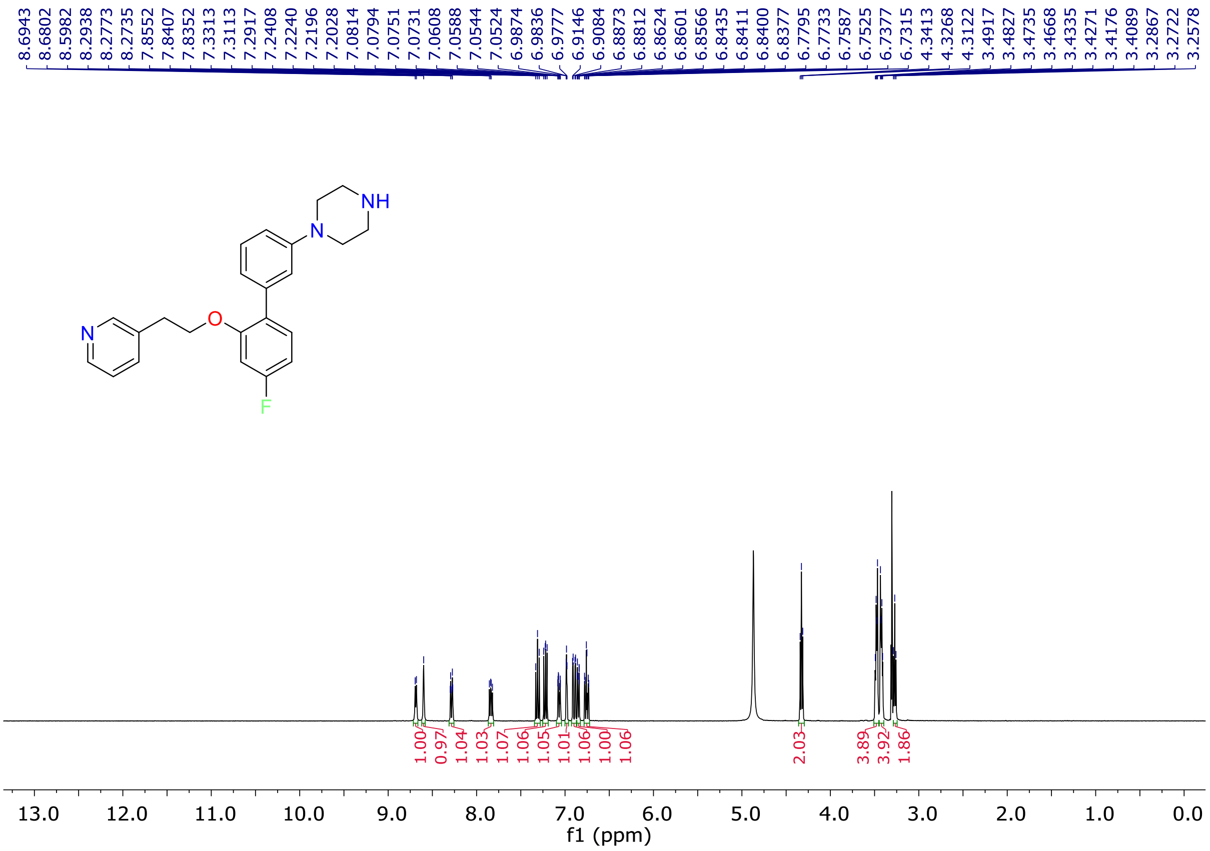


**Fig. S22.** $^{1}$H-NMR spectrum of compound **12k** ($CD_3OD$, 400 MHz).

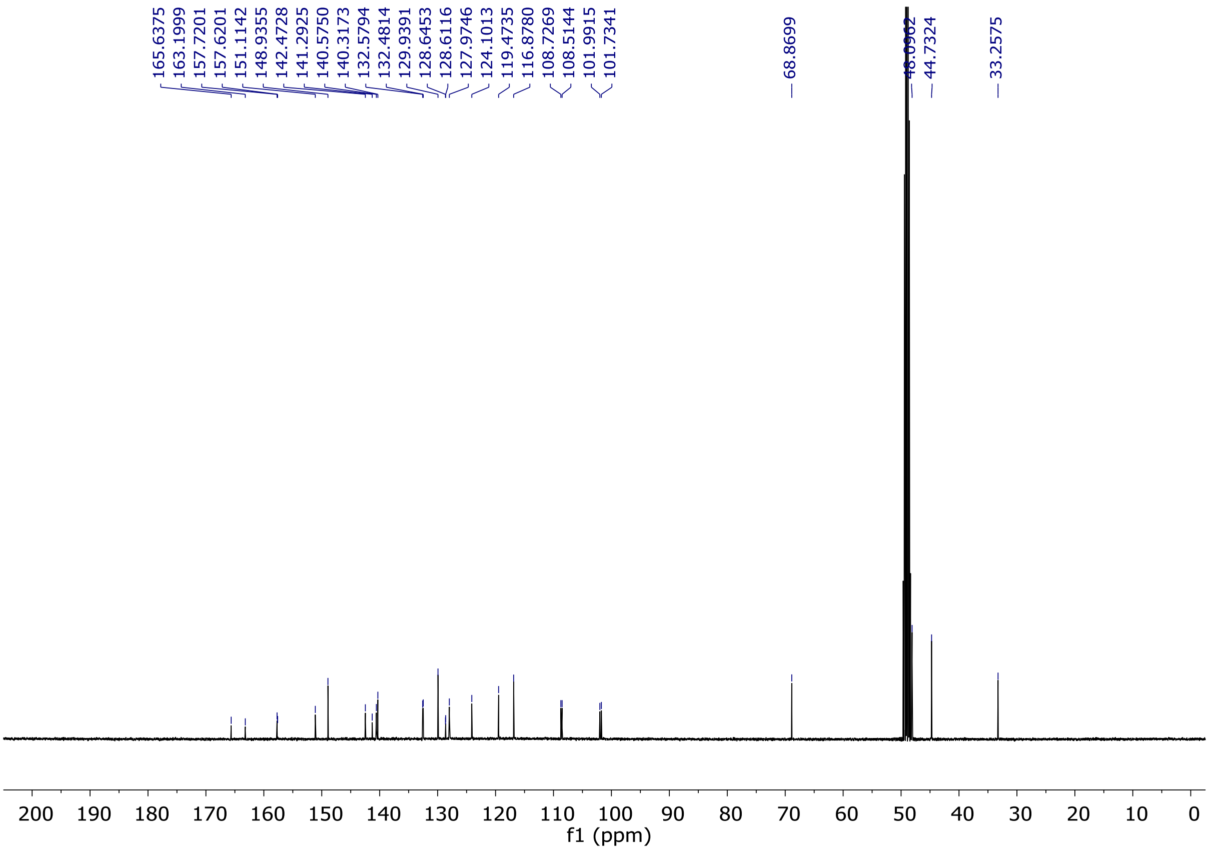


**Fig. S23.** $^{13}$C-NMR spectrum of compound **12k** ($CD_3OD$, 100 MHz).

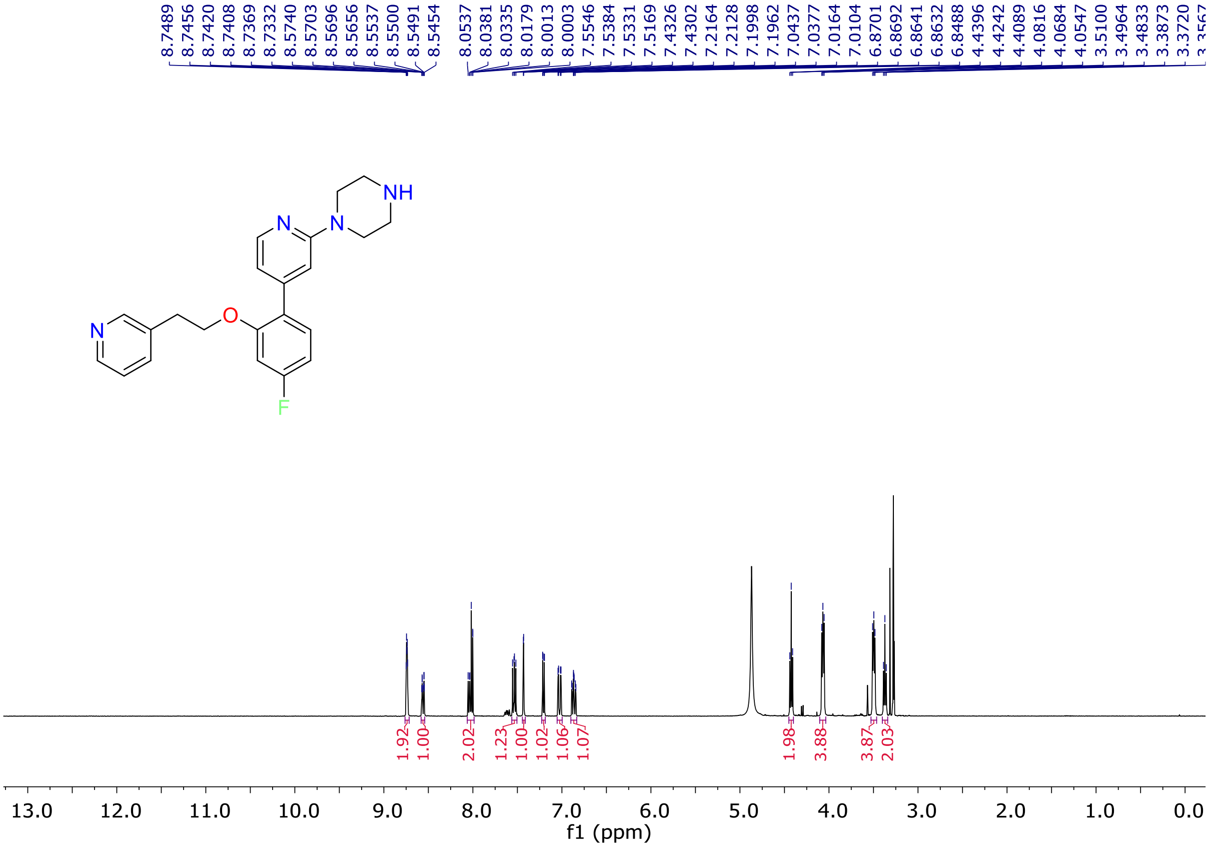


**Fig. S24.** $^{1}$H-NMR spectrum of compound **12l** ($CD_3OD$, 400 MHz).

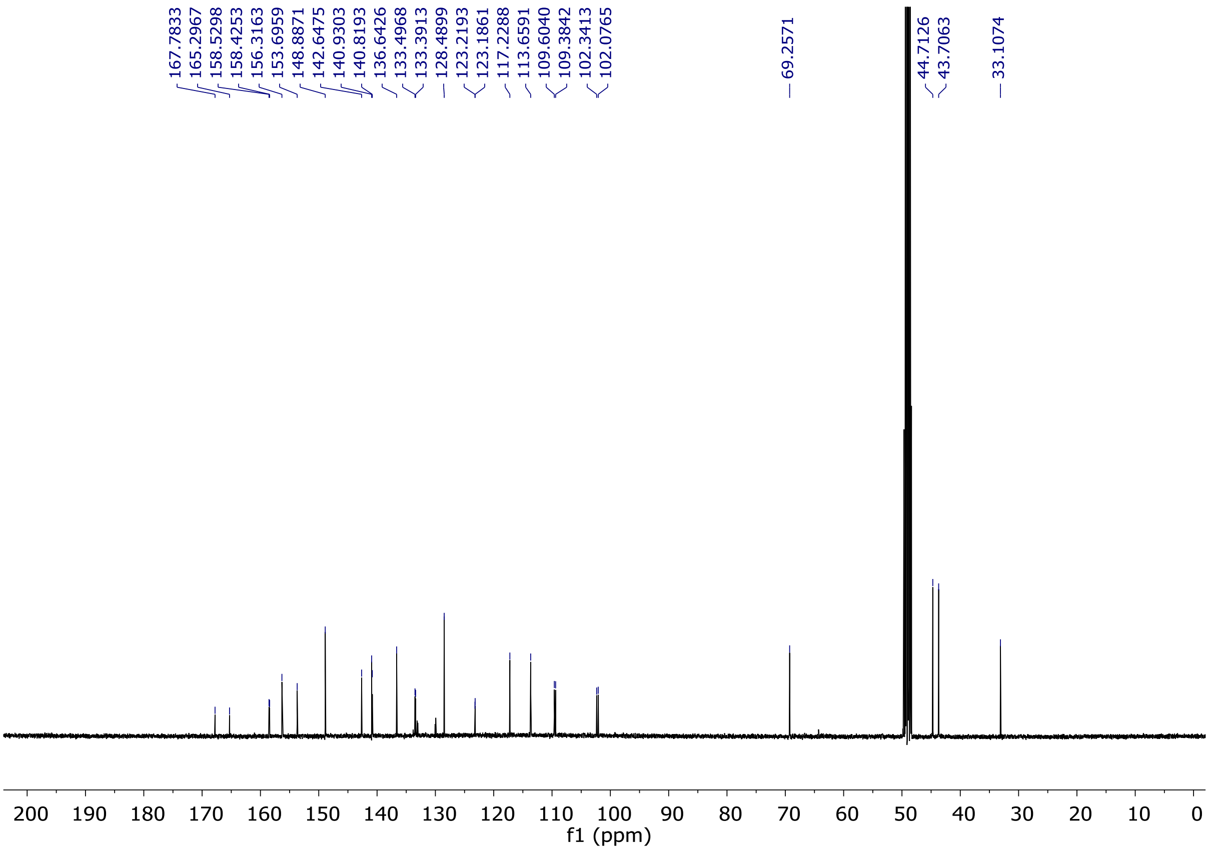


**Fig. S25.** $^{13}$C-NMR spectrum of compound **12l** ($CD_3OD$, 100 MHz).

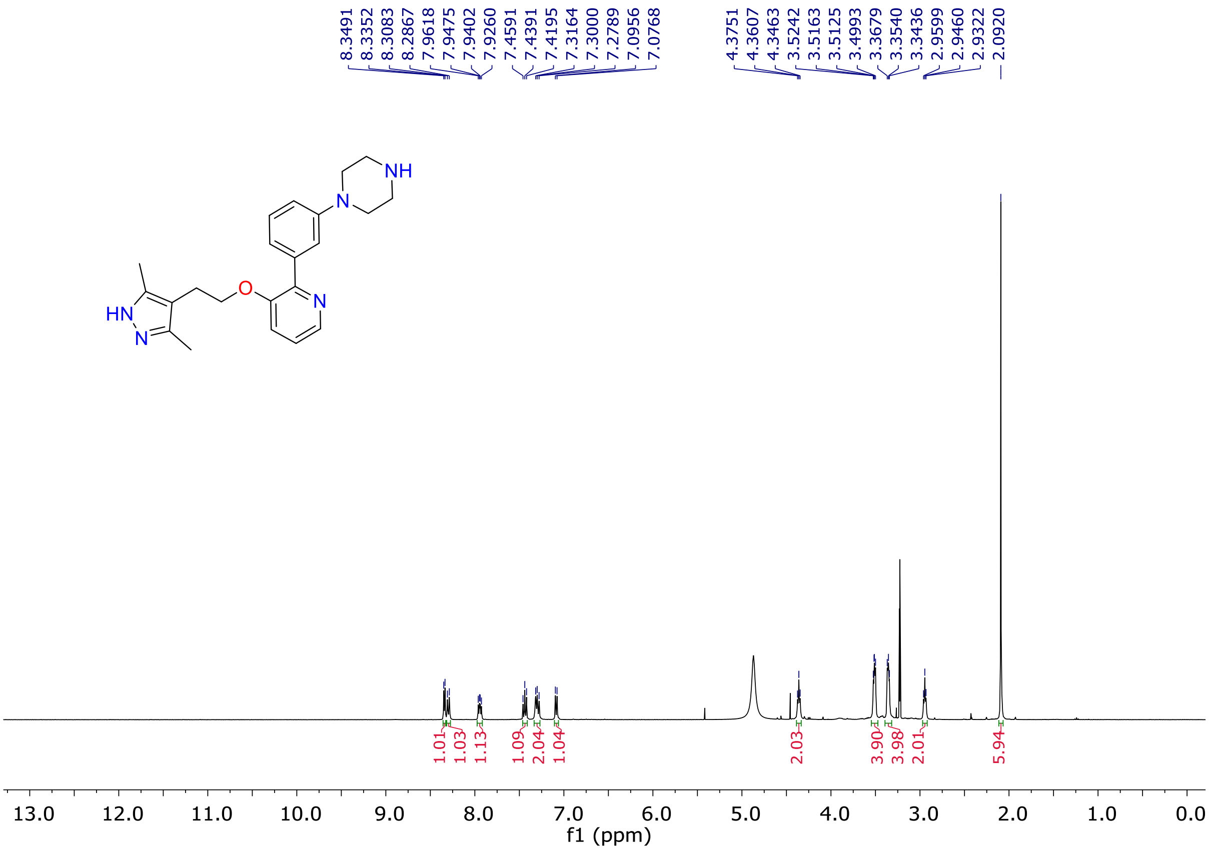


**Fig. S26.** $^{1}$H-NMR spectrum of compound **12m** ($CD_3OD$, 400 MHz).

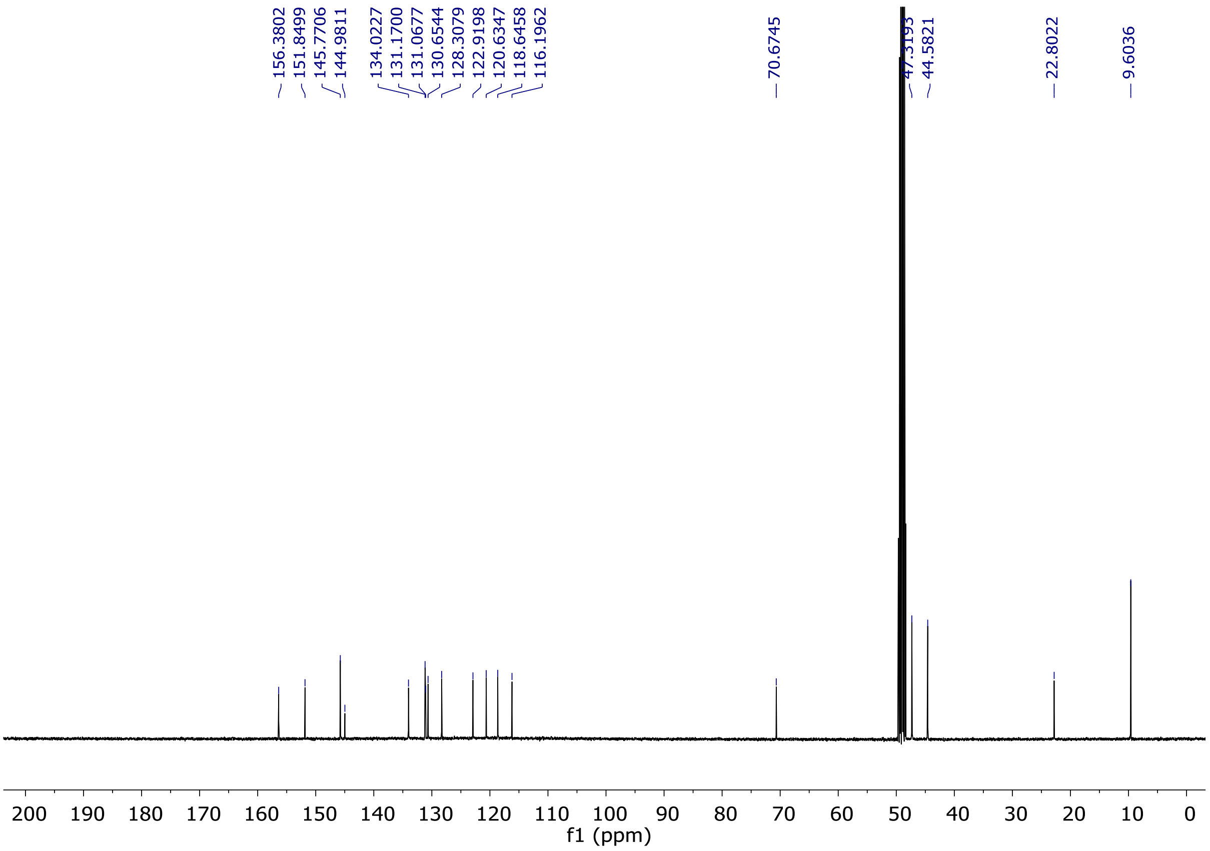


**Fig. S27.** $^{13}$C-NMR spectrum of compound **12m** ($CD_3OD$, 100 MHz).

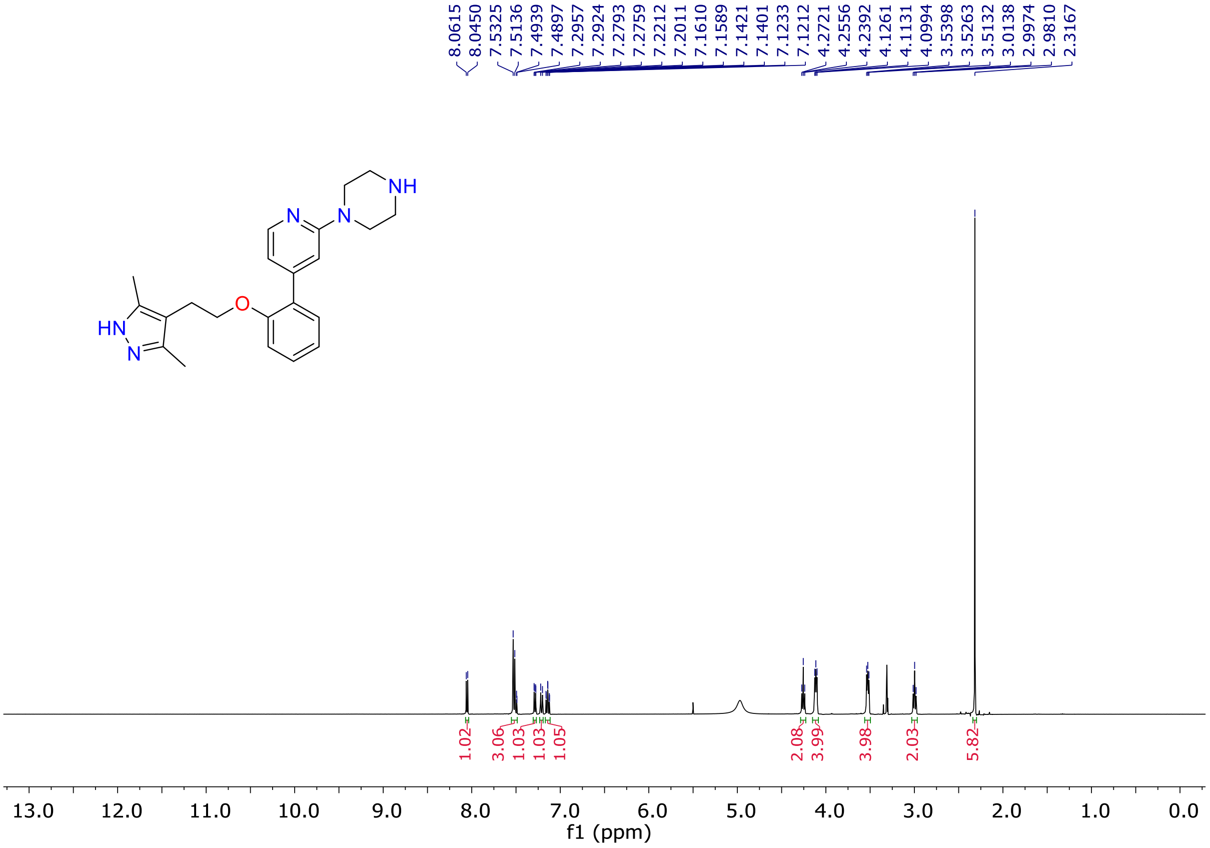


**Fig. S28.** $^{1}$H-NMR spectrum of compound **12n** ($CD_3OD$, 400 MHz).

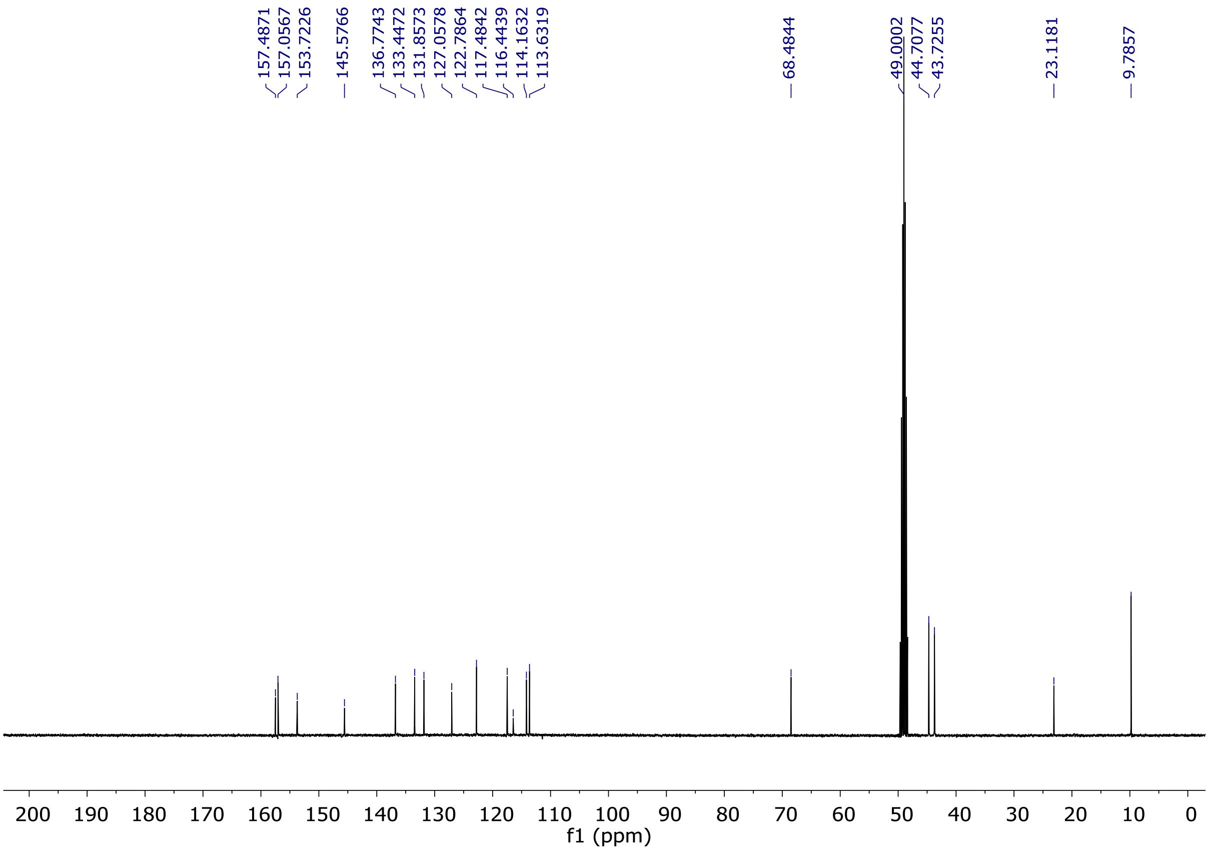


**Fig. S29.** [13]C-NMR spectrum of compound **12n** ($CD_3OD$, 100 MHz).

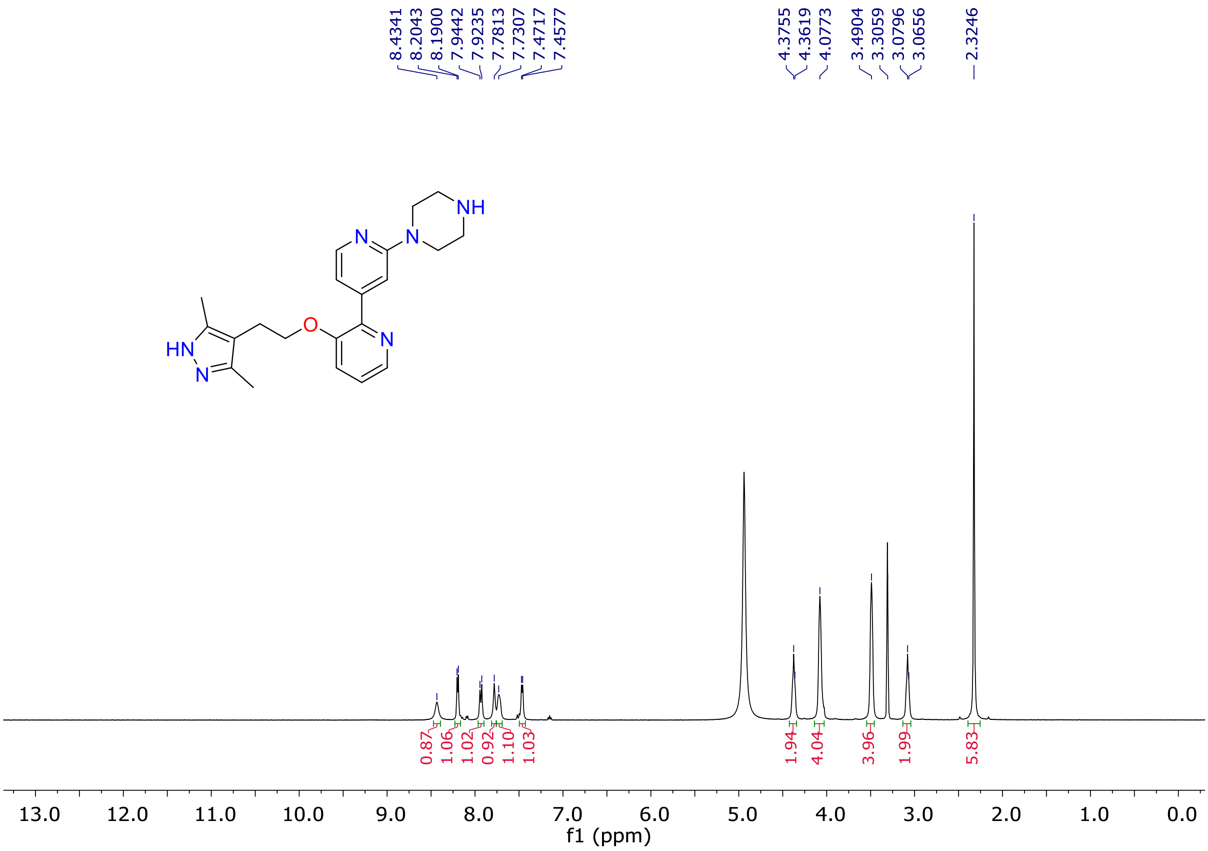


**Fig. S30.** [1]H-NMR spectrum of compound **12o** ($CD_3OD$, 400 MHz).

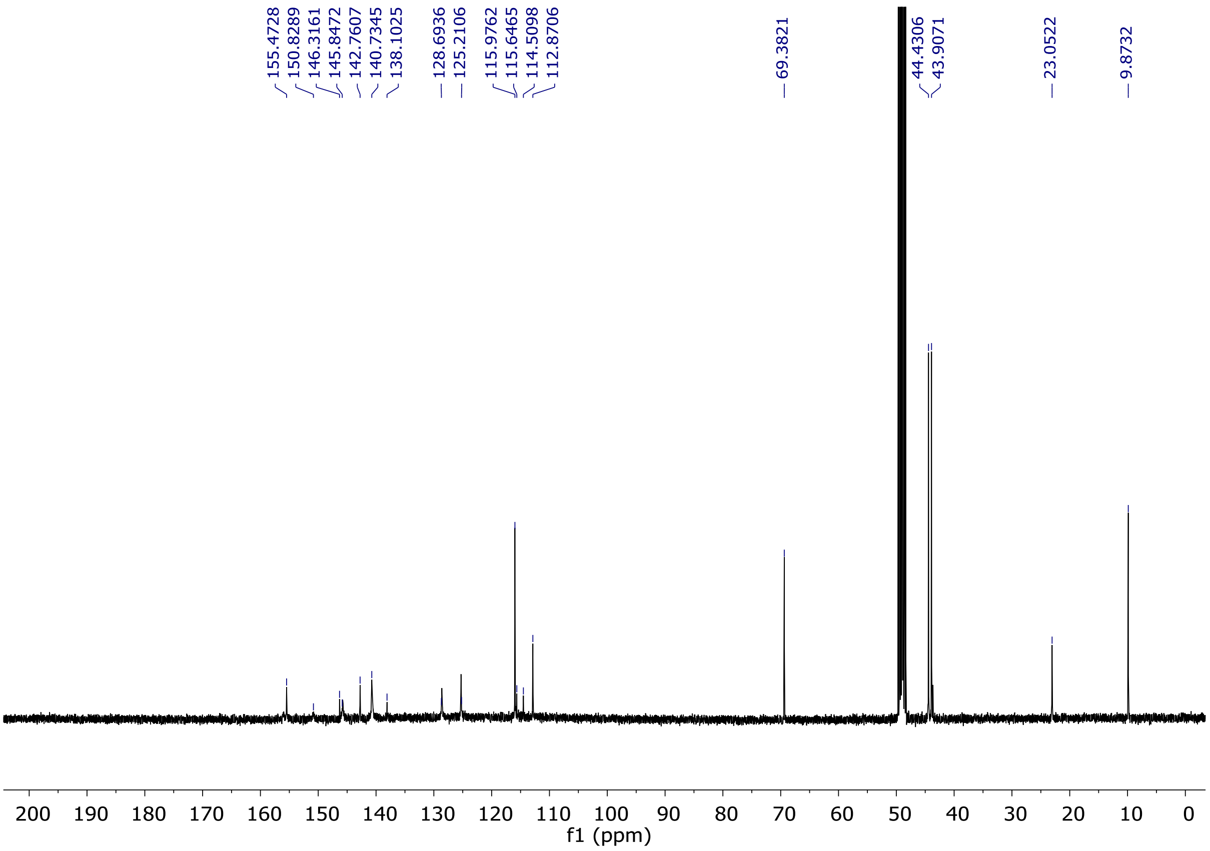


**Fig. S31.** $^{13}$C-NMR spectrum of compound **12o** ($CD_3OD$, 100 MHz).

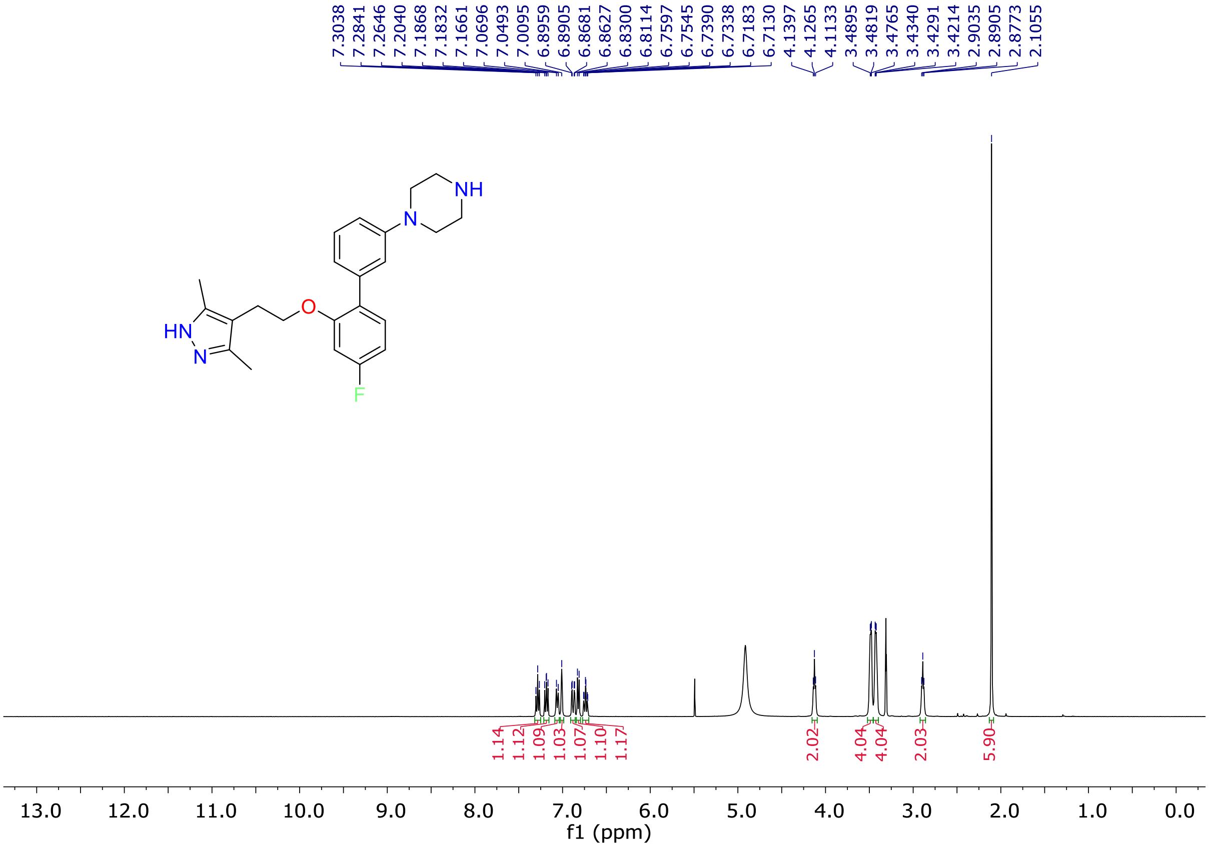


**Fig. S32.** $^{1}$H-NMR spectrum of compound **12p** ($CD_3OD$, 400 MHz).

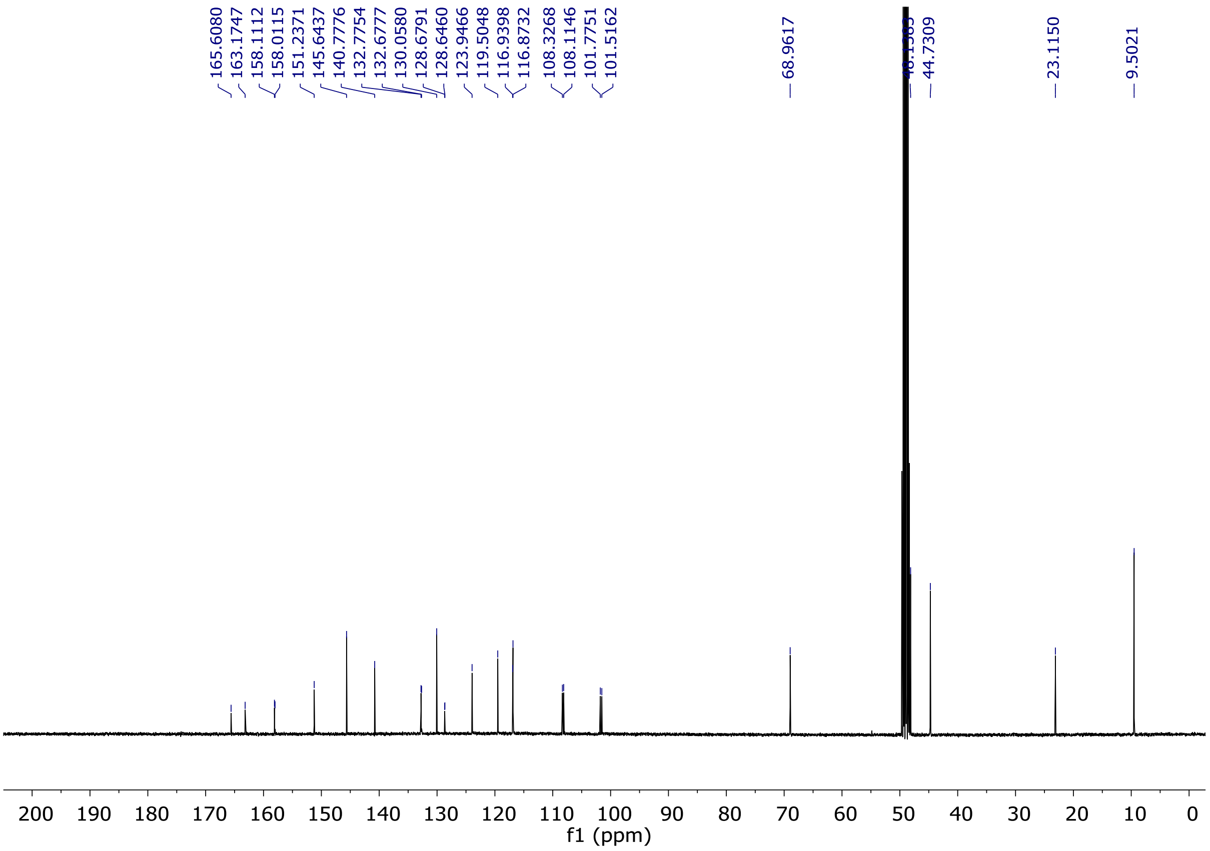


**Fig. S33.** $^{13}$C-NMR spectrum of compound **12p** ($CD_3OD$, 100 MHz).

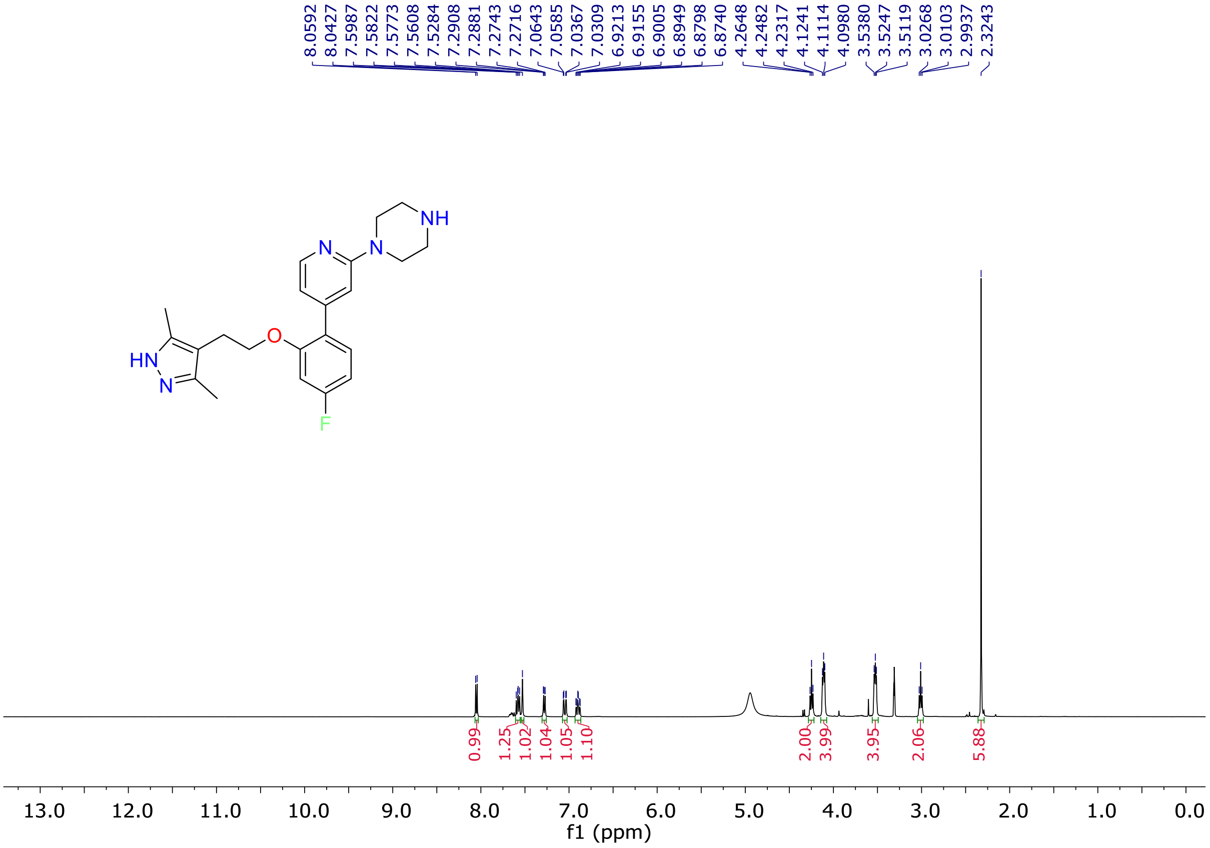


**Fig. S34.** $^{1}$H-NMR spectrum of compound **12q** ($CD_3OD$, 400 MHz).

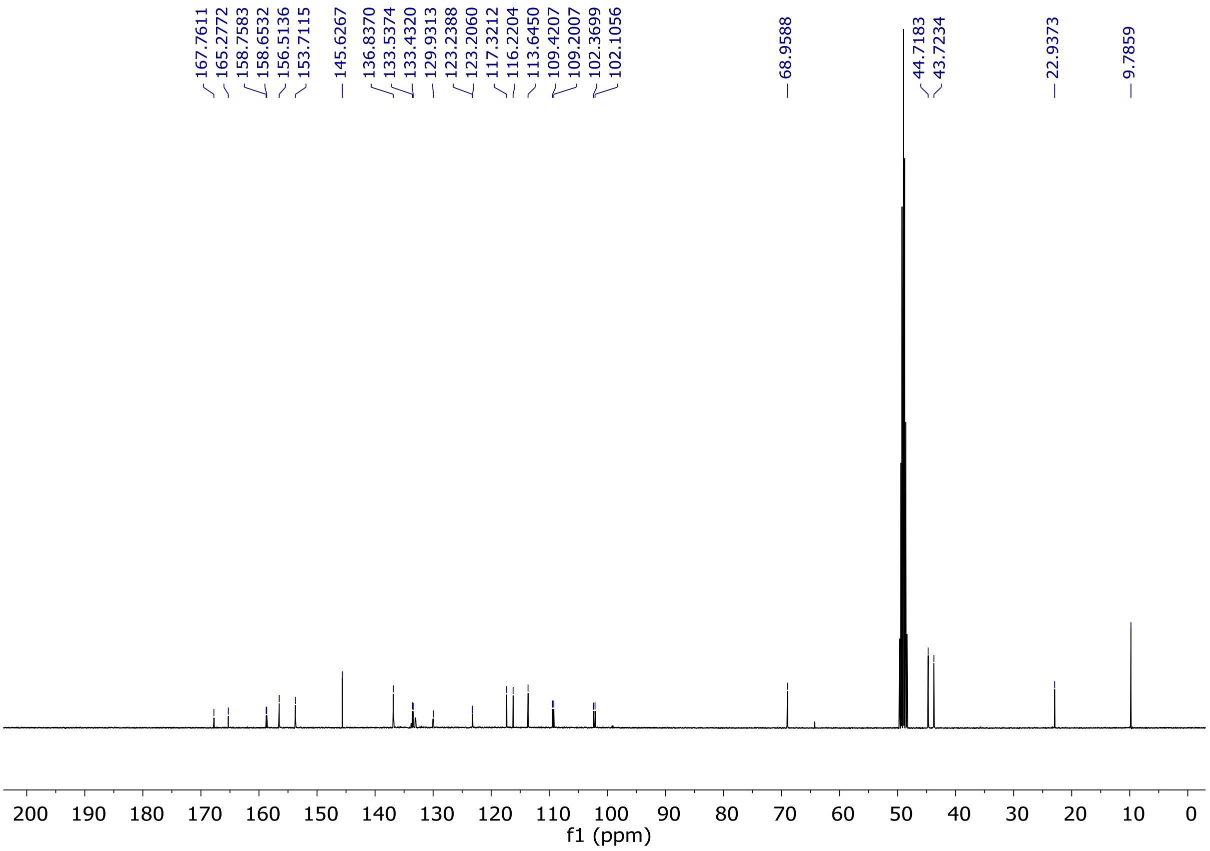


**Fig. S35.** $^{13}$C-NMR spectrum of compound **12q** ($CD_3OD$, 100 MHz).

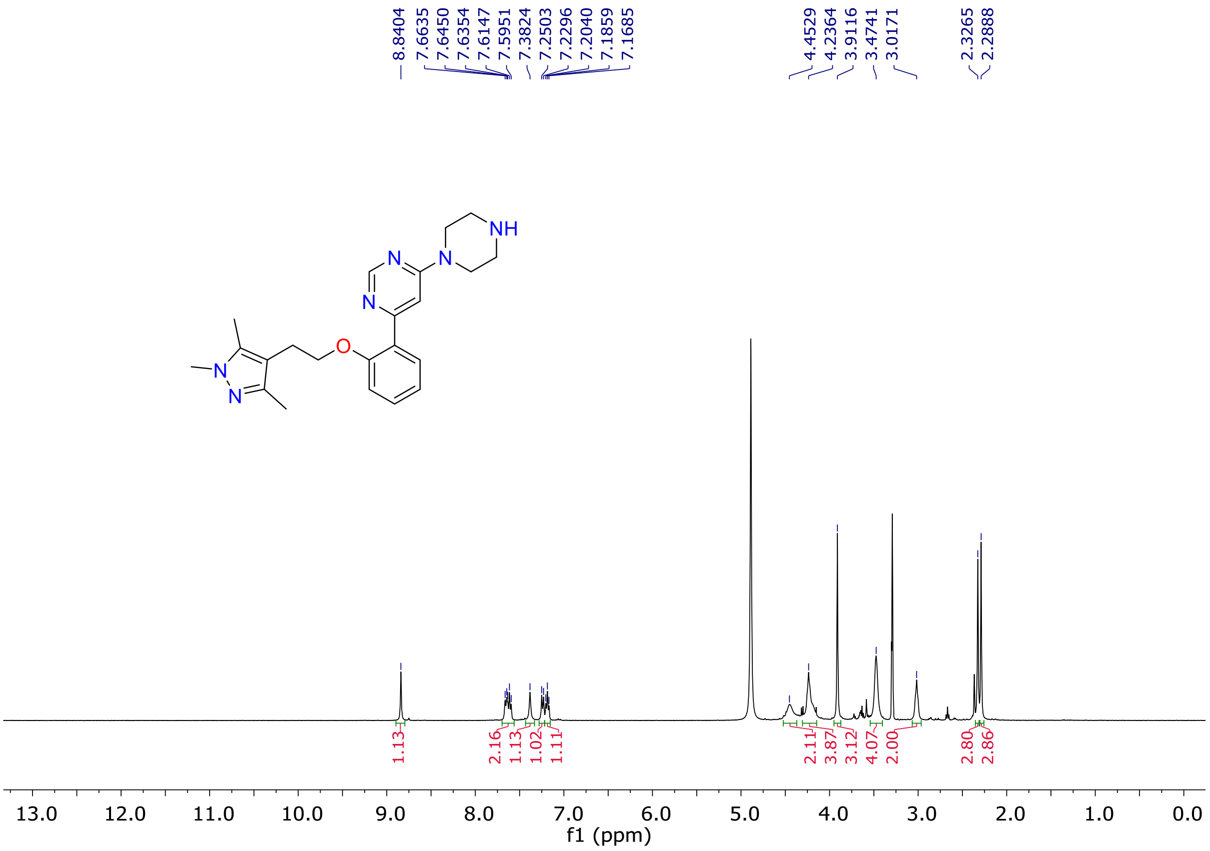


**Fig. S36.** $^{1}$H-NMR spectrum of compound **16a** ($CD_3OD$, 400 MHz).

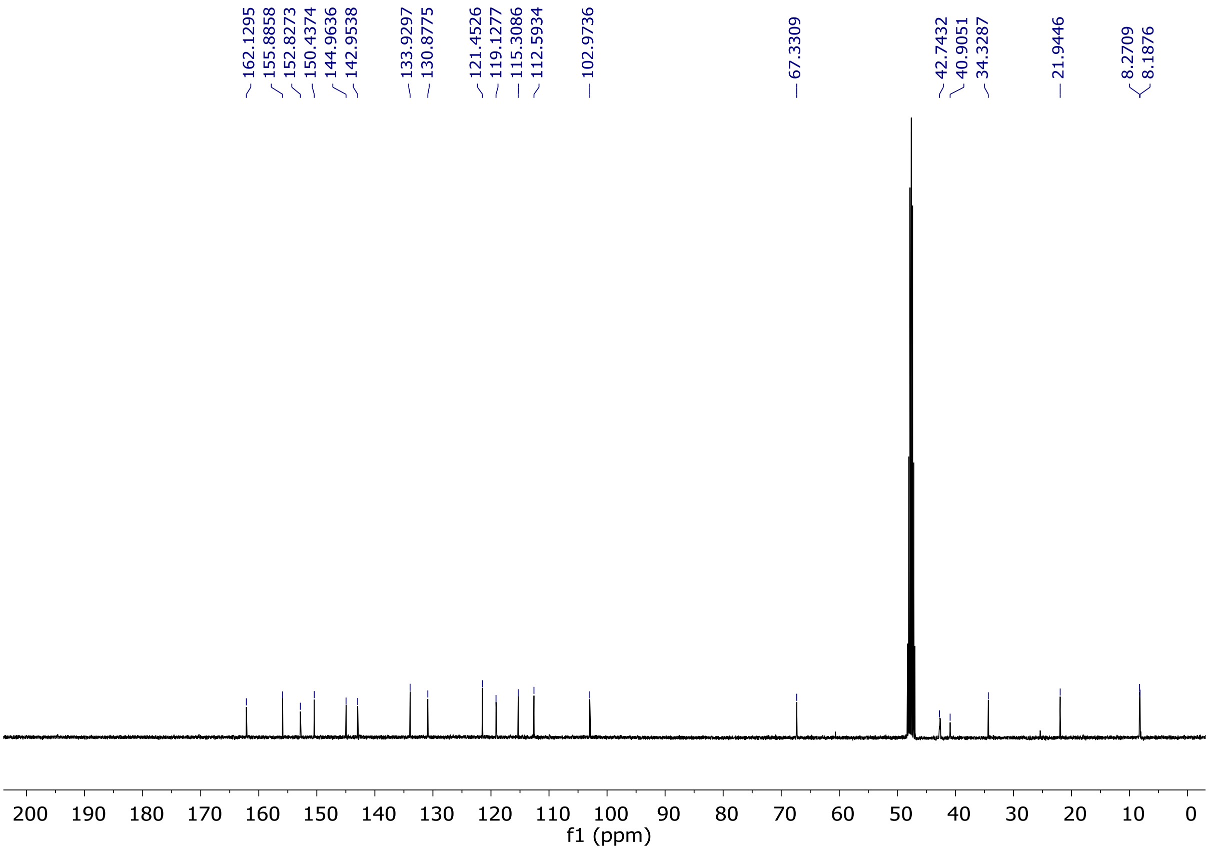


**Fig. S37.** $^{13}$C-NMR spectrum of compound **16a** ($CD_3OD$, 100 MHz).

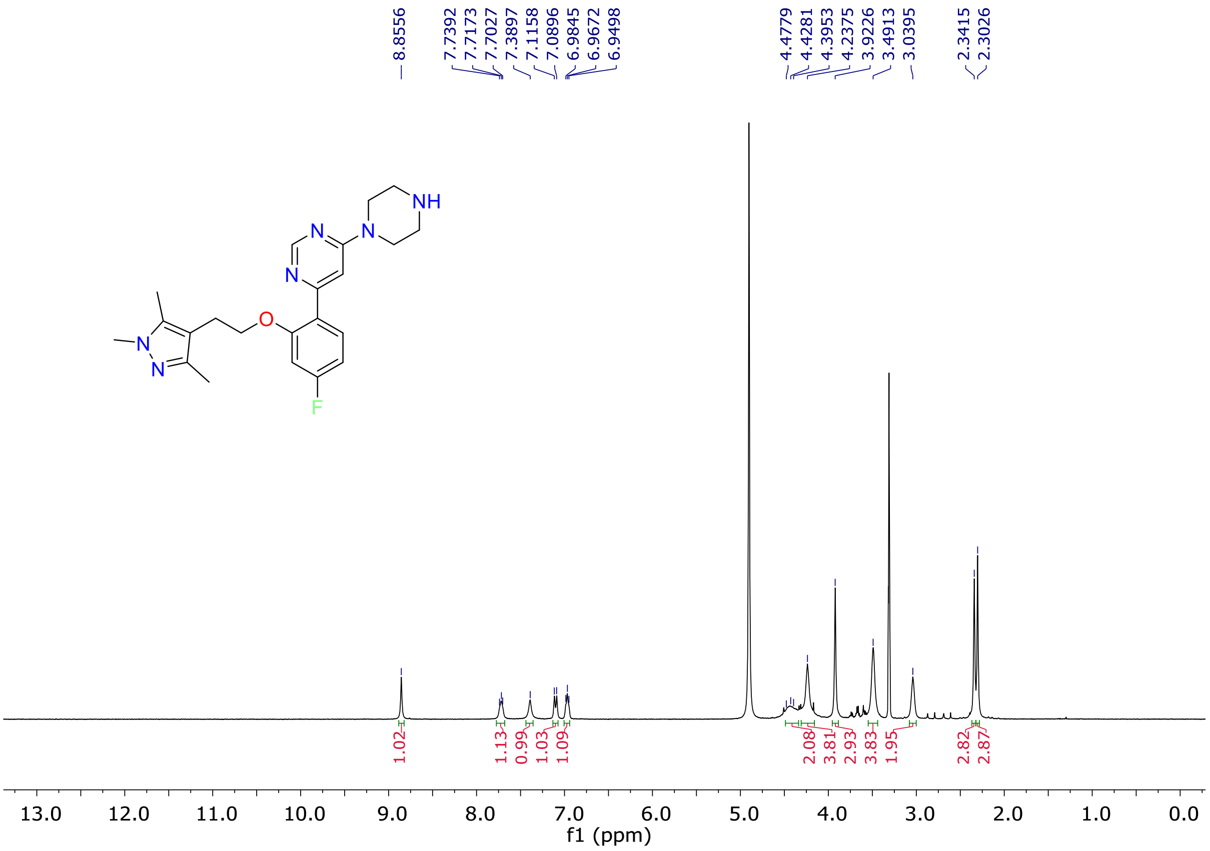


**Fig. S38.** $^{1}$H-NMR spectrum of compound **16b** ($CD_3OD$, 400 MHz).

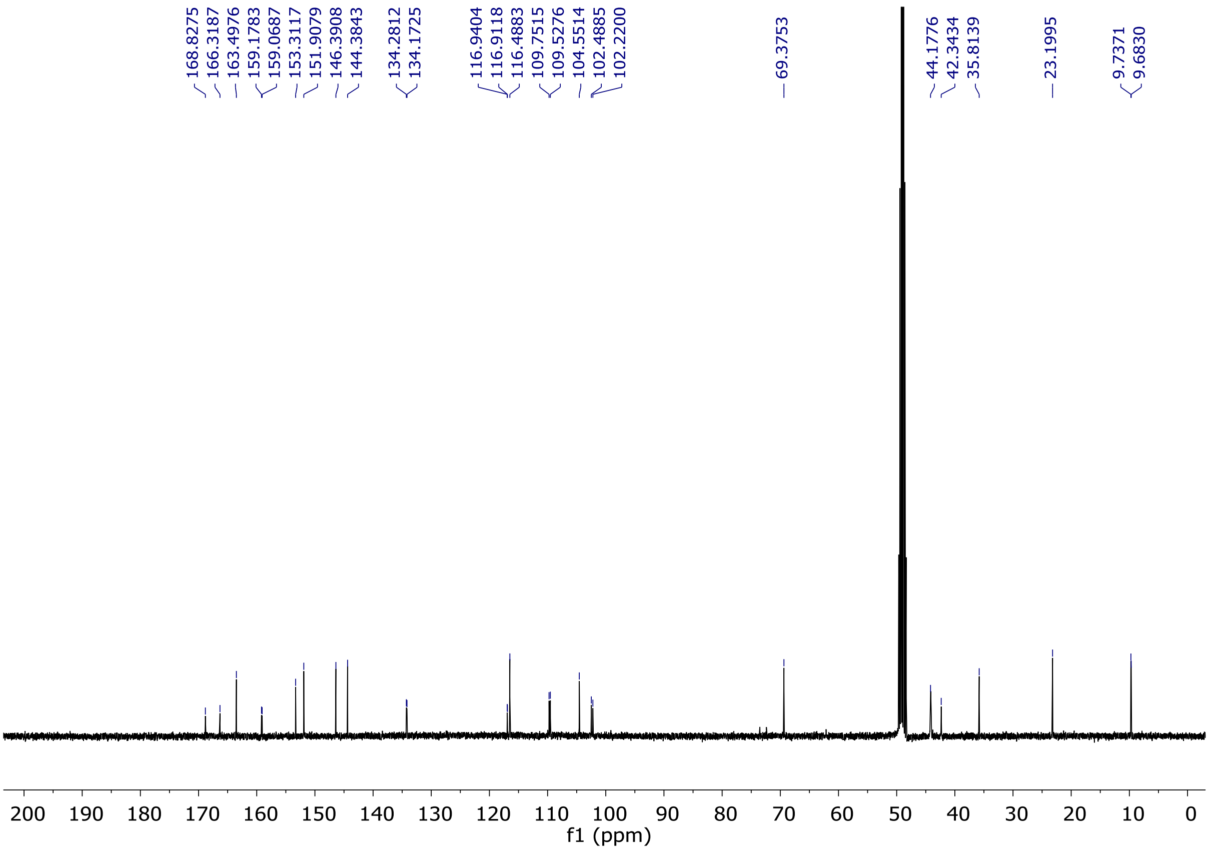


**Fig. S39.** [13]C-NMR spectrum of compound **16b** ($CD_3OD$, 100 MHz).

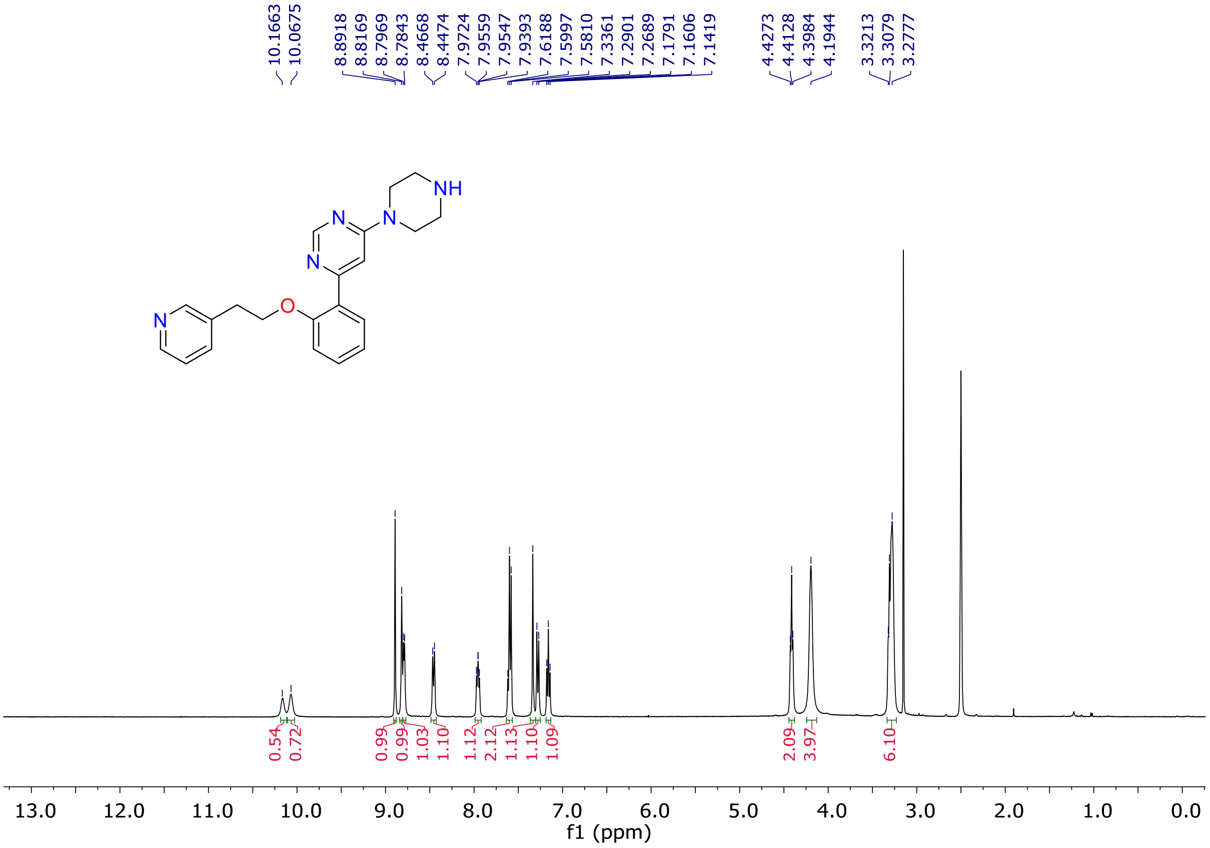


**Fig. S40.** [1]H-NMR spectrum of compound **16c** (DMSO-*d6*, 400 MHz).

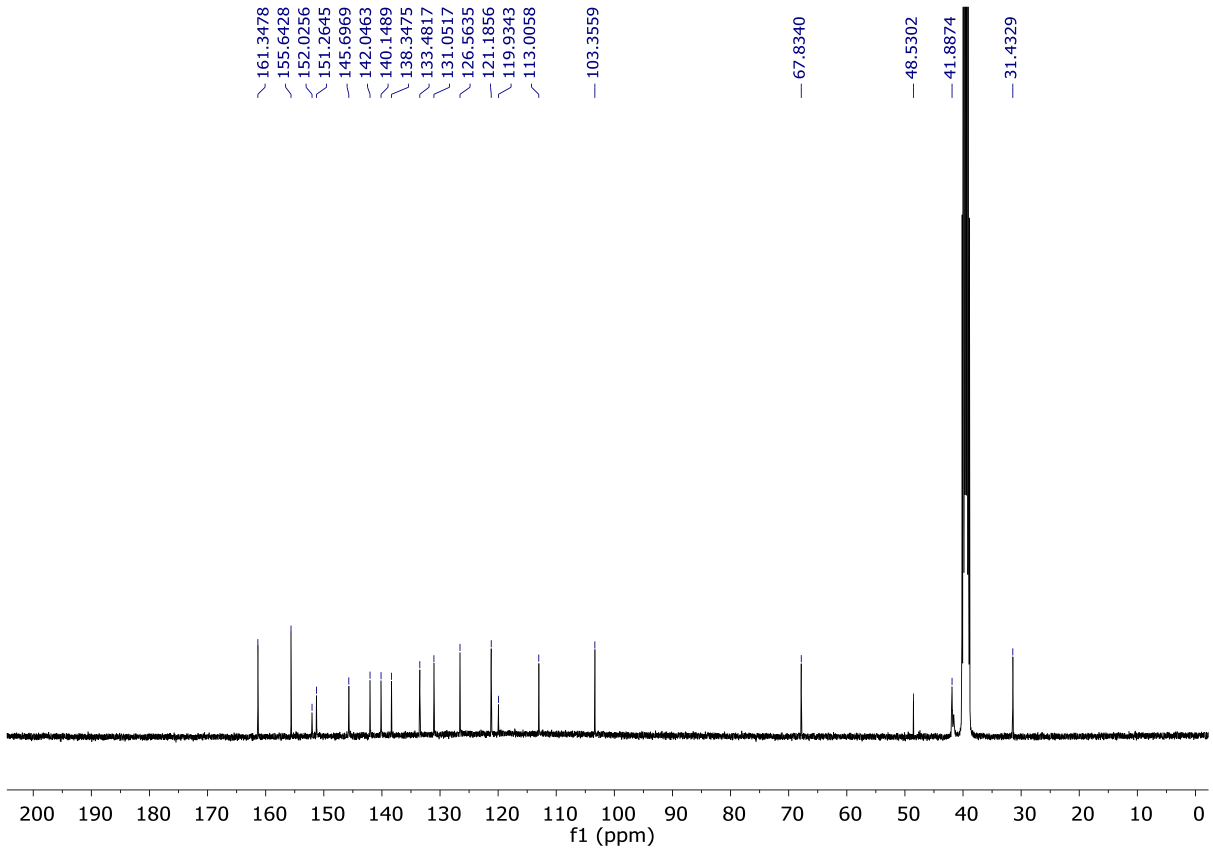


**Fig. S41.** $^{13}$C-NMR spectrum of compound **16c** (DMSO-*d6*, 100 MHz).

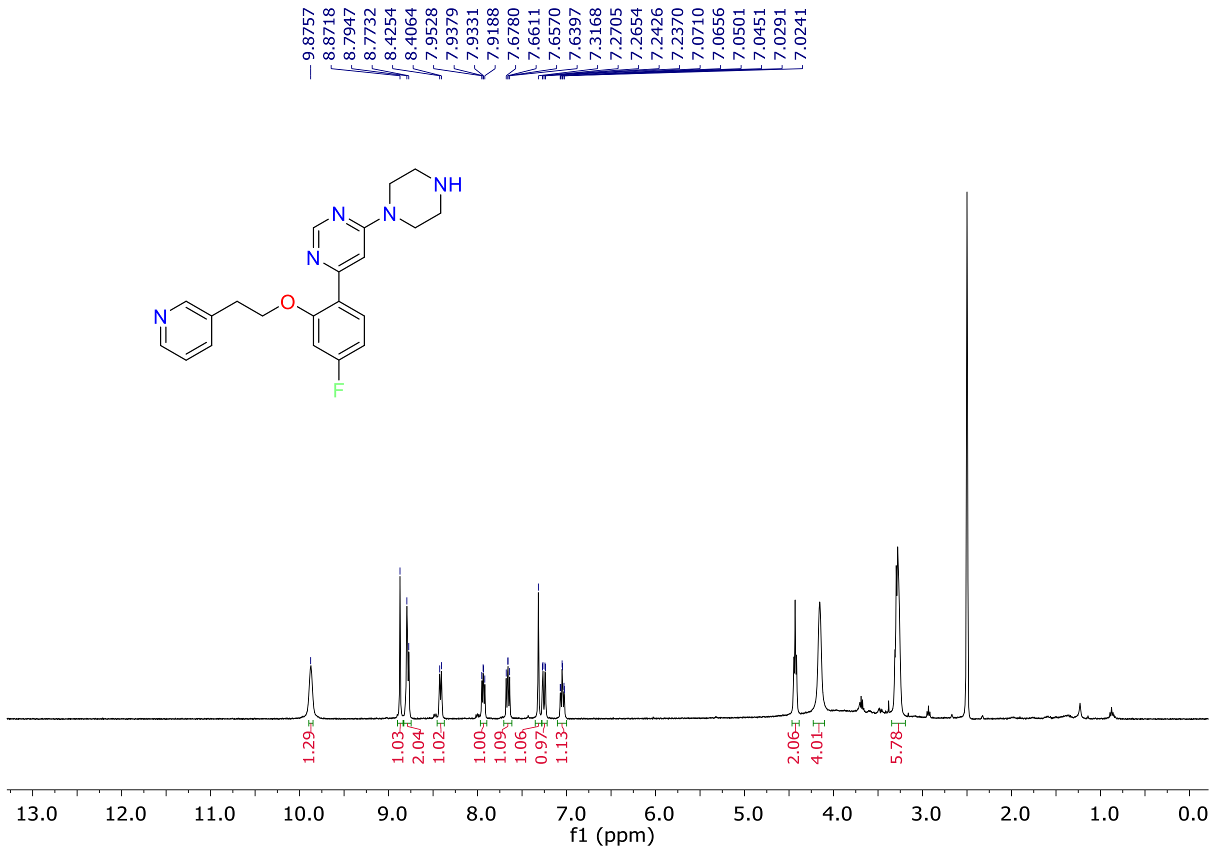


**Fig. S42.** $^{1}$H-NMR spectrum of compound **16d** (DMSO-*d6*, 400 MHz).

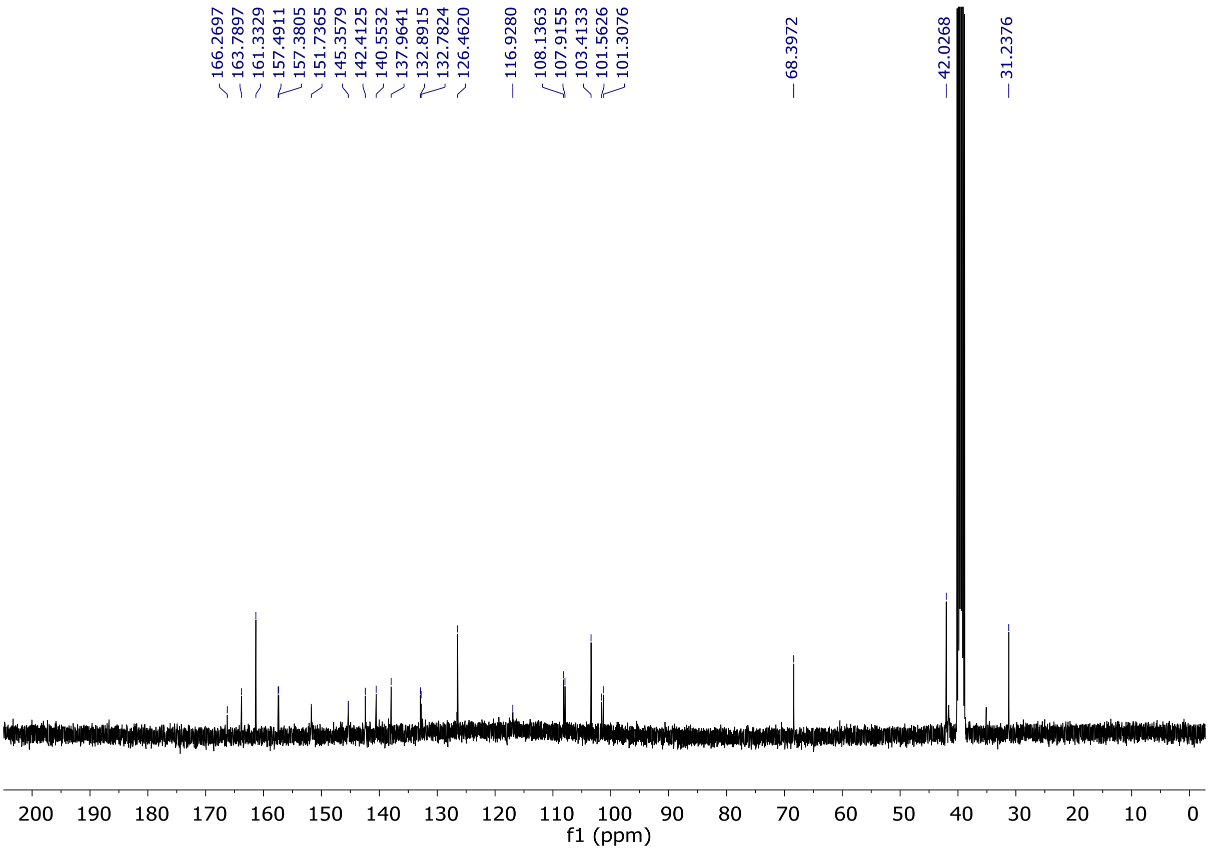


**Fig. S43.** $^{13}$C-NMR spectrum of compound **16d** (DMSO-*d6*, 100 MHz).

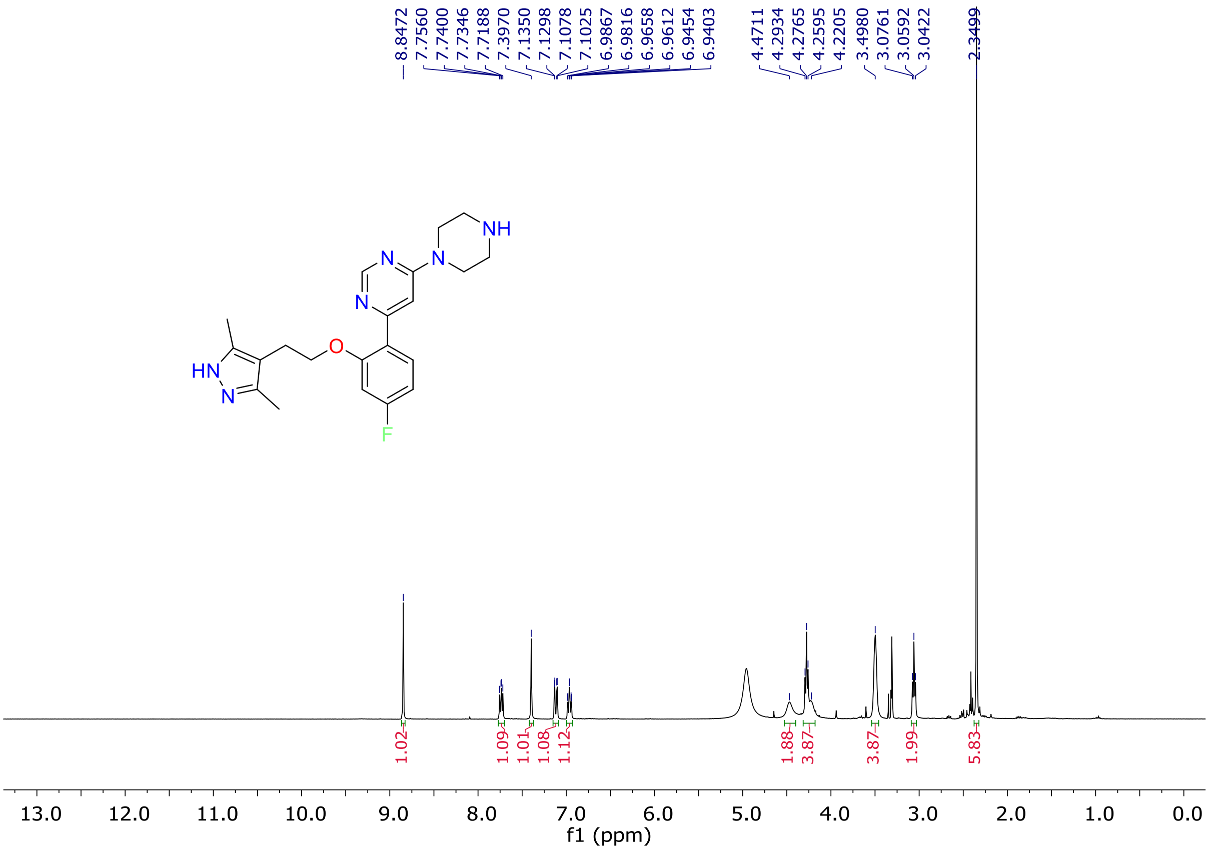


**Fig. S44.** $^{1}$H-NMR spectrum of compound **16e** ($CD_3OD$, 400 MHz).

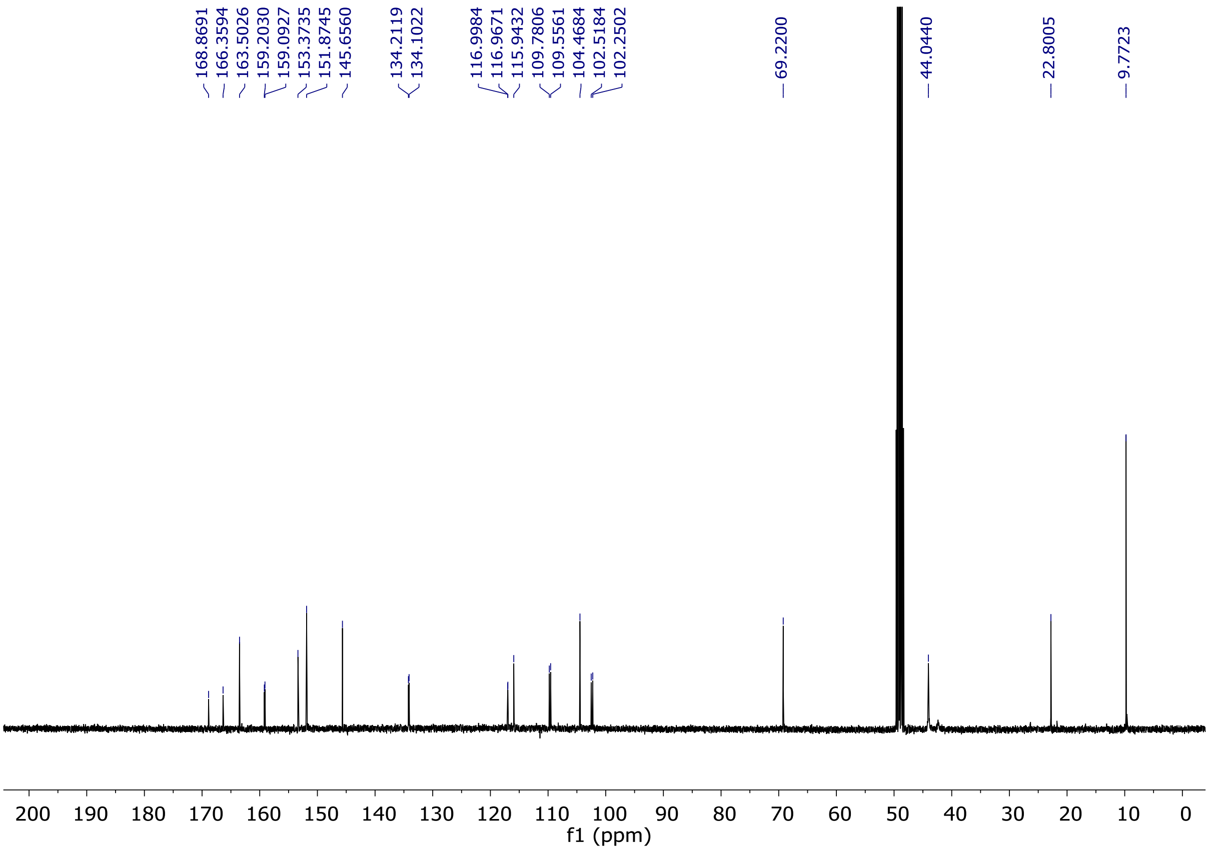


**Fig. S45.** $^{13}$C-NMR spectrum of compound **16e** ($CD_3OD$, 100 MHz).

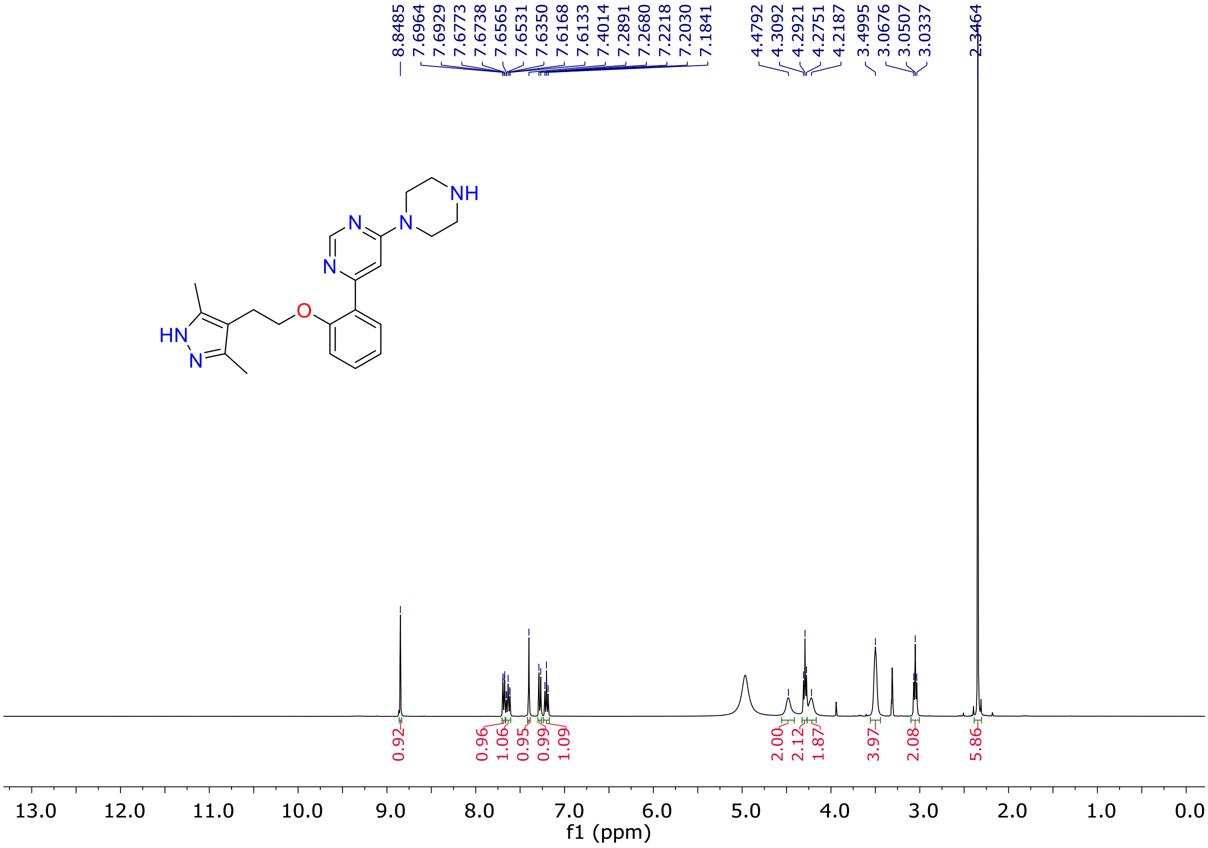


**Fig. S46.** $^{1}$H-NMR spectrum of compound **16f** ($CD_3OD$, 400 MHz).

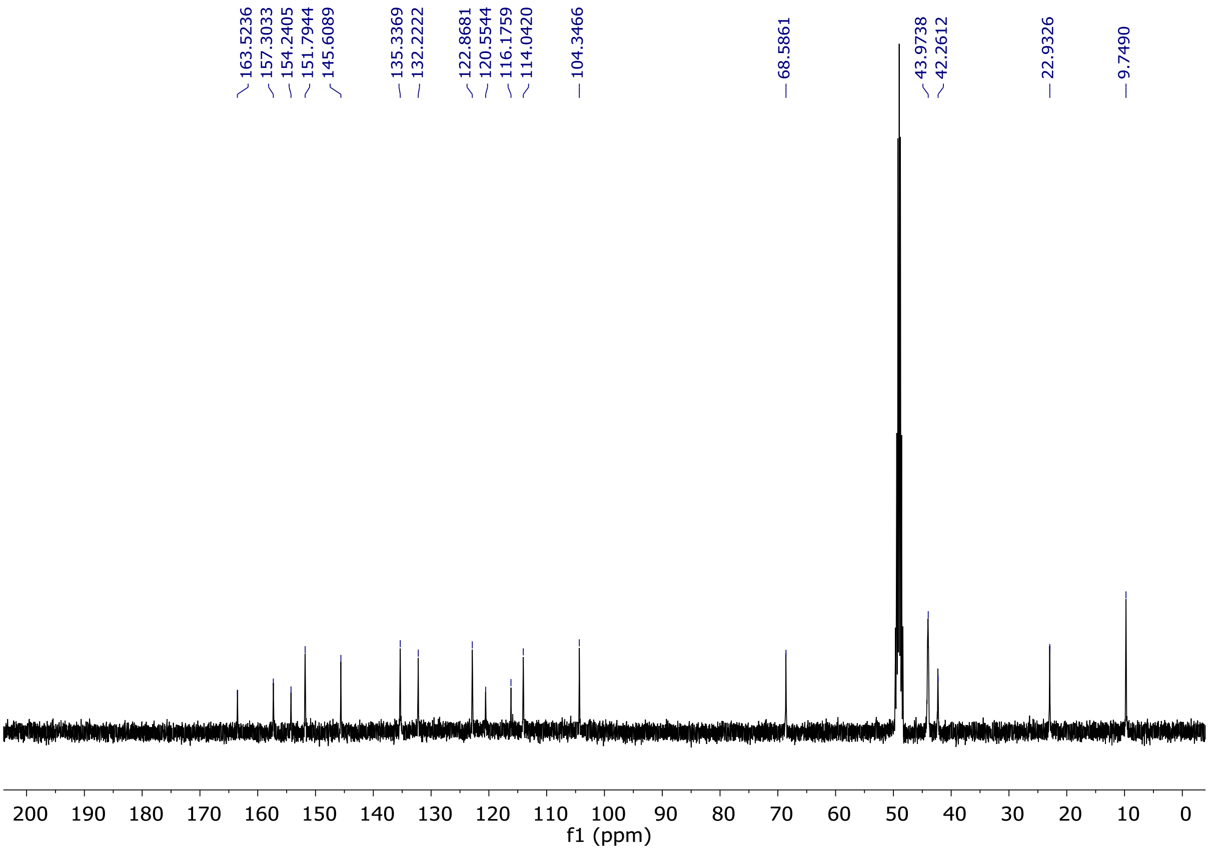


**Fig. S47.** $^{13}$C-NMR spectrum of compound **16f** ($CD_3OD$, 100 MHz).

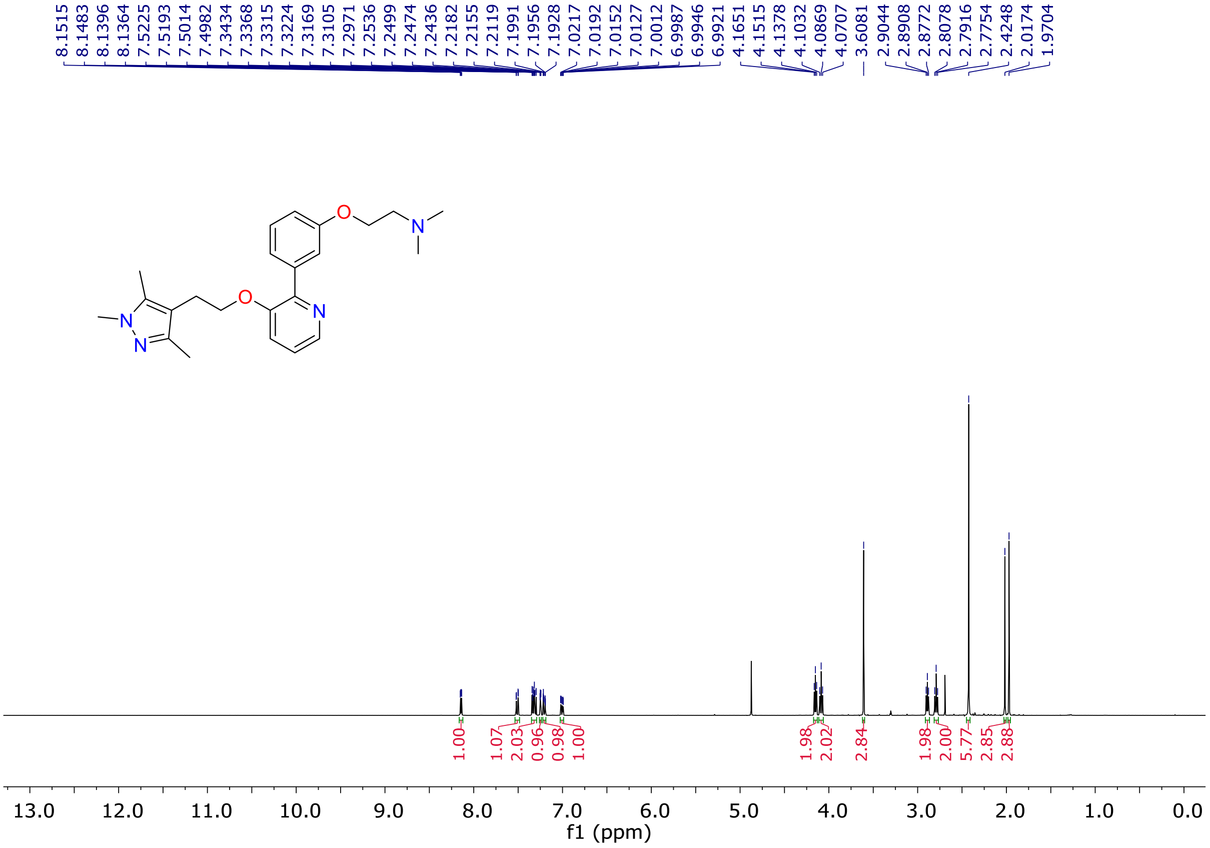


**Fig. S48.** $^{1}$H-NMR spectrum of compound **26a** ($CD_3OD$, 400 MHz).

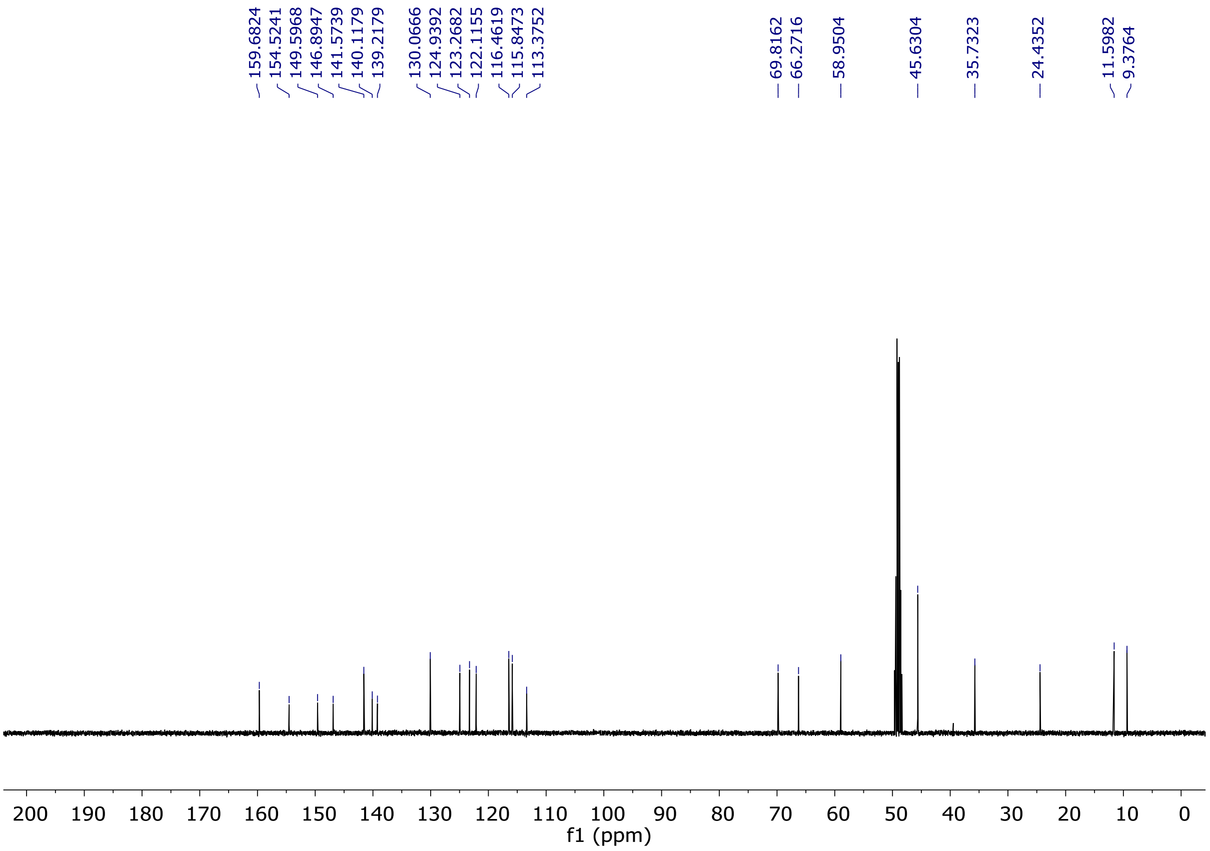


**Fig. S49.** [13]C-NMR spectrum of compound **26a** ($CD_3OD$, 100 MHz).

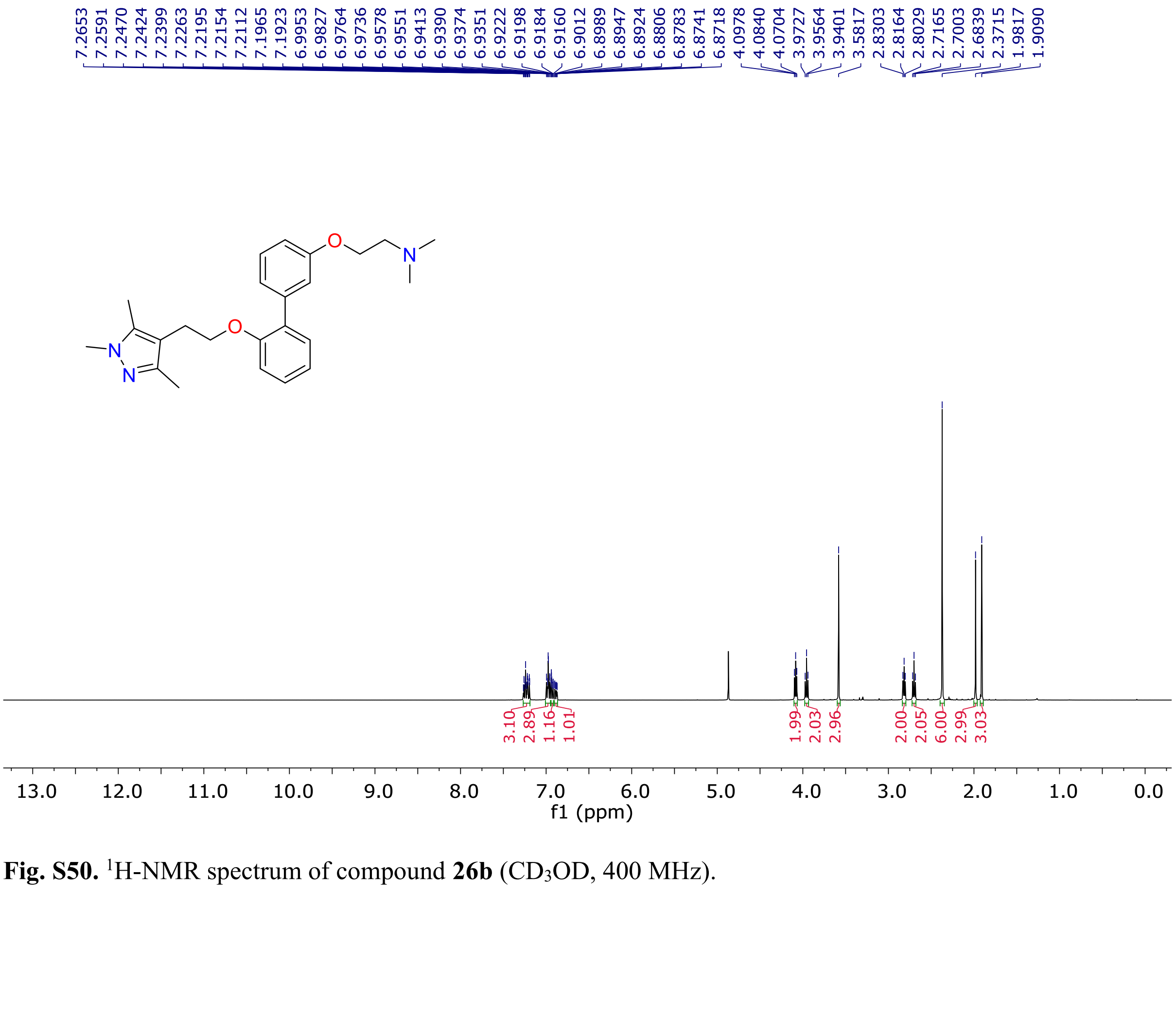


**Fig. S50.** [1]H-NMR spectrum of compound **26b** ($CD_3OD$, 400 MHz).

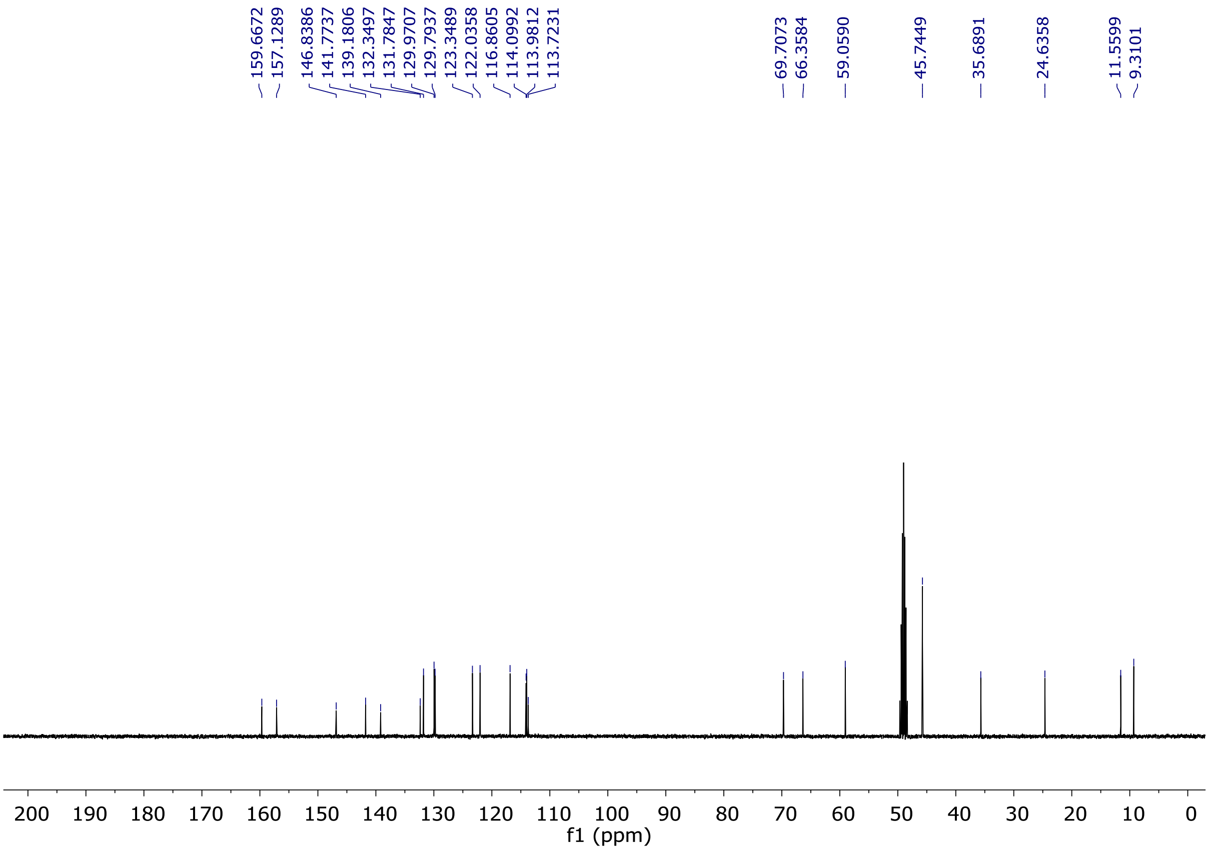


**Fig. S51.** $^{13}$C-NMR spectrum of compound **26b** ($CD_3OD$, 100 MHz).

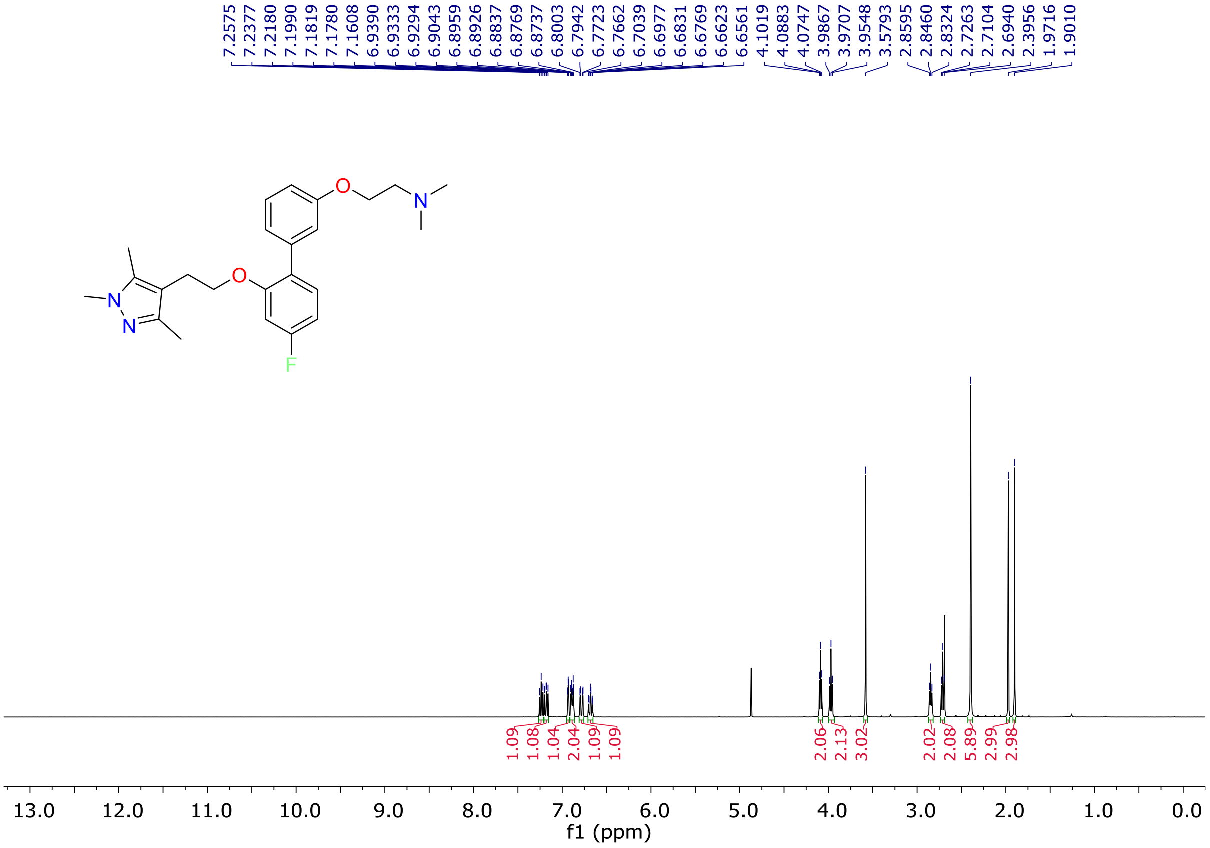


**Fig. S52.** $^{1}$H-NMR spectrum of compound **26c** ($CD_3OD$, 400 MHz).

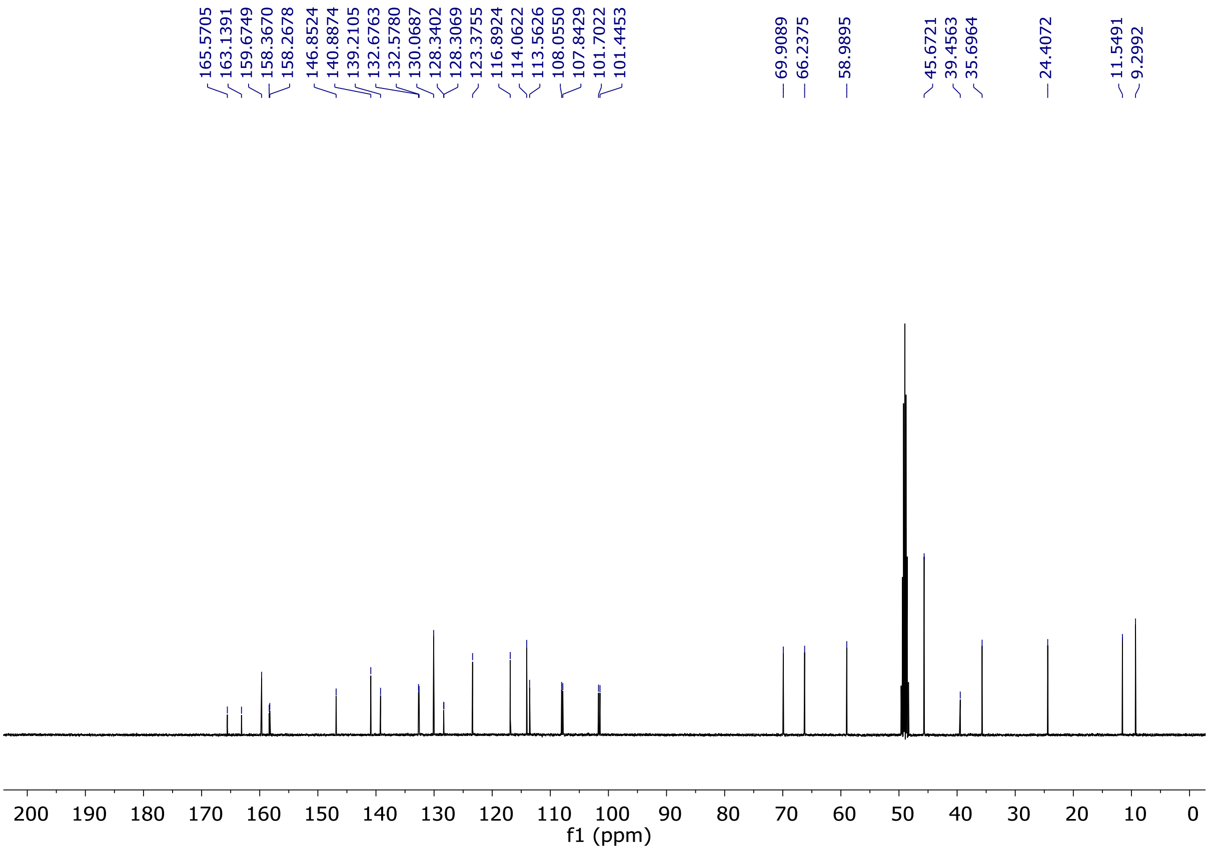


**Fig. S53.** $^{13}$C-NMR spectrum of compound **26c** ($CD_3OD$, 100 MHz).

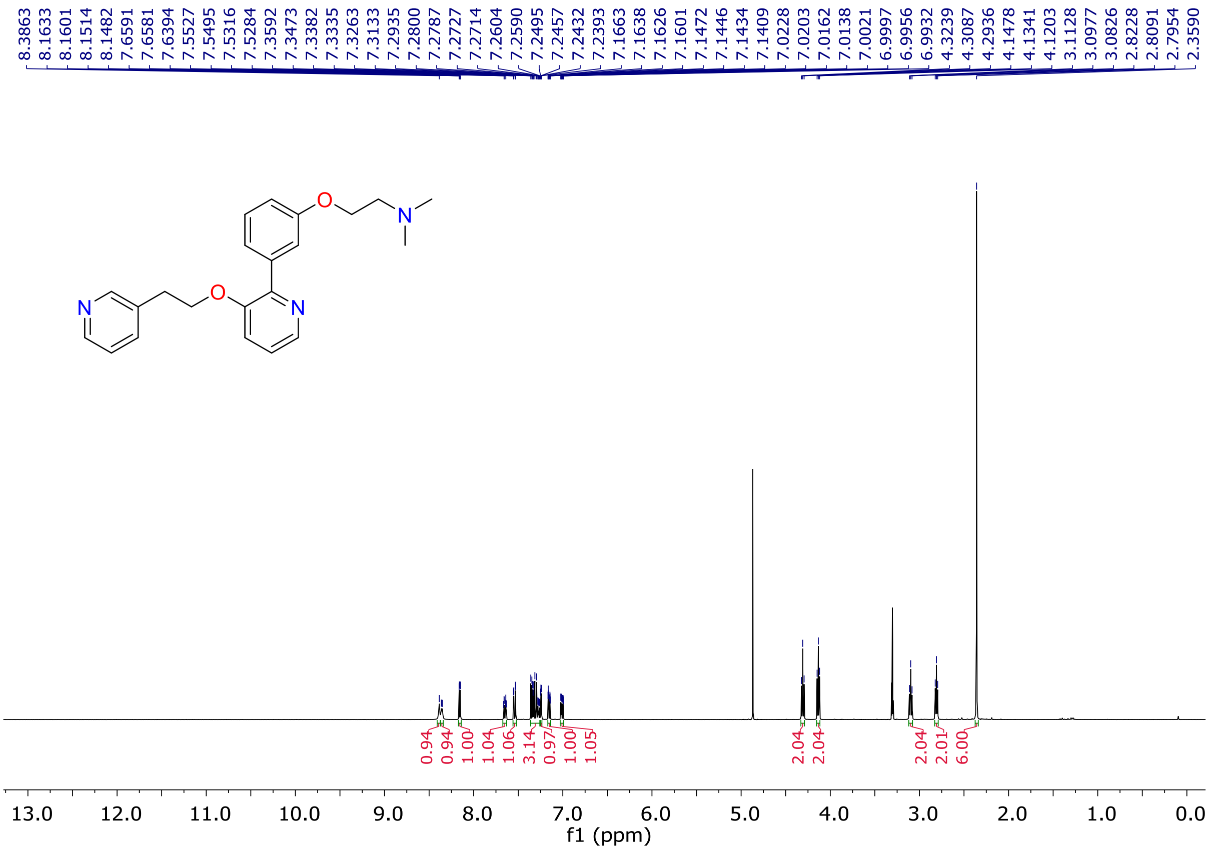


**Fig. S54.** $^{1}$H-NMR spectrum of compound **26d** ($CD_3OD$, 400 MHz).

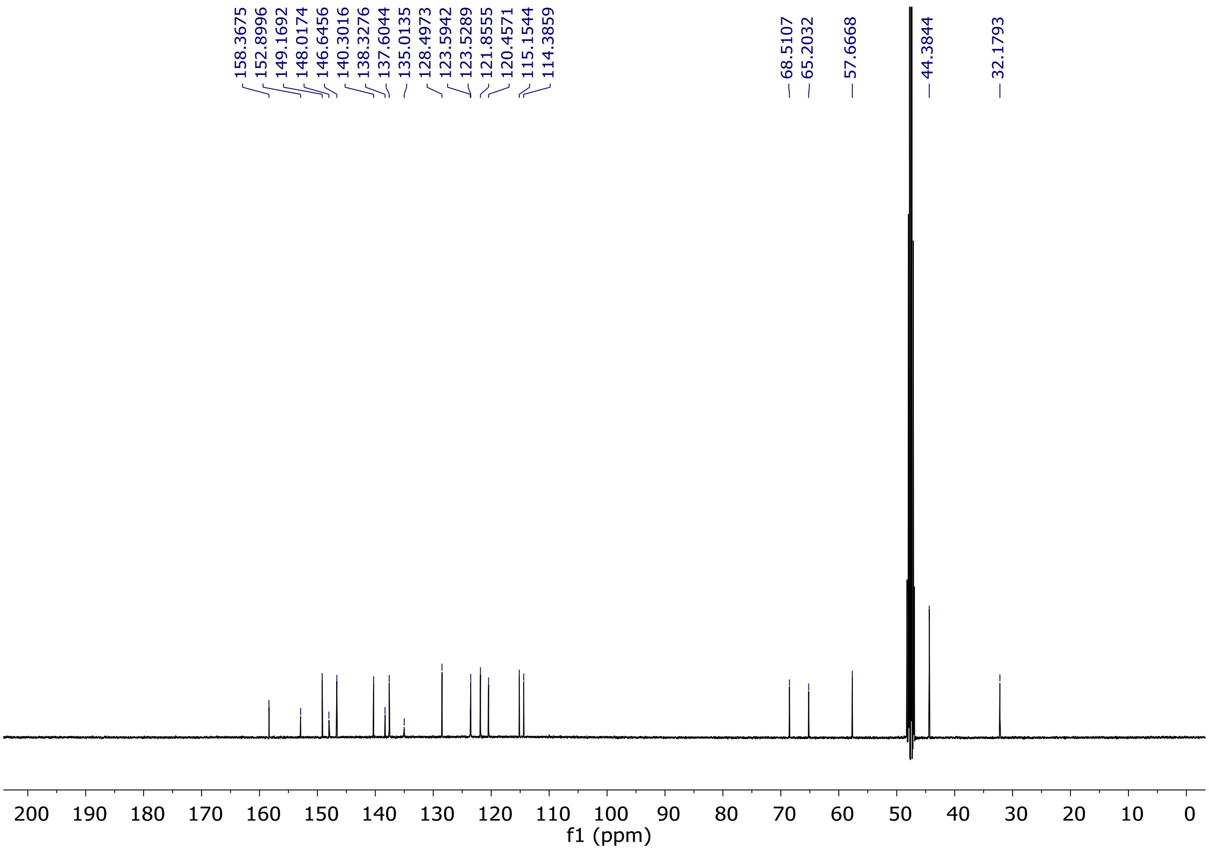

**Fig. S55.** $^{13}$C-NMR spectrum of compound **26d** ($CD_3OD$, 100 MHz).

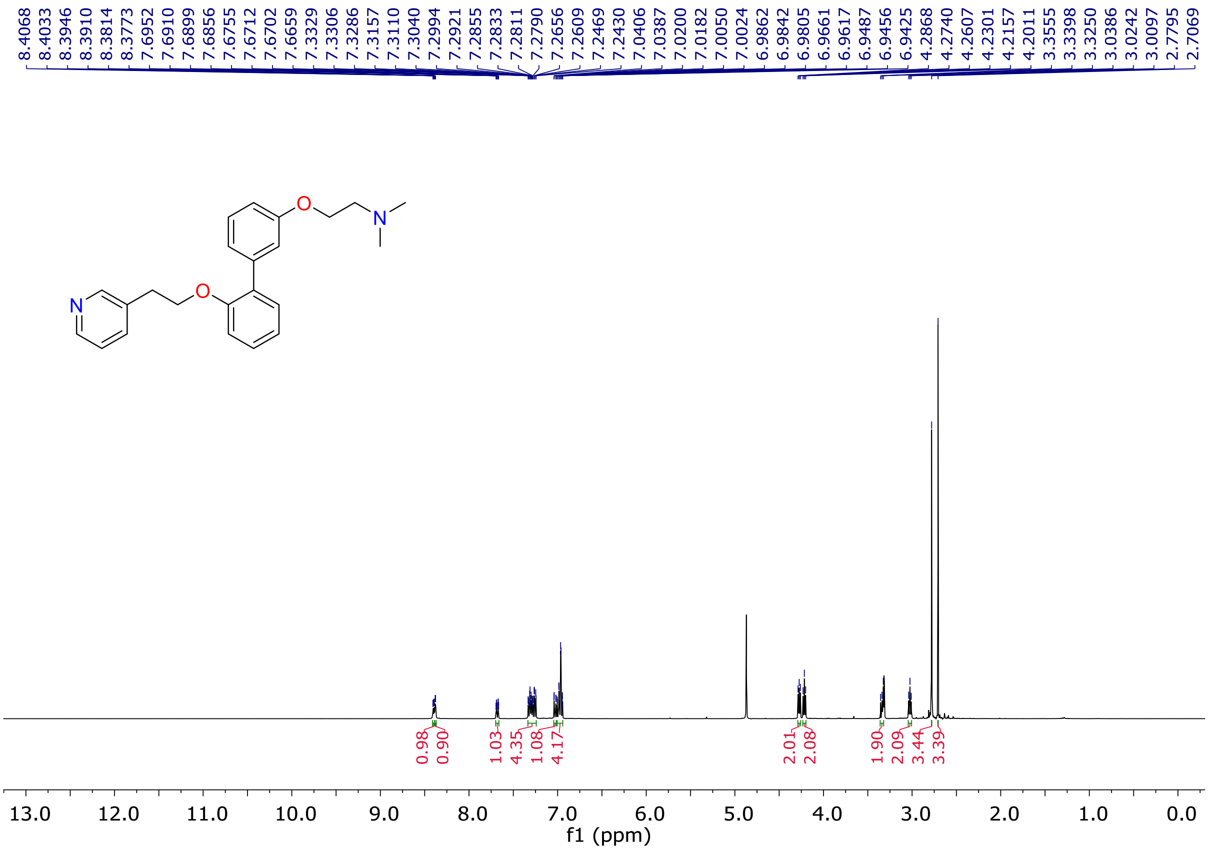

**Fig. S56.** $^{1}$H-NMR spectrum of compound **26e** ($CD_3OD$, 400 MHz).

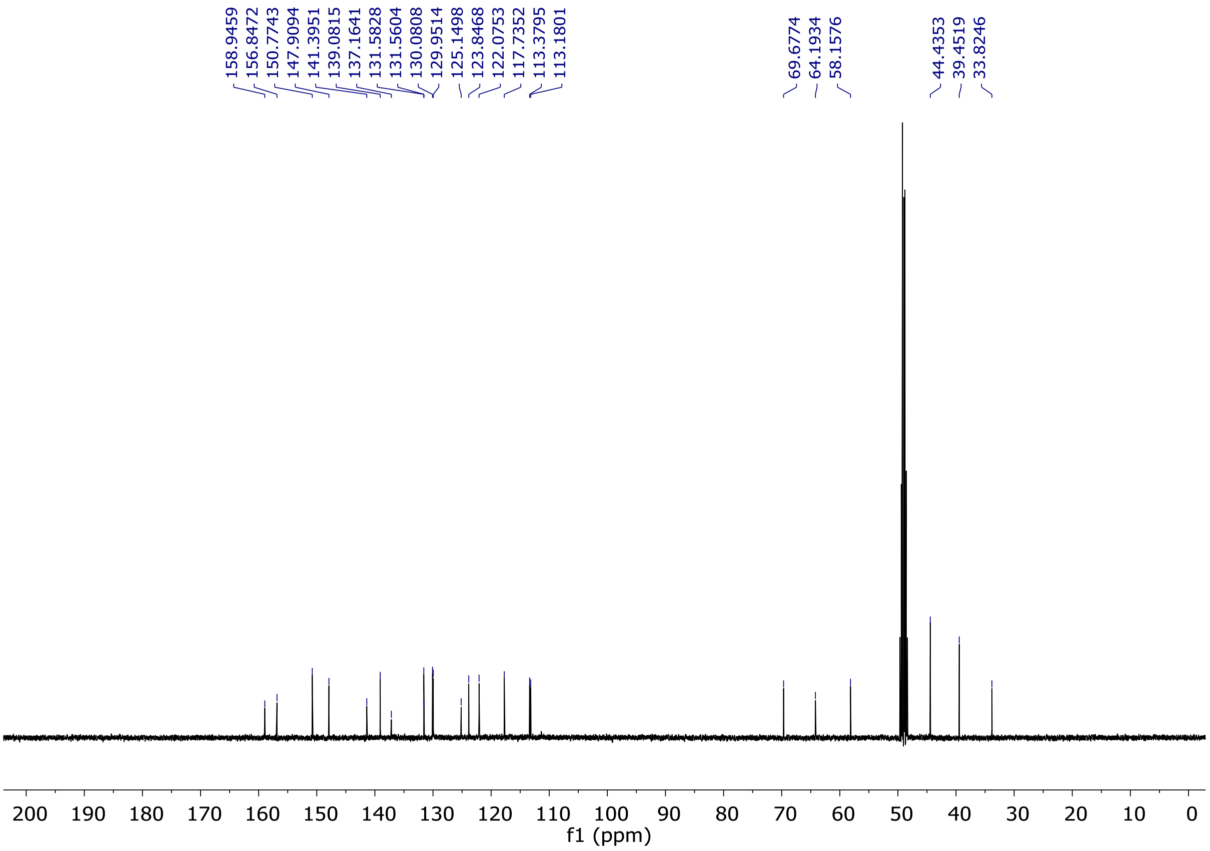


**Fig. S57.** [13]C-NMR spectrum of compound **26e** ($CD_3OD$, 100 MHz).

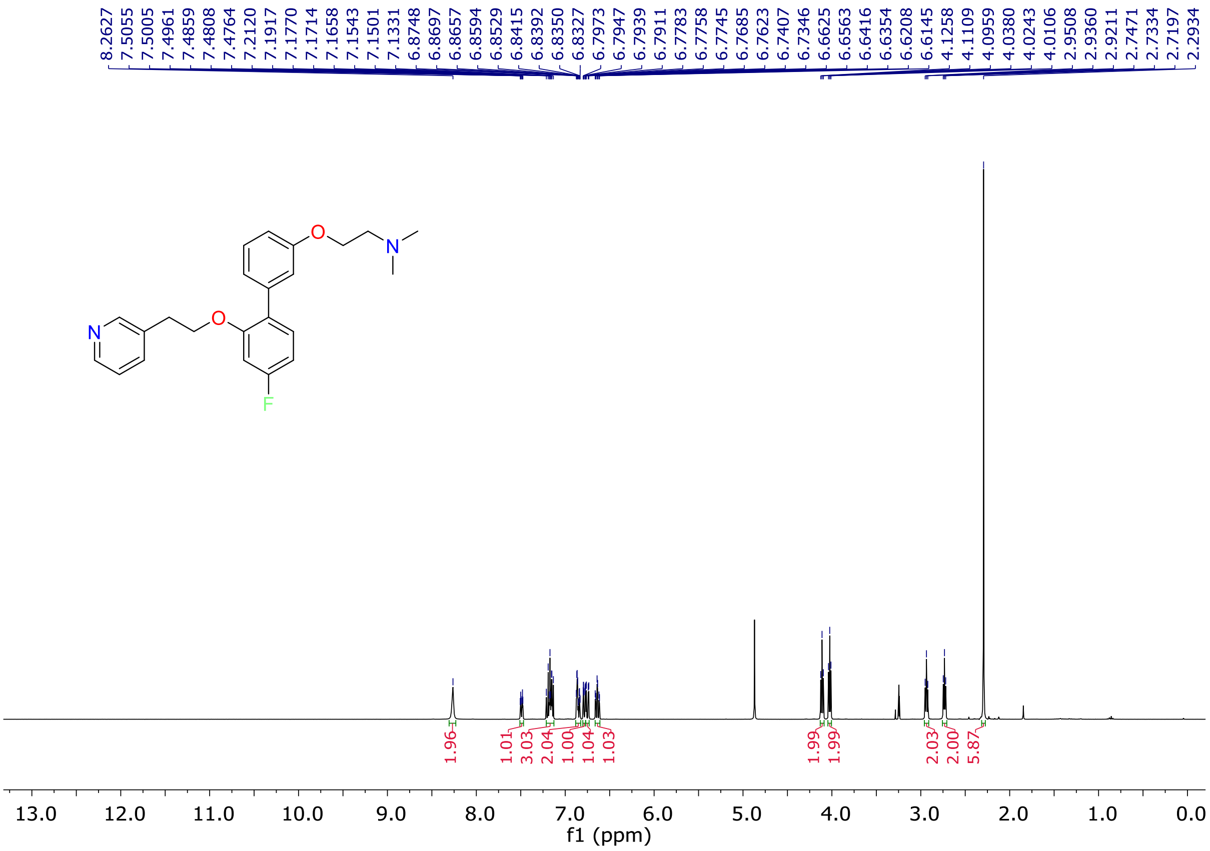


**Fig. S58.** [1]H-NMR spectrum of compound **26f** ($CD_3OD$, 400 MHz).

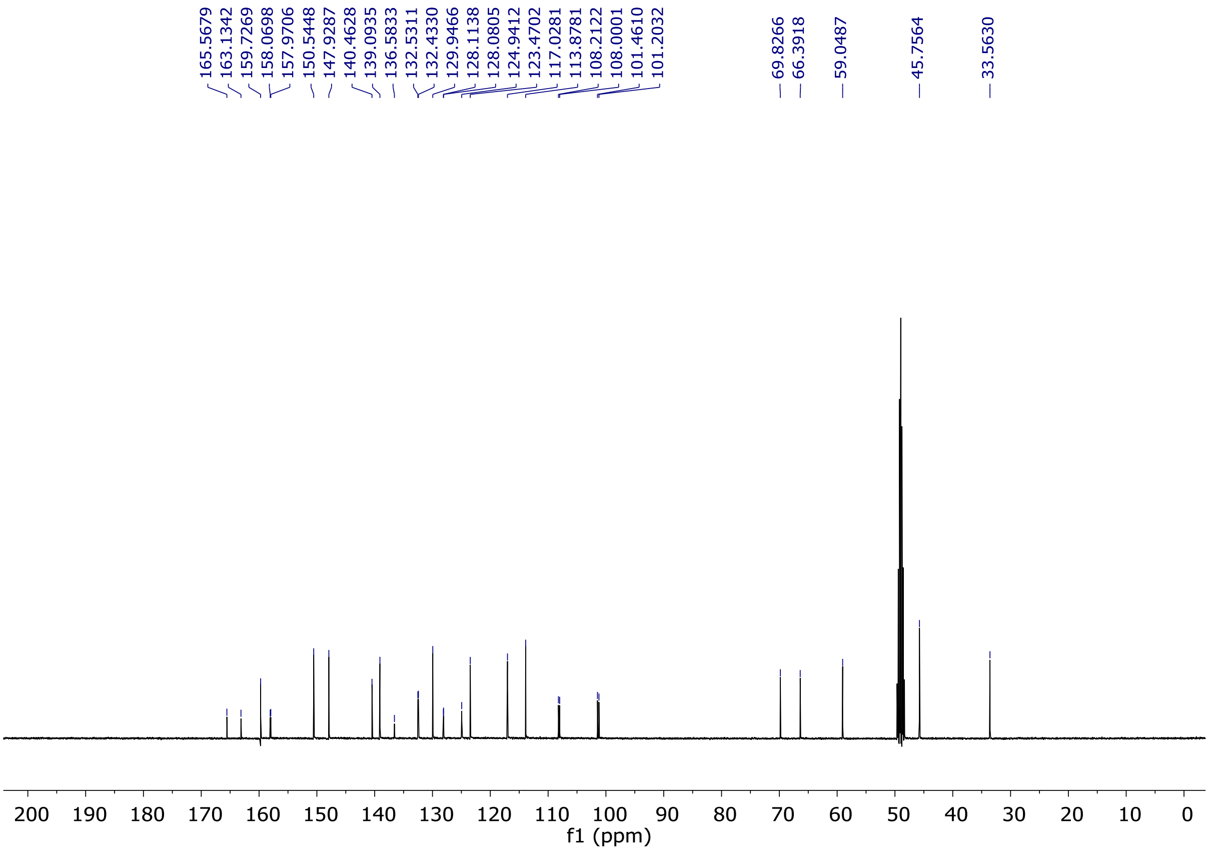


**Fig. S59.** $^{13}$C-NMR spectrum of compound **26f** ($CD_3OD$, 100 MHz).

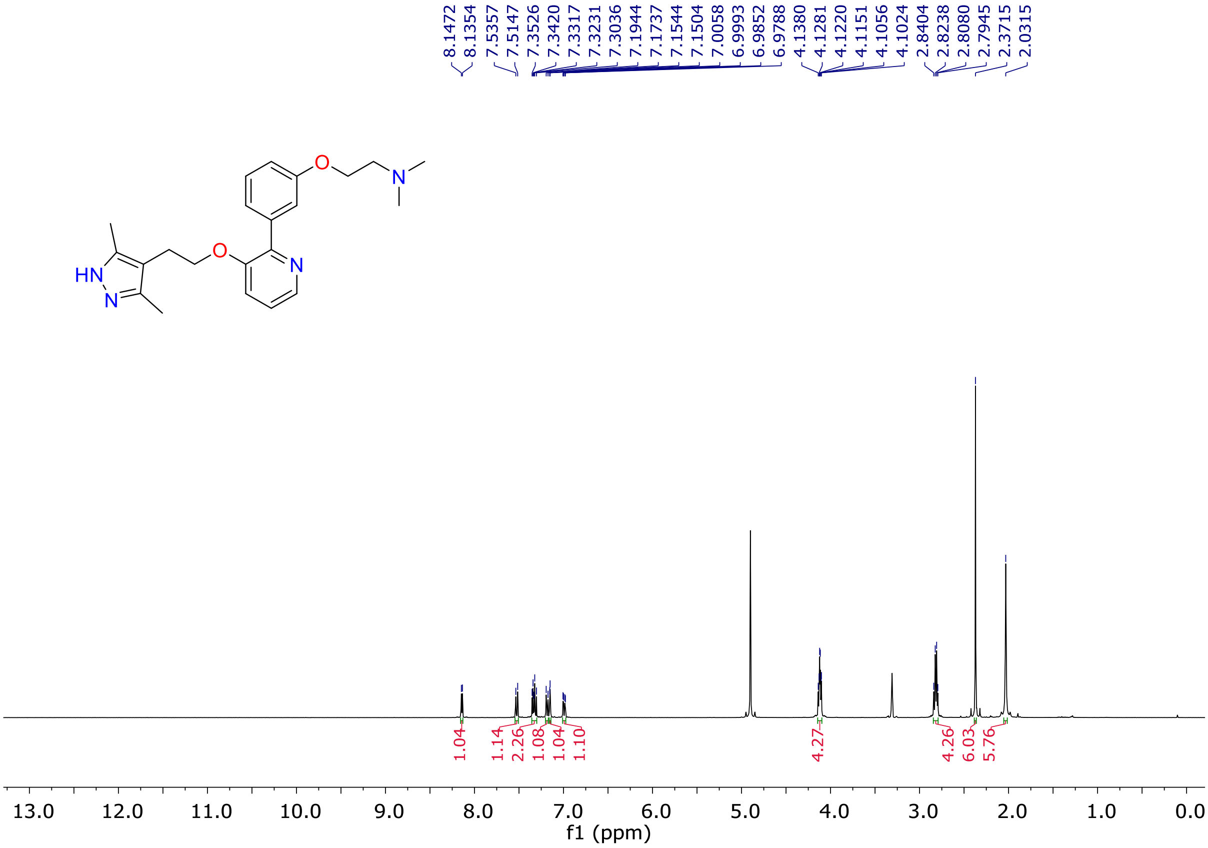


**Fig. S60.** $^{1}$H-NMR spectrum of compound **26g** ($CD_3OD$, 400 MHz).

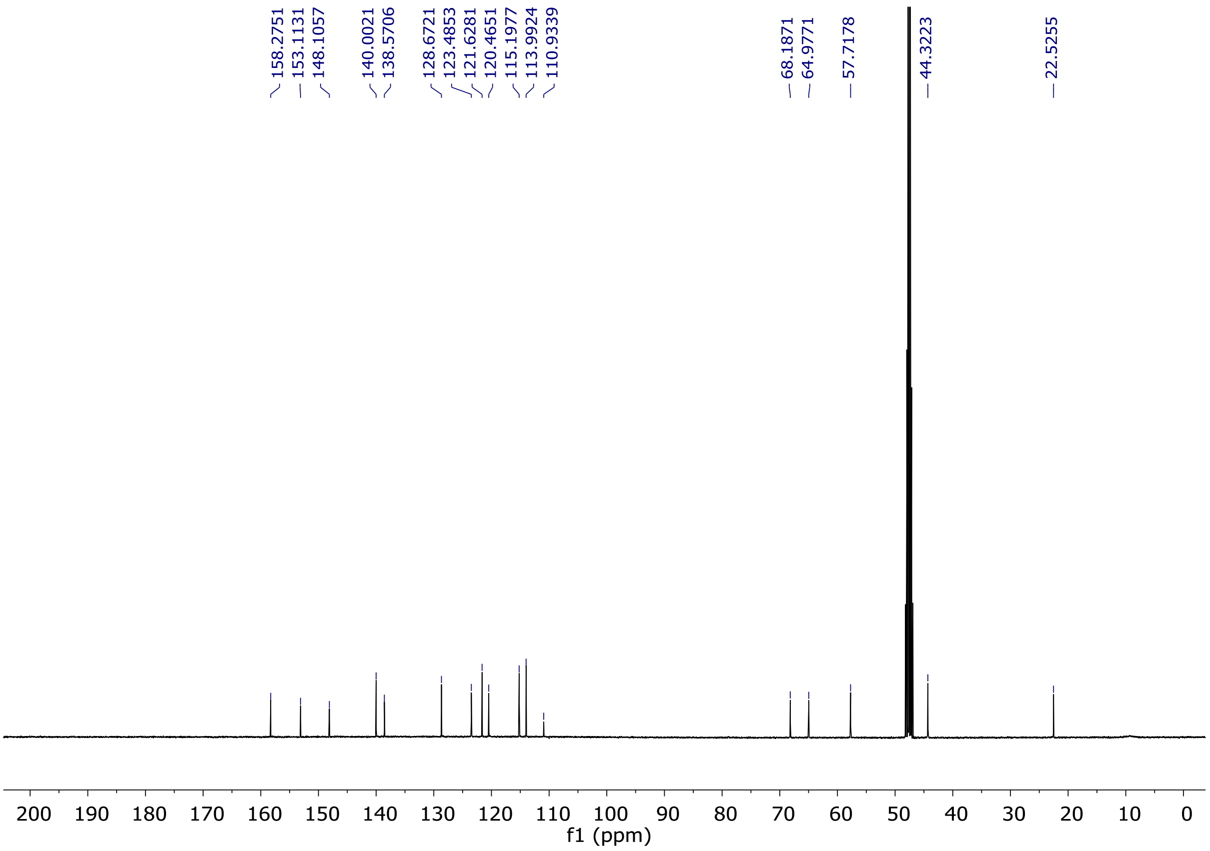


**Fig. S61.** $^{13}$C-NMR spectrum of compound **26g** ($CD_3OD$, 100 MHz).

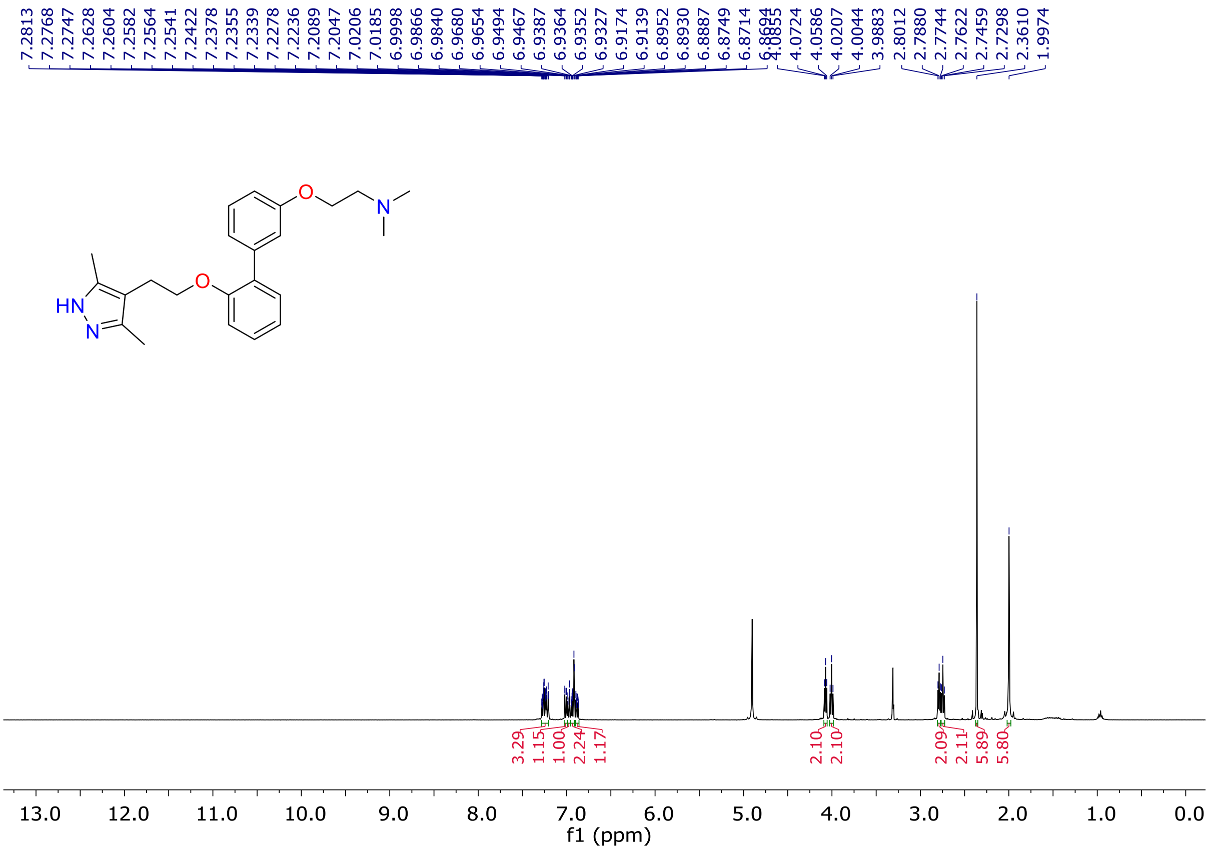


**Fig. S62.** $^{1}$H-NMR spectrum of compound **26h** ($CD_3OD$, 400 MHz).

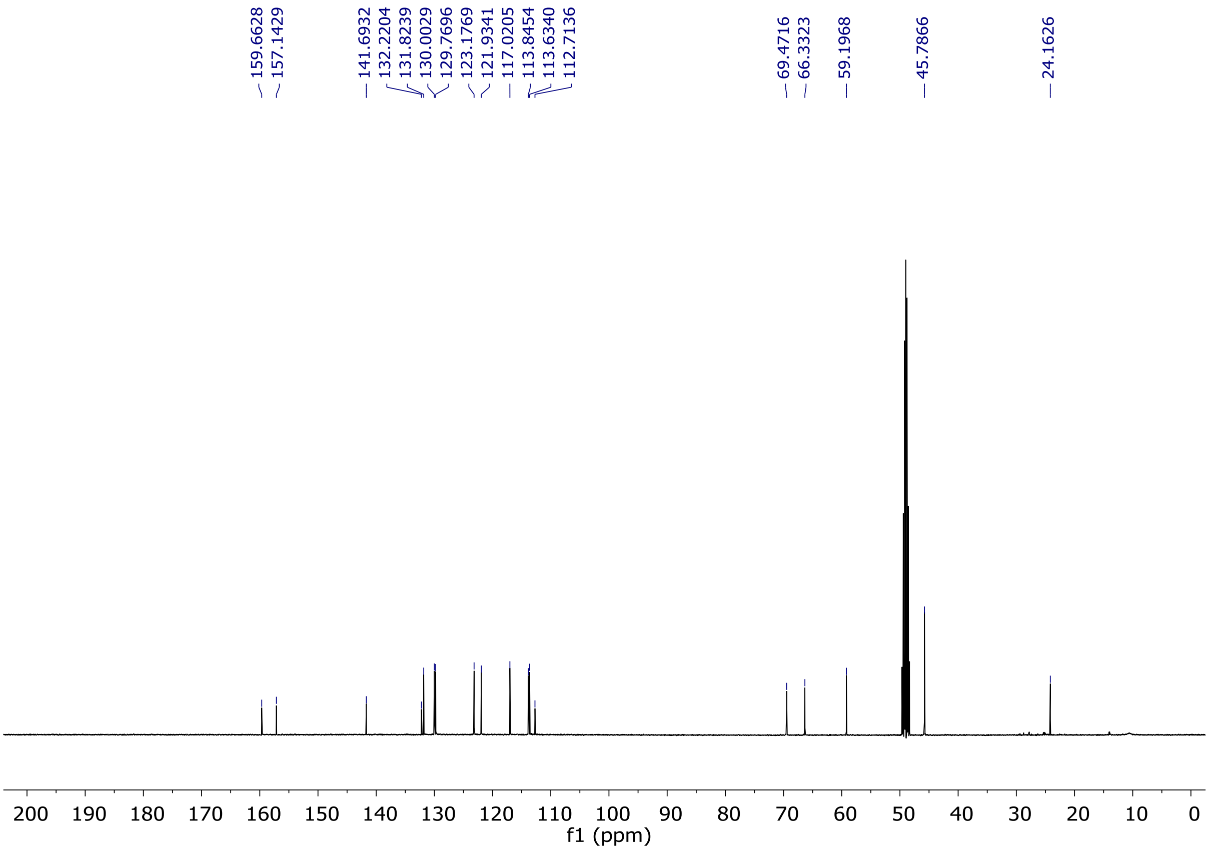


**Fig. S63.** $^{13}$C-NMR spectrum of compound **26h** ($CD_3OD$, 100 MHz).

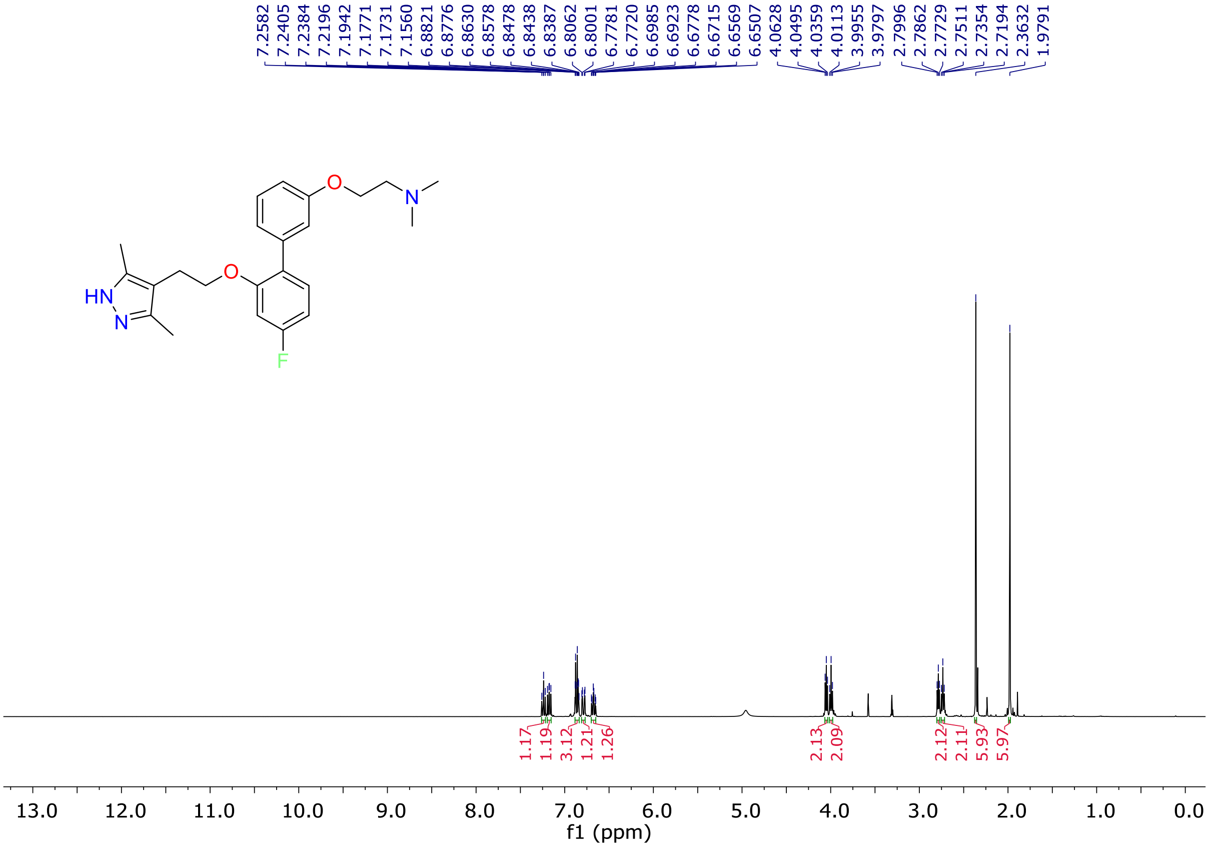


**Fig. S64.** $^{1}$H-NMR spectrum of compound **26i** ($CD_3OD$, 400 MHz).

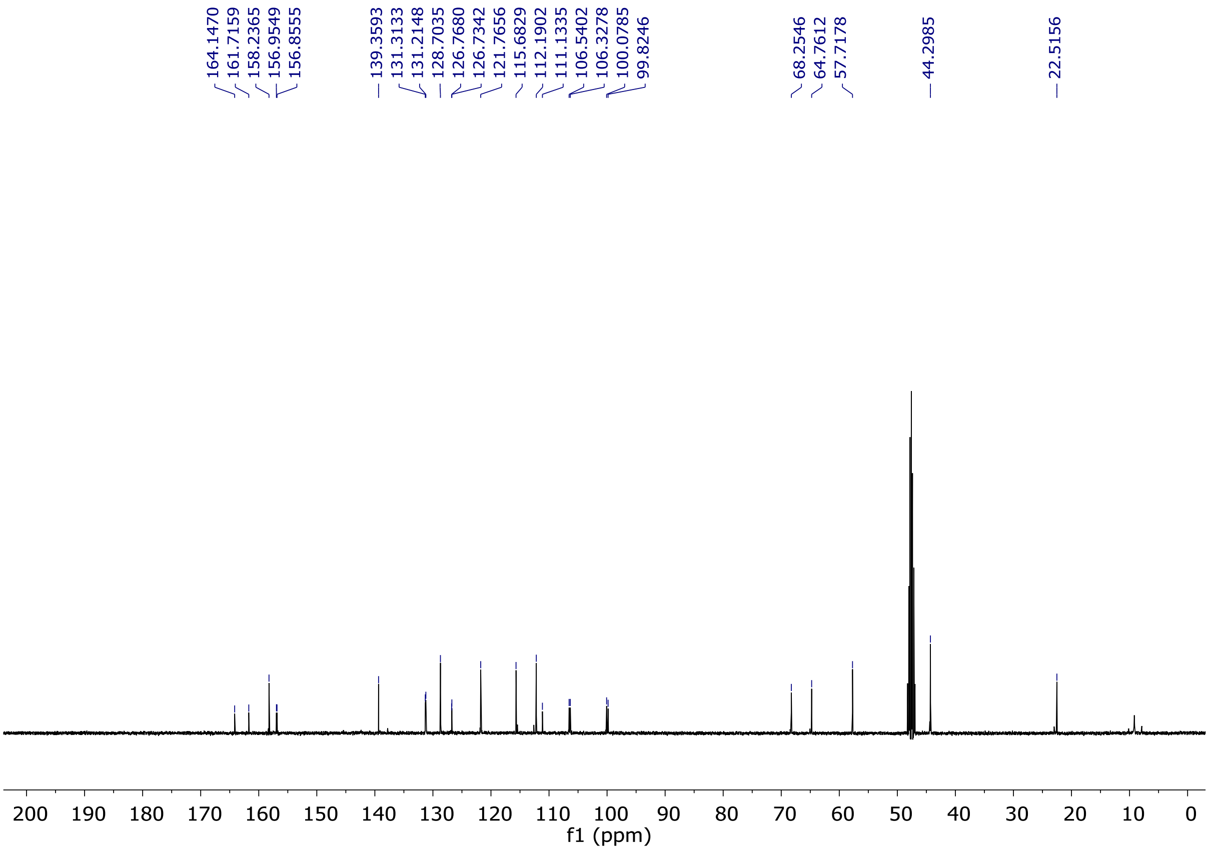


**Fig. S65.** $^{13}$C-NMR spectrum of compound **26i** ($CD_3OD$, 100 MHz).

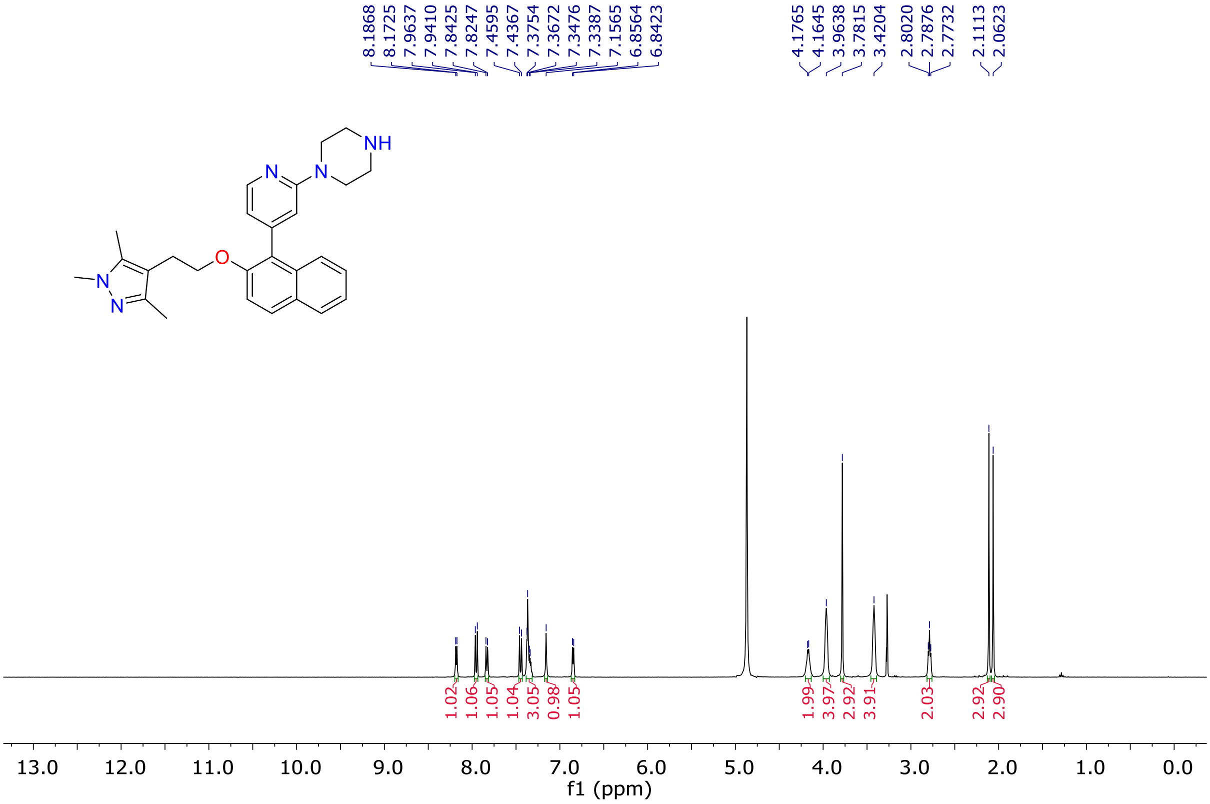


**Fig. S66.** $^{1}$H-NMR spectrum of compound **30a** ($CD_3OD$, 400 MHz).

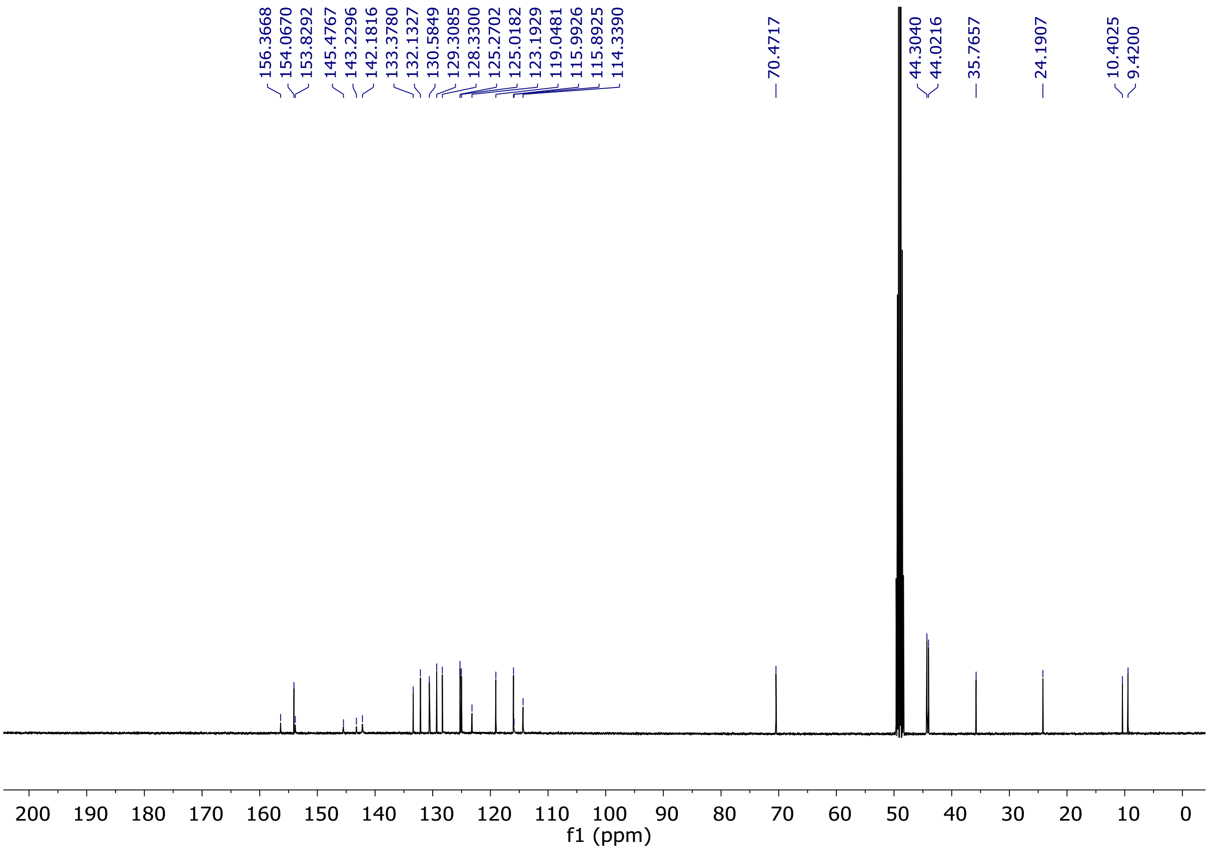


**Fig. S67.** [13]C-NMR spectrum of compound **30a** ($CD_3OD$, 100 MHz).

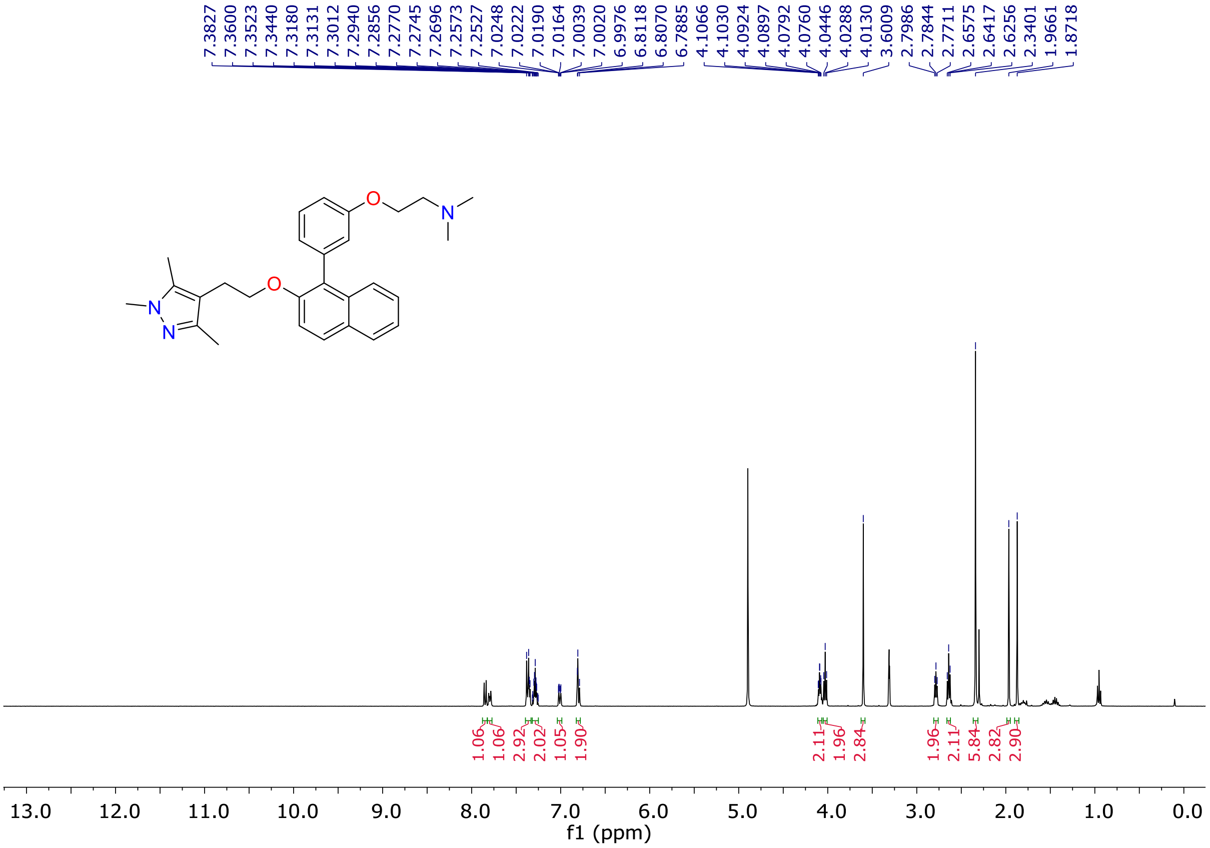


**Fig. S68.** [1]H-NMR spectrum of compound **30b** ($CD_3OD$, 400 MHz).

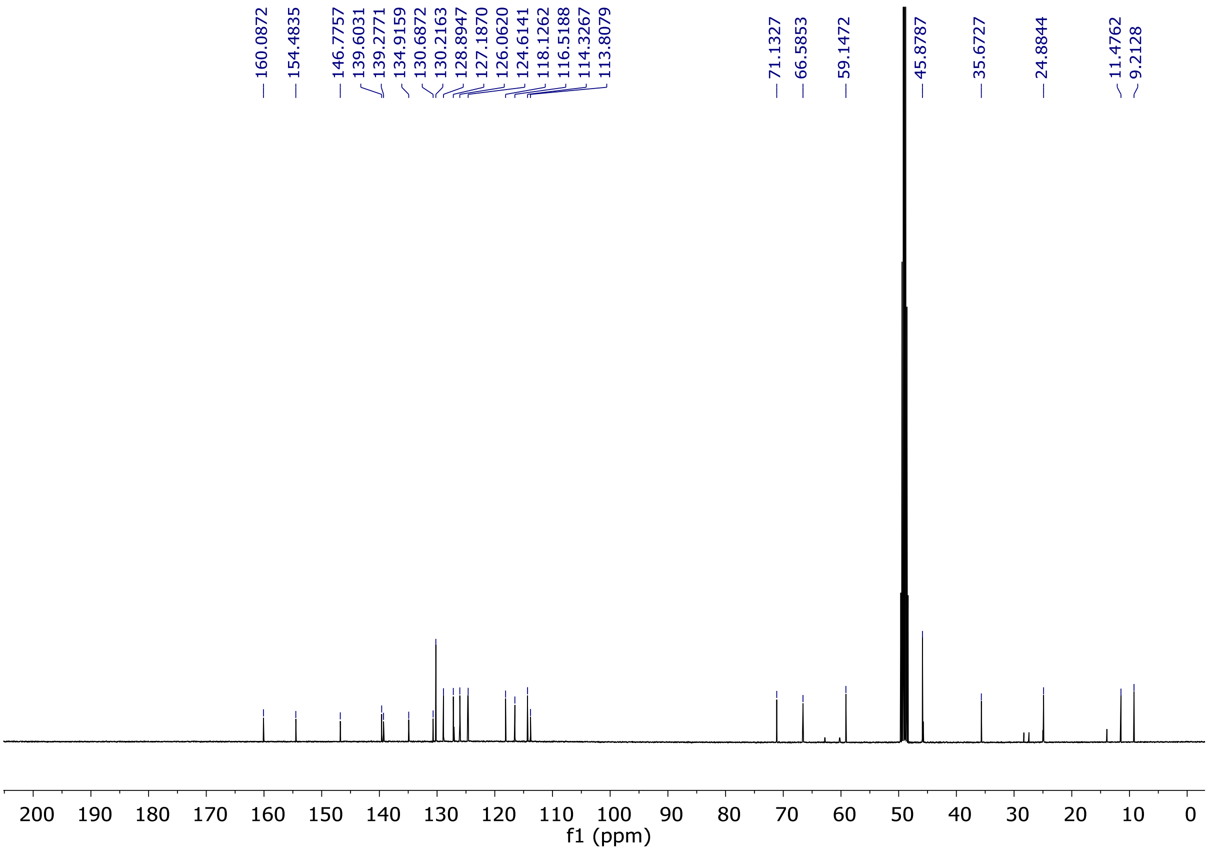


**Fig. S69.** $^{13}$C-NMR spectrum of compound **30b** ($CD_3OD$, 100 MHz).

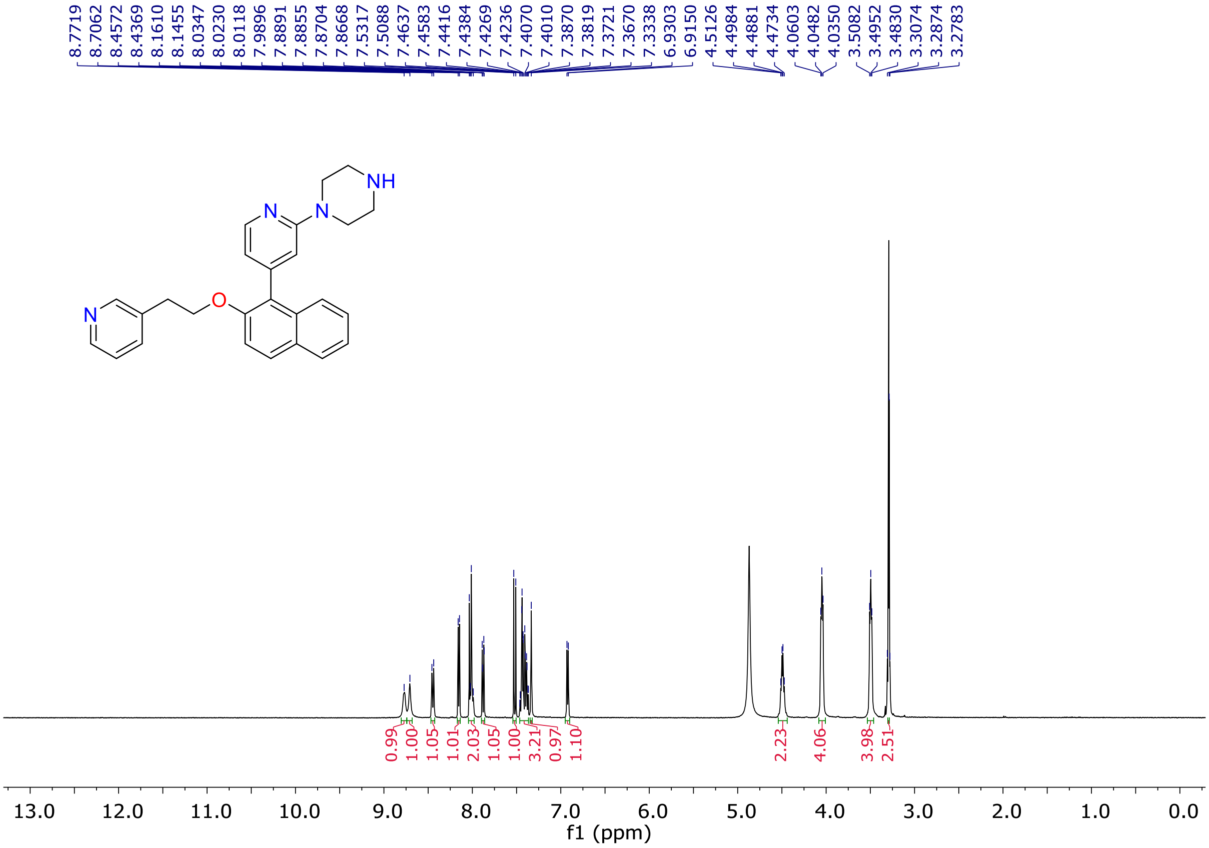


**Fig. S70.** $^{1}$H-NMR spectrum of compound **30c** ($CD_3OD$, 400 MHz).

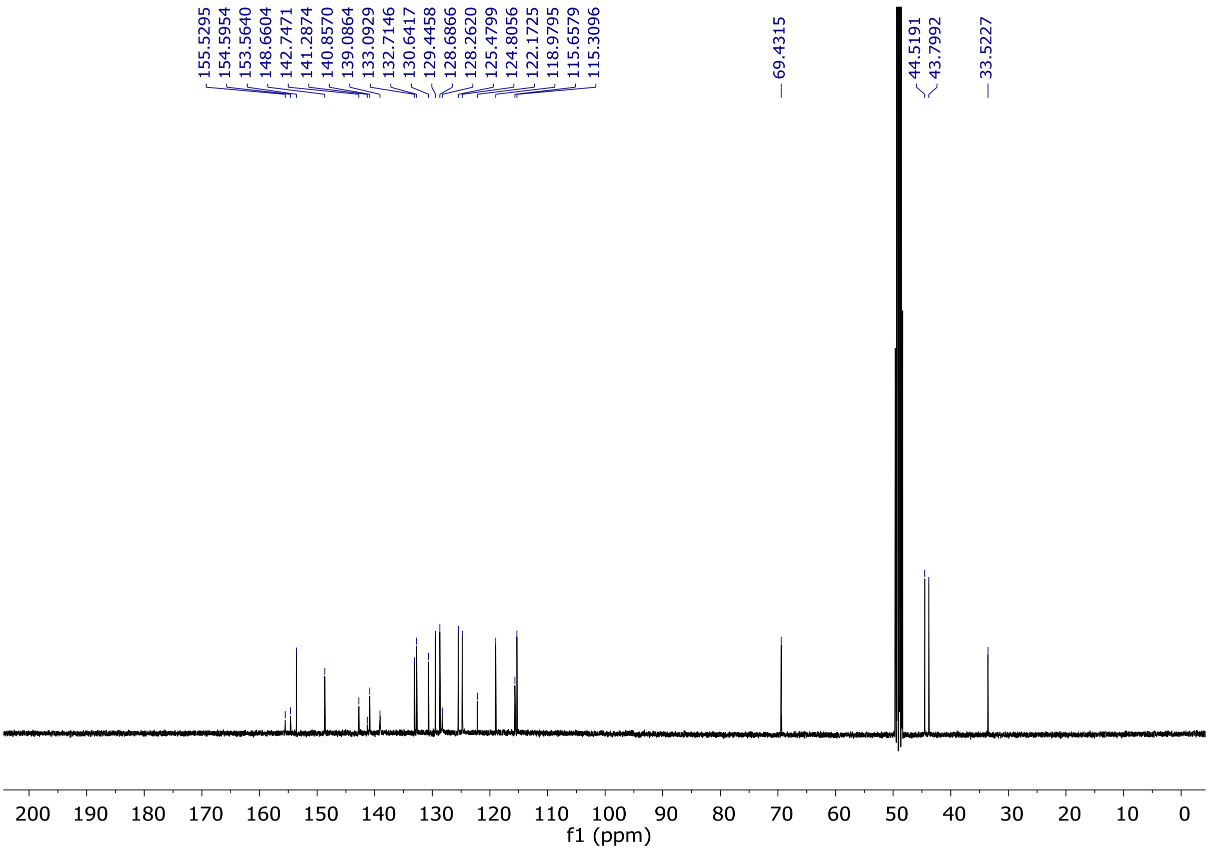


**Fig. S71.** [13]C-NMR spectrum of compound **30c** ($CD_3OD$, 100 MHz).

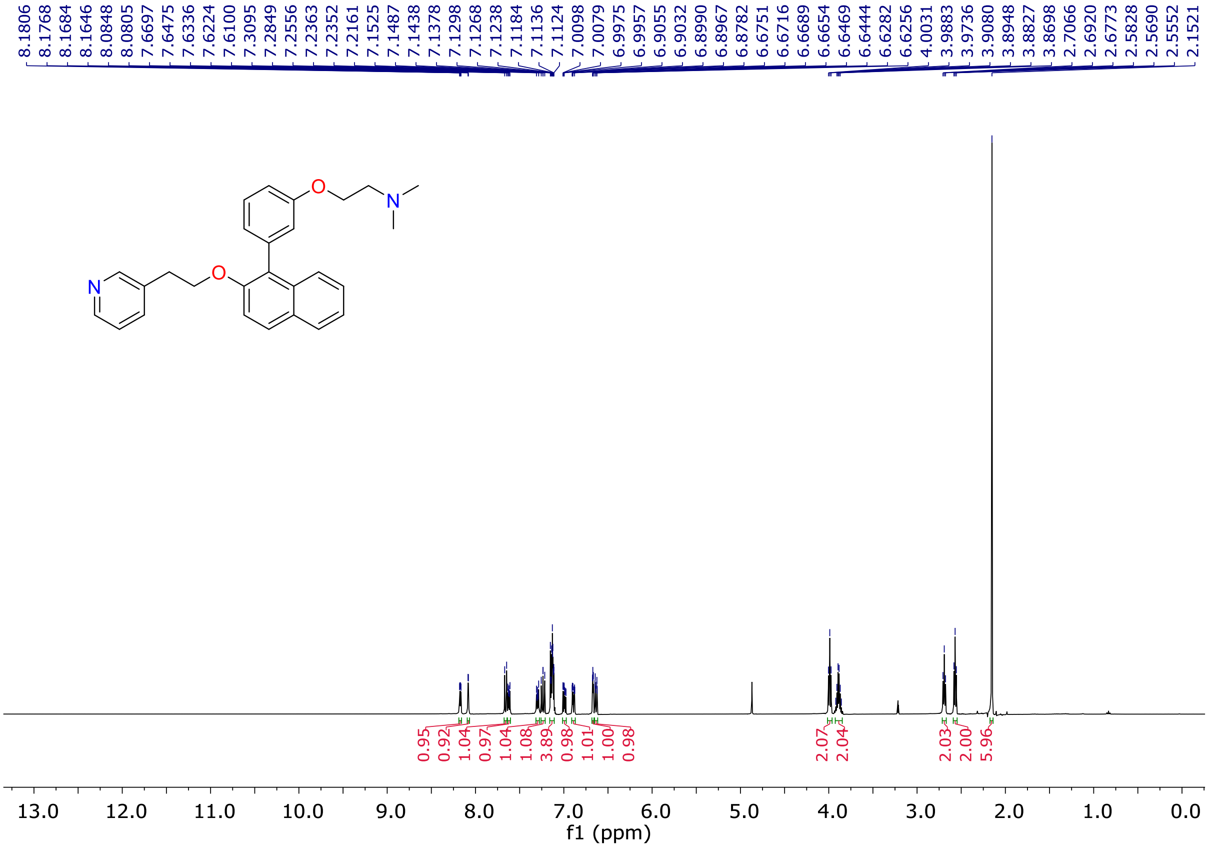


**Fig. S72.** [1]H-NMR spectrum of compound **30d** ($CD_3OD$, 400 MHz).

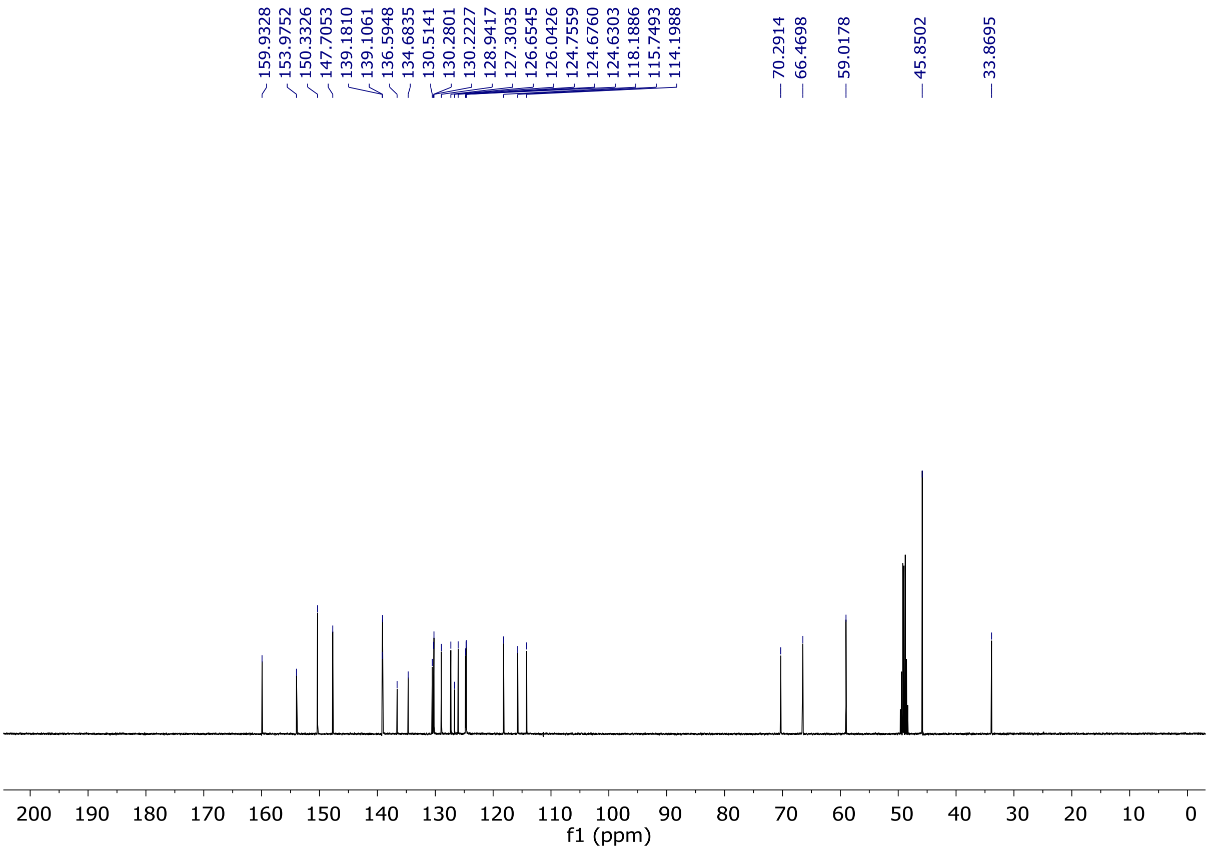


**Fig. S73.** $^{13}$C-NMR spectrum of compound **30d** ($CD_3OD$, 100 MHz).

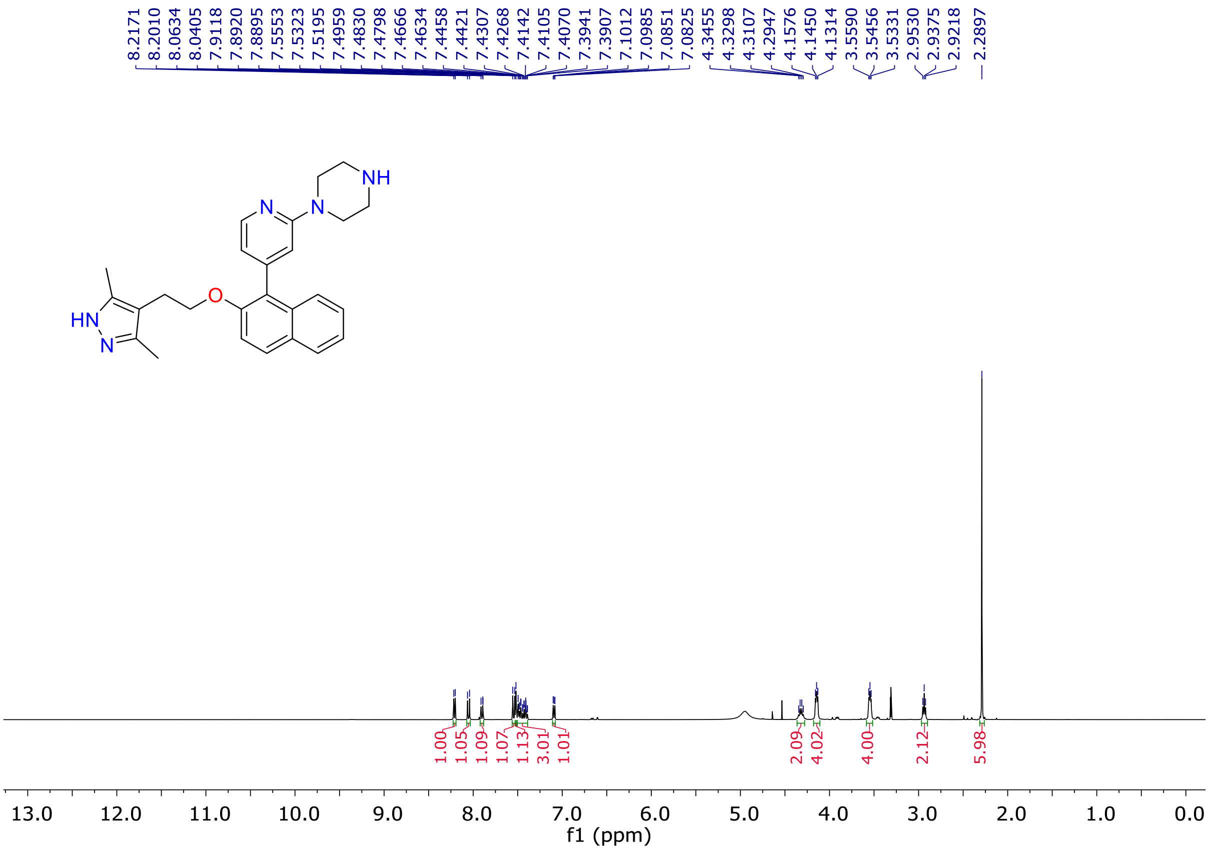


**Fig. S74.** $^{1}$H-NMR spectrum of compound **30e** ($CD_3OD$, 400 MHz).

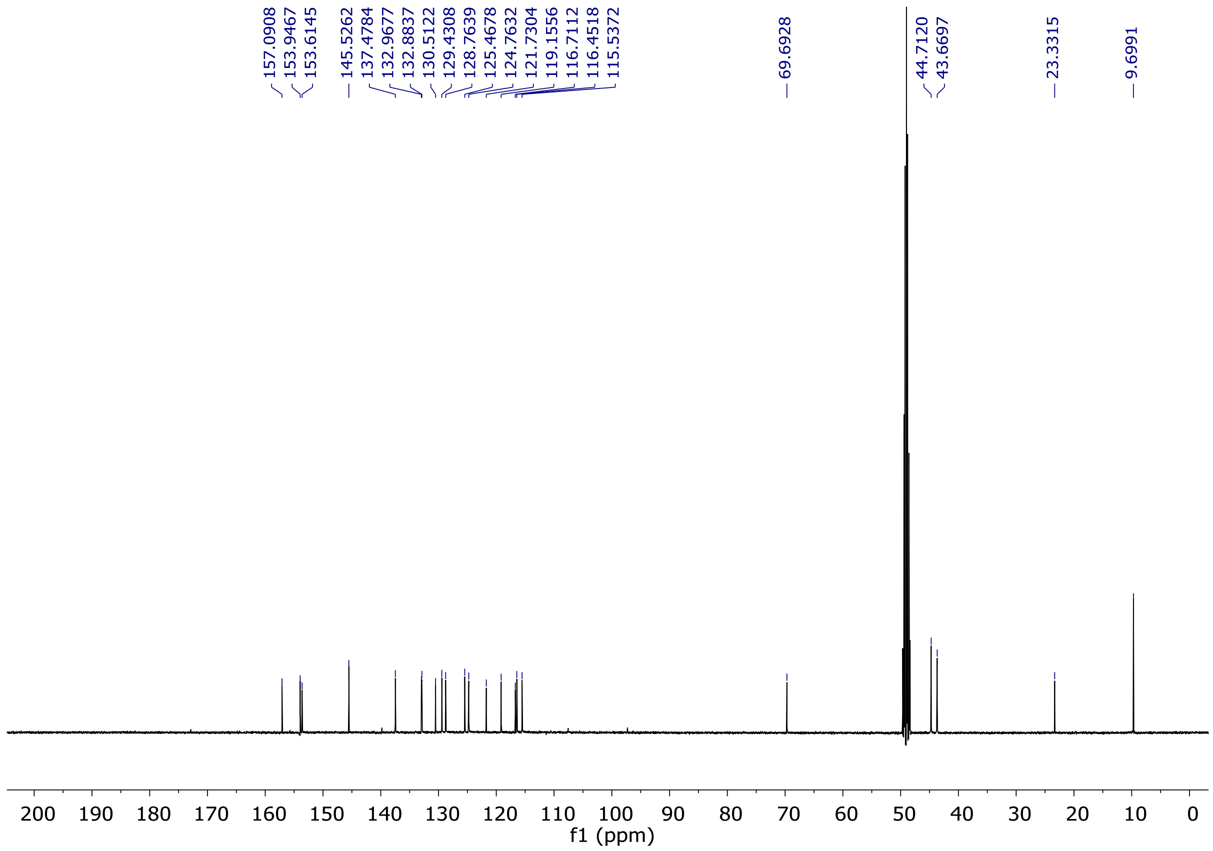

**Fig. S75.** $^{13}$C-NMR spectrum of compound **30e** ($CD_3OD$, 100 MHz).

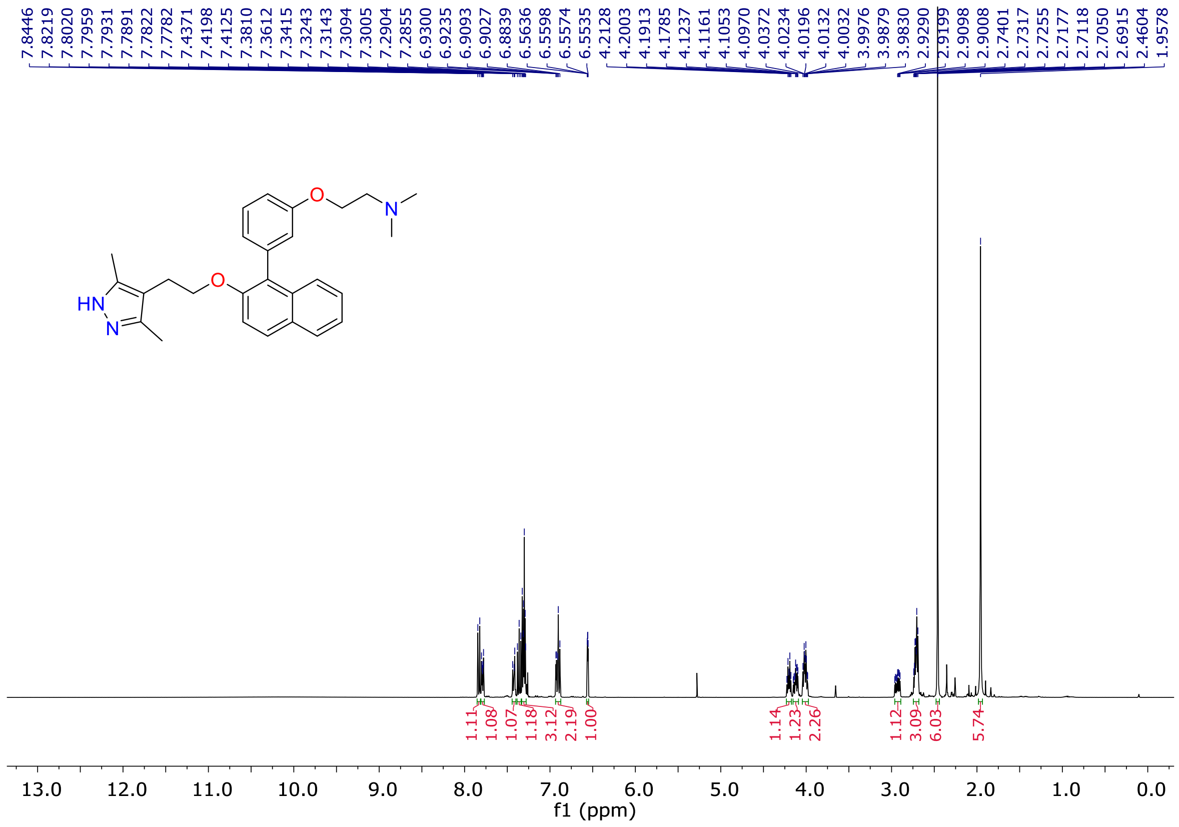

**Fig. S76.** $^{1}$H-NMR spectrum of compound **30f** ($CD_3OD$, 400 MHz).

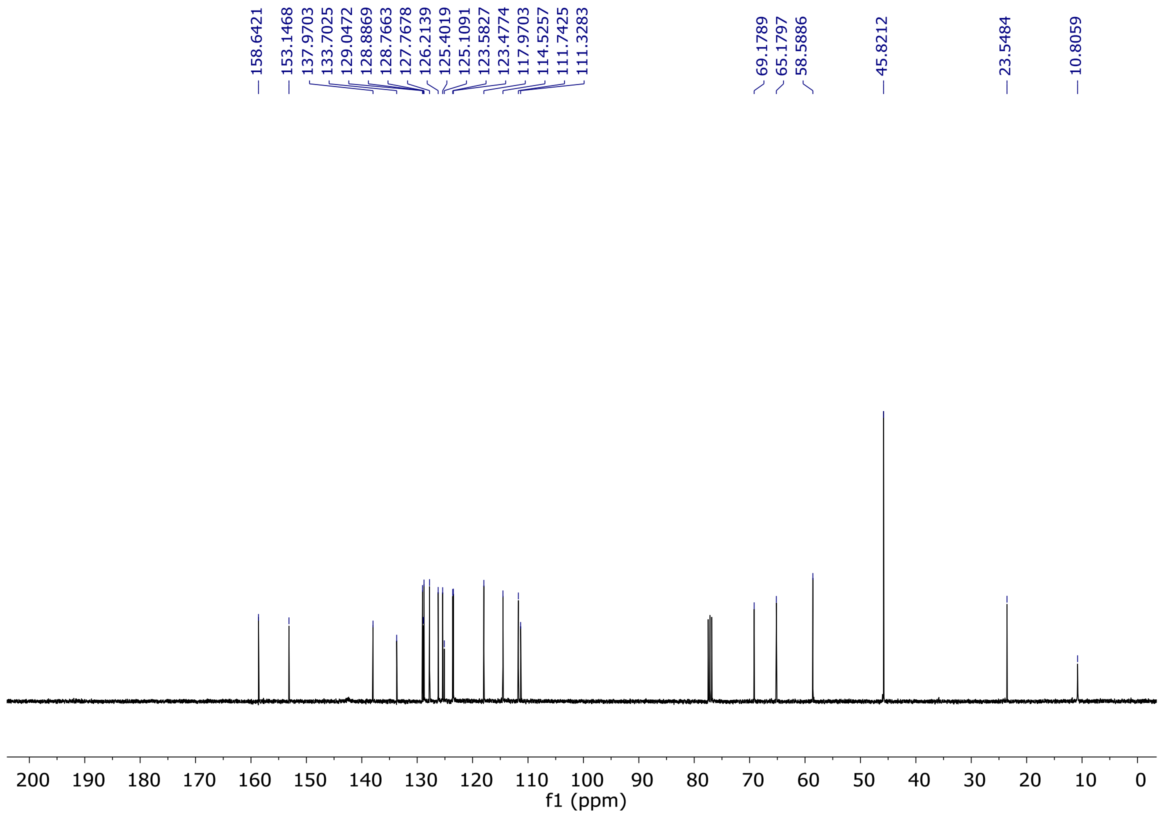


**Fig. S77.** $^{13}$C-NMR spectrum of compound **30f** ($CD_3OD$, 100 MHz).

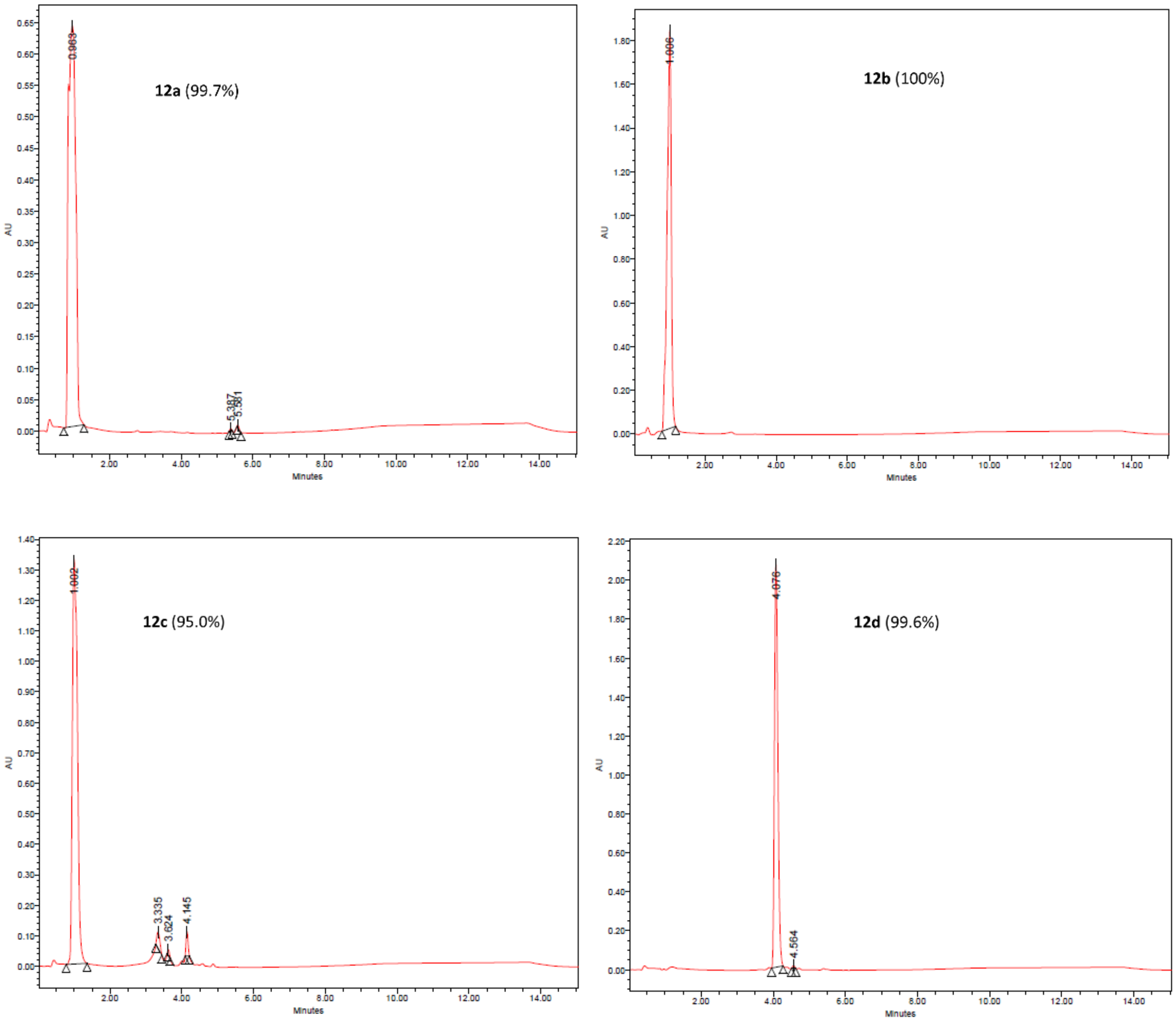

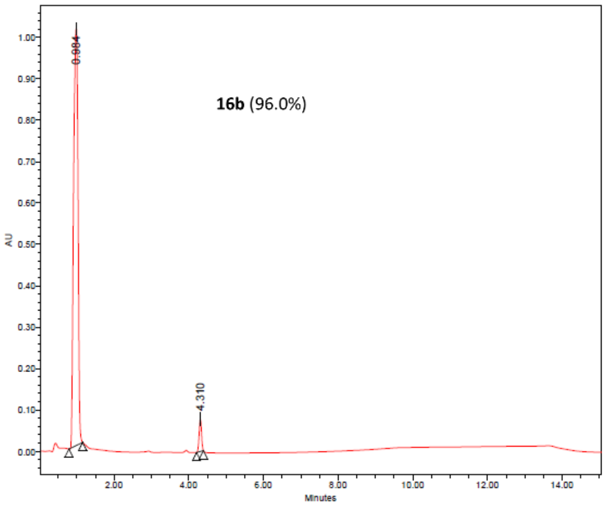
16b (96.0%)

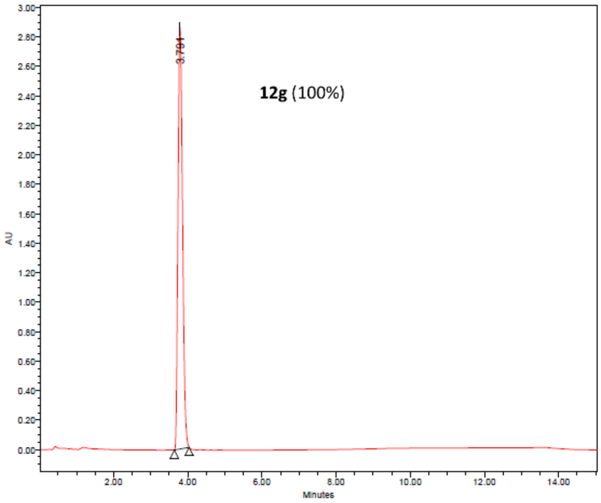
12g (100%)

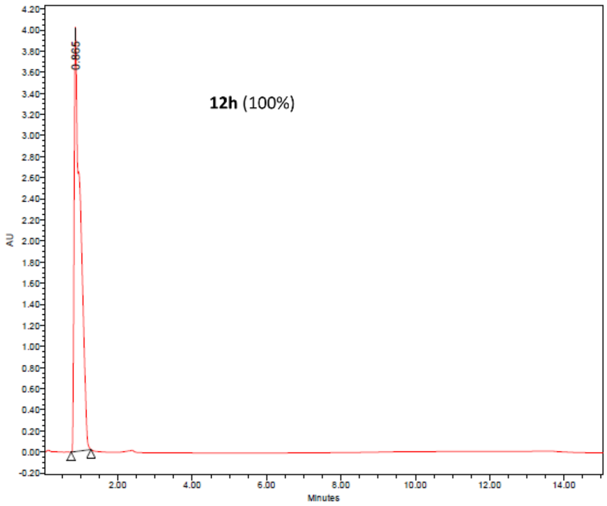
12h (100%)

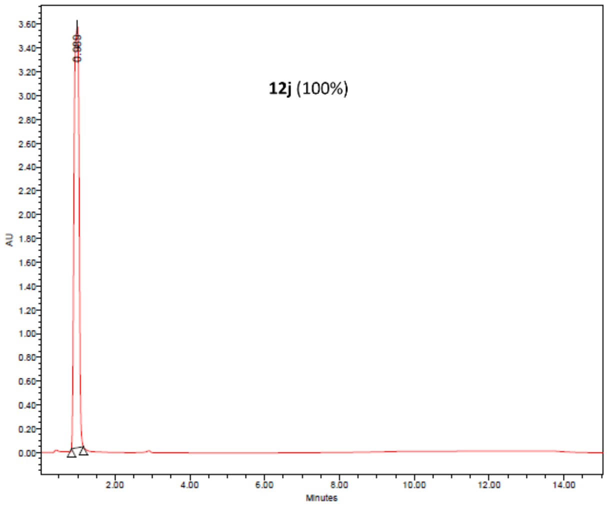
12j (100%)

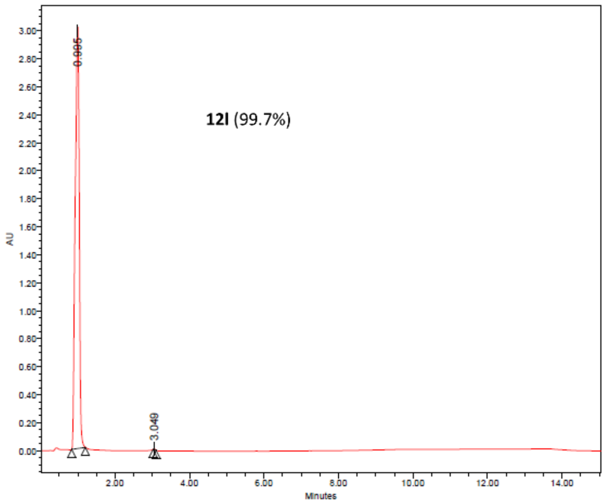
12l (99.7%)

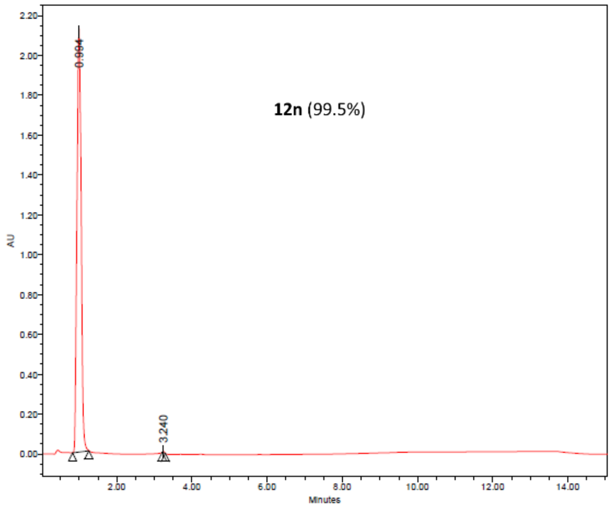
12n (99.5%)

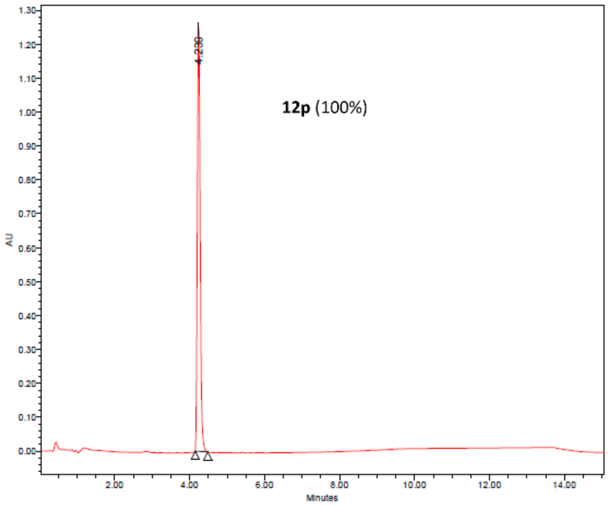
12p (100%)
4.230
AU
Minutes

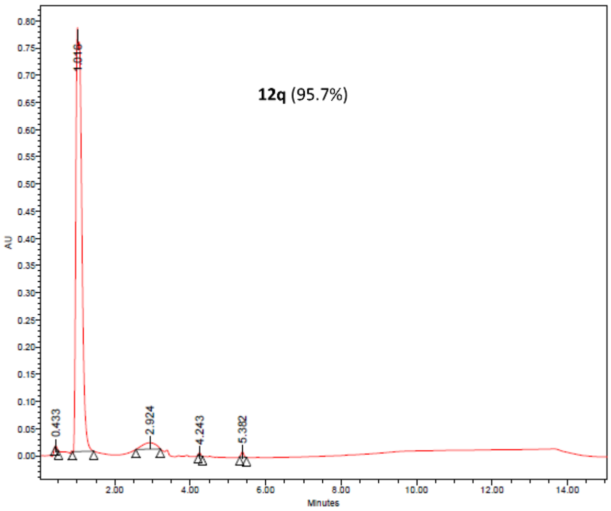
12q (95.7%)
1.016
0.433
2.924
4.243
5.382
AU
Minutes

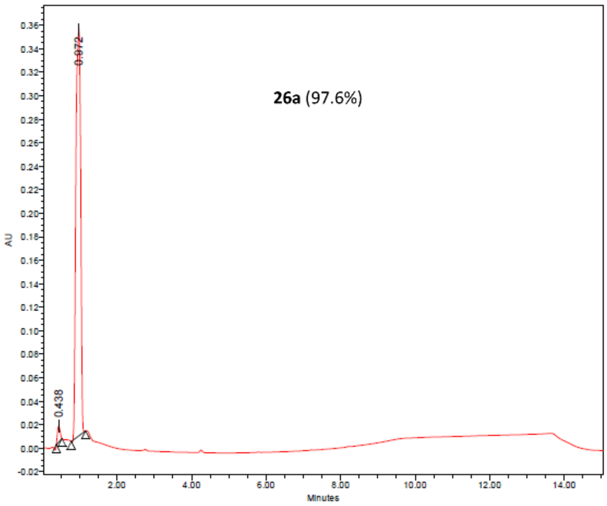
26a (97.6%)
0.972
0.438
AU
Minutes

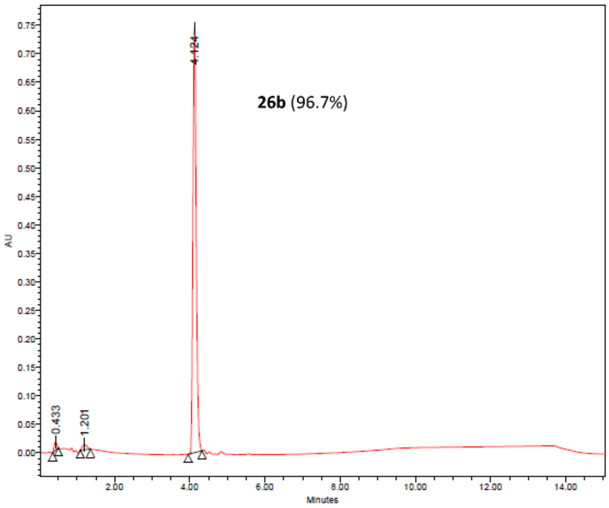
26b (96.7%)
4.124
0.433
1.201
AU
Minutes

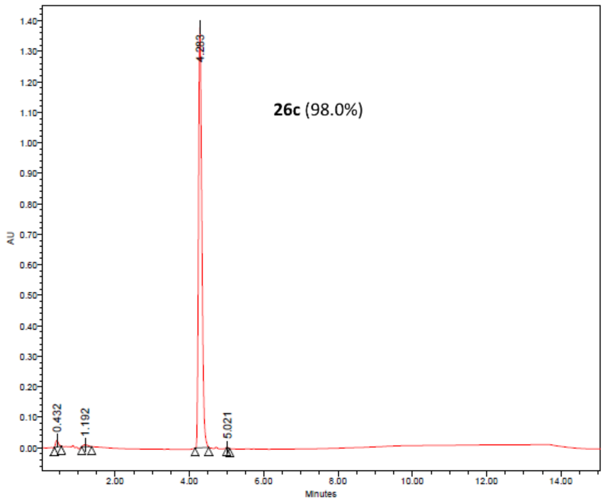
26c (98.0%)
4.283
0.432
1.192
5.021
AU
Minutes

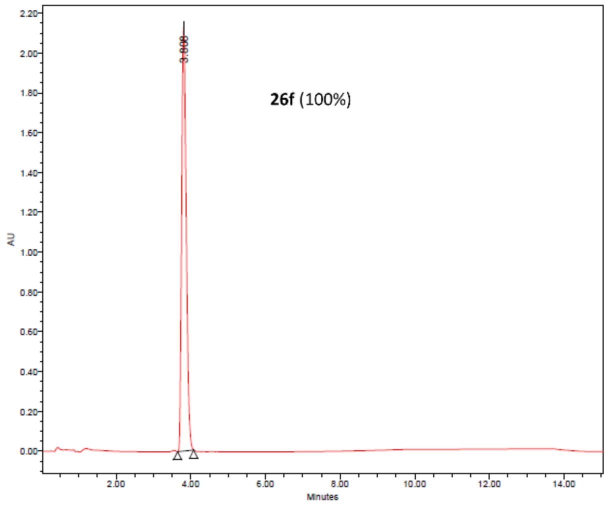
26f (100%)
3.808
AU
Minutes

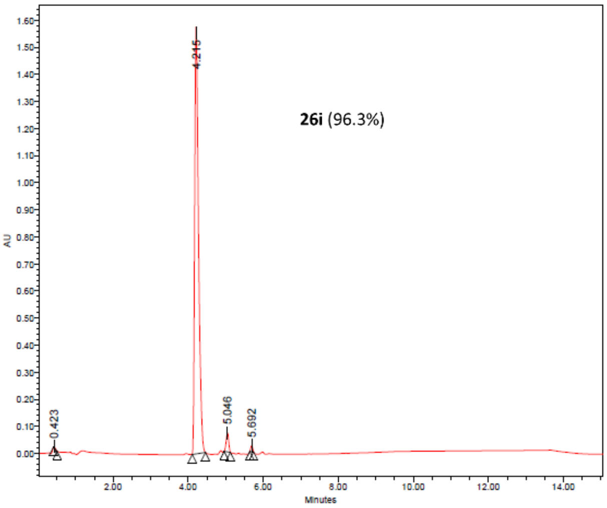


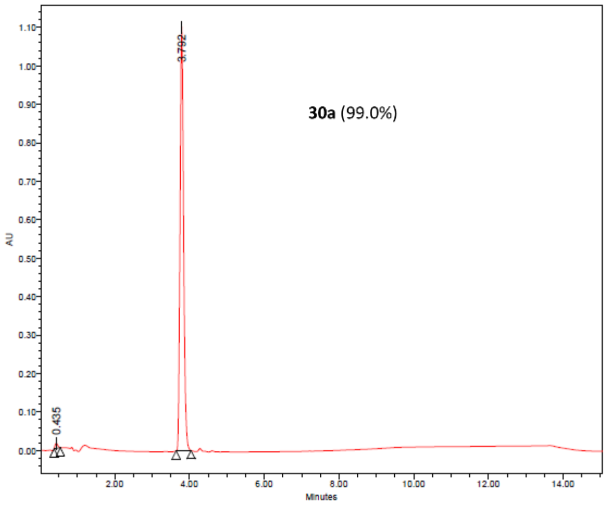


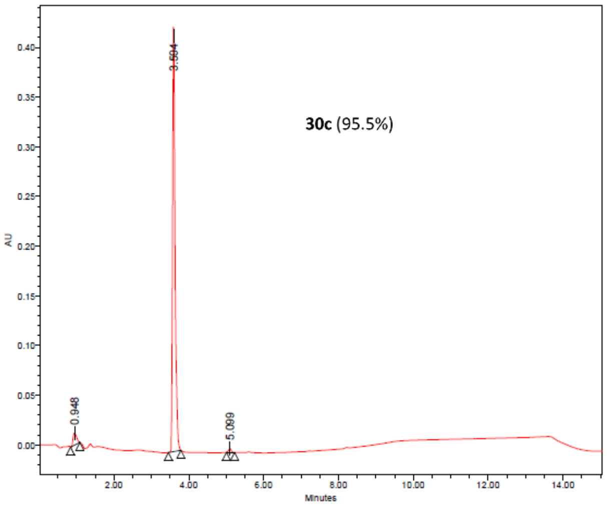


**Fig. S78.** HPLC Chromatogram of selected compounds.

### Supplementary Tables

**Table S1.** X-ray diffraction and structure refinement statistics.

| **PDB entry** | |
|---|---|
| SSGCID sample ID | PlviB.18219.a.FR2.GE44010 |
| X-ray source | ALS beamline 8.2.2 |
| Wavelength (Å) | 1.00000 |
| Resolution (Å) | 50.00 – 1.65 |
| Space group | $P4_32_12$ |
| **Unit cell** | |
| *a*, *c* (Å) | 91.930, 101.286 |
| No. reflections | 751,443 |
| Unique reflections | 52,868 |
| $R_{merge}$ (%) | 11.6 (88.9) |
| $R_{pim}$ (%) | 3.2 (24.6) |
| *CC1/2* | 0.998 (0.903) |
| Completeness (%) | 100.0 (100.0) |
| Redundancy | 14.2 (13.4) |
| $\langle I \rangle / \langle \sigma \rangle$ | 23.7 (2.2) |
| **Refinement** | |
| Resolution (Å) | 45.97 – 1.65 |
| No. reflections | 52,787 |
| Protein atoms | 6,928 |
| Ligand atoms | 402 |
| Water molecules | 435 |
| $R_{work}/R_{free}$ (%) | 13.61/16.48 |
| **RMS deviations** | |
| Bond lengths (Å) | 0.010 |
| Bond angles (°) | 1.085 |
| **Ramachandran plot** | |
| Favored (%) | 97.13 |
| Allowed (%) | 2.87 |
| Outlier (%) | 0.00 |
| **Molprobity (*39*)** | |
| All-atom clashscore | 1.91 |

*Values in parentheses correspond to the highest resolution shell

**Table S2**. Effect of NMTis on *P. vivax* liver stage infection

| compound | schizonts (fold change to DMSO) | | | | | | | hypnozoites (fold change to DMSO) | | | | | |
|---|---|---|---|---|---|---|---|---|---|---|---|---|---|
| | isolate 1 | | isolate 2 | | isolate 3 | | difference between isolates? | isolate 1 | | isolate 2 | | difference between isolates? | difference between forms? |
| | mean diff | p-value | mean diff | p-value | mean diff | p-value | | mean diff | p-value | mean diff | p-value | | |
| **12a** | 0.850 | 0.000 | 0.9245 | <0.0001 | 0.7273 | <0.0001 | no | 0.754 | 0.0015 | 0.897 | <0.0001 | no | no |
| **12b** | 0.950 | <0.0001 | 0.8868 | <0.0001 | 0.8977 | <0.0001 | no | 0.923 | <0.0001 | 0.966 | <0.0001 | no | no |
| **12d** | 0.967 | <0.0001 | 0.9434 | <0.0001 | 0.8977 | <0.0001 | no | 0.939 | <0.0001 | 0.862 | <0.0001 | no | no |
| **12e** | 0.983 | <0.0001 | 0.8302 | <0.0001 | 0.8295 | <0.0001 | no | 0.815 | 0.0005 | 1.000 | <0.0001 | no | no |
| **12f** | 1.000 | <0.0001 | 0.9811 | <0.0001 | 0.8295 | <0.0001 | no | 0.939 | <0.0001 | 0.931 | <0.0001 | no | no |
| **12g** | 1.000 | <0.0001 | 0.7547 | <0.0001 | 0.7955 | <0.0001 | no | 0.785 | 0.0009 | 0.966 | <0.0001 | no | no |
| **12j** | 0.983 | <0.0001 | 0.9434 | <0.0001 | 0.7614 | <0.0001 | no | 0.985 | <0.0001 | 0.897 | <0.0001 | no | no |
| **12l** | 0.983 | <0.0001 | 0.7736 | <0.0001 | 0.6932 | <0.0001 | no | 1.000 | <0.0001 | 0.828 | <0.0001 | no | no |
| **12q** | 0.983 | <0.0001 | 0.8491 | <0.0001 | 0.6591 | <0.0001 | yes | 0.939 | <0.0001 | 0.931 | <0.0001 | no | no |
| **26b** | 0.917 | <0.0001 | 0.8679 | <0.0001 | 0.625 | <0.0001 | no | 0.908 | <0.0001 | 0.793 | <0.0001 | no | no |
| **26c** | 1.000 | <0.0001 | 0.9434 | <0.0001 | 0.8977 | <0.0001 | no | 0.923 | <0.0001 | 0.862 | <0.0001 | no | no |
| **26i** | 0.933 | <0.0001 | 0.8491 | <0.0001 | 0.8295 | <0.0001 | no | 1.000 | <0.0001 | 0.310 | 0.0429 | yes | yes |
| **30a** | 1.000 | <0.0001 | 0.9245 | <0.0001 | 0.8636 | <0.0001 | no | 0.954 | <0.0001 | 1.000 | <0.0001 | no | no |
| **30c** | 0.900 | 0.000 | 0.7547 | <0.0001 | 0.8636 | <0.0001 | no | 0.939 | <0.0001 | 0.690 | <0.0001 | no | no |
